# Evaluation of altered cell-cell communication between glia and neurons in the hippocampus of 3xTg-AD mice at two time points

**DOI:** 10.1101/2024.05.21.595199

**Authors:** Tabea M. Soelter, Timothy C. Howton, Elizabeth J. Wilk, Jordan H. Whitlock, Amanda D. Clark, Allison Birnbaum, Dalton C. Patterson, Constanza J. Cortes, Brittany N. Lasseigne

**Author notes:** corresponding author Brittany N. Lasseigne. **Co-author email addresses:** Tabea M. Soelter, Timothy C. Howton, Elizabeth J. Wilk, Jordan H. Whitlock, Amanda D. Clark, Allison Birnbaum, Dalton C. Patterson, Constanza J. Cortes.

## Abstract

Alzheimer’s disease (AD) is the most common form of dementia and is characterized by progressive memory loss and cognitive decline, affecting behavior, speech, and motor abilities. The neuropathology of AD includes the formation of extracellular amyloid-β plaque and intracellular neurofibrillary tangles of phosphorylated tau, along with neuronal loss. While neuronal loss is an AD hallmark, cell-cell communication between neuronal and non-neuronal cell populations maintains neuronal health and brain homeostasis. To study changes in cellcell communication during disease progression, we performed snRNA-sequencing of the hippocampus from female 3xTg-AD and wild-type littermates at 6 and 12 months. We inferred differential cell-cell communication between 3xTg-AD and wild-type mice across time points and between senders (astrocytes, microglia, oligodendrocytes, and OPCs) and receivers (excitatory and inhibitory neurons) of interest. We also assessed the downstream effects of altered glia-neuron communication using pseudobulk differential gene expression, functional enrichment, and gene regulatory analyses. We found that glia-neuron communication is increasingly dysregulated in 12-month 3xTg-AD mice. We also identified 23 AD-associated ligand-receptor pairs that are upregulated in the 12-month-old 3xTg-AD hippocampus. Our results suggest increased AD association of interactions originating from microglia. Signaling mediators were not significantly differentially expressed but showed altered gene regulation and TF activity. Our findings indicate that altered glia-neuron communication is increasingly dysregulated and affects the gene regulatory mechanisms in neurons of 12-month-old 3xTg-AD mice.

## Introduction

Age is the primary known risk factor for Alzheimer’s disease (AD), with most patients being 65 or older.^1^ AD is the most prevalent neurodegenerative disease^2^ and the most common form of dementia.^3,4^ There are an estimated 55 million AD patients worldwide, and by the year 2050, AD prevalence is expected to increase to 139 million.^5^ AD is characterized by progressive memory loss and cognitive decline, affecting behavior, speech, and motor abilities.^6–8^ The neuropathology of AD includes the formation of extracellular amyloid-β plaques and intracellular neurofibrillary tangles of phosphorylated tau, as well as neuronal loss.^6^ Memory-associated brain regions like the entorhinal cortex and hippocampus are primarily affected in the early disease stages.^9,10^ During the later stages, the cerebral cortex, responsible for speech, social behavior, and reasoning, is also affected.^10^ AD is diagnosed through clinical evaluation of memory and motor skills and confirmation of tau and amyloid-β plaque formation using tau and amyloid PET imaging, respectively.^11–13^ In familial or early-onset AD, variants in the presenilin 1 (*PSEN1*), presenilin 2 (*PSEN2*), and amyloid precursor protein (*APP*) genes can cause AD.^14^ However, most patients have sporadic or late-onset AD, where the genetic causes are not fully understood.^15^ One approach to early AD diagnosis involves detecting nucleic acids and proteins like amyloid-β in cerebrospinal fluid and blood through liquid biopsies (reviewed in ^16–18^). However, the blood-brain barrier impacts detection sensitivity. Therefore, pre-clinical studies investigating the effects of amyloid-β and tau pathology on molecular mechanisms, such as cell-cell communication (CCC), in the brain are critical for establishing novel therapeutic targets.

While neuronal loss is an AD hallmark,^6^ CCC between neuronal and non-neuronal cell populations is crucial for maintaining neuronal health and brain homeostasis.^19–22^ CCC is mediated through the release of chemical intermediates, which are taken up by cell surface membrane receptors.^23,24^ Tripartite synapses, supported by microglia, facilitate bidirectional crosstalk between neurons and astrocytes, providing metabolic support and maintaining synaptic homeostasis of neurotransmitters.^21,25–27^ As the brain’s resident macrophages, microglia respond to injury by releasing chemical messengers, impacting neuronal activity, and modulating neurotransmission.^28–30^ In addition, oligodendrocytes facilitate myelin formation of neuronal axons,^31^ and oligodendrocyte precursor cells (OPCs) form synapses with neurons.^32^ Further, disruption of OPC-neuron communication during development has been shown to lead to social cognitive deficits.^33^ Using computational CCC inference tools, we and others have demonstrated that CCC is altered in the post-mortem human AD brain compared to healthy controls.^34–37^ These studies identified CCC dysregulation between all brain cell types in AD and underscored the significance of altered communication between neurons and glia, as neuron-astrocyte and neuron-microglia interactions had increased involvement of AD-risk genes. We previously presented how altered glia-neuron communication in the human prefrontal cortex might affect canonical signaling pathways, including WNT, NFkB, and p53 in inhibitory neurons.^36^ Although human studies are critical, their reliance on late-stage post-mortem brain tissue does not address how CCC is dysregulated throughout aging and AD progression. While cell membrane receptors have emerged as potentially druggable targets for disease, the effects of AD neuropathology on CCC between glia and neurons throughout disease progression remains understudied. Thus, using a mouse model that mimics AD pathology across multiple time points may further define changes in CCC throughout disease progression that cannot be investigated using postmortem human tissues.

To study how CCC changes during AD progression with respect to amyloid-β and tau pathology, we generated single-nucleus RNA sequencing (snRNA-seq) data from 3xTg-AD mice across two time points. The transgenic mouse model we studied, 3xTg-AD, harbors three human transgenes (*APP*, *PSEN1*, and *MAPT*) and exhibits both amyloid-β and tau pathology shown to occur as early as 6 and 12 months, respectively.^38^ It is a well-characterized and extensively phenotyped model of AD pathogenesis,^39^ and has been successfully utilized as a pre-clinical and drug discovery platform in the past.^40–42^ Importantly, sexually dimorphic characteristics are becoming a well-recognized feature of this transgenic model, although their prevalence and impact on AD-associated pathological hallmarks such as inflammation, plaque, and/or tangle accumulation and others remain poorly understood (reviewed in ^43^). Therefore, we sequenced the hippocampus from female 3xTg-AD and wild-type (WT) littermates at 6 and 12 months (**Fig 1A**) to evaluate altered glia-neuron communication. Given how critical CCC between glia and neurons is for brain health and AD pathology, we inferred differential CCC between 3xTg-AD and WT mice across time points and between senders (astrocytes, microglia, oligodendrocytes, and OPCs) and receivers (excitatory and inhibitory neurons) of interest. We predicted differentially expressed ligand-receptor pairs and their potential downstream target genes (**Fig 1B**). We also assessed the global downstream effects of altered glia-neuron communication using pseudobulk differential gene expression and functional enrichment analyses (**Fig 1C-D**). To determine AD-associated interactions, we compiled an AD risk gene set from the Molecular Signatures Database (MSigDB)^44,45^ and a recent Genome-wide association study (GWAS).^46^ Using gene regulatory information, we also predicted signaling mediators of AD-associated ligand-receptor-target (LRT) pairings and characterized their expression, differential gene targeting, and transcription factor (TF) activity (**Fig 1E-F**).

**Figure 1.**
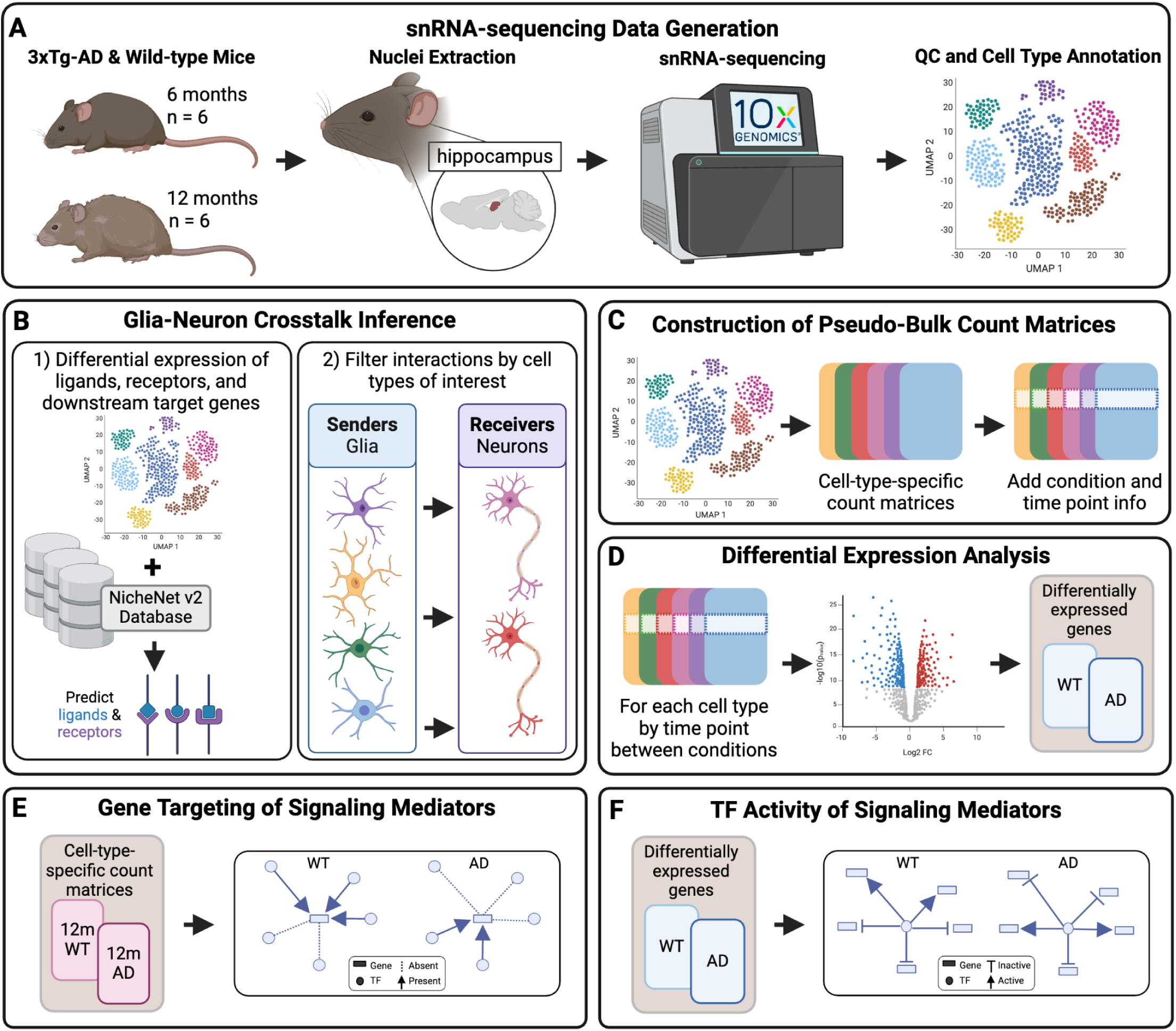
Schematic overview of our study design. (**A**) We sequenced the hippocampus of female 3xTg-AD and WT littermates at 6 and 12 months. (**B**) We inferred ligands and receptors between senders (astrocytes, microglia, oligodendrocytes, OPCs) and receivers (excitatory and inhibitory neurons) using differential gene expression (log2FC > 0.5) and known ligand-receptor information. We also predicted target genes potentially regulated by the predicted ligand-receptor pairs. (**C**) We generated cell-type-specific count matrices with condition and time point information for downstream analyses. (**D**) Using the cell-type-specific count matrices, we identified differentially expressed genes between 3xTg-AD and WT hippocampus for every cell type and at each time point (absolute log2FC > 0.2, Wald test padj < 0.05). (**E**) To determine changes in gene regulation of signaling mediators, we built cell-type- and time-point-specific gene regulatory networks (GRNs) and calculated the differential gene targeting score of signaling mediators between 12-month 3xTg-AD and WT GRNs. (**F**) Using differential expression information, we inferred the differential TF activity of signaling mediators that are TFs between conditions at 12 months.

## Materials & Methods

A more detailed version of the methods section is available in the **Supplementary Methods.**

### Animals

We bred and housed 3xTg-AD mice (B6;129-Tg(APPSwe,tauP301L)1Lfa *Psen1*^tm1Mpm^/Mmjax: JAX MMRRC Stock# 034830) and WT littermates.^38^ We anesthetized mice using Avertin intraperitoneal injections before perfusion with 1X PBS. We snap-froze and stored hippocampi at −70C until nuclei isolation.

### Nuclei isolation and snRNA sequencing of 3xTg-AD mouse hippocampus

We isolated nuclei from the left hippocampus of 12 female mice (n = 6/condition) from two time points (6 and 12 months; n = 3/time point and condition) as previously described.^47^ We prepared sequencing libraries using the Chromium Single Cell 3ʹ GEM, Library & Gel Bead Kit v3 (10x Stock #: PN-1000121). We sequenced libraries on an Illumina NovaSeq 6000 using an S4 flowcell to an average depth of ∼34,000 reads per nucleus with ∼15,000 nuclei per sample (**Table S1**).

### Protein quantification of amyloid-β 40, amyloid-β 42, and total tau

We used the human Amyloid-β 40 (ThermoFisher Cat No. KHB3481), Amyloid-β 42 (ThermoFisher Cat No. KHB3441), and Total Tau (ThermoFisher Cat No. KHB0041) ELISAs to quantify protein abundance in the hippocampus for both conditions and time points. We performed a Bicinchoninic acid assay (BCA) to quantify the total protein (**Table S2**) before performing ELISAs according to the manufacturer’s instructions and measured absorbance (**Table S2**). We used a 4-parameter algorithm to quantify protein concentration and compared groups using a one-way ANOVA followed by a Tukey test for multiple hypothesis correction.

### Data Processing and Quality Control

We aligned raw FASTQ files to GRCm38/mm10 using Cell Ranger^48^ v6.1.1. We performed all downstream data processing in docker^49^ R v4.2.3. We removed ambient RNA using soupX^50^ v1.6.2. We created a merged Seurat object using Seurat v4.3.0^51^. We filtered each dataset at the cell and at the gene level. We performed batch correction using harmony v0.1.0.^52^ We also performed Principal Component Analysis (PCA) without approximation (approx = FALSE), scaled, and normalized before plotting UMAPs to confirm successful integration across conditions.

### Clustering and Cell Type Identification

We compared multiple resolutions using clustree v0.5.0^53^ and chose 0.7 resolution, yielding 40 clusters using Leiden v0.4.3^54^. We identified differentially expressed marker genes for each cluster using the *FindAllMarkers* function from Seurat v4.3.0^51^ with a log2foldchange threshold > 0.2. We assigned cell types using differential expression of cell-type-specific genes identified through PanglaoDB^55^ and CellMarker 2.0^56^ (**Table S3**) and feature plots using canonical cell type markers (**Table S4**).

### Pseudobulking of snRNA-seq Data

We converted raw data from the Seurat object *counts* slot into Single Cell Experiment objects using SingleCellExperiment^57^ v1.20.1 and incorporated necessary metadata information. We aggregated counts across samples by cell type, resulting in cell-type-specific gene-by-sample count matrices.

### Differential Gene Expression Analysis

We performed differential expression analysis using DESeq2^58^ v1.38.3. We made pairwise comparisons for every cell type between conditions for each time point independently using the Wald test. Then we performed log2foldchange shrinkage using apeglm v1.23.1.^59^

### Inference of CCC Across Conditions and Time Points

To infer glia-neuron CCC between AD and WT across time points, we applied multinichenetr v1.0.3 and used the Nichenet v2 prior.^60^ We performed ligand-receptor pair prediction between all cell types for every time point across conditions before filtering for senders and receivers of interest. We identified differentially expressed ligands, receptors, and target genes using a log2FC threshold = 0.5 and non-adjusted p-values threshold = 0.05. When calculating ligand activity, we considered the top 250 targets with the highest regulatory potential. We prioritized ligand-receptor interactions using default MultiNicheNet parameters.

### Functional Enrichment Analysis of Predicted Target Genes

To determine the molecular function of predicted target genes from MultiNicheNet, we identified significantly enriched pathways using gprofiler2^61^ v0.2.1. We queried predicted target genes for every group (6mAD, 6mWT, 12mAD, 12mWT) to find overrepresented pathways. We included all genes measured as background genes. We filtered pathways by whether they were significant after multiple hypothesis correction using Bonferroni (q < 0.05) as well as the term size (> 10 and < 1000).

### AD risk gene set curation

We compiled an AD risk gene set by combining two human gene sets from the Molecular Signatures Database (MSigDB)^44,45^ with genes from a GWAS.^46^ The AD risk gene set comprised 245 mouse genes (**Table S5**).

### Prediction of LRT signaling mediators

To find AD-associated interactions, we used our AD risk gene set to filter LRTs targeting an AD risk gene. We used gene regulatory information from the NicheNet v2 database to generate weighted and directed igraph objects for AD-associated LRTs using igraph^62^ v1.4.2. We determined potential signaling mediators by identifying outgoing nodes from the predicted receptor in every igraph object.

### GRN Construction

To investigate regulatory relationships of signaling mediators, we constructed cell-type- and time-point-specific networks using Passing Attributes between Networks for Data Assimilation (PANDA)^63,64^ v1.30.0. This resulted in 4 GRNs (12mWT & 12mAD for excitatory neurons, 12mWT & 12mAD for inhibitory neurons). Network edge weights represent the strength of interactions between nodes, where nodes represent coexpressed genes, proteins, or pairs of TFs and genes. More positive weights correspond to higher confidence interactions, and negative weights indicate a lack of evidence.^64^

### TF Activity Analysis of Signaling Mediators

To determine differences in TF activity between conditions for receiver cell types, we performed a pseudobulk differential gene expression analysis with DESeq2. However, to preserve the Wald test statistic needed for our TF activity analyses, we used the method’s original shrinkage estimator (type = normal). We combined the Wald test statistic across time points for both receivers. We used the CollecTRI prior and calculated TF activity scores for all signaling mediators in receivers with a minimum of 5 targets using the Multivariate Linear Model (decoupleR v2.9.1).^65^

### Differential Gene Targeting of Signaling Mediators

We calculated differential gene targeting scores^66^ to determine whether signaling mediators in receivers were differentially regulated in the 3xTg-AD hippocampus using the previously generated PANDA GRNs. We quantified the in-degree edge weights for each mediator before determining the differential targeting score between conditions. We calculated quartiles of all gene targeting scores to prioritize the most differentially targeted mediators.^67^

## Results

### snRNA-seq data of 3xTgAD and WT mice across conditions and time points

To study CCC throughout AD progression, and given the higher neuropathological burden and extensive characterization of female individuals in the 3xTgAD model ^43^, we generated snRNA-seq data from female 3xTgAD and WT littermates across two time points: one considered early in the pathological progression in this model (6 months of age) and mid-way through (12 months of age).^39^ Using Leiden clustering, we identified 40 individual clusters, which we combined into 12 brain and vasculature cell types (**Fig 2A**) using canonical marker gene expression (**Fig 2B**). Next, we confirmed that all cell types integrated and were evenly distributed across conditions and time points (**Fig 2C-F**). However, we observed greater variation in our cell type proportions between time points (range of proportions: 0.35 - 0.65) than conditions (range of proportions: 0.45 - 0.55). Interestingly, microglia, the brain’s resident macrophages, had the greatest imbalance between time points, with almost 70% of microglia originating from 12-month 3xTg-AD and WT samples (**Fig 2F**). Increased inflammation due to microglia is well-known in AD (reviewed in ^68–70^) and the aging brain (reviewed in ^71^). Additionally, we confirmed the presence of amyloid-β and tau pathology in our mice by quantifying human amyloid-β 40, amyloid-β 42, and total tau in the hippocampus at both time points. We observed significant increases of all three proteins in 3xTg-AD compared to WT hippocampus at both time points (adjusted p-value < 0.01, one-way ANOVA followed by Tukey’s test). While we observed an increase in amyloid-β 40 and amyloid-β 42 abundance between 6 and 12 months in the 3xTg-AD hippocampus, this difference was not statistically significant (adjusted p-value > 0.05, one-way ANOVA followed by Tukey’s test; **Fig S1)**.

**Figure 2.**
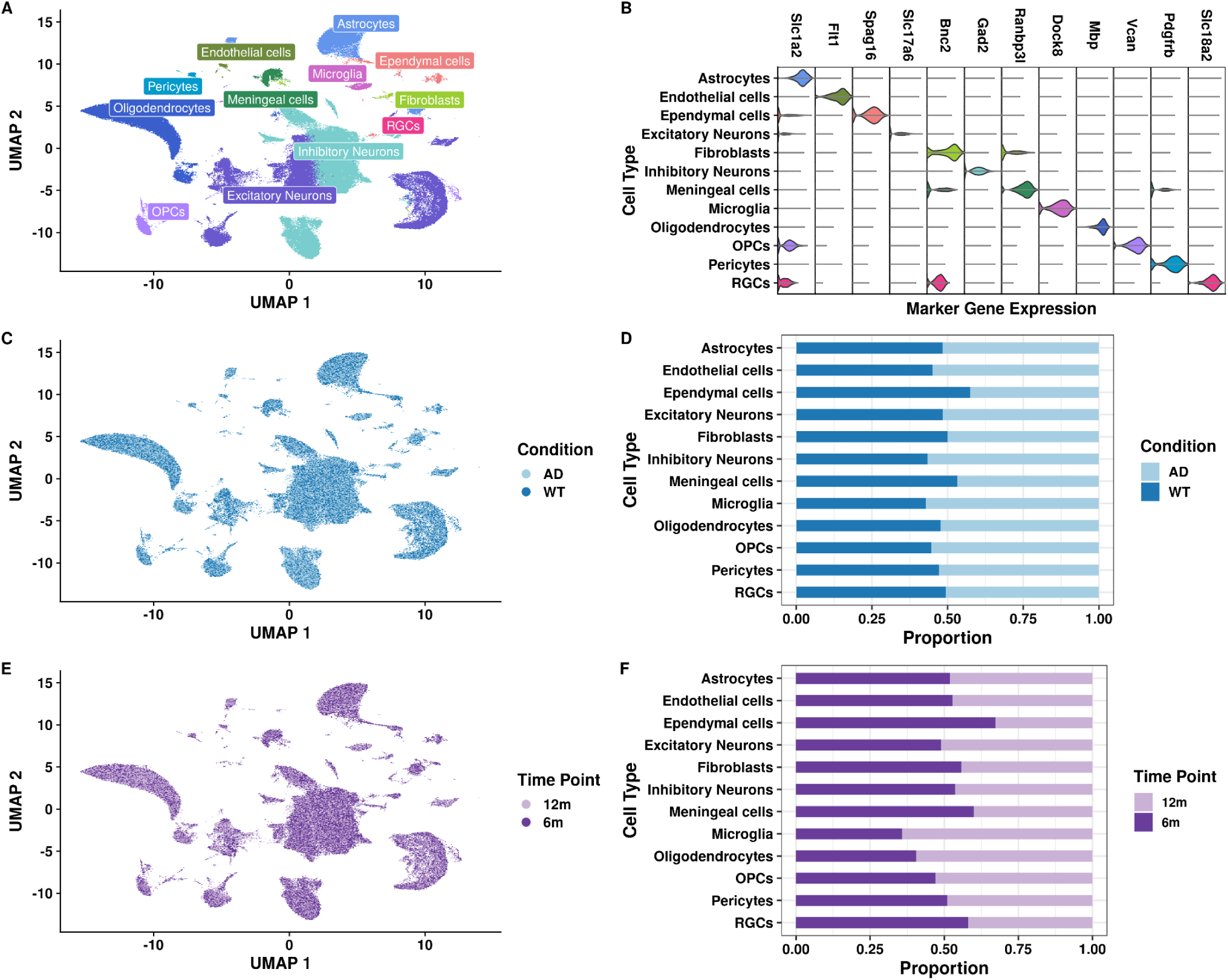
Overview of our 3xTgAD snRNA-seq data from the hippocampus across time points. **(A)** Uniform Manifold Approximation and Projection (UMAP) of 184,858 nuclei after quality control colored by cell type. **(B)** Stacked violin plot of the expression of representative marker genes used for cell type annotation. **(C)** Integrated UMAP split by condition (AD, WT). **(D)** Stacked barplot of the proportion of cell types annotated by condition. **(E)** UMAP split by time point (6 and 12 months) in both conditions. **(F)** Stacked barplot of cell type proportions by time point in both conditions.

### CCC is more dysregulated at 12 months across all cell types of interest

Using MultiNicheNet,^60^ we identified differentially expressed ligands and receptors between conditions for each time point across all cell types in our snRNA-seq dataset. We also predicted potential downstream target genes of inferred ligand-receptor pairs. In total, we identified 83,928 interactions (**Fig S2A**). Since glial cells are important for maintaining neuronal health and homeostasis and neuronal loss is an AD hallmark,^19–22^ we filtered interactions by senders (astrocytes, oligodendrocytes, OPCs, and microglia) and receivers (excitatory and inhibitory neurons) of interest and prioritized for higher signaling strength through ligand activity and regulatory potential of the downstream target genes. We retained 4,073 LRT pairings (log2FC > 0.5, Fisher’s exact test p-value < 0.05; **Fig S2B**). Most of the prioritized LRTs originated from astrocytes (n = 1,188), followed by OPCs (n = 1,147), microglia (n = 1,113), and oligodendrocytes (n = 625). Interestingly, 86% of predicted LRTs were dysregulated at the 12-month time point, with most interactions being upregulated in the 12-month 3xTg-AD hippocampus (**Fig 3A & 3E**). This indicates that while CCC is affected at 6 months in the 3xTg-AD mouse hippocampus, the disease progression and increased severity and duration of the AD pathology at 12 months exasperated CCC dysregulation in this AD mouse model. Most interactions originating from astrocytes, microglia, and OPCs were upregulated in 12-month 3xTg-AD hippocampus compared to WT (**Fig 3A**), potentially explained by reactive astrocyte activation through microglial secreted cytokines in AD.^72^ Most of the interactions originating from oligodendrocytes were downregulated in 12-month 3xTg-AD compared to WT (**Fig 3A**). Since CCC is predicted using differential gene expression, interactions upregulated in WT mice have decreased expression in the 3xTg-AD hippocampus; therefore, we inferred those interactions as downregulated or lost in AD. Accordingly, CCC originating from oligodendrocytes is primarily downregulated or lost in 3xTg-AD compared to WT hippocampus at 12 months (**Fig 3A**), indicating decreased signaling between oligondedrocytes and neurons, possibly contributing to neuronal demyelination, a pathophysiological feature in AD^73^ and throughout aging.^74^ Overall, we observe an age-dependent increase in CCC dysregulation for interactions originating from all senders in both conditions (**Fig 3A**). Since we predicted fewer interactions in 6-month-old mice (14% of all interactions; **Fig 3A-B & 3D**), despite their previously described amyloid-β pathology at this time point,^38^ we surmise that CCC in the 3xTg-AD hippocampus is likely influenced by the duration of amyloid-β and tau pathology and its long-term effects. Additionally, we observed 2,964 interactions targeting inhibitory neurons and 1,109 targeting excitatory neurons (**Fig S2C**). This bias towards inhibitory neurons was independent of the sender (**Fig S2C**), time point (**Fig 3B**), and the number of nuclei per receiver, as our dataset had 55,567 inhibitory neurons and 56,745 excitatory neurons. Interestingly, previous studies have indicated that inhibitory neurons are unaffected by amyloid-β pathology, and others revealed that disruption of inhibitory neuronal function in AD leads to memory deficits (reviewed in ^75^). To determine whether interactions overlapped between time points and conditions, we analyzed time point and condition pairings (groups: 6mAD, 6mWT, 12mAD, 12mWT). While the majority of interactions were group-specific, 132 LRTs overlapped between 6m and 12m AD samples, and 29 LRTs overlapped between 6m and 12m WT (**Fig 3B**), suggesting that some interactions are dysregulated during early amyloid-β pathology and remain dysregulated throughout the disease progression. Additionally, the 11 LRTs that overlapped between 6mAD and 12mWT, which included 7 ligand-receptor pairs with 6 target genes, might be markers of early senescence due to age and/or disease (**Table S6**). When disbanding shared LRTs by senders, OPCs had the highest number of LRTs shared between groups, and oligodendrocytes had the fewest (**Fig 3C**). Once we accounted for shared interactions, we observed that 30 of 39 interactions originating from microglia were downregulated or lost in 3xTg-AD mice at the 6-month time point, and interactions originating from other senders were upregulated in 3xTg-AD hippocampus (**Fig 3D**). Given that reactive microglia colocalize with amyloid-β plaques^76,77^ and neurofibrillary tau tangles^78^, these shared or age-associated interactions should be further investigated for their utility as early disease detection markers, as microglia’s ability to respond to injury becomes impaired throughout aging.^79,80^ Additionally, out of all downregulated LRTs at 12 months, 62.7% originated from oligodendrocytes (**Fig 3E**). Although microglia had the most downregulated interactions at 6 months (**Fig 3D**), they had the fewest at 12 months (**Fig 3E**). Though we identified the fewest interactions overall in oligodendrocytes (**Fig S2**), at the 6-month time point, all senders had similar numbers of predicted interactions (**Fig 3D**). In contrast, oligodendrocytes were the least dysregulated sender at 12 months (**Fig 3A & 3E**). Overall, our CCC analysis revealed that signaling between glia and neurons is increasingly dysregulated at 12 months in the 3xTg-AD mouse hippocampus, which coincides with both amyloid-β and tau pathology.^38^

**Figure 3.**
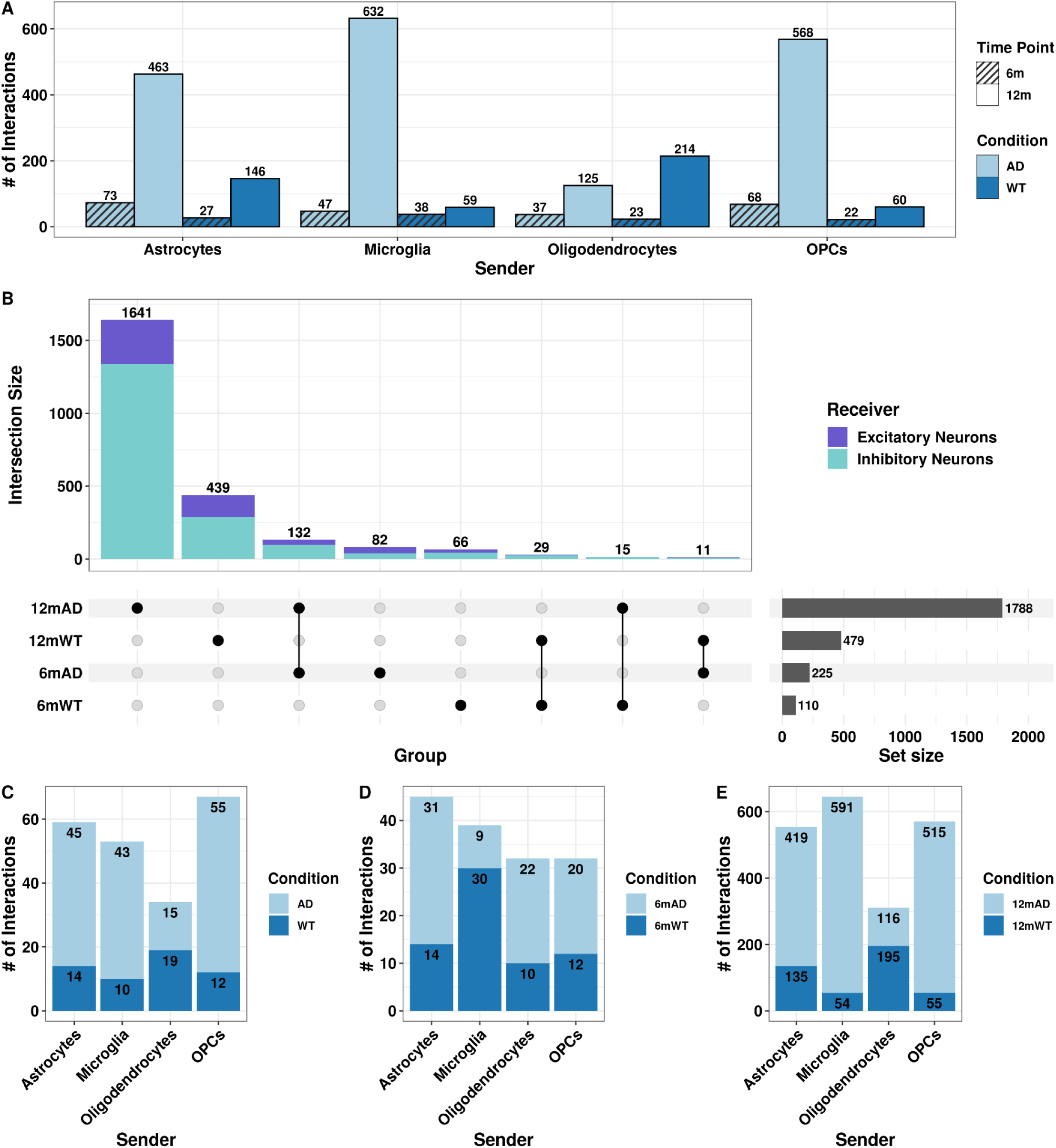
CCC dysregulation is increased at 12 months in the 3xTg-AD hippocampus. **(A)** Bar plot of the number of interactions up-regulated (log2FC > 0.5) in each time point and condition grouped by senders. Color and pattern indicate condition and time point, respectively. **(B)** UpSet plot of the overlap of interactions between individual groups (group = time point and condition pairs). The top bar plot denotes intersection size, circles represent which comparisons have overlap, and the set size reflects the total number of interactions for individuals groups. The intersection size bar plot is colored by the receiver. **(C)** Stacked bar plot of the interactions shared between time points and conditions colored by condition (3xTg-AD, WT). **(D)** Stacked bar plot of the interactions specific to the 6-month time point colored by group (6mAD, 6mWT). **(E)** Stacked bar plot of the interactions specific to the 12-month time point colored by group (12mAD, 12mWT).

### Target genes had greater differential expression and are associated with signaling and regulatory pathways in the 12-month-old 3xTg-AD hippocampus

To investigate the downstream effects of altered glia-neuron interactions in 3xTg-AD mouse hippocampus across time points, we performed pseudobulk differential gene expression analyses of predicted target genes in neurons between AD and WT at both time points (6mAD vs. 6mWT and 12mAD vs. 12mWT; **Table S7**).^58^ Since we predicted fewer altered interactions between 3xTg-AD and WT at 6 months (**Fig 3**), the number of predicted target genes was overall lower at this time point (**Fig 4A & 4B**), although the number of nuclei in our neuronal subpopulations was similar across time points (**Fig 2F**), further suggesting that CCC becomes increasingly dysregulated with age and/or due to prolonged exposure to amyloid-β and tau. Additionally, 6-month targets had smaller log2 gene expression fold changes (absolute log2FC = 2, **Fig 4A**) than 12-month targets between AD and WT (absolute log2FC = 4; **4B**). We found that caspase recruitment domain family member 10 (*Card10*) was the most downregulated gene in 3xTg-AD inhibitory (log2FC = −1.9) and excitatory neurons (log2FC = −1.25) at 6 months (**Fig 4A**). While *Card10* was also downregulated at 12 months in inhibitory and excitatory neurons, it was not the most downregulated gene at this later time point (log2FC = −2 and −1.75, respectively; **Fig 4B**). *Card10* is involved in activating the NF-kappa-B (NFkB) signaling pathway, known to have increased activity in AD due to its role in the inflammatory response and amyloid-β plaque formation.^81^ However, previous work in the blood of a transgenic AD mouse model indicated increased expression of *Card10* compared to WT mice.^82^ Protein tyrosine phosphatase N22 (*Ptpn22)*, an immune signaling regulator previously implicated in AD,^83,84^ was the only other target gene that was significantly differentially expressed in both neuronal subpopulations at both time points (absolute log2FC > 0.2 with a Wald test padj < 0.05, **Fig 4A & 4B**). While not all predicted target genes were significantly differentially expressed in excitatory and inhibitory neurons, many target genes showed changes in log2FC magnitude throughout aging (i.e., between 6 & 12 months; **Table S7**). Then, we performed a functional enrichment analysis (FEA) to infer the time point-specific function of predicted target genes in excitatory and inhibitory neurons. Unsurprisingly, the FEA terms for both 6- and 12-month targets were widely related to signaling (**Fig 4C & 4D**). Interestingly, 6-month targets also had decreased receptor ligand activity, signaling receptor activator activity, and receptor signaling activity in the 3xTg-AD hippocampus (**Fig 4C**). Growth factor binding was upregulated at 6 months in hippocampal neurons of 3xTg-AD mice compared to WT, and it was downregulated in neurons at 12 months in the hippocampus of 3xTg-AD mice (**Fig 4C & 4D**). Other 12-month 3xTg-AD FEA terms were associated with gene regulatory mechanisms through kinase activity (**Fig 4D**), indicating an age-associated increase of gene regulatory disruption which contributes to age-related disorders like AD.^85^ Overall, we found smaller log2 gene expression fold changes of predicted 3xTg-AD targets in 6 compared to 12 months, and FEA terms associated with these targets recapitulated signaling pathways at both time points, but gene regulatory disruption was specific to 12 months in the hippocampus of 3xTg-AD mice.

**Figure 4.**
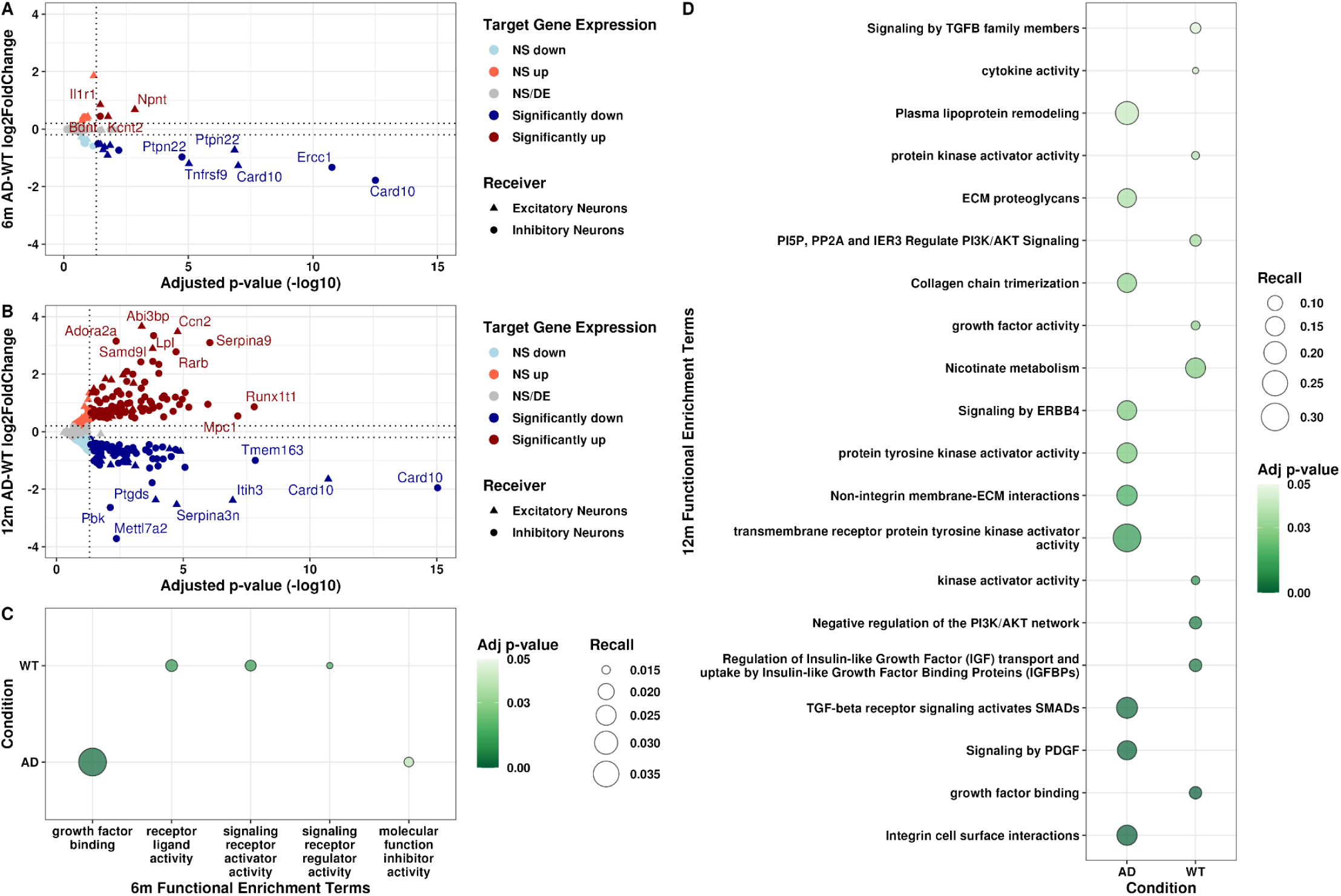
Increased target gene expression differences in the 3xTg-AD hippocampus at 12 months. Volcano plot of differential target gene expression in excitatory and inhibitory neurons between 3xTg-AD and WT mouse hippocampus at **(A)** 6 months and **(B)** 12 months. Color refers to differential gene expression direction and significance, and shape indicates receiver cell type. NS/DE = not significant or differentially expressed. Significant genes had an absolute log2FC > 0.2 with a Wald test padj < 0.05. **(C)** Bubble plot of functional enrichment terms of target genes at 6 months and **(D)** at 12 months. Condition denotes the context in which the target-associated interaction (i.e., the ligand-receptor pair) was upregulated. Plotted terms had Bonferroni adjusted p-values < 0.05.

### AD-risk gene-associated ligand-receptor pairs are 12-month time point-specific

To determine known AD associations of predicted interactions, we compiled an AD risk gene set of 245 genes from the Molecular Signatures Database (MSigDB)^44,45^ and a recent GWAS.^46^ Using this AD risk gene set, we filtered our pseudobulk differential expression analysis results. We found that 47 AD-risk genes were significantly differentially expressed across cell types and time points in 3xTg-AD vs. WT mice (Wald test padj < 0.05, absolute log2FC > 0.2; **Fig S3**). Interestingly, OPCs and microglia, which have previously been shown to highly express AD risk genes,^86^ did not have significant differential expression of any AD risk gene, and 12-month 3xTg-AD inhibitory neurons had the most significantly differentially expressed AD risk genes compared to WT (**Fig S3**). Overall, 12-month neurons (excitatory and inhibitory) had the most significantly differentially expressed AD-risk genes (34 and 13, respectively; **Fig S3**). Additionally, presenilin 1 (*Psen1*) was the only AD-risk gene upregulated in both time points in inhibitory and excitatory neurons between AD and WT, and apolipoprotein E (*Apoe*) was only differentially expressed in 6-month astrocytes.

After surveying AD-risk gene expression across cell types, we determined AD-risk gene-associated ligand-receptor pairs by filtering interactions using our AD-risk gene set. As none of the significantly differentially expressed AD-risk genes were among predicted ligands and receptors (**Fig S3**), we considered any ligand-receptor pair with a significantly differentially expressed AD-risk gene as a target as AD-associated. This resulted in 23 AD-risk gene-associated ligand-receptor pairs (**Fig 5A**) specific to 12-month-old mice. While we aimed to investigate the effects of altered glia-neuron communication and therefore focused on AD-risk gene-associated targets in receivers, we also noticed that 4 of 23 ligand-receptor pairs had either a ligand (*Calm2*) or receptor (*App*) that was in the AD-risk gene set. Interestingly, these interactions originated from microglia, with the highest AD-risk gene involvement overall, as we predicted 11 of 23 AD-risk gene-associated LRTs in microglia. Additionally, microglia were the only sender cell type to target two AD-risk genes uniquely (*Mme* and *Inpp5d;* **Fig S4**). Our findings corroborate those from a previous study that determined enrichment for AD-risk genes in neuron-microglia interactions in the human postmortem superior parietal cortex.^35^ The remaining 12 of 23 AD-risk gene-associated ligand-receptor pairs were predicted in multiple sender cell types (**Fig 5A**). Even though OPCs themselves did not significantly differentially express AD risk genes (**Fig S3**), they had the second most AD-risk target gene-associated ligand-receptor pairs (6 of 23), followed by astrocytes (4 of 23) and oligodendrocytes (1 of 23; **Fig 5A**). The 23 AD-associated ligand-receptor pairs had 6 predicted target genes which were significantly differentially expressed in the 12-month 3xTg-AD hippocampus: *Lpl*, *Ptk2b*, *Mme*, *Inpp5d*, *Cacna1c*, and *Adamts1* (**Fig 5B**). Out of 6 AD-risk associated targets, 5 were differentially expressed and predicted only in inhibitory neurons (**Top of Fig 5B**). *Adamts1* was the only AD-risk target gene downregulated in AD and was only predicted to be so in the 3xTg-AD excitatory neurons at 12 months. Moreover, *Lpl*, *Ptk2b*, *Mme*, *Inpp5d,* and *Cacna1c* were upregulated in 12-month 3xTg-AD inhibitory neurons compared to WT (**Fig 5B**).

**Figure 5.**
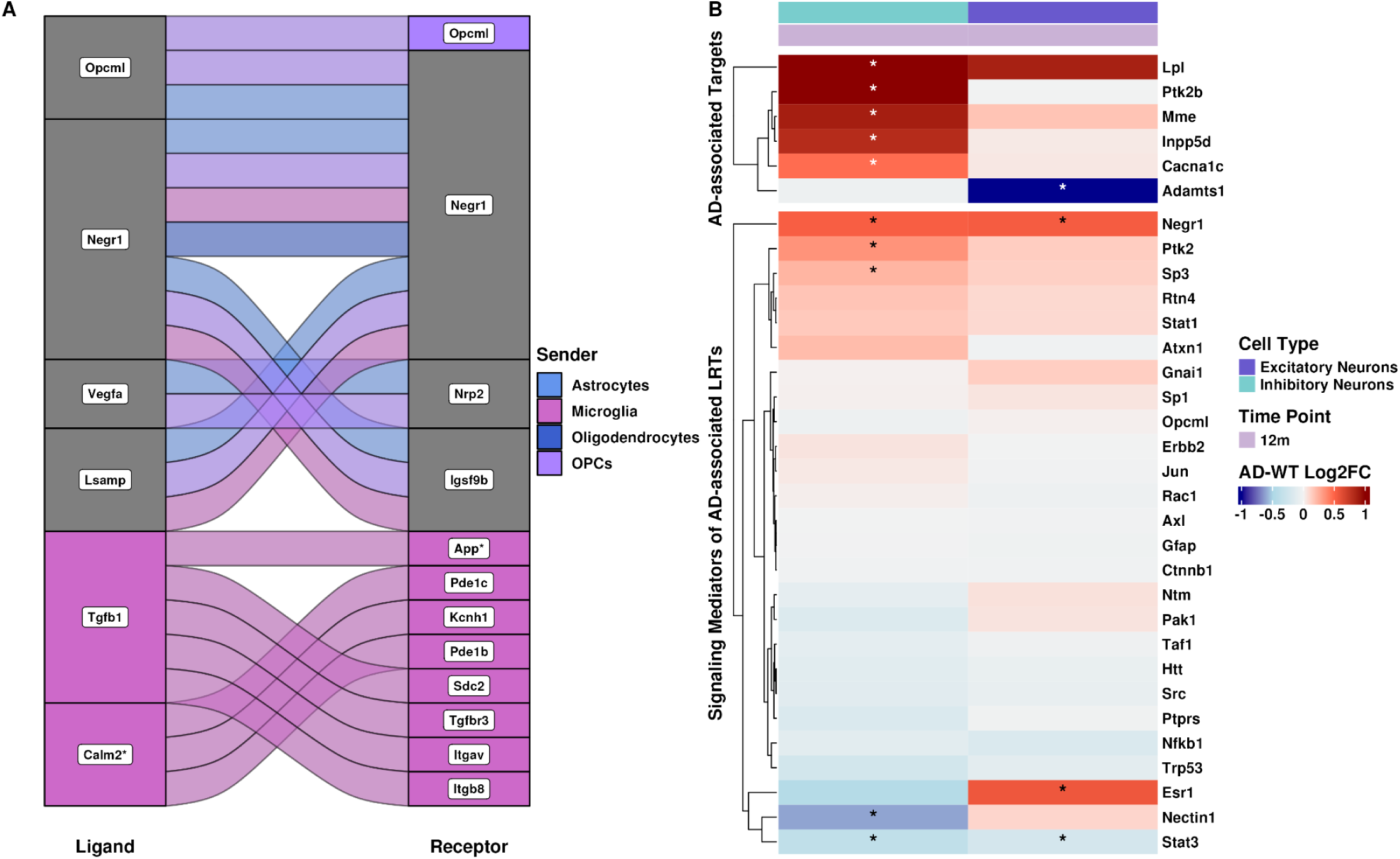
Significant AD-risk gene-associated interactions are specific to the 12-month 3xTg-AD hippocampus. **(A)** Alluvial plot of ligand-receptor pairs with an AD risk gene as a target. Color refers to the sender, and grey indicates the involvement of more than one sender. Asterisks indicate ligands and receptors that are an AD-risk gene. **(B)** Split heatmap of log2FC of target genes that are AD-risk genes (top) and their predicted signaling mediators (bottom) in 12-month receiver cell types. Asterisks indicate genes with an absolute log2FC > 0.2 and Wald test padj < 0.05 in the 3xTg-AD hippocampus.

To further investigate AD-risk gene-associated interactions, we constructed GRNs (**Supplementary File 1**) using ligand-receptor and target gene signaling information from the NicheNet v2 prior, a database that includes ligand, receptor, signaling mediator, and target gene information.^60^ By considering any node adjacent to our predicted receptors as potential signaling mediators of AD-associated LRTs, we identified 26 signaling mediators in our receiver cell types (**Bottom of Fig 5B, Table S8**). We used these AD-associated mediators to filter our pseudobulked inhibitory and excitatory neuron DEGs to investigate whether signaling mediators were differentially expressed in the 3xTg-AD hippocampus. Similar to our predicted target genes, most signaling mediators were not significantly differentially expressed in 12-month inhibitory and excitatory neurons (**Fig 5B**). However, *Negr1* and *Stat3* were significantly differentially expressed in both 12-month excitatory and inhibitory neurons, while *Ptk2*, *Sp3*, and *Nectin1* were significantly differentially expressed in 12-month inhibitory neurons and *Esr1* in 12-month excitatory neurons (Wald test padj < 0.05, absolute log2FC > 0.2; **Fig 5B**).

Overall, we found that AD-risk gene-associated ligand-receptor pairs were specific to 12-month 3xTg-AD neuronal subtypes. Additionally, LRTs predicted in microglia had the highest association with AD-risk genes. Since most signaling mediators were not significantly differentially expressed, this suggests that gene regulatory mechanisms might be driving downstream effects of altered glia-neuron communication in the 3xTg-AD hippocampus.

### 3xTg-AD signaling mediators were differentially regulated and had altered TF activity in excitatory and inhibitory neurons at 12 months

Since almost 81% of our predicted signaling mediators were not significantly differentially expressed in 3xTg-AD excitatory and inhibitory neurons, we investigated if these signaling mediators had perturbed GRNs and/or TF activity. We generated receiver-specific GRNs for each condition at 12 months using Passing Attributes between Networks for Data Assimilation (PANDA).^63,64^ Then, we calculated gene targeting scores (the sum of all inbound edge weights) for all genes in the network to quantify whether signaling mediators were differentially regulated in 3xTg-AD compared to WT hippocampus. We calculated the difference between 3xTg-AD and WT GRNs at 12 months for both receivers, where positive and negative scores indicate increased and decreased gene targeting in 3xTg-AD brain samples, respectively. When we investigated all genes detected in excitatory and inhibitory neurons, almost 64% of genes in excitatory neurons exhibited increased gene targeting, but only 41% of genes in inhibitory neurons exhibited increased gene targeting in the 3xTg-AD hippocampus (**Fig S5**). To determine whether our predicted signaling mediators were among the most differentially targeted genes, we calculated gene targeting score quartiles of all genes by receiver and further examined signaling mediators in the top and bottom quartiles (i.e., the most different signaling mediators).^67^ Interestingly, 17 of the 26 signaling mediators were among the most differentially targeted genes in 3xTg-AD excitatory and inhibitory neurons (**Fig 6A**). *Esr1*, which is a trigger of AD-associated neuroinflammation^87^, and *Nectin1* were the most differentially targeted mediators, indicating increased regulation by TFs in 3xTg-AD excitatory neurons and decreased regulation in 3xTg-AD inhibitory neurons (**Fig 6A**). Additionally, *Esr1* had a significant increase in gene expression in excitatory neurons and had non-significant decreased gene expression in inhibitory neurons (absolute log2FC > 0.2, padj < 0.05; **Fig 5B**). Considering the role of Esr1, glia-neuron communication may impact neuroinflammation through activation of this TF. In parallel, *Nectin1* had significantly less expression in 3xTg-AD inhibitory neurons and increased expression in excitatory neurons. Finally, *Stat3*, a transcriptional enhancer of autophagy-related genes^88^, had a significant decrease in expression in both receivers in AD (**Fig 5B**), and its overexpression ameliorates cognitive deficits in mice.^89^ *Stat3* was also only among the top quartile of differential gene targeting scores in 3xTg-AD inhibitory neurons, indicating differential regulation in this receiver (**Fig 6A**). However, mediators like *Ptk2*, *Stat1*, and *Gfap* in excitatory neurons had gene targeting scores close to 0, indicating very little change in their regulation in the 3xTg-AD hippocampus compared to WT. This may suggest that increased expression in excitatory neurons of *Ptk2* and *Stat1* was due to post-transcriptional, post-translational, or epigenetic modifications. Similarly, *Negr1*, which was significantly differentially expressed in inhibitory neurons (absolute log2FC > 0.2, padj < 0.05; **Fig 5B**), had a gene targeting score close to 0 in inhibitory neurons (**Fig 6A**). The signaling mediators *Sp1*, *Rtn4*, *Gfap,* and *Jun* also had gene targeting scores close to 0 in inhibitory neurons (**Fig 6A**). Even though gene targeting scores are not directly correlated with gene expression^66^, we observe a lack of strong differential expression of *Sp1*, *Gfap,* and *Jun* (absolute log2FC < 0.2; **Fig 5B**). Altogether, our findings imply that a set of the predicted AD-associated signaling mediators, which have known associations to neuroinflammation and autophagy, undergo changes in gene regulation in the 3xTg-AD hippocampus.

**Figure 6.**
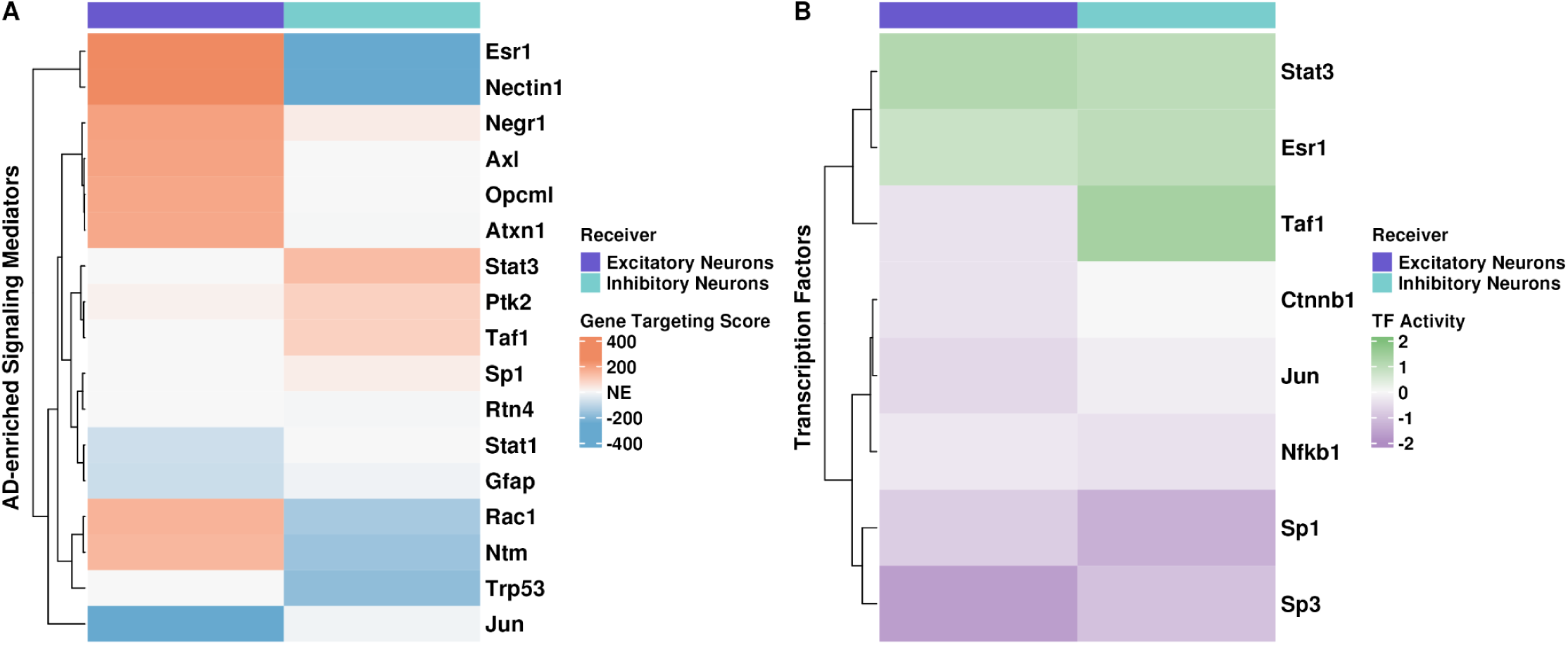
Signaling mediators were differentially regulated and had altered TF activity at 12 months. **(A)** Heatmap of differential gene targeting scores of signaling mediators in 12-month receivers. Differential gene targeting scores are in reference to AD. Red and blue represent increased (i.e., top quartile) and decreased (i.e., bottom quartile) gene targeting in the 3xTg-AD hippocampus, respectively. NE = not enriched in 3xTg-AD hippocampus (i.e., not among the highest and lowest quartiles/targeting scores). **(B)** Heatmap of differential TF activity scores of signaling mediators between 3xTg-AD and WT hippocampus, annotated by receiver, at 12 months. Green and purple represent TF activity as increased (i.e., active) or decreased (i.e., inactive), respectively.

In addition, since even small changes in TF gene expression can cause downstream regulatory effects,^90^ we investigated the TF activity of mediators that were TFs. We used pseudobulked gene expression information from excitatory and inhibitory neurons to calculate differential TF activity scores between 3xTg-AD and WT hippocampus for the 8 signaling mediators we identified that were also TFs. Positive values represent an active TF, and negative values represent an inactive TF in 3xTg-AD receiver cells compared to WT. The only TFs that were significantly expressed were *Stat3*, *Esr1*, and *Sp3* (absolute log2FC > 0.2, padj < 0.05; **Fig 5B**). The TFs Stat3 and Esr1 had increased activity in 12-month excitatory and inhibitory neurons (**Fig 6B**). Activation of Stat3 and Esr1 are associated with decreased autophagy^91^ and increased inflammation^87^, respectively. *Stat3* had significantly decreased gene expression in both receivers in the 3xTgAD hippocampus compared to WT, while *Esr1* and *Sp3* had a significant increase in gene expression in excitatory and inhibitory neurons, respectively (absolute log2FC = 0.2, padj < 0.05; **Fig 5B**). Except for the TF Taf1, TF activity scores were consistent in both receivers (**Fig 6B**). *Taf1* was not significantly differentially expressed (absolute log2FC < 0.2, padj < 0.05; **Fig 5B**) but had the highest TF activity in inhibitory neurons (TF activity score = 2; **Fig 6B**) and had decreased TF activity in 3xTg-AD excitatory neurons (TF activity score = −0.5; **Fig 6B**). Likewise, despite its increased expression in the 3xTg-AD hippocampus, Sp3 had the lowest TF activity score in excitatory neurons (TF activity score = −2; Fig 5B & 6B). Though *Sp3* was significantly differentially expressed (absolute log2FC > 0.2, padj < 0.05; **Fig 5B**) in inhibitory neurons, the TF Sp3 had the lowest TF activity score in inhibitory neurons (**Fig 5B & 6B**). We also see decreased TF activity in Ctnnb1 and Nfkb1, which are associated with WNT and NFkB canonical signaling. The TF encoded by Jun, a proto-oncogene, also had decreased TF activity in the 3xTg-AD mouse hippocampus (**Fig 6B**). Overall, we find changes in gene regulation of signaling mediators in hippocampal excitatory and inhibitory neurons of 3xTg-AD compared to WT mice at 12 months. This further suggests gene regulatory consequences in neurons of altered glia-neuron communication in the hippocampus of 3xTg-AD mice.

## Discussion

We generated snRNA-seq data from the hippocampus of female 3xTg-AD and age-matched WT mice at two clinically relevant time points for amyloid-β plaque formation^38^ and cognitive impairment^92^ (by 6 months) and gliosis^93^ and tau tangle formation^38^ (by 12 months) to investigate altered glia-neuron communication and their downstream effects throughout disease progression. Glial cells maintain neuronal homeostasis and health by interacting with neurons and regulating synaptic connectivity.^19–22,94^, and glial dysfunction has arisen as a powerful modulator of AD pathogenesis (reviewed in ^95^). Specifically, astrocytes fail to provide neurons with metabolic and nutritional support in models of AD.^96,97^ During aging^71^ and in the context of AD, ^68–70^ microglia also enter a chronic proinflammatory state, affecting neuronal function^28–30^ and disrupting phagocytosis in response to amyloid and tau pathology.^69^ Finally, OPCs facilitate synapse formation and directly communicate with neurons.^98^ This highlights the high cellular complexity of AD pathogenesis, and multiple studies have recently attempted to describe cell- and non-cell autonomous contributions to AD pathobiology.^95^ Indeed, multiple snRNA-seq studies on postmortem brain tissue from patients with AD and healthy age-matched controls have described dysregulated CCC in the postmortem human brain in AD.^34–37^ These studies emphasized the importance of CCC between glia and neurons, as astrocytes and microglia had increased involvement of AD-risk genes as ligands or receptors. Although these human studies have high disease relevance, they fail to capture early/pre-symptomatic CCC patterns relevant to disease progression, as extensive neuronal loss has already occurred. Using our newly generated snRNA-seq data in a well-established and highly relevant transgenic mouse model of AD pathology, we predicted 4,073 differentially expressed LRTs in the 3xTg-AD hippocampus compared to WT. This transgenic mouse, which includes three familial AD variants (APP Swedish, MAPT P301L, and PSEN1 M146V), exhibits amyloid-β and tau pathology as early as 6 and 12 months, respectively.^38^ Though plaque formation has been described to induce synaptic dysfunction in 3xTg-AD mice^38^ and other models (reviewed in ^99^), additional evidence suggests tau-induced synaptic dysfunction in AD.^100^ Interestingly, 86% of the dysregulated LRTs we predicted were in the 12-month 3xTg-AD hippocampus, which we showed had a significant increase of amyloid-β and tau pathology compared to 12-month-old WT mice. Since we did not see a significant neuropathological increase between 6- and 12-month-old 3xTg-AD hippocampus, the increase of dysregulated CCC at 12 months could be explained by some of the known ‘hallmarks of AD’, including age-associated loss of proteostasis, extended duration of amyloid-β and tau pathology, and/or widespread pathology as cortical regions show increased neuropathology^39^ that may affect hippocampal glia-neuron communication. While amyloid-β accumulates during the prodromal period of AD, patients do not exhibit cognitive decline until decades after initial depositions can be detected, indicating additional biological processes that can bolster cognitive function even in the face of amyloid-β toxicity. Therefore, our results suggest an effect of prolonged duration of amyloid pathology on CCC in the 3xTg-AD hippocampus but do not disqualify age-associated loss of proteostasis as a driver of the increased dysregulation of CCC at 12 months in 3xTg-AD mice.

Our differential gene expression analysis indicated that 6-month mice have smaller target gene log2FC values compared to 12-month mice, a finding independent of the number of nuclei per condition and cell type. This further demonstrates that impactful changes in neuronal gene expression profiles occur after the presence of amyloid-β and tau in 3xTg-AD mice. These findings are opposite of those in a bulk RNA-seq study in the insular cortex of 3xTg-AD mice, where the number of differentially expressed genes decreased over time.^101^ The MODEL-AD bulk RNA-seq data from the 3xTg-AD hippocampus revealed an increase in the number of differentially expressed genes between 4 and 12 months, followed by a decrease between 12 and 18 months.^102^ However, since bulk RNA-seq is performed on an entire tissue, cell-type-specific changes may be masked. Finally, we found that FEA terms of predicted targets were associated with signaling pathways at both time points, but 12-month targets included highly biologically relevant terms of signaling regulation through protein and tyrosine kinase activity. It is interesting that our predicted targets at 12 months in the 3xTg-AD hippocampus are associated with protein and tyrosine kinase activity in neurons. Tyrosine kinase inhibitors have previously been investigated as potential therapeutics in models of AD with neuroprotective effects.^103^ This suggests that our predicted interactions may produce novel therapeutic targets for drug repurposing or discovery in pre-clinical AD studies. Finally, our predicted ligand-receptor pairs should be further investigated for their utility as druggable targets, as previous work has identified drug repurposing candidates targeting dysregulated ligand-receptor pairs in the postmortem human AD prefrontal cortex.^34^

Our AD-associated LRTs were upregulated in the 12-month 3xTg-AD hippocampus and had significant differential expression of AD risk genes in either inhibitory or excitatory neurons. Importantly, many of these LRTs are known disease-modifying genes involved in key AD signaling pathways. For example, *Lpl*, which was significantly upregulated in 3xTg-AD inhibitory neurons, encodes a lipoprotein kinase, a key enzyme in lipid and lipoprotein metabolism.^104^ While we detected significant differential expression of *Lpl* in our transgenic mice, RNA-seq studies revealed its primary expression in human macrophages and microglia, in addition to OPCs from non-diseased/transgenic mice.^105,106^ Similarly, *Inpp5d*, associated with autophagy and inflammasome activation in microglia,^107^ is significantly differentially upregulated in our inhibitory neurons, has previously been described in microglia using 5xFAD transgenic mice and INPP5D-disrupted iPSC-derived human microglia.^107–109^ *Inpp5d* and *Mme* were the only two target genes uniquely targeted by interactions from microglia. MME encodes neprilysin, an enzyme responsible for amyloid-β degradation.^110–112^ Neprilysin injections decreased amyloid-beta oligomers in an AD mouse model.^110^ *Cacna1c*, a psychiatric risk gene identified as an AD-risk gene by GWAS,^113^ encodes a calcium channel. Expression of the CACNA1C protein was previously identified as increased in the hippocampus of the transgenic AD mouse model APP/PS1.^114^ The gene also plays a role in oxidative stress pathways, which have been demonstrated to contribute to AD pathology (reviewed in ^115^). Finally, we describe significantly increased expression of *Ptk2b* in inhibitory neurons, but previous studies on *Ptk2b* have been inconclusive. Deletion of *Ptk2b* in APP/PS1 transgenic mice rescued synaptic loss and memory deficits,^116^ while overexpression rescued behavior and increased amyloid plaque number in 5xFAD mice.^117^ A human-derived HEK293T cell line study indicated that *Ptk2b* contributed to tau phosphorylation, a finding that could not be recapitulated in iPSC-derived human neurons.^118^ Lastly, *Adamts1* was significantly downregulated in our excitatory neurons at 12 months. The gene’s protein product has been suggested as a marker protein for neurodegeneration,^119^ and hippocampal amyloid-β load was alleviated after introducing ADAMTS1 in a mouse model of AD.^120^ The robust representation of AD-associated gene targets identified by this screen highlights the relevance of altered glia-neuron communication. Considering that five of six predicted AD-risk target genes were significantly differentially expressed in inhibitory neurons, it is plausible that altered glia-neuron communication affects inhibitory neurons more severely than excitatory neurons in the 12-month 3xTg-AD hippocampus, although this hypothesis remains to be tested. Four ligand-receptor pairs with AD risk genes also included an AD risk gene as a ligand (*Calm2*) or receptor (*App*) of interactions originating from microglia, another previously identified key node of AD-pathogenesis. Therefore, our results corroborate previous findings indicating increased association to AD-risk genes of microglia-associated CCC.^34–37^

Most signaling mediators of predicted AD-associated interactions were not significantly differentially expressed in the 3xTg-AD hippocampus. Due to processes like post-transcriptional modifications and downstream differential expression of additional mediators, signaling mediators can be differentially regulated, regardless of the lack of significant differential expression in inhibitory and excitatory neurons. Our differential gene targeting analysis revealed that some signaling mediators were differentially regulated but not significantly differentially expressed. For example, we found that *Stat1* and *Ptk2*, which had increased expression in inhibitory neurons, had gene targeting scores close to 0, indicating they were not differentially regulated in AD. Along with previous work, this suggests that changes in their gene expression may be due to post-transcriptional, post-translational, or epigenetic modifications.^121–123^ Additionally, eight signaling mediators that are also TFs had differential activity in both 3xTg-AD excitatory and inhibitory neurons at 12 months. In line with the FEA of predicted target genes, we found that altered glia-neuron communication may affect gene regulatory mechanisms in excitatory and inhibitory neurons of 12-month 3xTg-AD mice. Using human single-nucleus multi-ome data, Gupta et al. identified TFs that disrupt cell-type-specific GRNs in AD patients and determined their utility in predicting drug repurposing candidates.^124^ Interestingly, Gupta et al. identified ESR1 as a master regulator across microglia, oligodendrocytes, and excitatory and inhibitory neurons in their study. We predicted *Esr1* as a differentially regulated signaling mediator in excitatory and inhibitory neurons (increased and decreased differential gene targeting, respectively) despite being only significantly differentially expressed in excitatory neurons. However, the TF *Esr1* had increased TF activity in both receivers. This further suggests changes in gene regulatory mechanisms due to glia-neuron communication in the 3xTg-AD mouse hippocampus, especially at the later stages of the disease progression.

Although models like the 3xTg-AD mouse mimic AD pathology, they do not fully recapitulate human disease pathology and progression. We have previously shown that CCC between glia and neurons is altered in AD in the postmortem human prefrontal cortex using snRNA-seq data.^36^ Previously, we validated two ligand-receptor pairs across three independent human patient AD cohorts that were semaphorin-plexin interactions, which are associated with tau tangle colocalization^125^ and phosphorylation^125,126^ in AD. Interestingly, we also predicted two semaphorin-plexin interactions in our 3xTg-AD hippocampus data (Sema4c - Plxnb2 and Sema4a - Plxna4), which originated from microglia and astrocytes, respectively. These findings, in addition to those from our human study, further suggest the need for additional investigation of changes in semaphorin-plexin signaling throughout aging and in AD. Additionally, six ligand-receptor pairs overlapped between our postmortem human prefrontal cortex and 3xTg-AD hippocampus studies. However, the ligand-receptor pair TGFB1-APP/Tgfb1-App was the only interaction originating from the same sender (microglia) and upregulated in AD in both studies. This interaction was among our high-confidence and sex-specific interactions, as we predicted it in two independent AD patient cohorts before accounting for patient sex,^36^ which had an AD-risk gene as a downstream target in the 3xTg-AD hippocampus at 12 months. Many drugs targeting amyloid-β have been tested for AD in the last 20 years with conflicting results.^127–129^ Interestingly, Tamoxifen, an estrogen modulator, and Benazepril, an angiotensin-converting enzyme (ACE) inhibitor, are predicted drug repurposing candidates that inverse the activity of the TGFB1 ligand. Women receiving Tamoxifen are less likely to be diagnosed with AD and show improved cognitive performance.^130,131^ Moreover, Tamoxifen prevents learning and memory impairments in ovariectomized rats.^132^ Finally, patients treated with ACE inhibitors like Benazepril have a slowed cognitive decline.^133^ While the TGFB1-APP/Tgfb1-App interaction validates across human datasets and in our 3xTg-AD mice, whether this interaction occurs in additional human datasets and AD models remains to be seen.

While our study provides insight into dysregulated glia-neuron interactions and their downstream effects across two time points in a transgenic mouse model of AD, there are a few limitations, such as using a single AD mouse model and curated priors for CCC, GRN, and TF activity analyses. We used the hippocampus from female 3xTg-AD mice; therefore, future studies should expand this work to the hippocampus of male 3xTg-AD mice since amyloid-β and tau pathology has been shown to differ between male and female 3xTg-AD mice.^134^ Future work should also expand our work to include additional brain regions, such as the cerebral cortex, to determine whether changes in glia-neuron communication are brain region-specific since AD pathology affects multiple brain regions throughout AD progression in patients.^9,10^ We focused on two time points, corresponding to the onset of amyloid-β and tau pathology; however, including additional time points from presymptomatic (3 months) or aged (20 months) mice might provide further insight into the effects of disease progression and aging on altered CCC. Additionally, mice do not fully recapitulate human disease presentation, therefore, future studies should include human tissues to confirm our findings. Additionally, CCC inference analysis relies on the use of curated priors. Therefore, we cannot predict biologically relevant but previously undescribed interactions, limiting our ability to identify novel interactions. Furthermore, CCC inference methodologies use gene expression information to deduce ligand and receptor protein abundance. However, due to protein degradation and post-translational modifications, mRNA and protein levels are not directly affiliated. Finally, future studies should further investigate the effects of amyloid-β and tau pathology on glia-neuron communication to pinpoint whether the lack of differential CCC at 6 months in our study is due to a lack of tau pathology or low levels of amyloid-β at this time point. Overall, our study lays the groundwork for additional validation in future studies to confirm that the predicted LRTs affect cell-type-specific gene regulatory mechanisms, as they remain largely understudied in AD, especially throughout disease progression.

## Conclusion

We report that glia-neuron communication is altered in a time point-specific manner in the hippocampus of 3xTg-AD mice using snRNA-seq. We find that CCC is increasingly dysregulated in 12-month AD mice. Additionally, we identify 23 ligand-receptor pairs that are upregulated in the 12-month-old 3xTg-AD hippocampus and have an AD risk gene as a downstream target. We also find increased AD association in interactions originating from microglia. Finally, we describe altered regulation and TF activity of predicted signaling mediators, which were not significantly differentially expressed. Therefore, our findings suggest that altered glia-neuron communication affects the gene regulatory mechanisms in neurons of 3xTg-AD mice.

## Abbreviations

snRNA-seq: single-nucleus RNA sequencing
AD: Alzheimer’s disease
PSEN1: presenilin 1
PSEN2: presenilin 2
APP: amyloid precursor protein
CCC: cell-cell communication
OPC: oligodendrocyte progenitor cell
WT: wild-type
MSigDB: Molecular Signatures Database
GWAS: Genome-wide association study
LRT: ligand-receptor-target
TF: transcription factor
FACS: fluorescence-activated cell sorting
BCA: Bicinchoninic acid assay
PCA: Principal component analysis
log2FC: log2FoldChange
PANDA: Passing Attributes between Networks for Data Assimilation
CIS-BP: Catalog of Inferred Sequence Binding Preferences
UMAP: Uniform Manifold Approximation and Projection
NFkB: NF-kappa-B
FEA: Functional enrichment analysis
ACE: angiotensin-converting enzyme
GRN: Gene regulatory network

## Author contributions

TMS and BNL conceptualized the project. DCP and AB maintained mouse colonies and collected tissues. TMS, JHW, and TCH extracted nuclei for snRNA-sequencing. TMS, TCH, and ADC performed protein quantification. TMS performed alignment and pre-processing. JHW constructed gene regulatory networks and aided in differential gene targeting analyses. All other analyses were coded and performed by TMS. EJW, TCH, and ADC reviewed and validated the code. BNL provided supervision and project administration. BNL, TMS, and JHW acquired funding. TMS wrote the first draft. TMS, TCH, EJW, JHW, ADC, DCP, AB, CJC, and BNL reviewed and edited the manuscript. All authors read and approved the final manuscript.

## Acknowledgments

The authors thank the Lasseigne Lab members for their feedback throughout this study. Specifically, Vishal H. Oza for his expertise in differential gene targeting analyses and Emma F. Jones for aiding with data submission and graphical abstract design. Furthermore, the authors thank Dr. Anna Thalacker-Mercer’s lab at UAB for providing equipment and reagents for the BCA and amyloid-β and total tau protein quantification using ELISA experiments. In addition, we thank Shanrun Liu at the UAB CFCC for his expertise and preparation of snRNA-seq libraries for this study, as well as Michael Crowley and the team at the UAB Heflin Center for Genomic Sciences at the UAB Sequencing Core for sequencing the samples.

## Data availability statement

The 3xTg-AD snRNA-seq dataset is deposited on NCBI’s Gene Expression Omnibus (GSE261596). The intermediate outputs of this study are available on Zenodo (doi: https://zenodo.org/records/11043321). The code supporting the results of this study is available on Zenodo (doi: https://zenodo.org/records/11040825) and GitHub (https://github.com/lasseignelab/230418_TS_AgingCCC). Docker images used for these analyses are publicly available on Docker Hub (https://hub.docker.com/repository/docker/tsoelter/rstudio_aging_ccc/general) and Zenodo (doi: https://zenodo.org/records/11042577).

## Ethics approval statement

We carried out all animal experiments in this study according to the Institutional Animal Care and Use Committee at the University of Alabama at Birmingham.

## Patient consent statement

Not applicable

## Permission to reproduce material from other sources

Not applicable

## Clinical trial registration

Not applicable

