## Supplementary Information for "Evaluation of altered cell-cell communication between glia and neurons in the hippocampus of 3xTg-AD mice at two time points"

**Table S1: Post-alignment sequencing quality metrics by sample.**

| Sample | Estimated Number of Cells | Mean Reads per Cell | Median Genes per Cell |
|---|---|---|---|
| S01_6m_AD | 14 200 | 39 745 | 2 667 |
| S02_12m_AD | 15 159 | 34 991 | 2 514 |
| S03_6m_WT | 12 154 | 43 051 | 2 960 |
| S04_12m_WT | 18 205 | 27 331 | 2 068 |
| S05_6m_AD | 15 897 | 30 440 | 2 404 |
| S06_12m_AD | 15 765 | 30 703 | 2 403 |
| S07_6m_WT | 16 040 | 28 838 | 2 505 |
| S08_12m_WT | 13 268 | 30 075 | 2 364 |
| S09_6m_AD | 17 907 | 32 224 | 2 703 |
| S10_12m_AD | 19 846 | 28 446 | 2 300 |
| S11_6m_WT | 15 834 | 30 410 | 2 402 |
| S12_12m_WT | 11 368 | 48 714 | 2 940 |
| **Total** | **185 643** | **404 968** | **30 230** |
| **Average per Sample** | **15 470** | **33 747** | **2 519** |

**Table S2: Optical density values from BCA and amyloid-β 40, amyloid-β 42, and total tau ELISAs.**
Table_s2.xlsx

**Figure S1: Protein quantification of human amyloid-β 40, amyloid-β 42, and total tau using ELISAs**
Box plots of protein concentration across groups (age and condition) of (**A**) amyloid-β 40 protein (μg/mL), (**B**) amyloid-β 42 (μg/mL), and (**C**) total tau protein (log10 transformed μg/mL). Light and dark blue refer to AD and WT, respectively. N = 3 mice per group. **adjusted p-value < 0.01, ***adjusted p-value < 0.001 one-sided ANOVA followed by Tukey's test for multiple hypothesis correction.

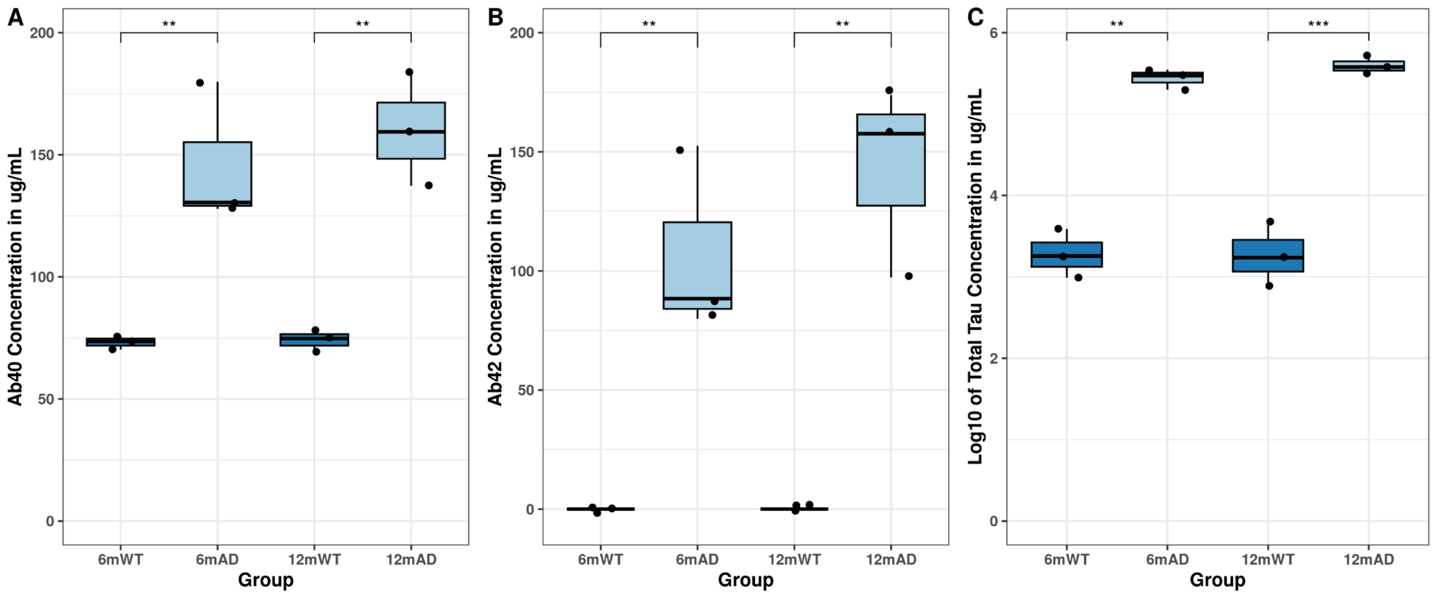


**Figure S2: Number of predicted cell-cell communication interactions.**

(**A**) Barplot showing the total number of predicted interactions (83,928) across all cell types in the dataset. (**B**) Total number of interactions filtered by sender and receiver (excitatory and inhibitory neurons) cell types of interest, and after prioritization using ligand activity and regulatory potential scores (Pearson and Spearman correlation values of greater than 0.33 and smaller than -0.33) colored by sender (**C**) and colored by receivers in a stacked bar plot.

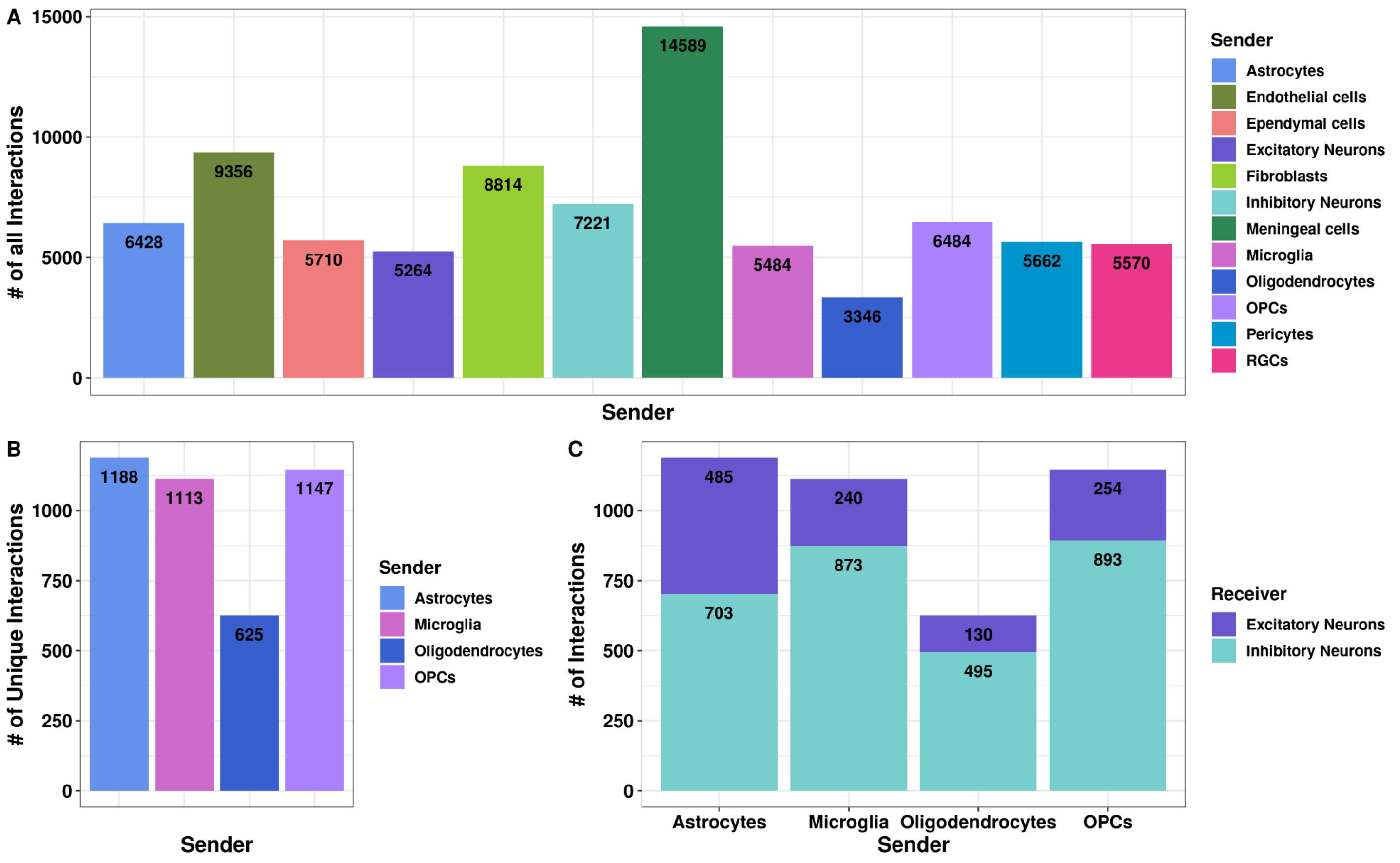

**Table S3: Table of marker genes.** Marker genes were upregulated in their respective clusters and had an average log2fc > 0.2. Table_s3.xlsx

**Table S4: Canonical marker genes**

Table includes marker genes that were used for assignment of major cell types in our 3xTg-AD dataset using feature plots.

| **Cell type** | **Marker(s)** |
|---|---|
| Excitatory Neurons | Slc17a6, Slc17a7 |
| Inhibitory Neurons | Gad1, Gad2 |
| Astrocytes | Slc1a2, Gpc5 |
| Oligodendrocytes | Mbp, Plp1, St18 |
| OPCs | Vcan, Tnr, Lhfpl3 |
| Meningeal cells | Ranbp3l, Slc6a20a |
| Microglia | Dock8, Tgfbr1 |
| Ependymal cells | Spag16, Dnah12 |
| Fibroblasts | Bnc2 |
| Pericytes | Pdgfrb, Cald1 |
| Endothelial cells | Flt1, Mecom |
| RGCs | Slc18a2, Klhl1 |

**Table S5: AD risk gene set.**

Table of AD risk genes along with their dataset of origin.

Table_s5.xlsx

**Table S6: LRTs and their overlap across groups**

Table of LRTs showing their overlap between groups and group-specificity.

Table_s6.xlsx

**Table S7: Group-specific target gene expression**

Table of group-specific target genes (12mAD, 12mWT, 6mAD, and 6mWT) log2FoldChanges and adjusted p-values. This excludes targets that were shared across groups or time points.

Table_s7.xlsx

**Figure S3: Gene expression of AD risk genes across senders and receivers.**
Heatmap of significantly differentially expressed AD risk genes across senders and receivers (padj < 0.05, absolute log2FoldChange > 0.2 ) in AD compared to WT in 6- and 12-month samples. AD risk genes in OPCs did not meet cut-offs and were excluded from this plot.

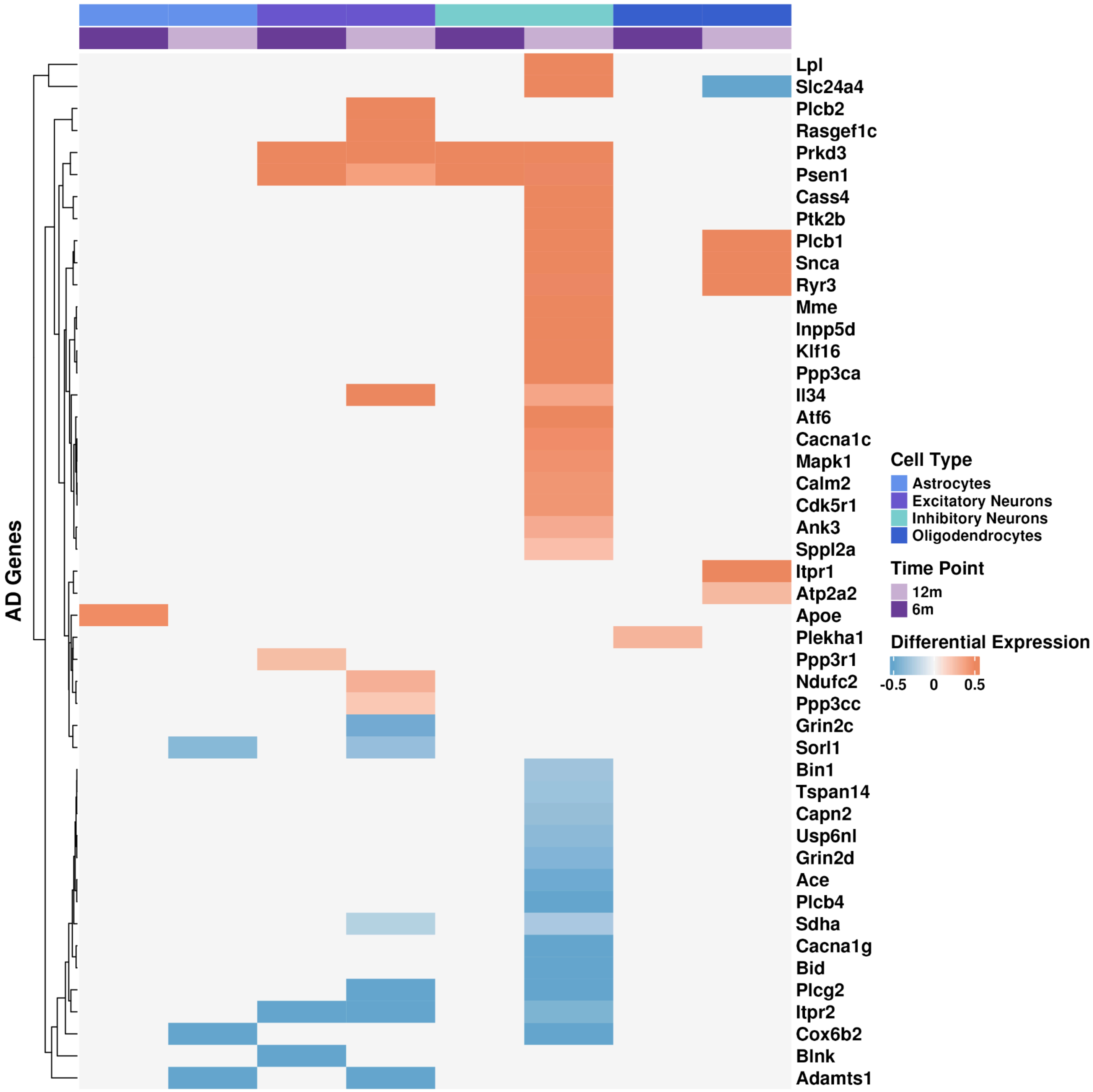

**Figure S4: AD-associated ligand-receptor-target pairings.**
Alluvial plot of ligand-receptor pairs and their target gene, which is an AD risk gene. The color indicates the sender, and grey refers to ligands, receptors, or targets that we predicted in more than one sender.

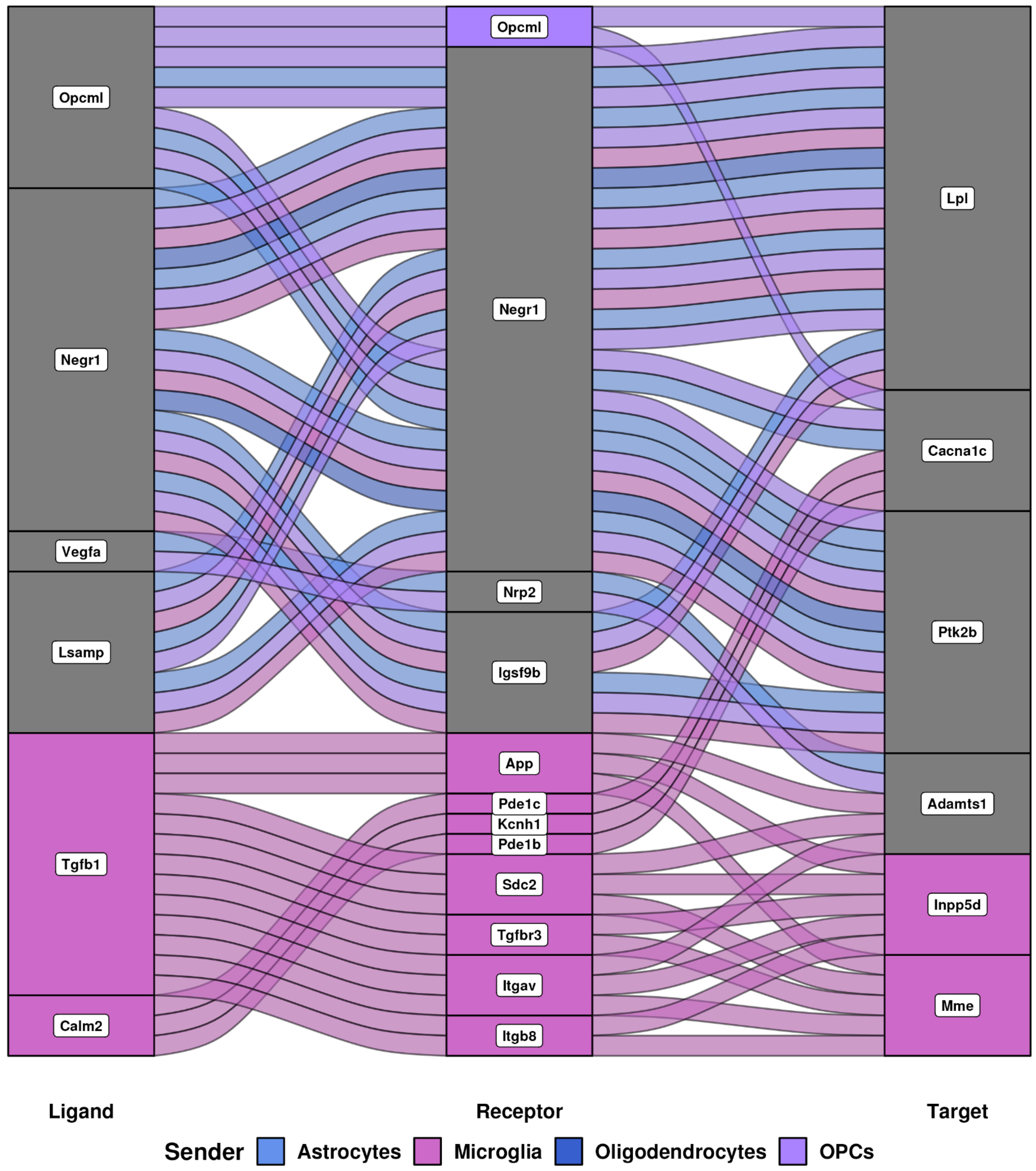


**Table S8: Signaling mediators of AD-associated interactions across receivers.**

| L-R Pair | Inhibitory Target | Excitatory Target | Inhibitory Mediators | Excitatory Mediators |
|---|---|---|---|---|
| Lsamp - Negr1 | *Lpl*, *Ptk2b* | *Lpl*, *Gm3839* | *Htt* | *Htt* |
| Opcml - Opcml | *Cacna1c, Fas* | N/A | *Erbb2, Ntm, Opcml, Atxn1* | N/A |

| | | | | |
|---|---|---|---|---|
| Opcml - Negr1 | *Cacna1c, Fas, Lpl, Ptk2b* | N/A | *Htt, Negr1, Ntm* | N/A |
| Negr1 - Negr1 | *Itpr1, Lpl, Ptk2b, Fas* | *Lpl, Gm3839* | *Gfap, Htt, Negr1, Taf1* | *Gfap, Htt, Negr1* |
| Negr1 - Igsf9b | *Itpr1, Lpl, Ptk2b, Fas* | N/A | N/A | N/A |
| Tgfb1 - App | *Inpp5d, Mme* | *Adamts1* | *Trp53* | *Jun* |
| Tgfb1 - Sdc2 | *Inpp5d, Mme* | *Adamts1* | N/A | *Jun* |
| Tgfb1 - Tgfbr3 | *Inpp5d, Mme* | N/A | N/A | N/A |
| Tgfb1 - Itgav | *Inpp5d, Mme* | *Adamts1* | *Trp53* | N/A |
| Tgfb1 - Itgb8 | *Inpp5d, Mme* | N/A | N/A | N/A |
| Calm2 - Pde1c | *Cacna1c, Fas* | N/A | N/A | N/A |
| Calm2 - Pde1c | *Cacna1c, Fas* | N/A | N/A | N/A |
| Calm2 - Kcnh1 | *Cacna1c, Fas* | N/A | *Trp53* | N/A |
| Vegfa - Nrp2 | N/A | *Adamts1* | N/A | N/A |
| Efnb3 - Epha10 | *Fas* | N/A | *Rac1* | N/A |
| Efnb3 - Ephb2 | *Fas* | N/A | *Pak1, Rac1, Stat3* | N/A |
| Kcna1 - Cntnap1 | *Fas* | N/A | *Rtn4* | N/A |
| Ptgfrn - Tmem59l | *Fas* | N/A | N/A | N/A |
| Nectin1 - Nectin1 | *Fas* | N/A | *Ctnnb1, Nectin1, Nfkb1* | N/A |
| Sema4a - Plxna4 | N/A | *Gm3839* | N/A | N/A |
| Sema4c - Plxnb2 | *Itpr1* | N/A | N/A | N/A |
| Ntm - Negr1 | *Fas* | *Gm3839* | N/A | N/A |
| Ntm - Thsd7a | *Fas* | N/A | N/A | N/A |
| Ntm - Lrrn3 | *Fas* | N/A | N/A | N/A |
| Ncam1 - Nptn | *Fas* | N/A | N/A | N/A |
| Edil3 - Itgav | *Fas* | N/A | *Ptk2, Stat1, Trp53* | N/A |
| Edil3 - Itga5 | *Fas* | N/A | *Ptk2, Trp53* | N/A |
| Pros1 - Tyro3 | *Fas* | N/A | *Axl, Src* | N/A |

| | | | | |
|---|---|---|---|---|
| Tcn2 - Cnr1 | *Fas* | N/A | *Esr1, Gnai1, Sp1, Sp3* | N/A |
| Slitrk5 - Ptprd | N/A | *Gm3839* | N/A | *Ptprs* |
| C4b - Nrp1 | N/A | *Gm3839* | N/A | *Ptk2* |

**Figure S5: Global differential gene targeting in 3xTg-AD mice at 12 months.**
Dotplot showing the number of genes that have increased (red) or decreased (blue) gene targeting in 12-month 3xTg-AD mouse hippocampus in receiver cell types.

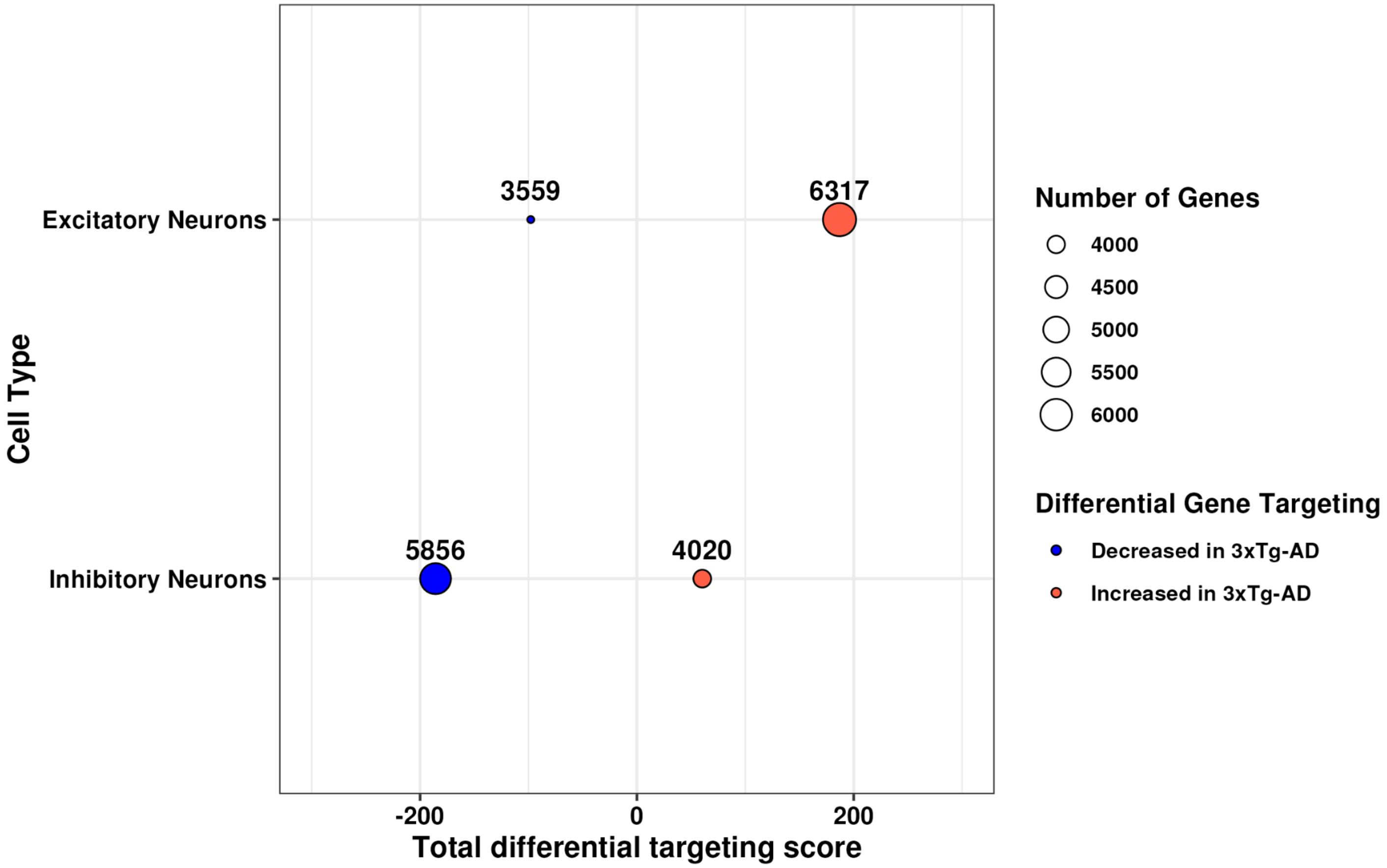
