## Supplementary Materials & Methods for "Evaluation of altered cell-cell communication between glia and neurons in the hippocampus of 3xTg-AD mice at two time points"

### Supplementary Materials and Methods:

Animals

We carried out all animal experiments in this study according to the Institutional Animal Care and Use Committee at the University of Alabama at Birmingham. 3xTg-AD mice (B6;129-Tg(APPSwe,tauP301L)1Lfa Psen1tm1Mpm/Mmjax: JAX MMRRC Stock# 034830) and WT littermates were bred and housed at animal facilities at the University of Alabama at Birmingham with standard light/dark cycles and ad libitum access to food and water. Before collection, animals were anesthetized using Avertin intraperitoneal injections before perfusion with 1X PBS. We dissected relevant brain regions and then snap-froze and stored hippocampi at -70C until nuclei isolation of all samples for sequencing to minimize batch effects and decouple tissue collection and processing. Due to our sample collection method[1], we performed snRNA-seq.

Nuclei isolation and snRNA sequencing of 3xTg-AD mouse hippocampus

We adapted a previously published rat nuclei isolation protocol for our 3xTg-AD mouse brain.[2] We isolated nuclei from the left hippocampus of 12 female mice (6 AD, 6 WT) from two time points (6 and 12 months; n = 3 per time point and condition). Following this, we performed fluorescence-activated cell sorting (FACS) on an Atlas BD ARIA II to remove debris and retain a pure nuclei suspension. We sorted for 200,000 nuclei per sample to account for nuclei loss during nuclei partition and library preparation. The UAB Flow Cytometry and Single Cell Core Facility prepared sequencing libraries according to the manufacturer's instructions using the Chromium Single Cell 3' GEM, Library & Gel Bead Kit v3 (10x Stock #: PN-1000121). Finally, we sequenced all twelve libraries in a single batch on an Illumina NovaSeq 6000 using an S4 flowcell at the UAB Heflin Center for Genomic Sciences to an average depth of ~34,000 reads per nucleus with ~15,000 nuclei per sample (see ranges in **Table S1**).

Protein quantification of amyloid-β 40, amyloid-β 42, and total tau

We used the human Amyloid-β 40 (ThermoFisher Cat No. KHB3481), human Amyloid-β 42 (ThermoFisher Cat No. KHB3441), and human Total Tau (ThermoFisher Cat No. KHB0041) ELISA kits to quantify protein abundance in the hippocampus for both conditions and time points (n = 3 per time point and condition). We weighed the right hippocampi and combined tissues with 1X PBS with AEBSF proteinase inhibitor at a 1:10 dilution. We homogenized hippocampi before extracting and centrifuging homogenates at 16,000 x g for 20 minutes at 4ºC. We combined 200uL of our samples with 200uL of 1X PBS with AEBSF proteinase inhibitor for all samples and performed a Bicinchoninic acid assay (BCA) to quantify the total protein in our samples (**Table S2**). We further diluted samples for the human Total Tau ELISA (1:100 dilution) using total protein abundance measured using the BCA. We loaded samples and blanks into all three pre-coated ELISA microwell plates and performed incubation and manual washing as described in the human Amyloid-β 40, human Amyloid-β 42, and human Total Tau ELISA kit protocols. We measured absorbance using a Tecan Infinite M Plex plate reader at 450 nm with 10 flashes (**Table S2**). By fitting the absorbance with a 4-parameter algorithm, we quantified protein concentration in ug/mL for every ELISA. We compared groups using a one-way ANOVA followed by a Tukey's test for multiple hypothesis correction and to determine which groups were statistically different from each other.

Data Processing and Quality Control

We aligned raw FASTQ files on UAB's high-performance computer using 10x Genomics Cell Ranger[3] 6.1.1 to the 10x Genomics mouse reference genome (mm10). We performed all downstream data processing in docker[4] R v4.2.3. We removed ambient RNA using soupX[5] v1.6.2 on Cell Ranger h5 files, resulting in filtered Cell Ranger output matrices. We converted the filtered Cell Ranger output matrices to expression matrices using the *Read10x* function from the Seurat v4.3.0[6]. We created a Seurat object using the *CreateSeuratObject* function for each sample, time point, and condition (AD and WT) before merging them into a single Seurat object. We filtered the dataset at the cell level using multiple quality control metrics (mitochondrial ratio, number of genes per cell, and log10 of the number of genes per UMI) and at the gene level by removing zero count values to prevent skewing of average expression values. We removed nuclei with a mitochondrial ratio higher than 5% and a complexity score greater than 80% (number of genes per UMI). We also filtered based on the number of genes per cell, removing nuclei with less than 500 and more than 10000 genes per nucleus. We performed batch correction using harmony v0.1.0[7], as it preserves biological variation while reducing variation due to technical noise.[8] We also performed Principal Component Analysis (PCA) using the *RunPCA* function from Seurat v4.3.0[6] without approximation (approx = FALSE), scaled and normalized before plotting UMAPs to confirm successful integration across conditions.

Clustering and Cell Type Identification

We compared multiple resolutions (0.5 - 1.5) using clustree v0.5.0[9]. We chose 0.7 resolution as its clustering showed the greatest stability (i.e., fewest switches of nuclei between clusters), which yielded 40 clusters using Leiden v0.4.3[10]. We identified differentially expressed marker genes for each cluster using the *FindAllMarkers* function on the RNA assay from Seurat v4.3.0[6] with a log fold change threshold > 0.2. We assigned cell types using differential expression of cell-type-specific genes identified through PanglaoDB[11] and CellMarker 2.0[12] (**Table S3**) and feature plots using canonical cell type markers (**Table S4**).

Pseudo-bulking of snRNA-seq Data

We pseudo-bulked to generate cell-type-specific count matrices to overcome the data sparsity in snRNA-seq.[13] We converted raw data from the Seurat object *counts* slot into Single Cell Experiment objects using SingleCellExperiment[14] v1.20.1 and incorporated metadata with information on sample origin (sample_id), condition (group_id), cell type (cluster_id), and time point (timepoint). We aggregated counts across samples by cell type using the *aggregate.Matrix* function (Matrix.utils R package v0.9.7), resulting in cell-type-specific gene by sample count matrices.

Differential Gene Expression Analysis

We performed differential expression analysis between conditions (AD vs. WT) at both time points (6 and 12 months) using DESeq2[15] v1.38.3 for each cell type. First, we created dds objects using the *DESeqDataSetFromMatrix* function. We made pairwise comparisons for every cell type between conditions for each time point independently using the Wald test. Then we performed log fold change shrinkage to estimate effect size for calculation of log2FoldChange changes using apeglm v1.23.1.[16]

Inference of CCC Across Conditions and Time Points

To infer CCC between glia and neurons between AD and WT across time points (6 and 12 months), we applied multinichenetr v1.0.3 and used the Nichenet v2 prior.[17] As MultiNicheNet requires inputs in Single Cell Experiment format, we converted our Seurat objects using the SingleCellExperiment R package[14] v1.20.1. Then, we performed ligand-receptor pair prediction between all cell types for every time point between conditions before we filtered for senders (astrocytes, microglia, oligodendrocytes, and OPCs) and receivers (excitatory and inhibitory neurons). Since MultiNicheNet uses pseudobulk aggregation for its differential expression analysis, we ensured that all of our samples had a minimum of 10 cells per cell type. During count aggregation, MultiNicheNet normalizes pseudobulk count matrices by library size (i.e., the number of cells per cell type) to account for differences in cell type proportions. As recommended by the package developers, we identified differentially expressed ligands, receptors, and target genes using a 0.5 log2FC threshold and non-adjusted p-values of 0.05. When calculating ligand activity, we considered the top 250 targets with the highest regulatory potential (calculated for every ligand-target pairing within the NicheNet v2 prior). The calculation of the ligand-target regulatory potential is described in the original NicheNet publication (see Supplementary Note 3).[18] We prioritized ligand-receptor interactions by min-max scaling the scores for all comparisons of the following metrics: differential expression of ligand, differential expression of the receptor, the fraction of ligand-receptor pairs expressed across samples, expression of ligand, expression of the receptor, scaled ligand activity, the abundance of the sender in the condition of interest, and abundance of the receiver in the condition of interest. Then, we performed weighted aggregation of the scaled scores using default MultiNicheNet prioritization weights. We calculated expression correlation information between ligand-receptor pairs and their targets by calculating Pearson and Spearman correlation coefficients to further prioritize LRT interactions. We filtered LRTs and included only those with Spearman and Pearson correlation coefficients greater than 0.33 and less than -0.33, as recommended by MultiNicheNet.

JI of Ligands, Receptors, and Target Genes Across Cell Types
We calculated the JI to determine the degree of overlap of ligands, receptors, and target genes between sender or receiver cell types. We included interactions from both time points and conditions to get a global view of the degree of overlap instead of calculating JI for each time point and condition independently. First, we calculated JI between receiver cell types (excitatory and inhibitory neurons) based on the overlap of receptors and target genes. Then, we calculated JI between sender cell types (astrocytes, microglia, oligodendrocytes, and OPCs) based on the overlap of ligands, receptors, and target genes.

Functional Enrichment Analysis of Predicted Target Genes
To determine the molecular function of predicted target genes from MultiNicheNet, we identified significantly enriched pathways using gprofiler2[19] v0.2.1 from Gene Ontology Molecular Function (GO:MF), Reactome, and Transfac source gene sets. We queried predicted target genes from both receiver cell types combined (excitatory and inhibitory neurons) to find overrepresented functional enrichment of receiver cell types. We identified pathways for targets shared across time points and conditions and targets specific to each group (time point and condition pairings: 6-month AD, 6-month WT, 12-month AD, 12-month WT). We included all genes measured in our receivers without filtering as our background gene set. We filtered pathways by whether they were significant after multiple hypothesis correction using Bonferroni ($q < 0.05$) as well as the term size (> 10 and < 1000) to remove arbitrary pathways, similar to Wilk et al. 2023.[20] We plotted the top 10 terms per condition based on recall.

AD risk gene set curation
We compiled an AD risk gene set by combining two human gene sets from the Molecular Signatures Database (MSigDB)[21,22] with genes from a GWAS.[23] We downloaded the human gene sets KEGG_ALZHEIMERS_DISEASE and HP_ALZHEIMER_DISEASE from MSigDB (accessed on January 26, 2024). Our final AD risk gene set comprised 243 human genes, which we mapped to 245 mouse orthologs using gprofiler2[19] (**Table S5**).

Prediction of LRT signaling mediators
To find AD-associated LRTs, we used our curated AD risk gene set to filter LRTs targeting any AD risk genes. We used these AD-associated LRTs to generate active signaling networks using the gene regulatory information from the NicheNet v2 prior and the *get_ligand_signaling_path_with_receptor* function from multinichenetr[17] v1.0.0. Our gene regulatory networks included the top two regulators (based on whether they were upstream of the target and downstream of the ligand) in each receiver cell type. Using MultiNicheNet's prior knowledge, which includes ligand-receptor to target signaling path information, we generated weighted and directed igraph objects for AD-associated LRTs using igraph[24] v1.4.2. We determined potential signaling mediators by identifying outgoing nodes from the predicted receptor in every igraph object.

Gene Regulatory Network Construction
To investigate regulatory relationships of signaling mediators in excitatory and inhibitory neurons from 12-month mice, we constructed cell-type- and time-point-specific networks using Passing Attributes between Networks for Data Assimilation (PANDA)[25,26] v1.30.0, which infers regulatory networks through an intersection

of gene expression, protein-protein binding, and known TF-motifs. First, we split our pseudo bulk expression matrices by condition and time point. We combined pseudo bulk gene expression inputs with protein-protein and non-enriched TF motif inputs that we previously generated in Whitlock et al. 2023.[27] We downloaded the protein-protein input for the mouse from StringDB (version 11.0)[28] and TF-motif input compiled from the Catalog of Inferred Sequence Binding Preferences (CIS-BP).[29] This resulted in 4 gene regulatory networks for each time point and condition pairing (12mWT & 12mAD for excitatory neurons, 12mWT & 12mAD for inhibitory neurons). Here, network edge weights represent the strength of interactions between nodes, where nodes represent coexpressed genes, proteins, or pairs of TFs and genes. More positive weights correspond to higher confidence interactions in each cell-type- and time-point-specific network, and negative weights indicate a lack of evidence.[26] We performed the gene regulatory network construction in R v4.2.1 using docker (https://hub.docker.com/repository/docker/jordanwhitlock/setbp1_manuscript_panda_1.0.1/general).[30]

TF Activity Analysis of Signaling Mediators

To determine differences in TF activity between conditions for receiver cell types, we again performed a differential gene expression analysis using previously pseudo bulked count matrices with DESeq2 (see *Differential Gene Expression Analysis* section for details). However, to preserve the Wald test statistic (*stat*) needed for our TF activity analyses, we used the method's original shrinkage estimator (type = normal). We combined the Wald test statistic (*stat*) across time points for both receivers. We downloaded the CollecTRI prior, which includes TF information such as their direction of regulation on their targets (accessed in February 2024).[31] We calculated TF activity scores for all signaling mediators in receivers with a minimum of 5 targets using the Multivariate Linear Model (*run_mlm* function decoupleR v2.9.1).[32] The Multivariate Linear Model provided t-values that represent regulator activity.[32] Positive values represent an active TF, and negative values represent an inactive TF in 3xTg-AD compared to WT.

Differential Gene Targeting of Signaling Mediators

We calculated differential gene targeting scores[33] to determine whether signaling mediators in inhibitory and excitatory neurons were differentially targeted in 3xTg-AD compared to WT hippocampus using the gene regulatory networks we generated using PANDA. We quantified the in-degree edge weights for each signaling mediator before determining the differential targeting score between conditions by calculating the difference in targeting scores. Positive differential targeting scores indicated increased targeting in 3xTg-AD compared to the WT hippocampus. Similarly to our previous work, we calculated quartiles of all gene targeting scores by receivers to determine whether mediators were among the highest or lowest differentially targeted genes.[27]

4. Merkel, D. Docker: lightweight Linux containers for consistent development and deployment. *Linux J.*

5. Young, M. D. & Behjati, S. SoupX removes ambient RNA contamination from droplet-based single-cell

RNA sequencing data. *Gigascience* **9**, (2020).

19. Kolberg, L., Raudvere, U., Kuzmin, I., Vilo, J. & Peterson, H. gprofiler2 -- an R package for gene list functional enrichment analysis and namespace conversion toolset g:Profiler. *F1000Res.* **9**, (2020).
20. Wilk, E. J. *et al.* Prioritized polycystic kidney disease drug targets and repurposing candidates from pre-cystic and cystic mouse Pkd2 model gene expression reversion. *Mol. Med.* **29**, 67 (2023).
21. Subramanian, A. *et al.* Gene set enrichment analysis: a knowledge-based approach for interpreting genome-wide expression profiles. *Proc. Natl. Acad. Sci. U. S. A.* **102**, 15545–15550 (2005).
22. Liberzon, A. *et al.* Molecular signatures database (MSigDB) 3.0. *Bioinformatics* **27**, 1739–1740 (2011).
23. Bellenguez, C. *et al.* New insights into the genetic etiology of Alzheimer's disease and related dementias. *Nat. Genet.* **54**, 412–436 (2022).
24. Csárdi, G. *et al. Igraph for R: R Interface of the Igraph Library for Graph Theory and Network Analysis*. (Zenodo, 2023). doi:10.5281/ZENODO.7682609.
25. Glass, K., Huttenhower, C., Quackenbush, J. & Yuan, G.-C. Passing messages between biological networks to refine predicted interactions. *PLoS One* **8**, e64832 (2013).
26. Schlauch, D., Paulson, J. N., Young, A., Glass, K. & Quackenbush, J. Estimating gene regulatory networks with pandaR. *Bioinformatics* **33**, 2232–2234 (2017).
27. Whitlock, J. H. *et al.* Cell-type-specific gene expression and regulation in the cerebral cortex and kidney of atypical Setbp1S858R Schinzel Giedion Syndrome mice. *J. Cell. Mol. Med.* **27**, 3565–3577 (2023).
28. Szklarczyk, D. *et al.* STRING v11: protein-protein association networks with increased coverage, supporting functional discovery in genome-wide experimental datasets. *Nucleic Acids Res.* **47**, D607–D613 (2019).
29. Sonawane, A. R. *et al.* Understanding Tissue-Specific Gene Regulation. *Cell Rep.* **21**, 1077–1088 (2017).
30. Creators Jordan H. Whitlock1 Tabea M. Soelter1 Timothy C. Howton1 Elizabeth J. Wilk1 Vishal H. Oza1 Brittany N. Lasseigne1 Show affiliations 1. The University of Alabama at Birmingham, Heersink School of Medicine. *Docker Images for Setbp1 (p.S858R) Variant Analysis*. doi:10.5281/zenodo.8190923.
31. Müller-Dott, S. *et al.* Expanding the coverage of regulons from high-confidence prior knowledge for accurate estimation of transcription factor activities. *bioRxiv* 2023.03.30.534849 (2023) doi:10.1101/2023.03.30.534849.
32. Badia-I-Mompel, P. *et al.* decoupleR: ensemble of computational methods to infer biological activities from

omics data. *Bioinform Adv* **2**, vbac016 (2022).

33. Weighill, D. *et al.* Gene Targeting in Disease Networks. *Front. Genet.* **12**, 649942 (2021).
