## Supplementary File 1 for "Evaluation of altered cell-cell communication between glia and neurons in the hippocampus of 3xTg-AD mice at two time points"

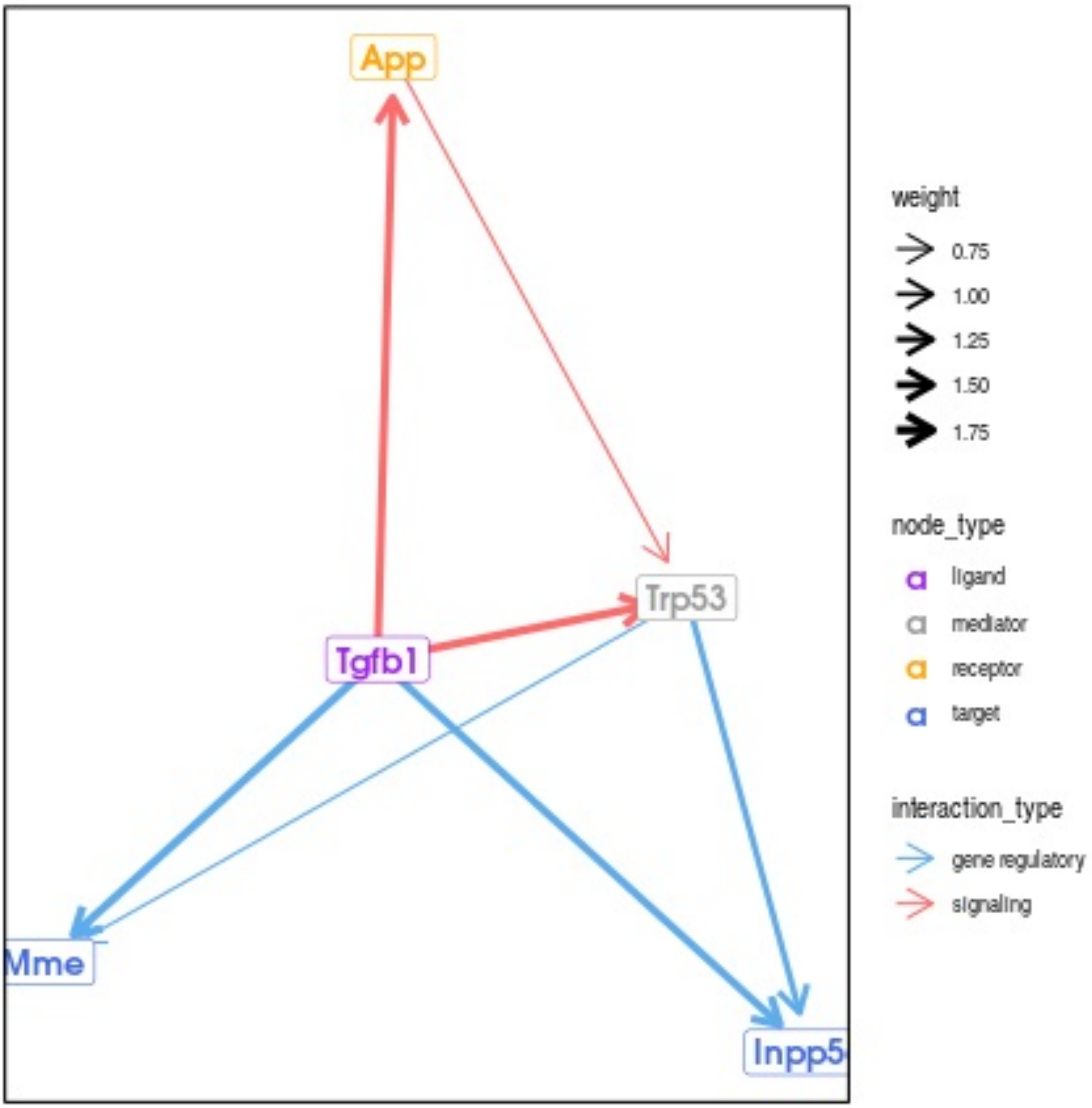
App
Trp53
Tgfb1
Mme
Inpp5
weight
0.75
1.00
1.25
1.50
1.75
node_type
ligand
mediator
receptor
target
interaction_type
gene regulatory
signaling

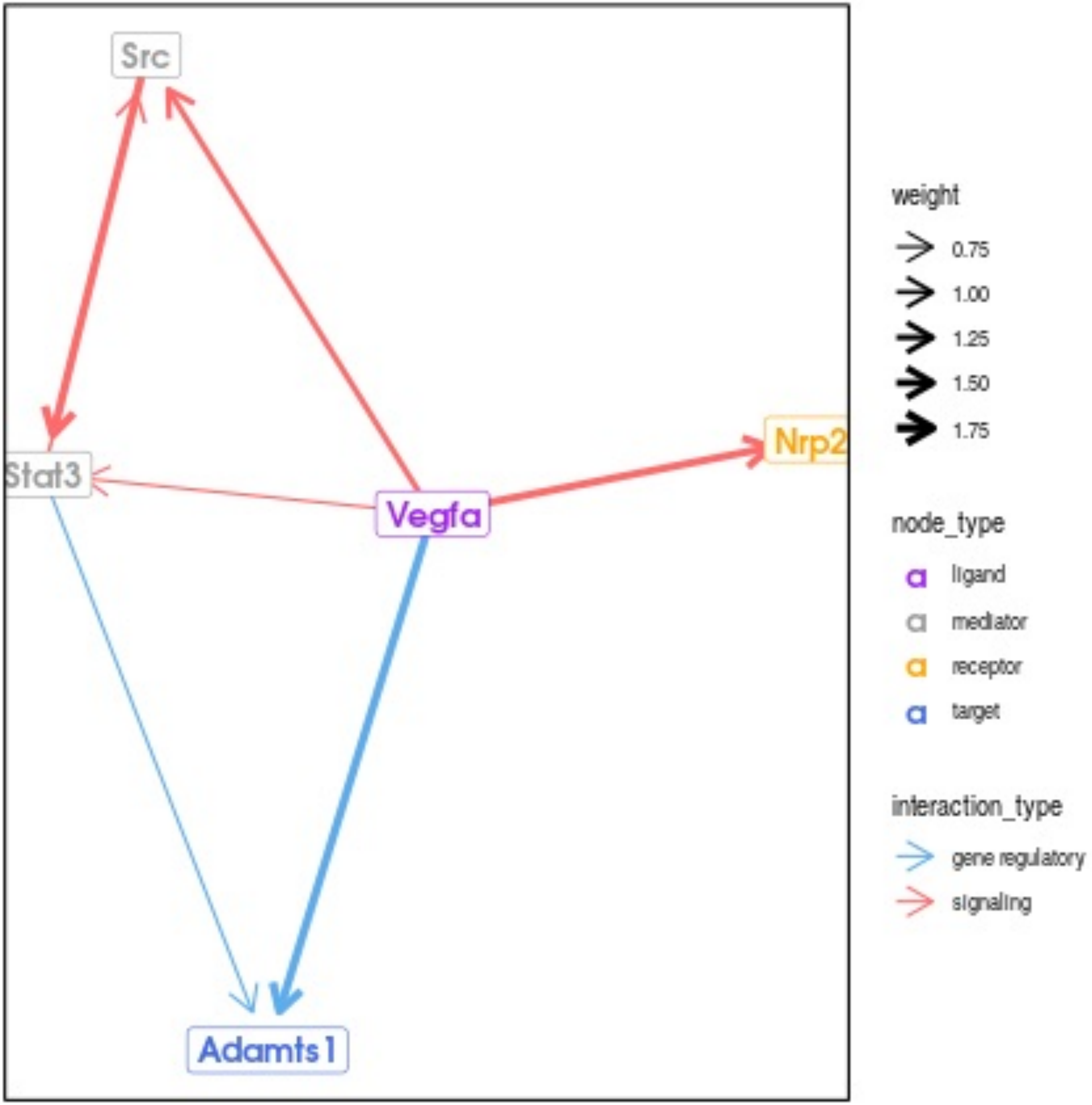

Src
Stat3
Vegfa
Nrp2
Adamts1
weight
0.75
1.00
1.25
1.50
1.75
node_type
ligand
mediator
receptor
target
interaction_type
gene regulatory
signaling

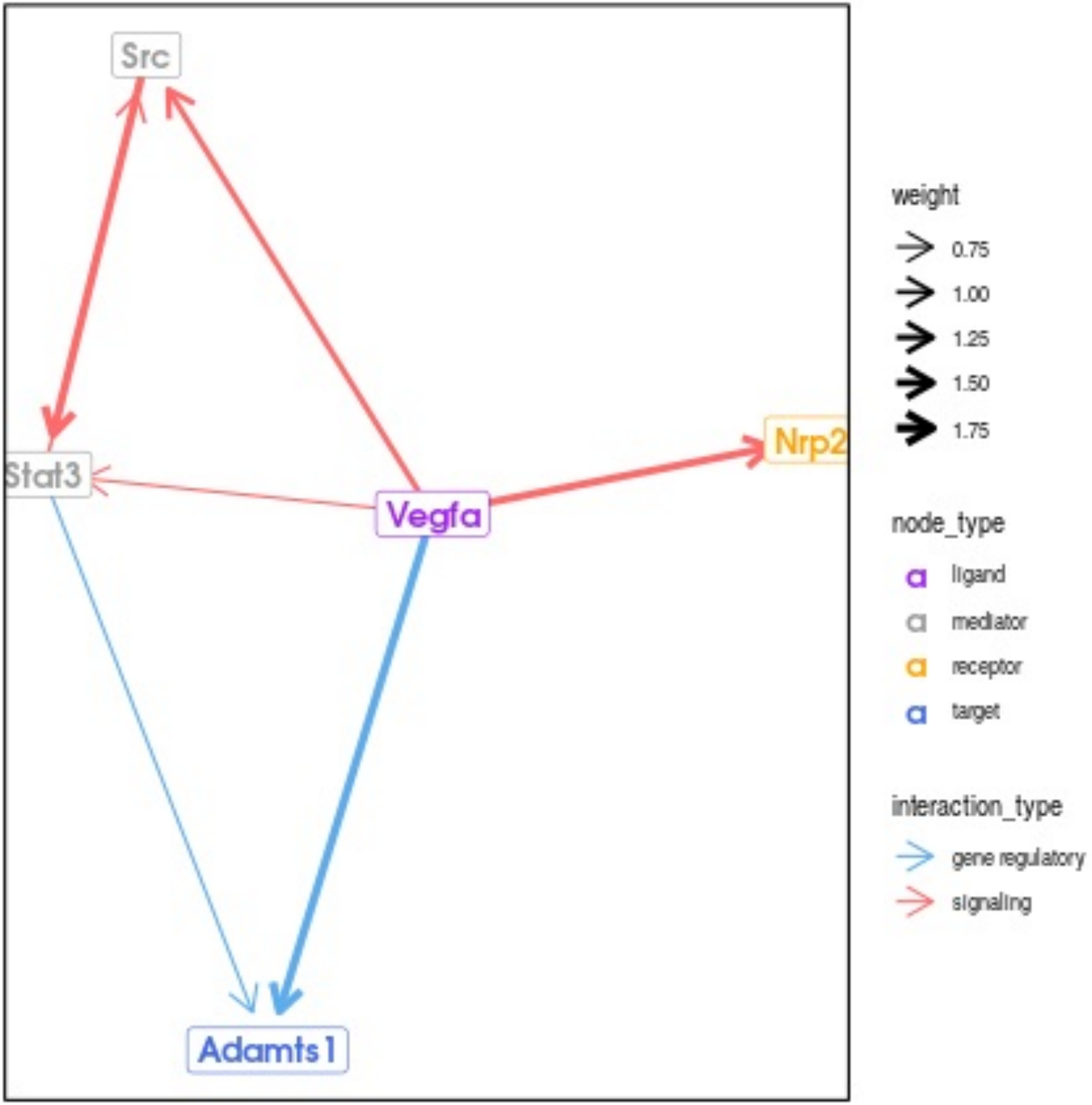
Src
Stat3
Vegfa
Nrp2
Adamts1
weight
0.75
1.00
1.25
1.50
1.75
node_type
ligand
mediator
receptor
target
interaction_type
gene regulatory
signaling

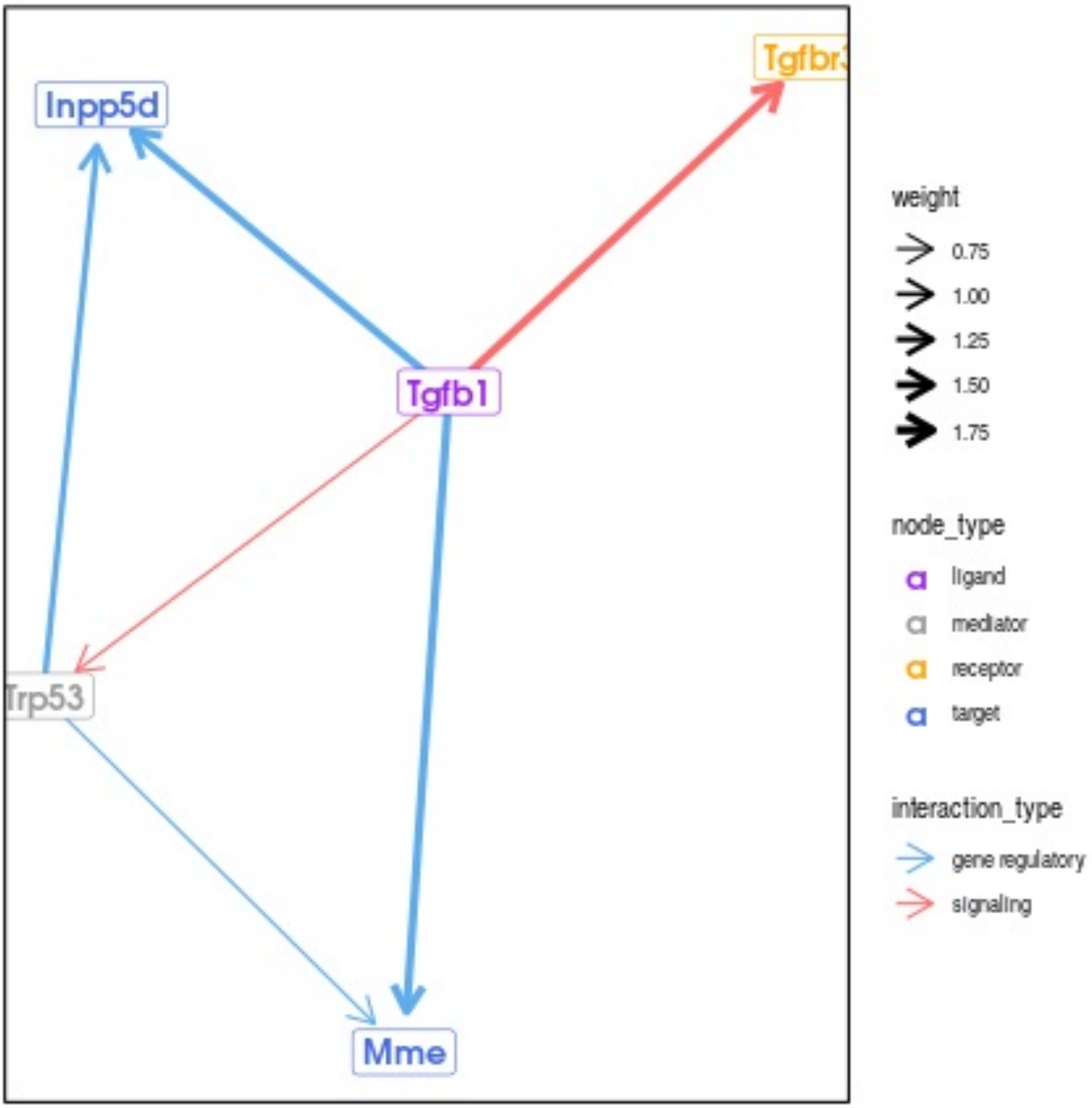
Inpp5d
Tgfbr3
Tgfb1
Trp53
Mme
weight
0.75
1.00
1.25
1.50
1.75
node_type
ligand
mediator
receptor
target
interaction_type
gene regulatory
signaling

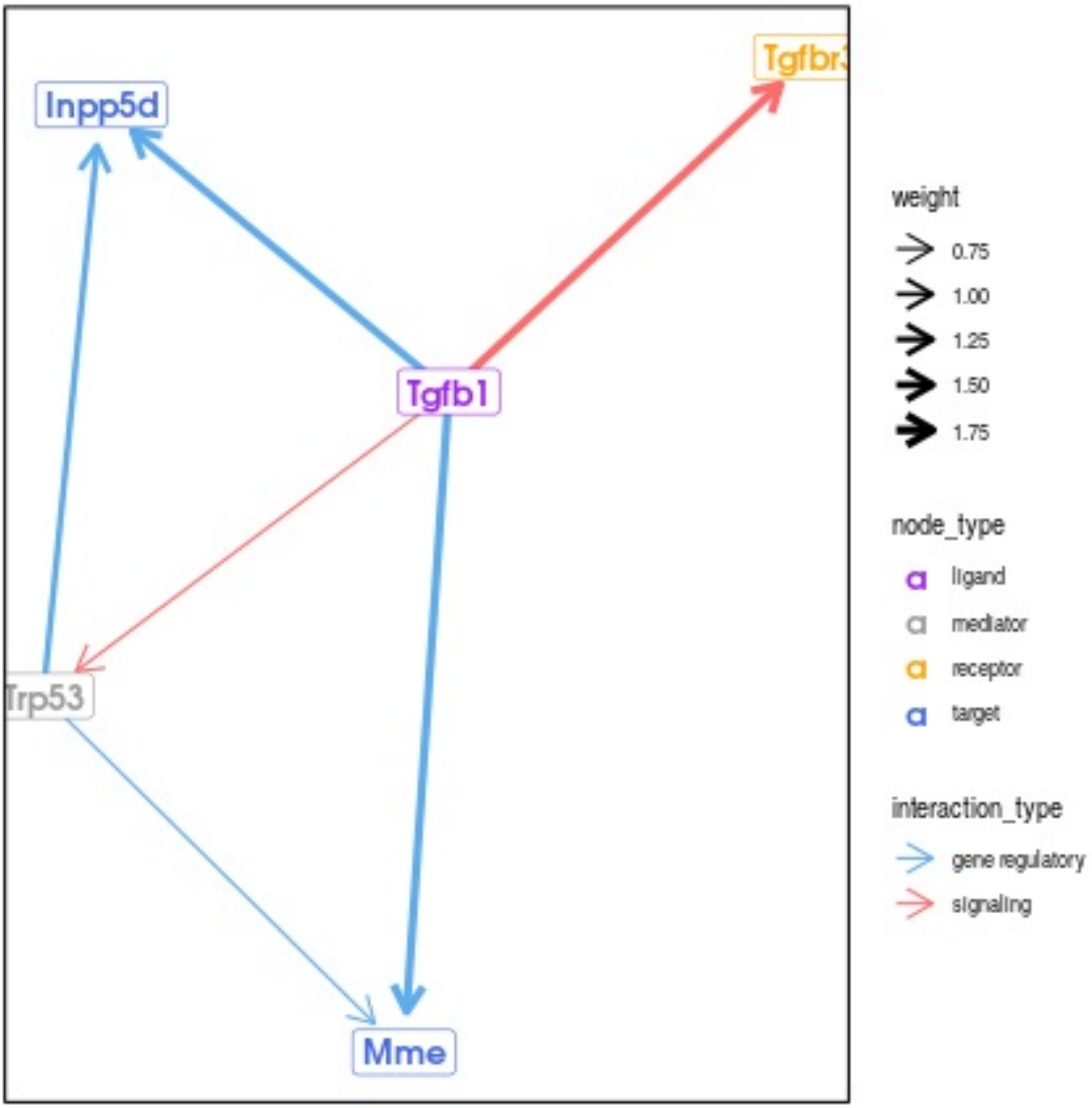
Inpp5d
Tgfbr3
Tgfb1
Trp53
Mme
weight
0.75
1.00
1.25
1.50
1.75
node_type
ligand
mediator
receptor
target
interaction_type
gene regulatory
signaling

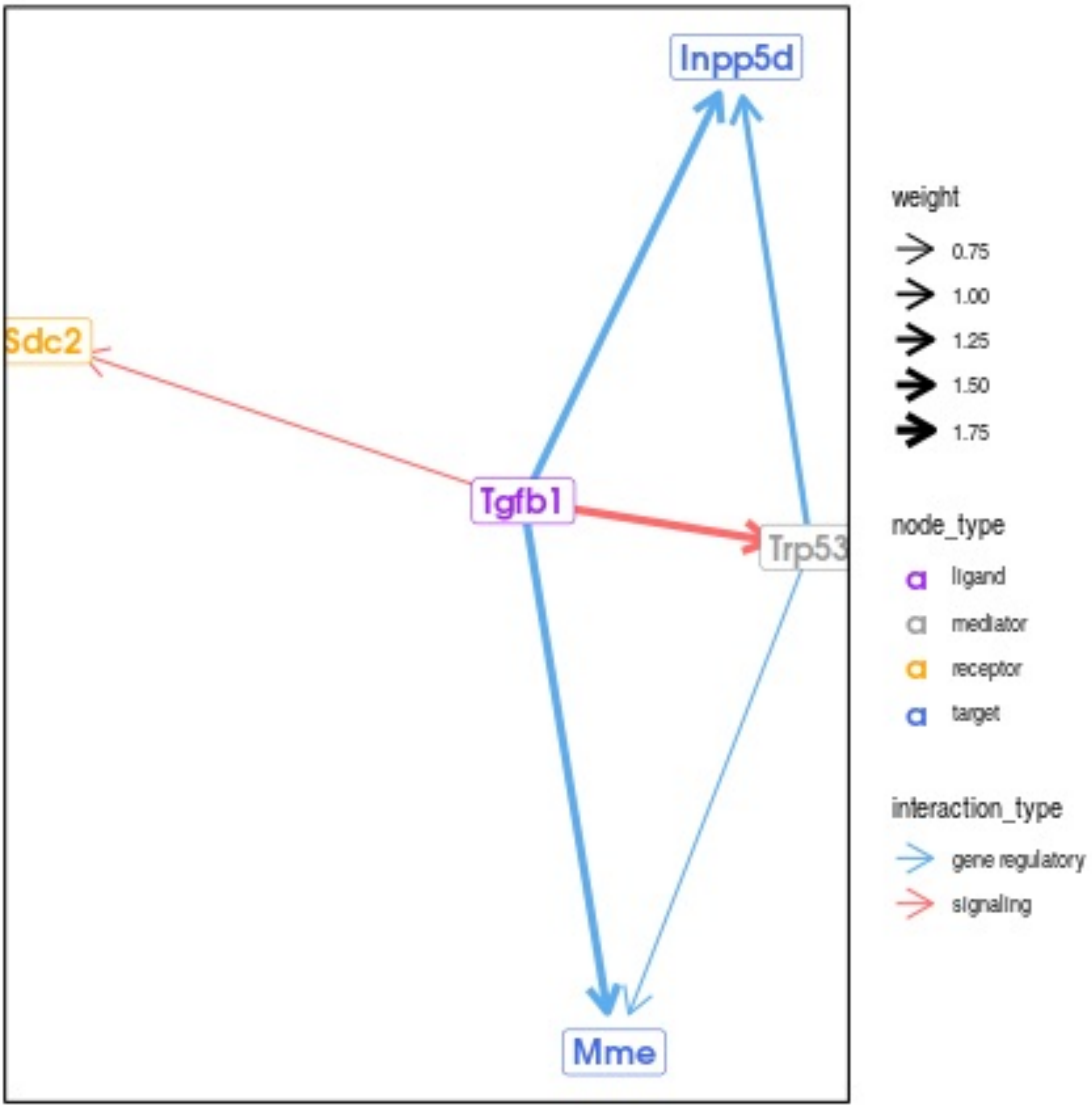
Inpp5d
Sdc2
Tgfb1
Trp53
Mme
weight
0.75
1.00
1.25
1.50
1.75
node_type
ligand
mediator
receptor
target
interaction_type
gene regulatory
signaling

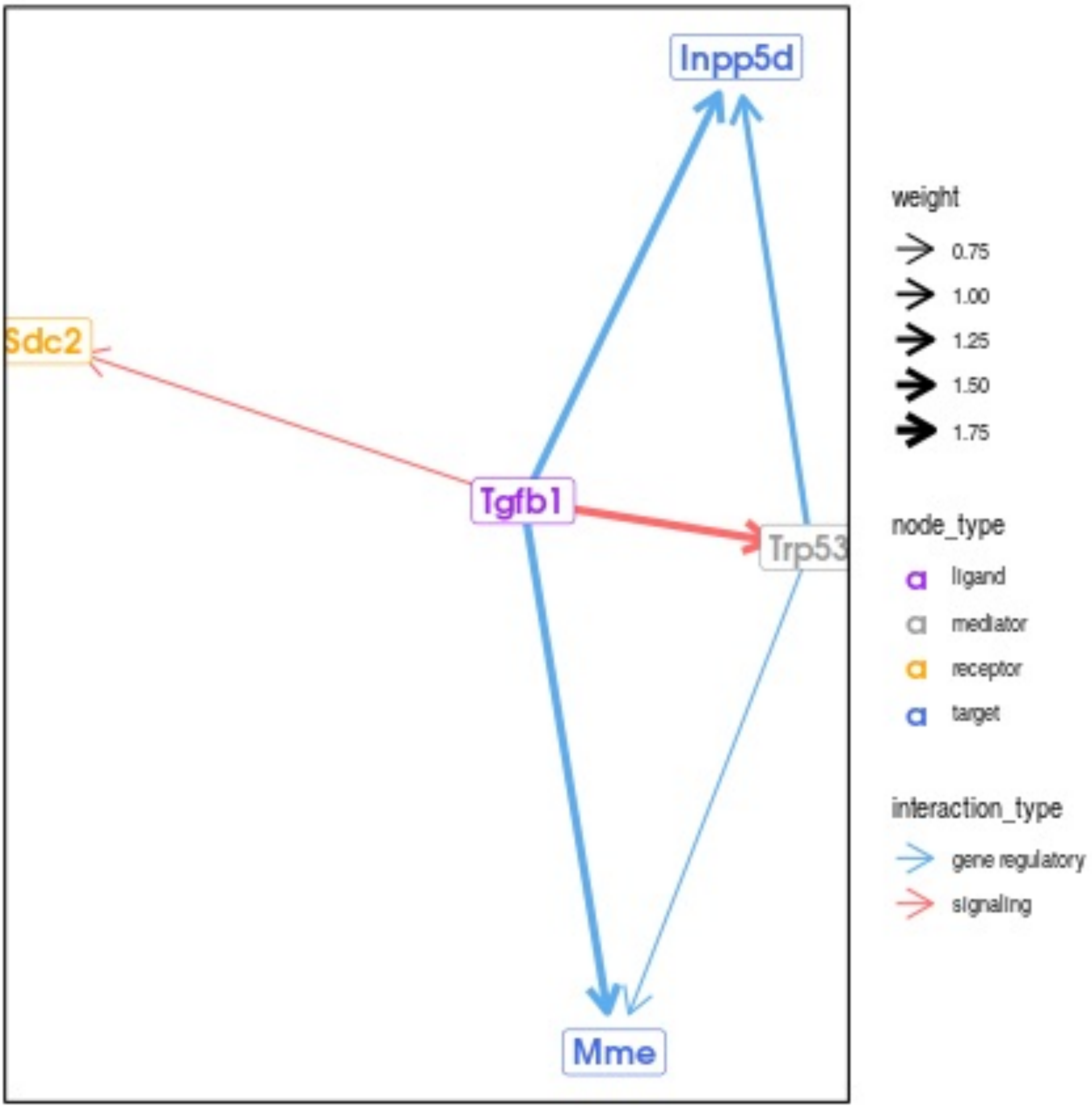
Inpp5d
Sdc2
Tgfb1
Trp53
Mme
weight
0.75
1.00
1.25
1.50
1.75
node_type
ligand
mediator
receptor
target
interaction_type
gene regulatory
signaling

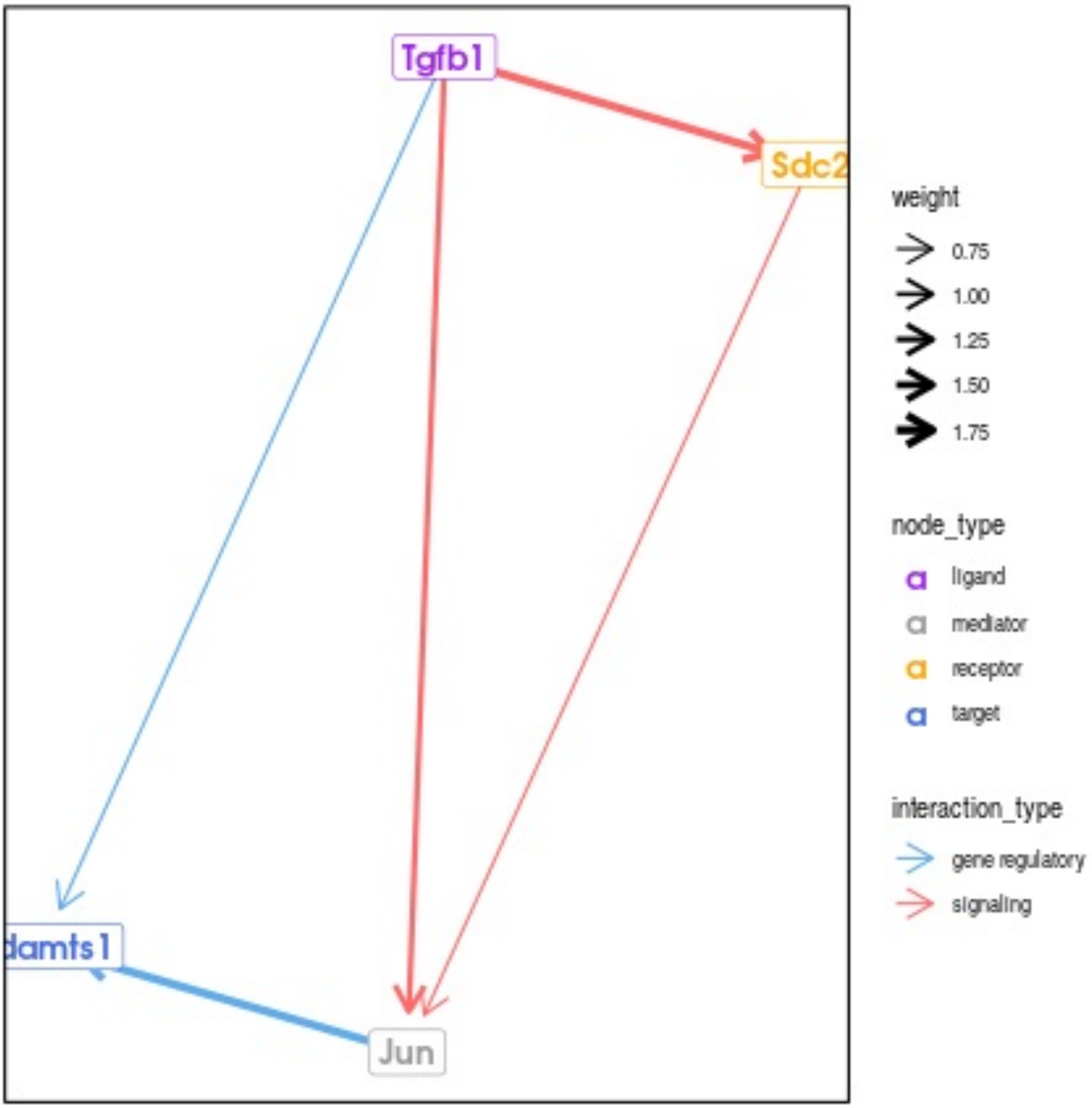

Tgfb1
Sdc2
damts1
Jun
weight
0.75
1.00
1.25
1.50
1.75
node_type
a
ligand
a
mediator
a
receptor
a
target
interaction_type
gene regulatory
signaling

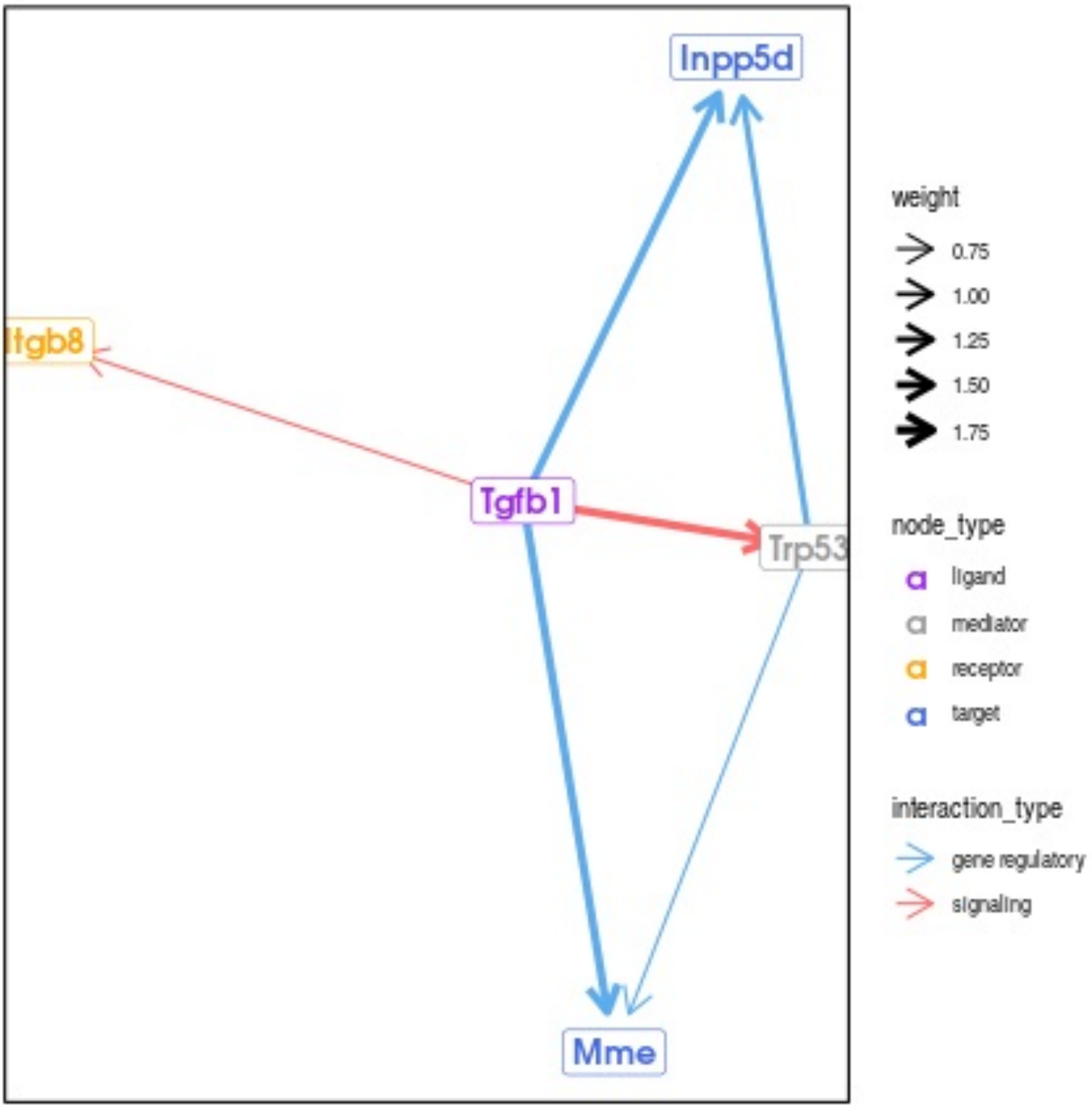
Inpp5d
Itgb8
Tgfb1
Trp53
Mme
weight
0.75
1.00
1.25
1.50
1.75
node_type
ligand
mediator
receptor
target
interaction_type
gene regulatory
signaling

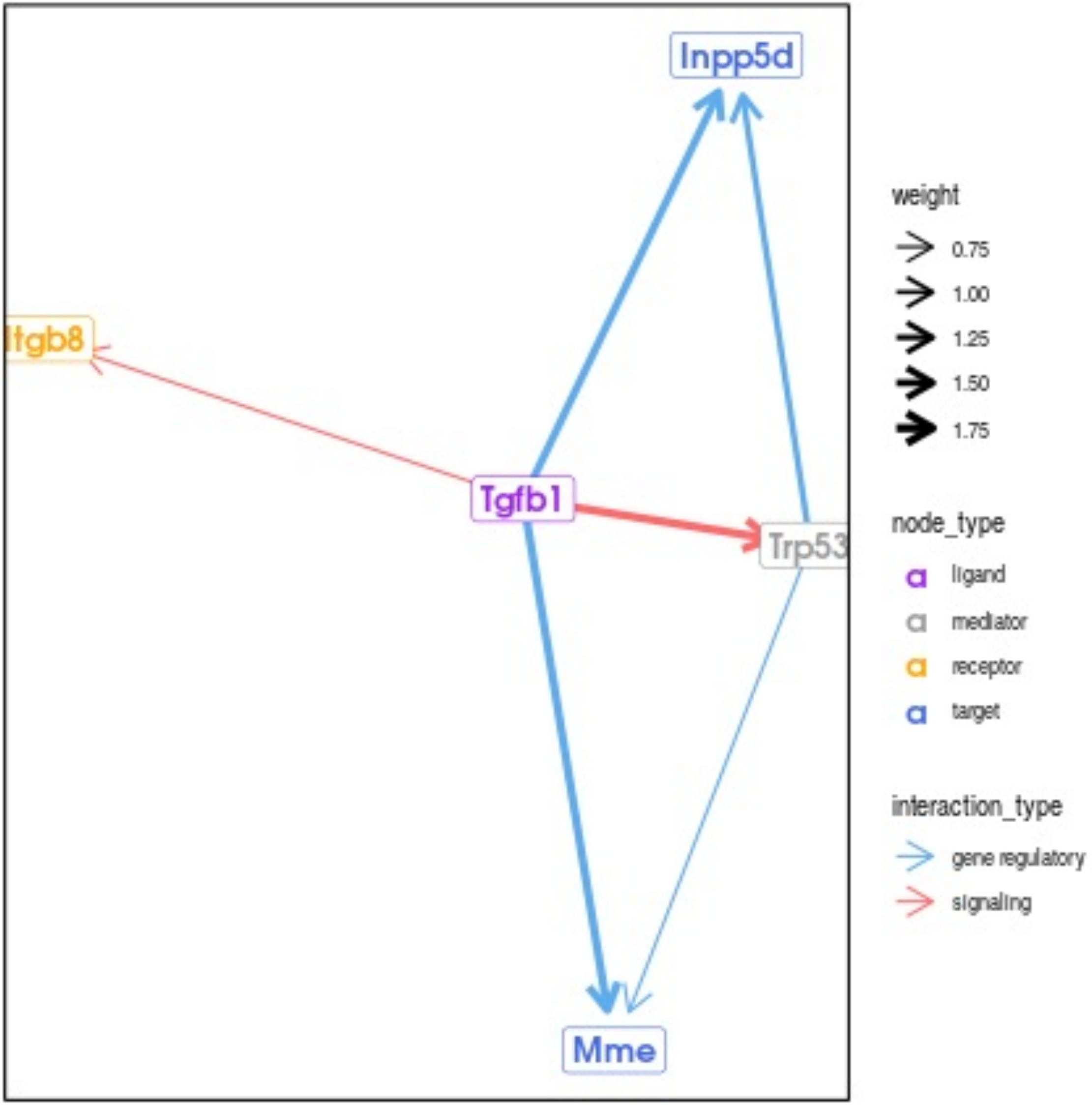
Inpp5d
Itgb8
Tgfb1
Trp53
Mme
weight
0.75
1.00
1.25
1.50
1.75
node_type
ligand
mediator
receptor
target
interaction_type
gene regulatory
signaling

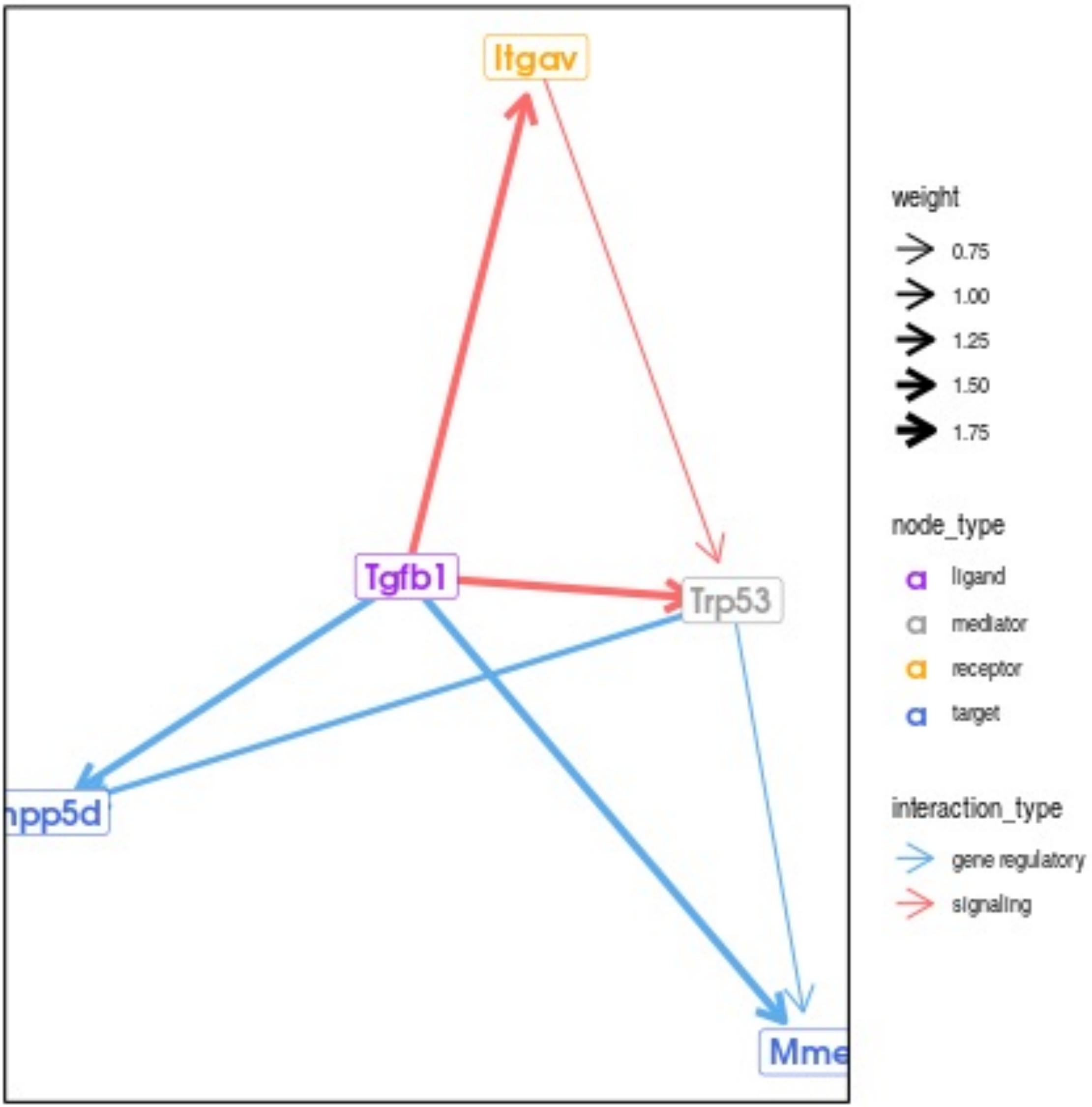

Itgav
Tgfb1
Trp53
npp5d
Mme
weight
0.75
1.00
1.25
1.50
1.75
node_type
ligand
mediator
receptor
target
interaction_type
gene regulatory
signaling

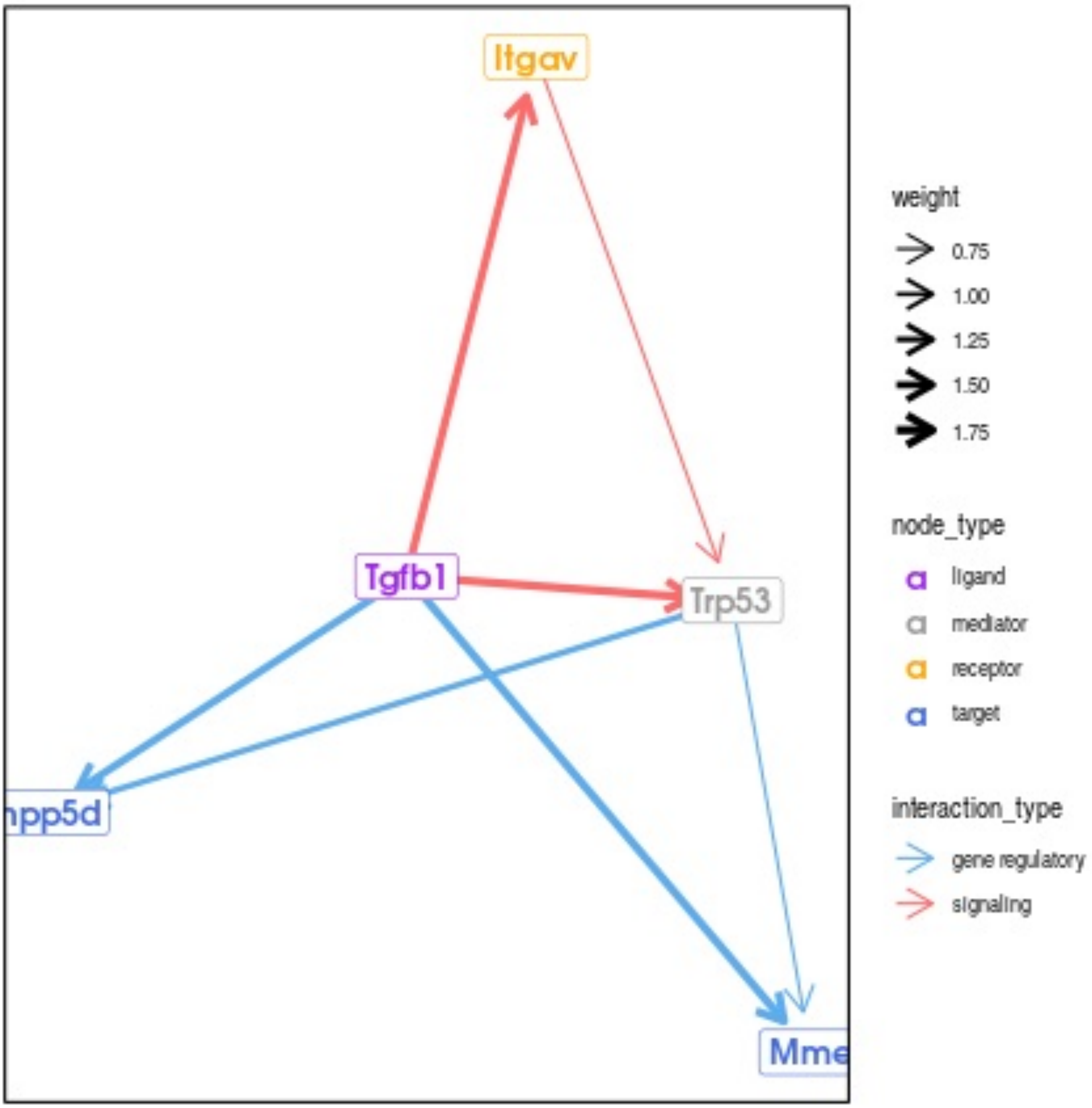
Itgav
Tgfb1
Trp53
npp5d
Mme
weight
0.75
1.00
1.25
1.50
1.75
node_type
ligand
mediator
receptor
target
interaction_type
gene regulatory
signaling

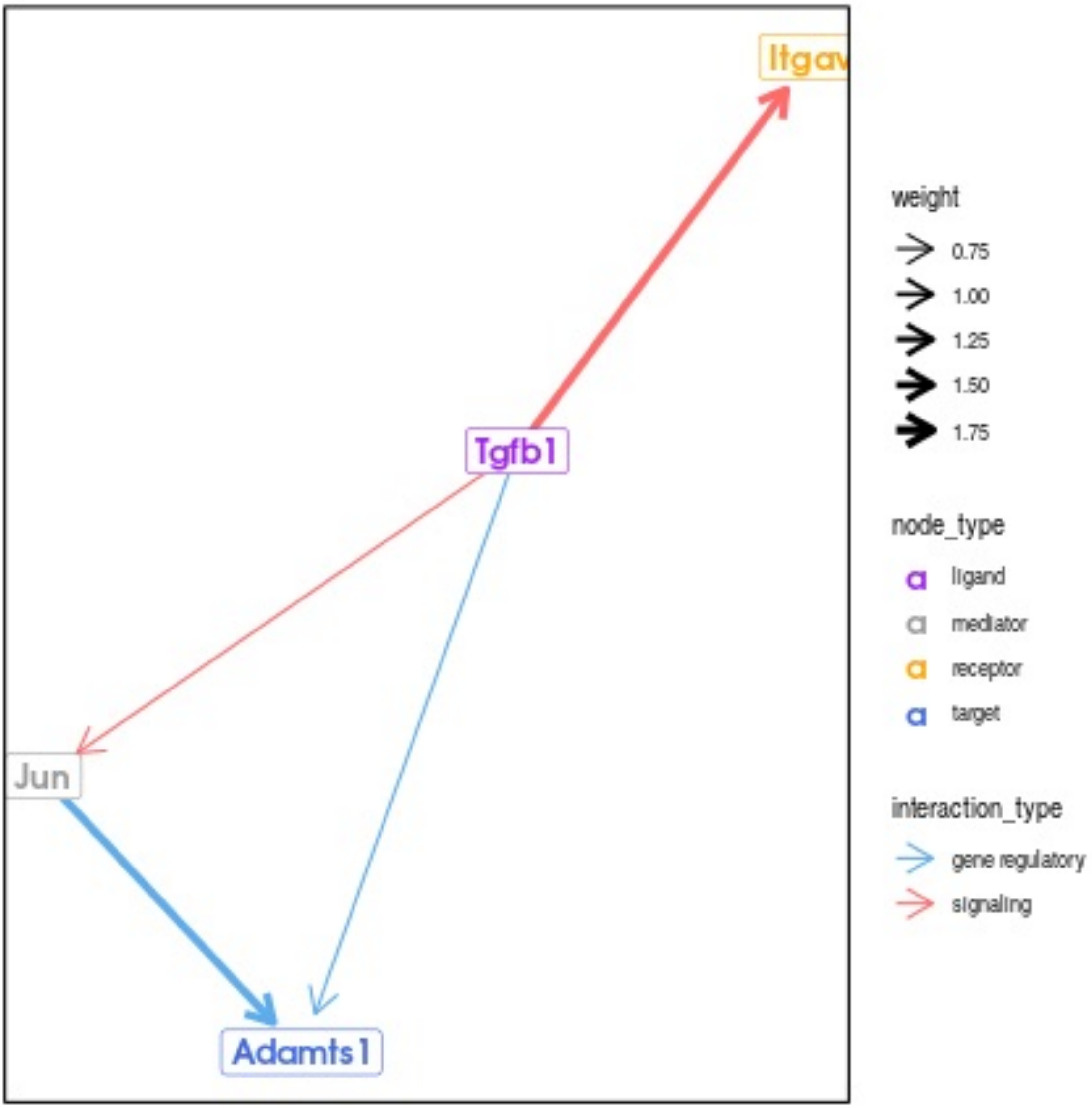
Itgav
Tgfb1
Jun
Adamts1
weight
0.75
1.00
1.25
1.50
1.75
node_type
ligand
mediator
receptor
target
interaction_type
gene regulatory
signaling

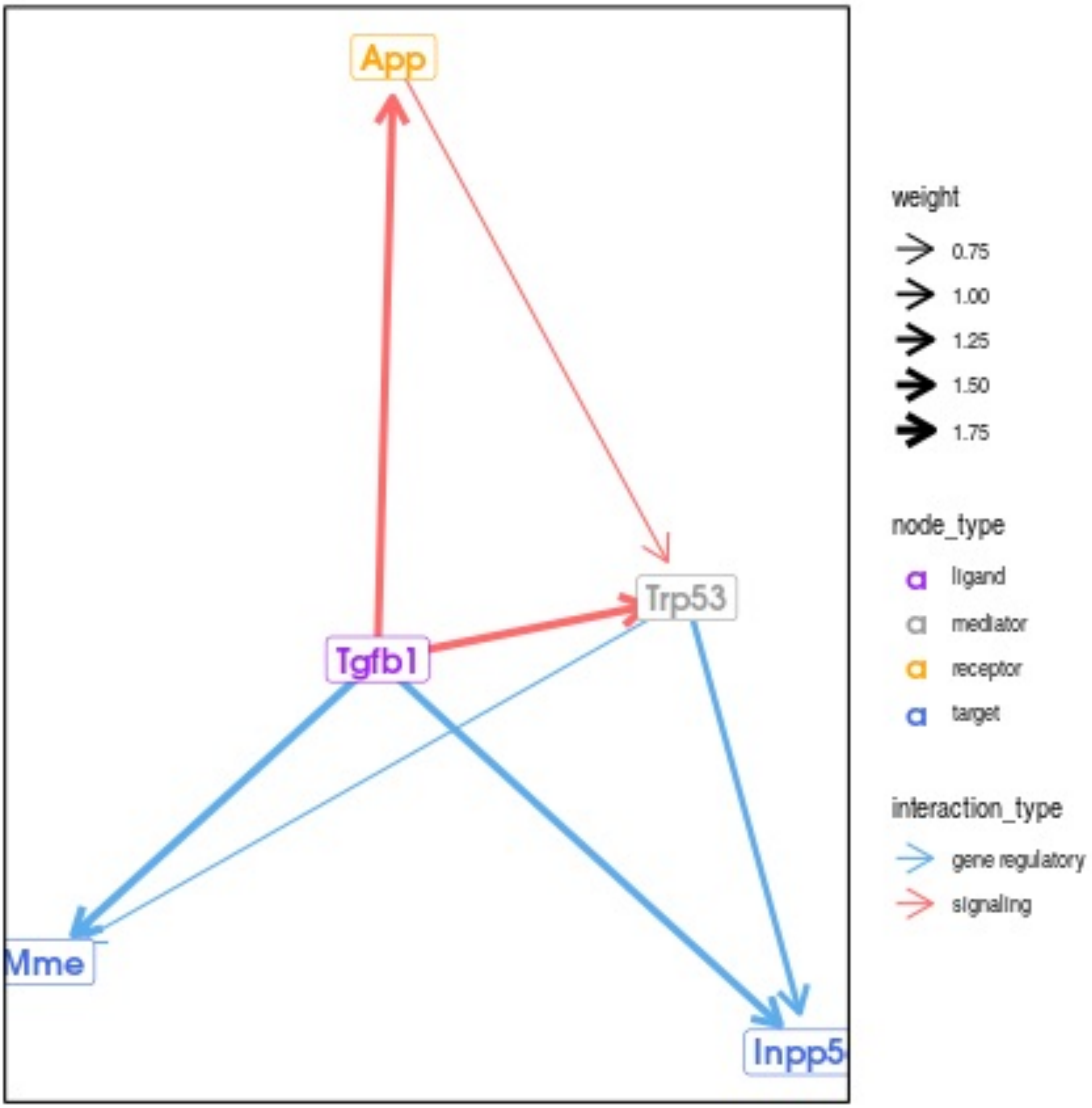

App
Trp53
Tgfb1
Mme
Inpp5
weight
0.75
1.00
1.25
1.50
1.75
node_type
ligand
mediator
receptor
target
interaction_type
gene regulatory
signaling

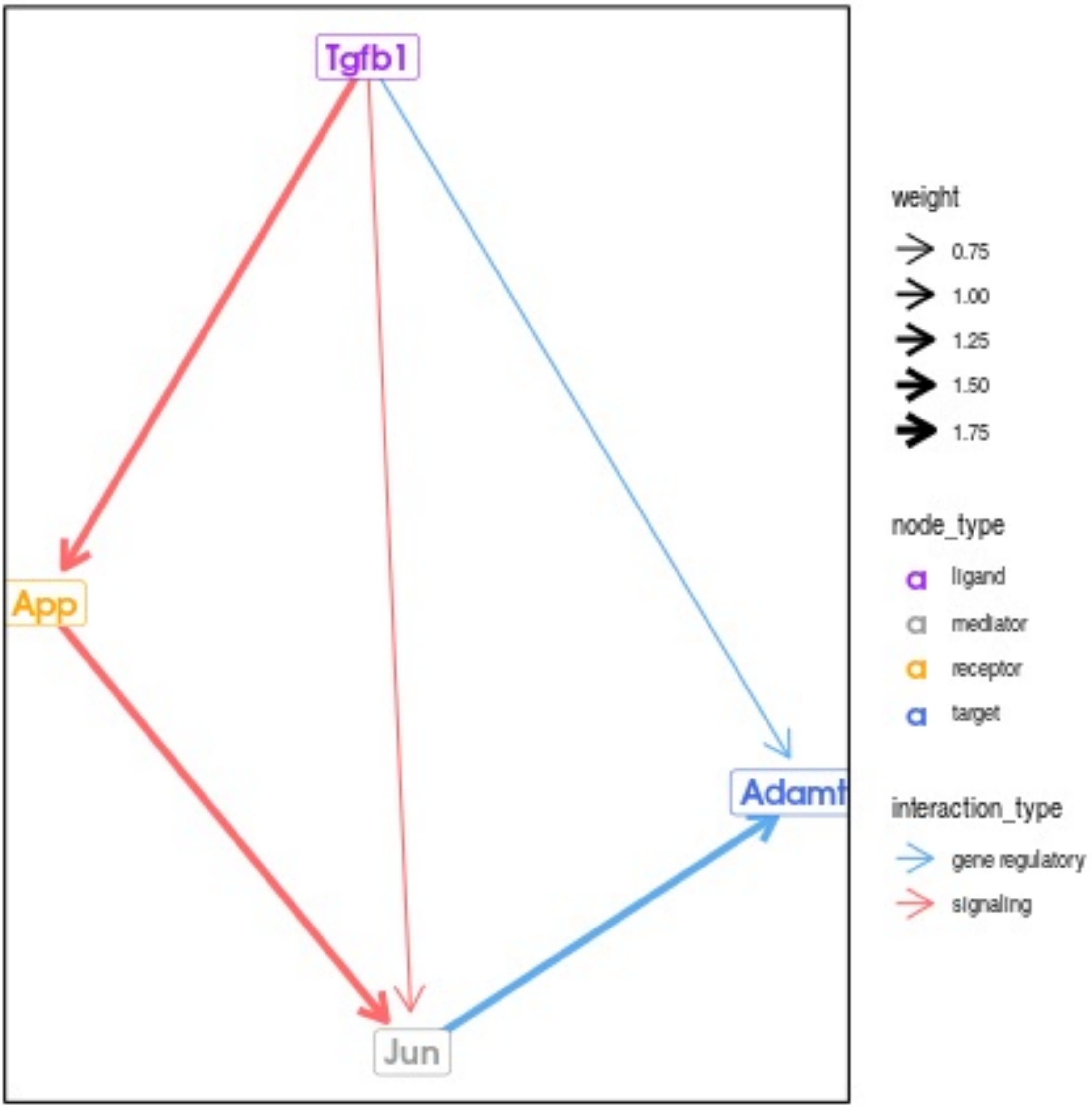
Tgfb1
App
Jun
Adamt
weight
0.75
1.00
1.25
1.50
1.75
node_type
a ligand
a mediator
a receptor
a target
interaction_type
gene regulatory
signaling

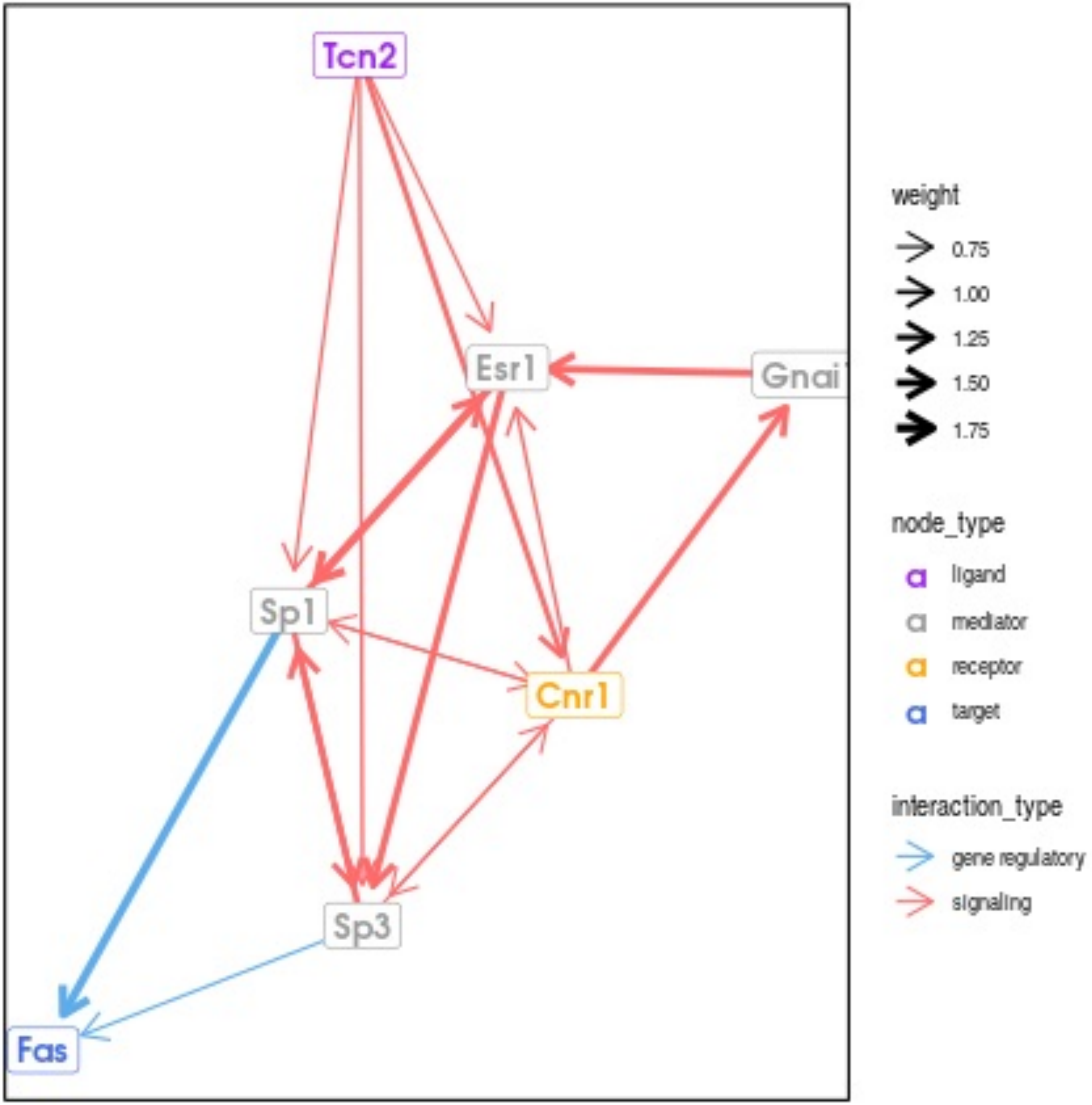
Tcn2
Esr1
Gnai
Sp1
Cnr1
Sp3
Fas
weight
0.75
1.00
1.25
1.50
1.75
node_type
ligand
mediator
receptor
target
interaction_type
gene regulatory
signaling

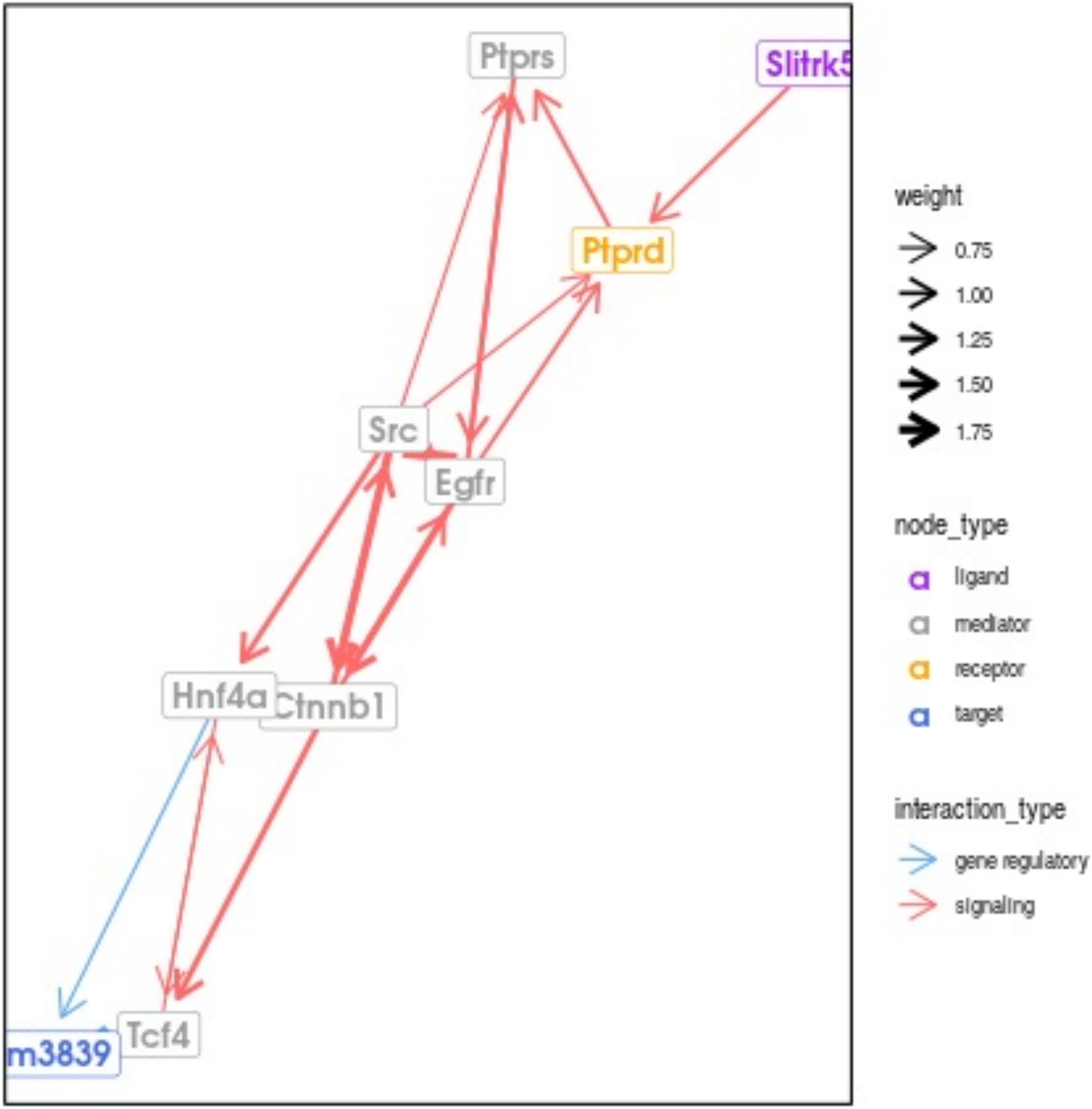

Ptprs
Slitrk5
Ptprd
Src
Egfr
Hnf4a
Ctnnb1
m3839
Tcf4
weight
0.75
1.00
1.25
1.50
1.75
node_type
ligand
mediator
receptor
target
interaction_type
gene regulatory
signaling

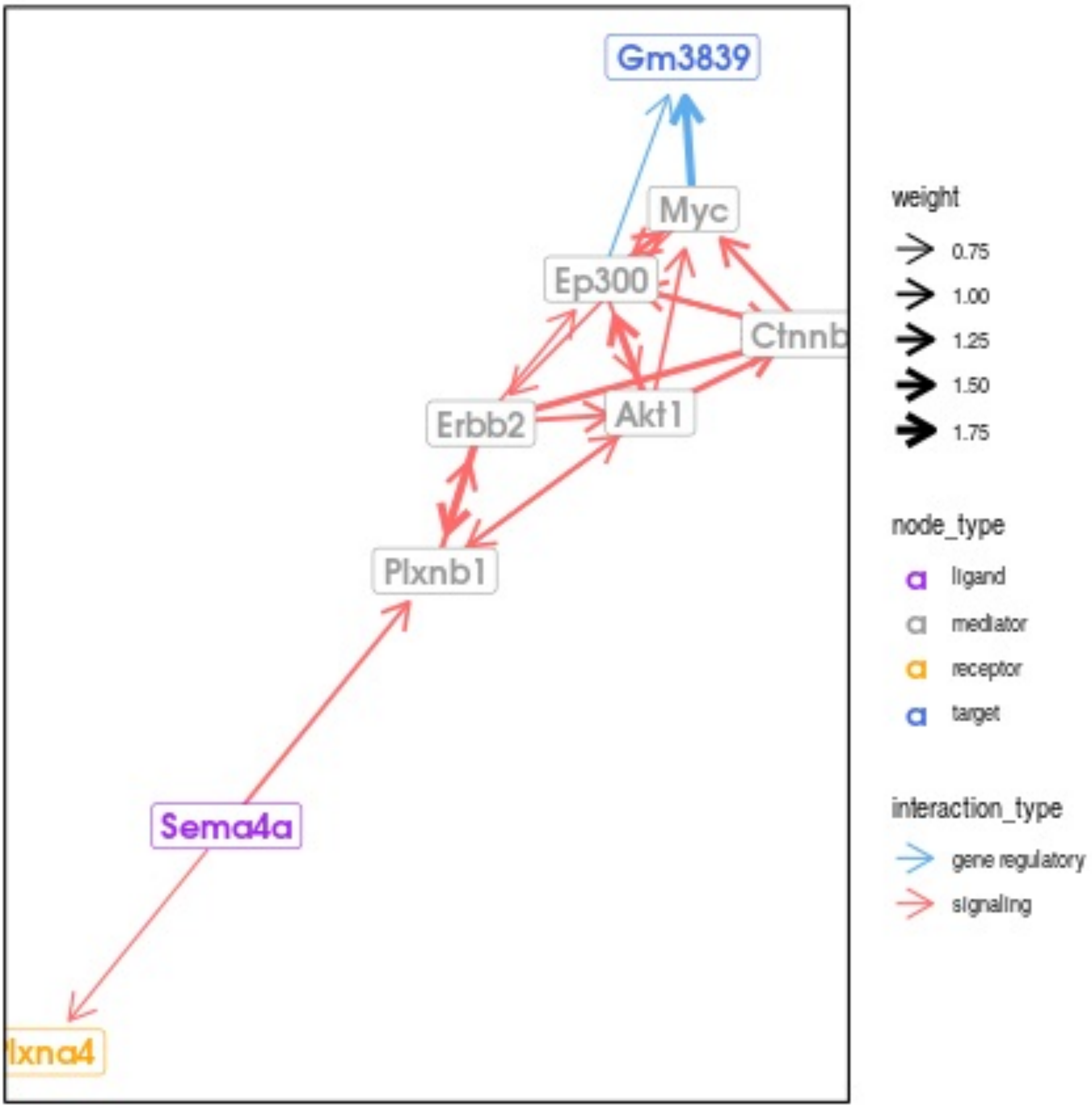

Gm3839
Myc
Ep300
Ctnnb
Erbb2
Akt1
Plxnb1
Sema4a
lxna4
weight
0.75
1.00
1.25
1.50
1.75
node_type
ligand
mediator
receptor
target
interaction_type
gene regulatory
signaling

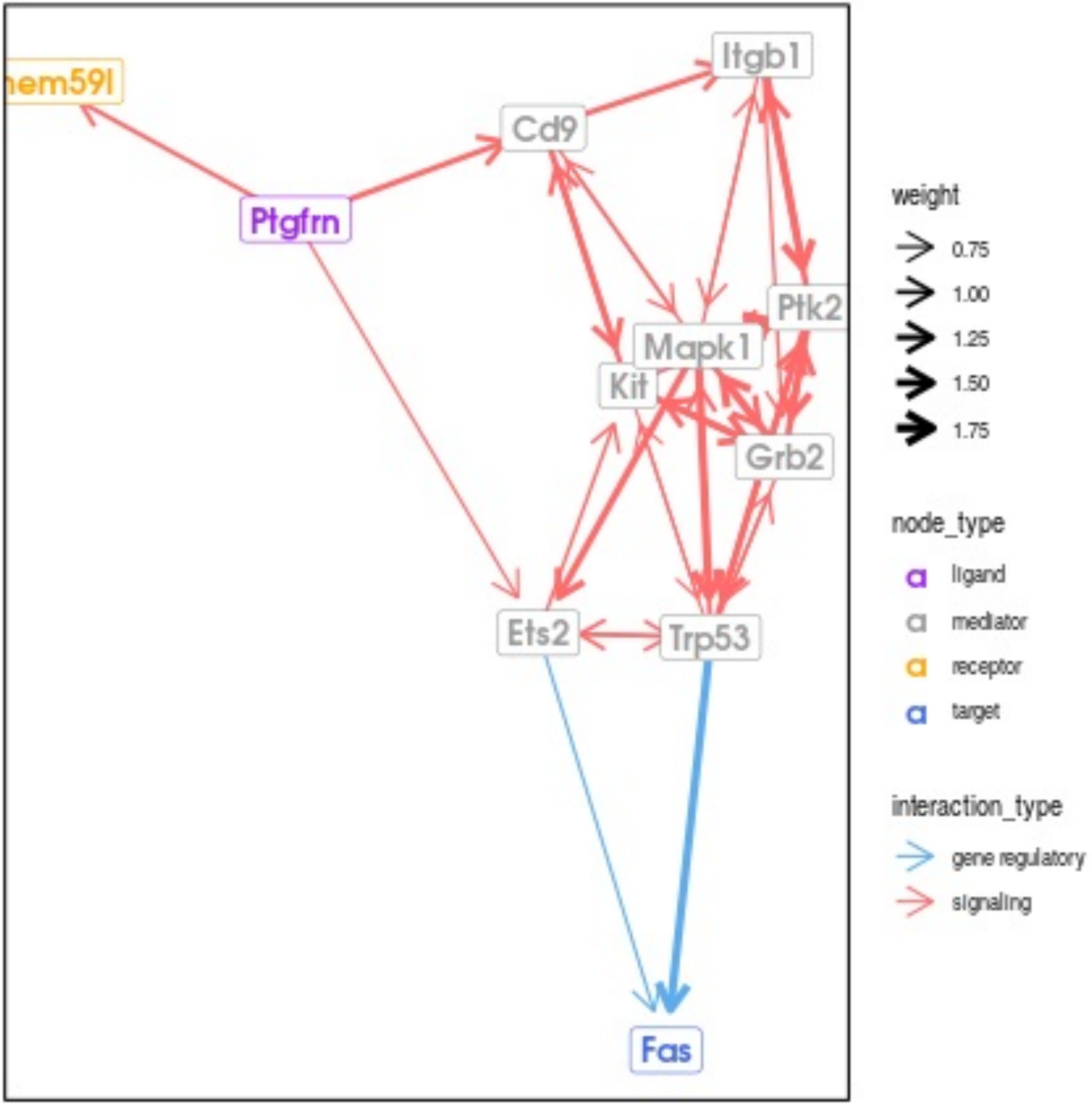

hem59l
Itgb1
Cd9
Ptgfrn
Ptk2
Mapk1
Kit
Grb2
Ets2
Trp53
Fas
weight
0.75
1.00
1.25
1.50
1.75
node_type
ligand
mediator
receptor
target
interaction_type
gene regulatory
signaling

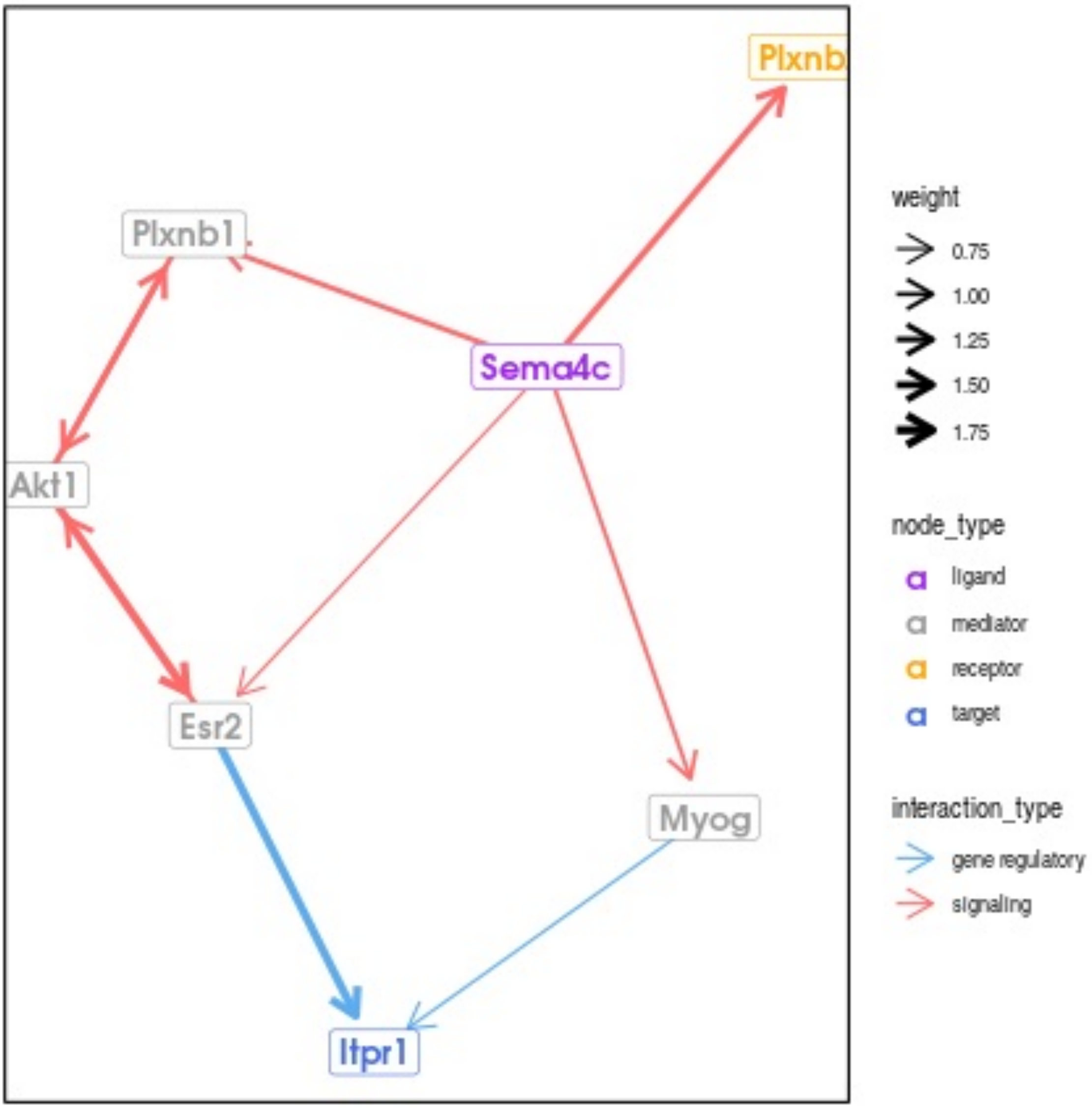

Plxnb
Plxnb1
Sema4c
Akt1
Esr2
Myog
Itpr1
weight
0.75
1.00
1.25
1.50
1.75
node_type
ligand
mediator
receptor
target
interaction_type
gene regulatory
signaling

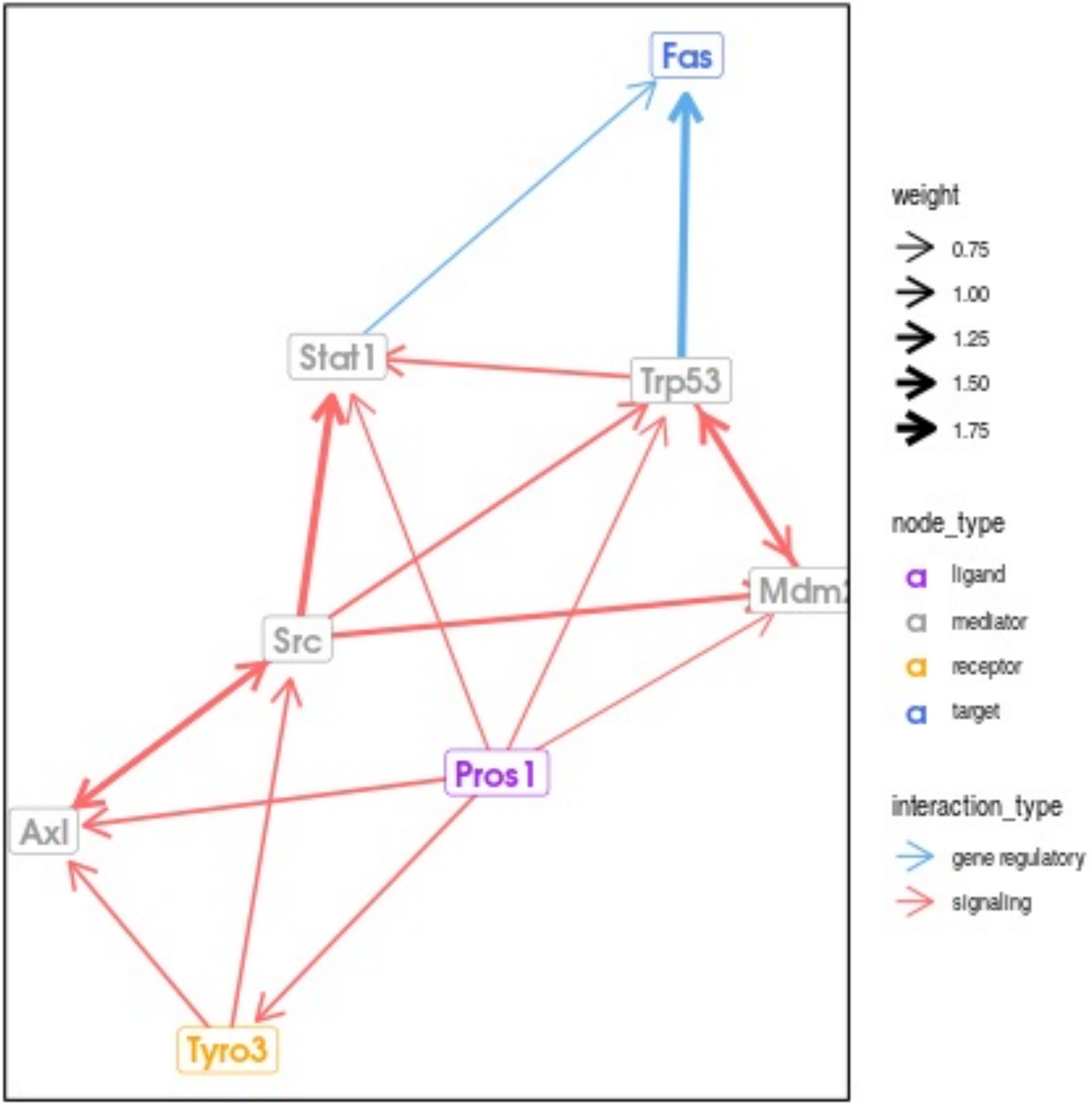
Fas
Stat1
Trp53
Mdm2
Src
Pros1
Axl
Tyro3
weight
0.75
1.00
1.25
1.50
1.75
node_type
ligand
mediator
receptor
target
interaction_type
gene regulatory
signaling

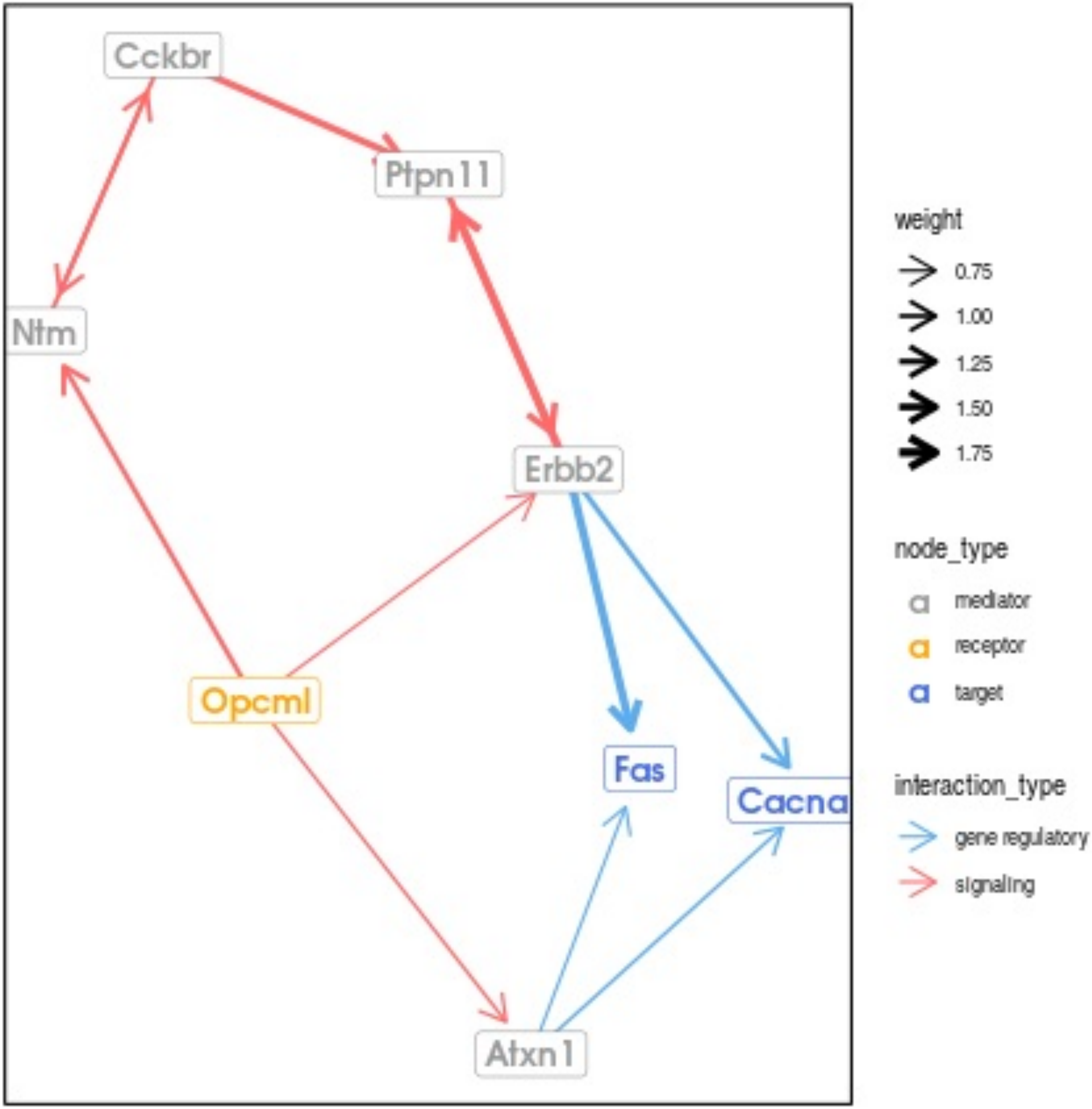

Cckbr
Ptpn11
Ntm
Erbb2
Opcml
Fas
Cacna
Atxn1
weight
0.75
1.00
1.25
1.50
1.75
node_type
mediator
receptor
target
interaction_type
gene regulatory
signaling

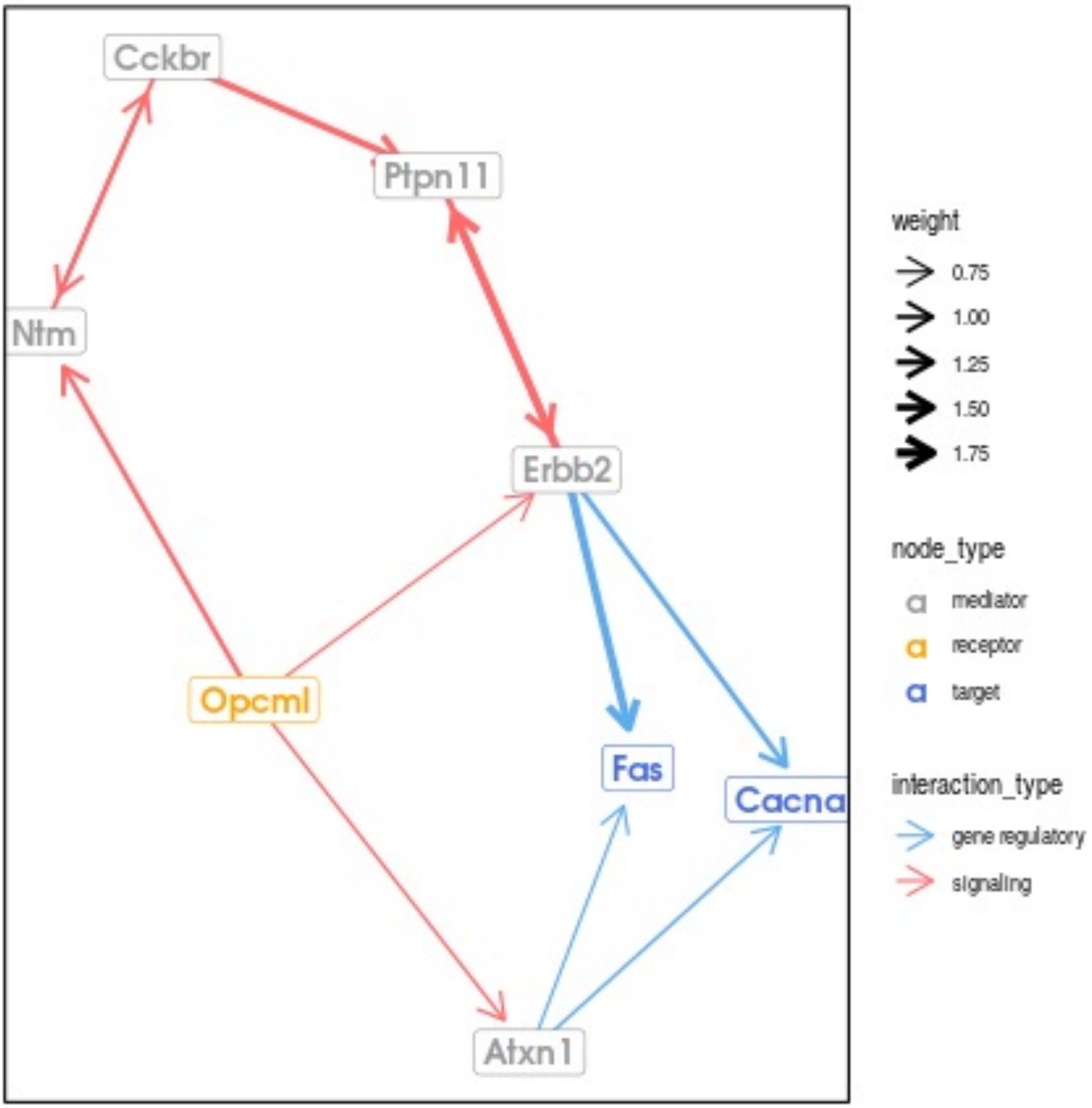

Cckbr
Ptpn11
Ntm
Erbb2
Opcml
Fas
Cacna
Atxn1
weight
0.75
1.00
1.25
1.50
1.75
node_type
mediator
receptor
target
interaction_type
gene regulatory
signaling

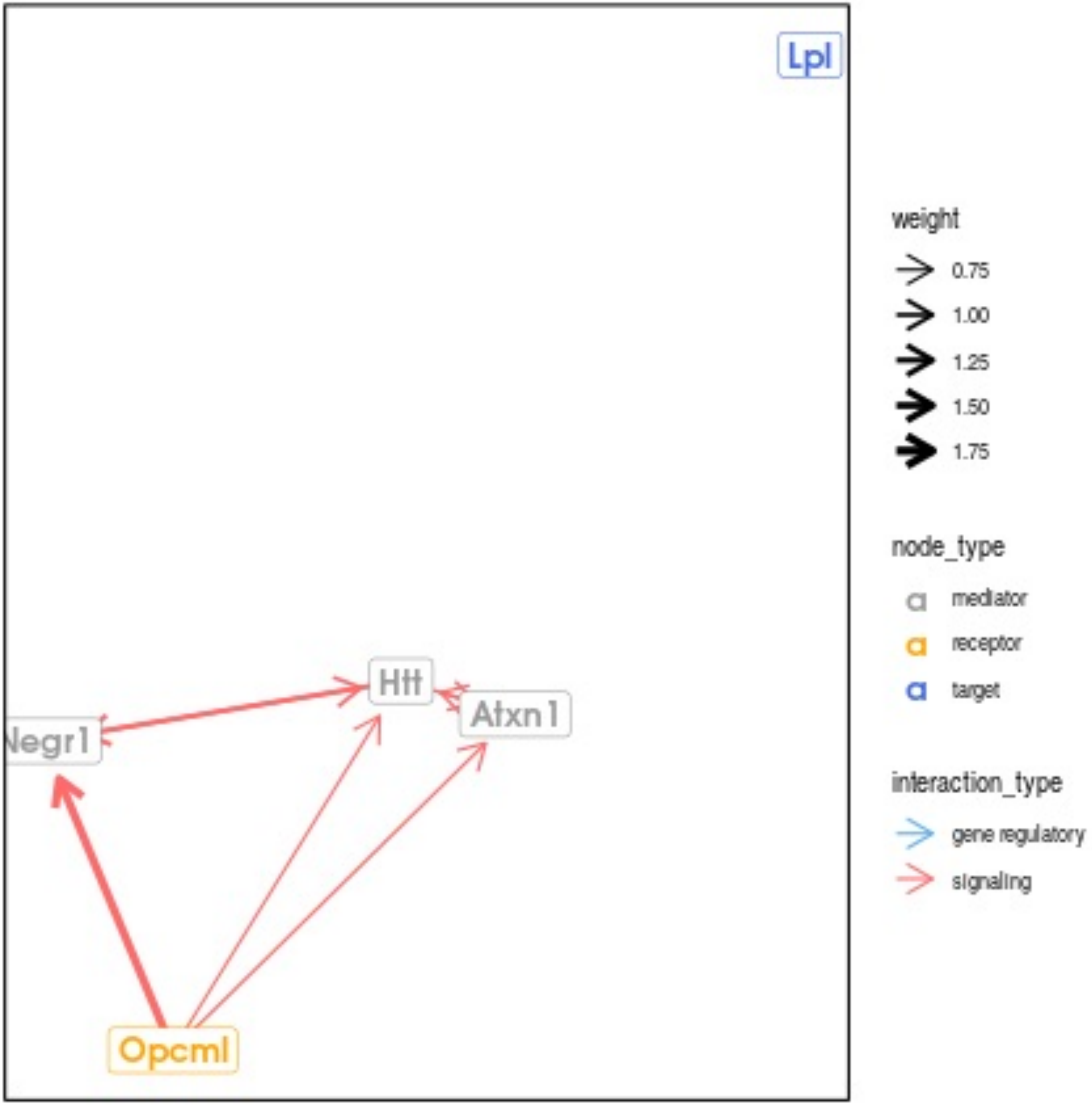

Lpl
Htt
Atxn1
Negr1
Opcml
weight
0.75
1.00
1.25
1.50
1.75
node_type
mediator
receptor
target
interaction_type
gene regulatory
signaling

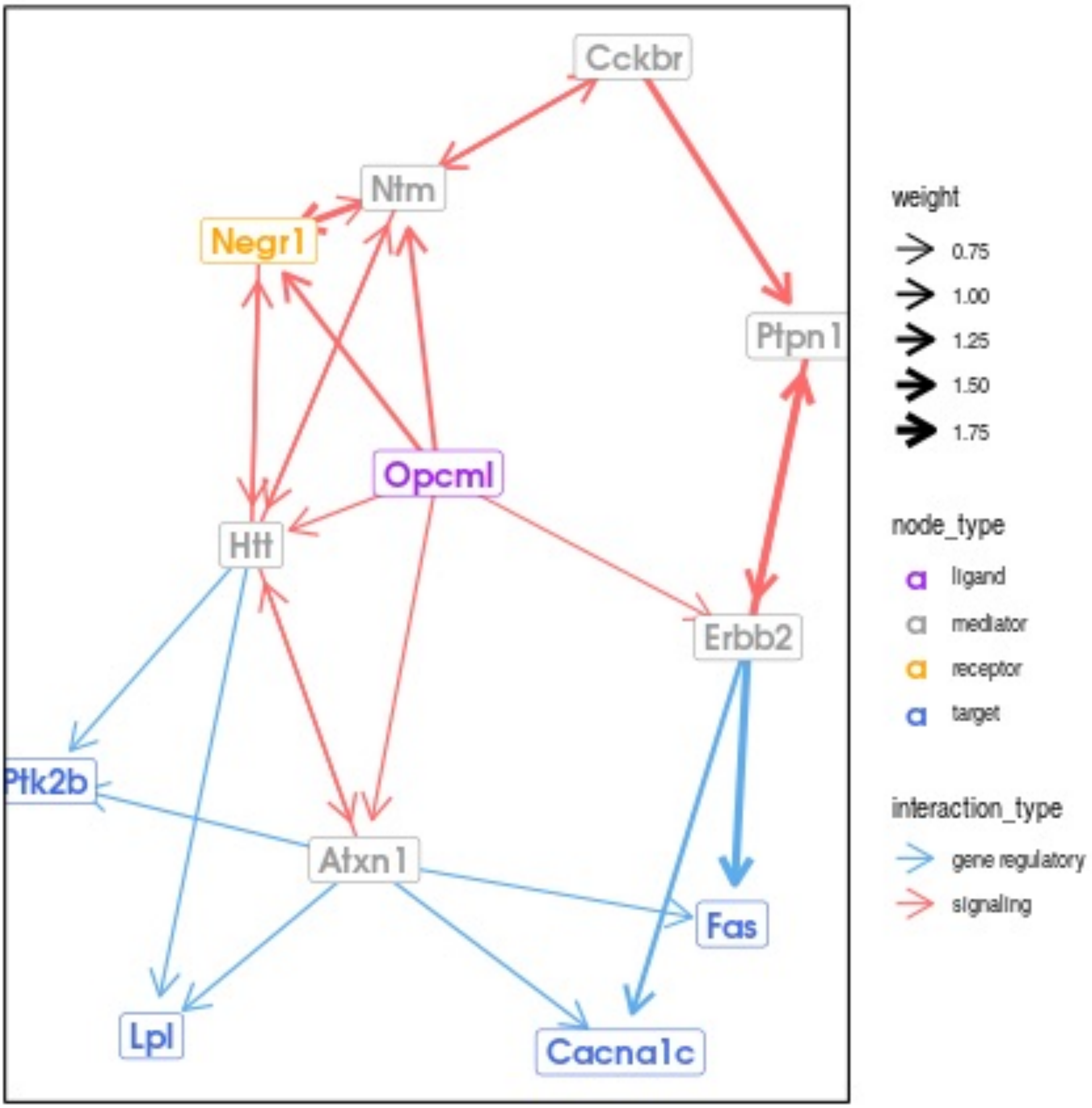
Cckbr
Ntm
Negr1
Ptpn1
Opcml
Htt
Erbb2
Ptk2b
Atxn1
Fas
Lpl
Cacna1c
weight
0.75
1.00
1.25
1.50
1.75
node_type
ligand
mediator
receptor
target
interaction_type
gene regulatory
signaling

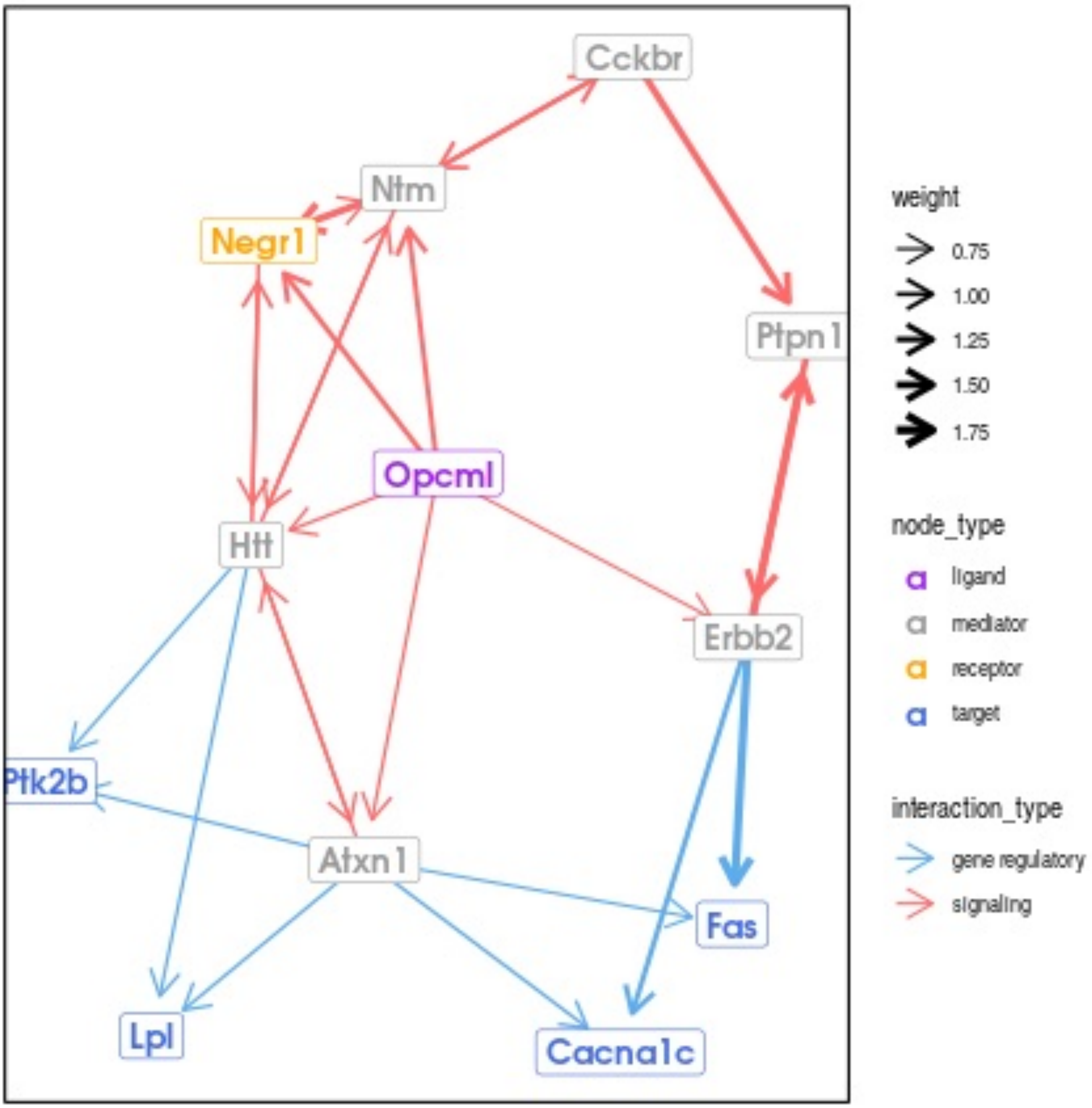
Cckbr
Ntm
Negr1
Ptpn1
Opcml
Htt
Erbb2
Ptk2b
Atxn1
Fas
Lpl
Cacna1c
weight
0.75
1.00
1.25
1.50
1.75
node_type
ligand
mediator
receptor
target
interaction_type
gene regulatory
signaling

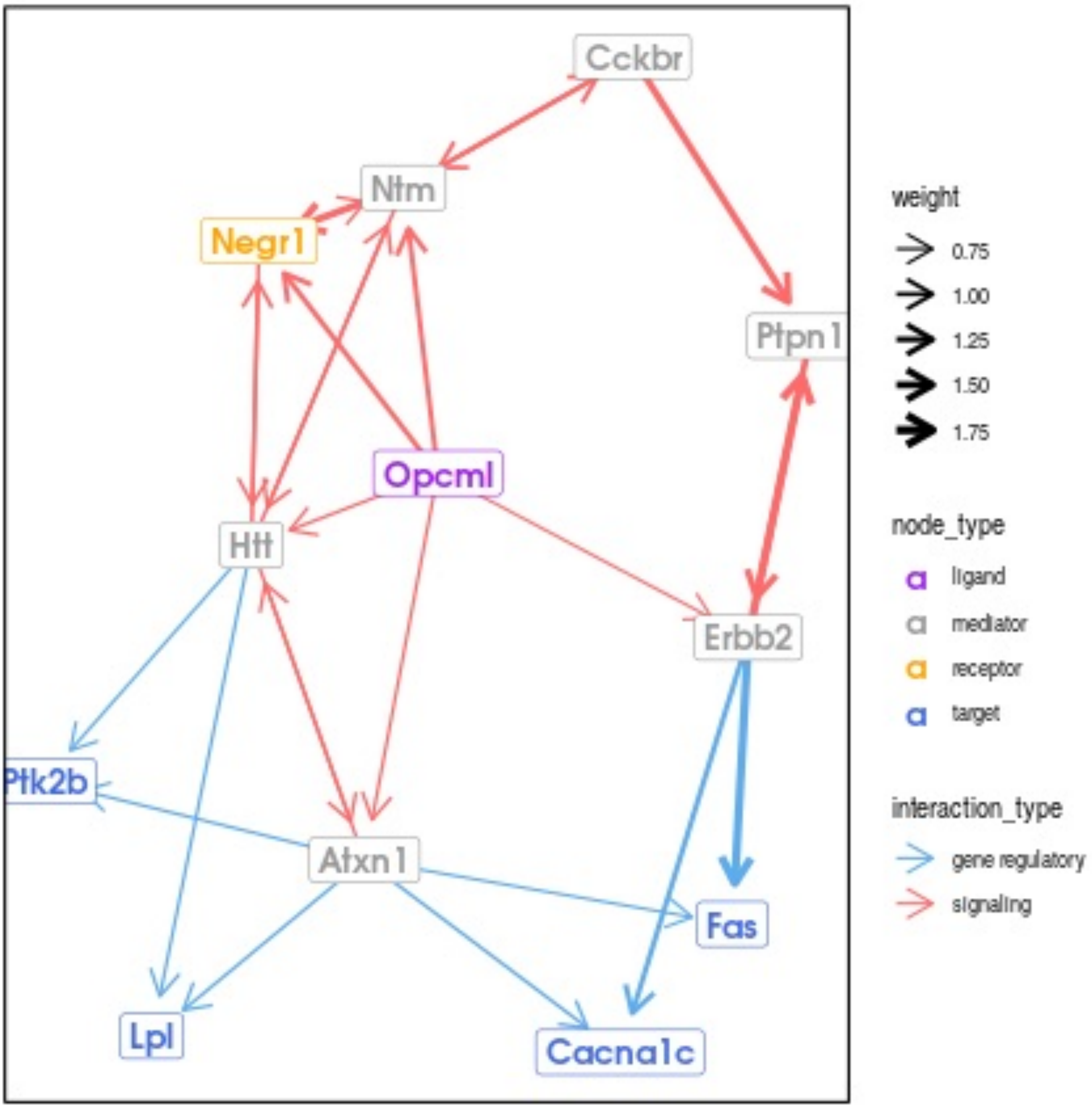

Cckbr
Ntm
Negr1
Ptpn1
Opcml
Htt
Erbb2
Ptk2b
Atxn1
Fas
Lpl
Cacna1c
weight
0.75
1.00
1.25
1.50
1.75
node_type
ligand
mediator
receptor
target
interaction_type
gene regulatory
signaling

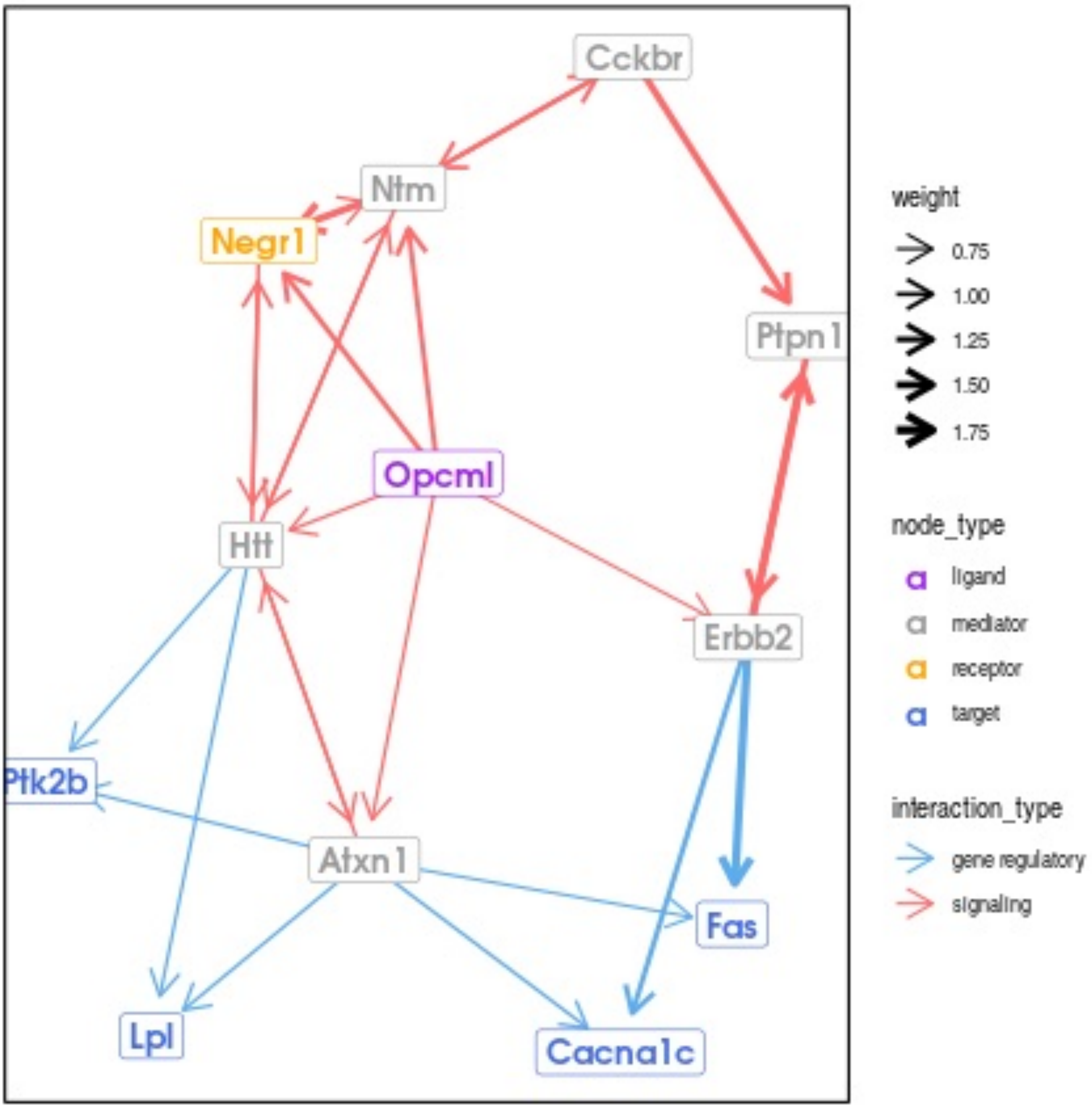
Cckbr
Ntm
Negr1
Ptpn1
Opcml
Htt
Erbb2
Ptk2b
Atxn1
Fas
Lpl
Cacna1c
weight
0.75
1.00
1.25
1.50
1.75
node_type
ligand
mediator
receptor
target
interaction_type
gene regulatory
signaling

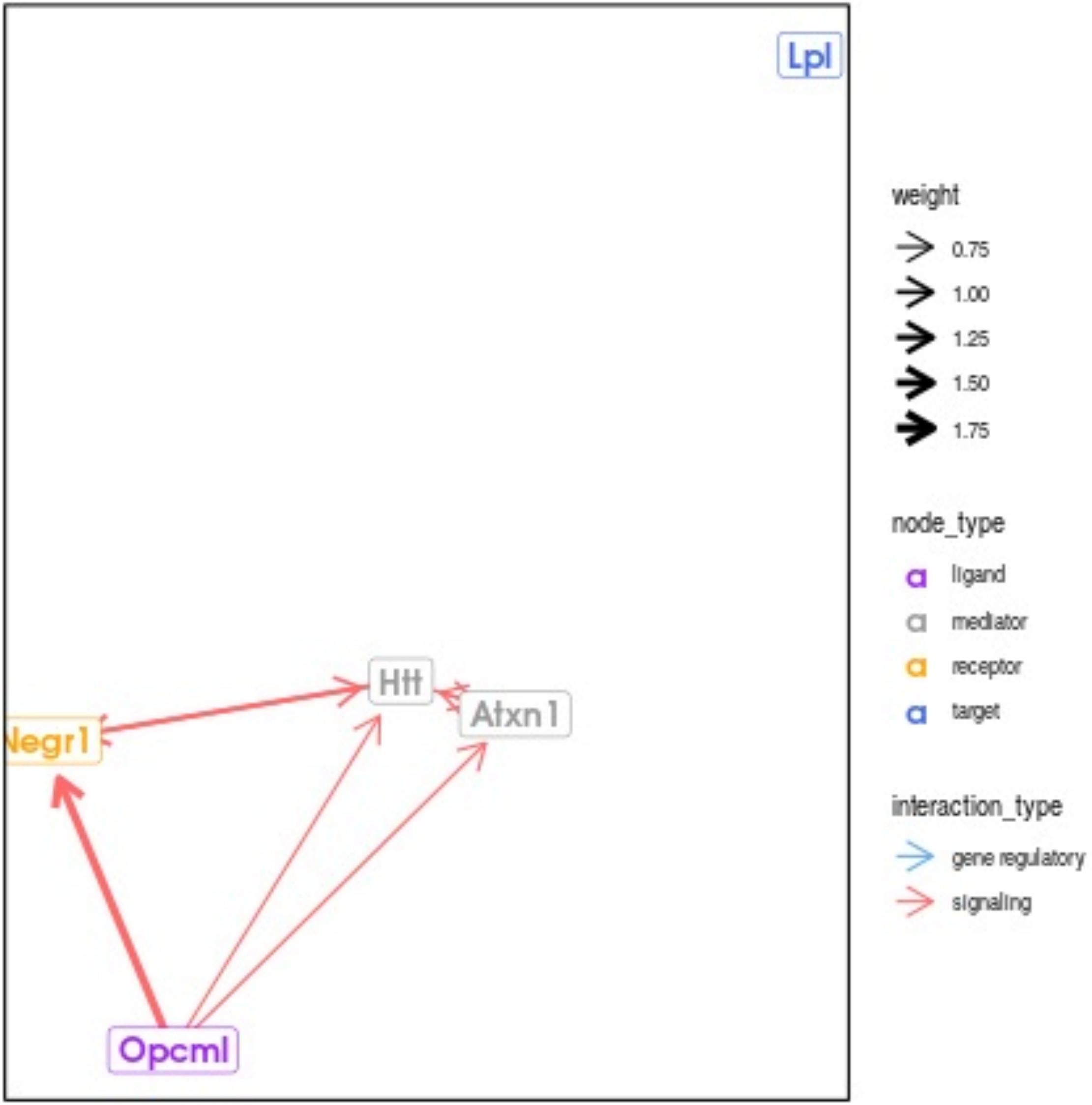
Lpl
Htt
Atxn1
Negr1
Opcml
weight
0.75
1.00
1.25
1.50
1.75
node_type
ligand
mediator
receptor
target
interaction_type
gene regulatory
signaling

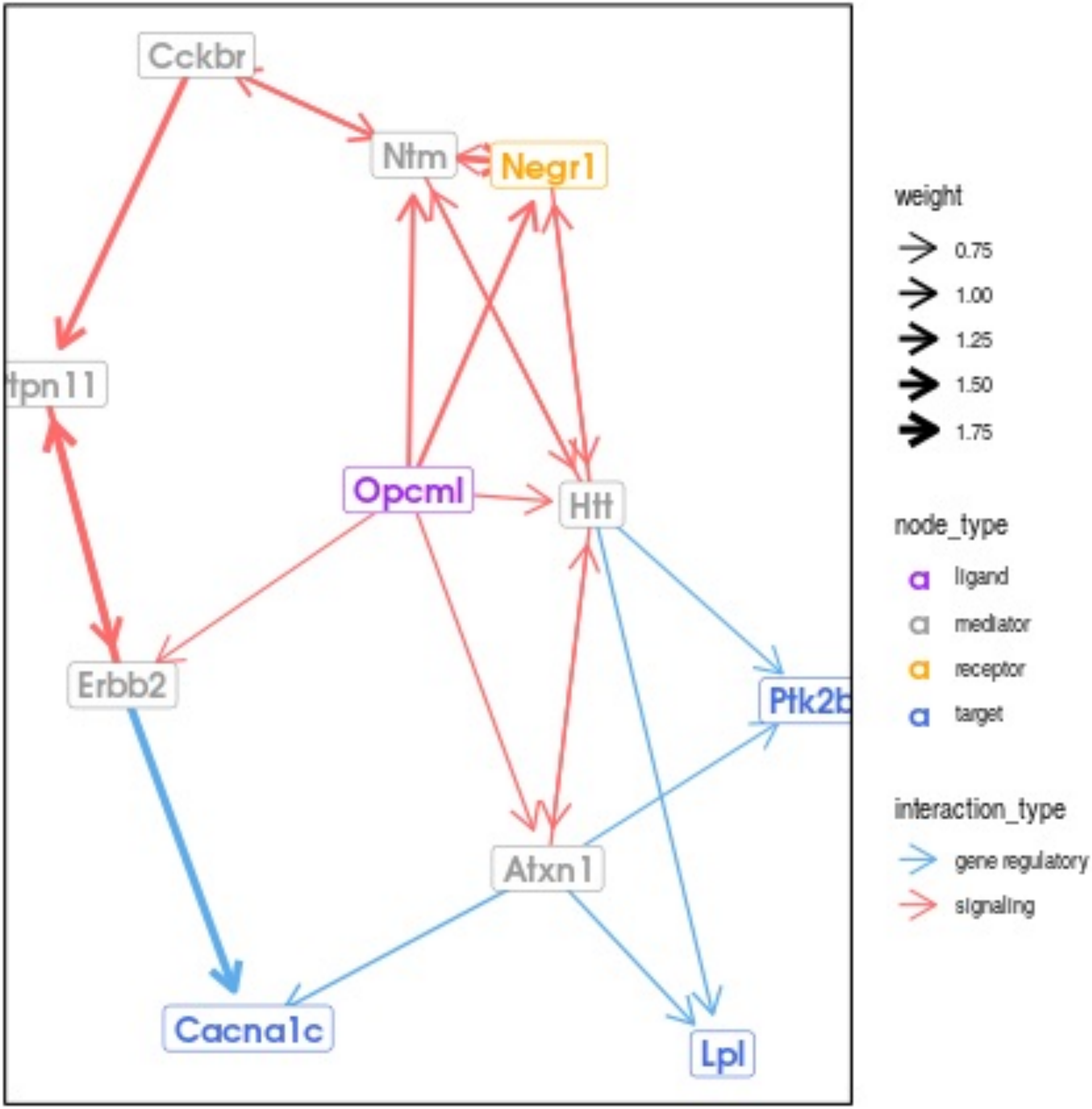
Cckbr
Ntm
Negr1
tpn11
Opcml
Htt
Erbb2
Ptk2b
Atxn1
Cacna1c
Lpl
weight
0.75
1.00
1.25
1.50
1.75
node_type
ligand
mediator
receptor
target
interaction_type
gene regulatory
signaling

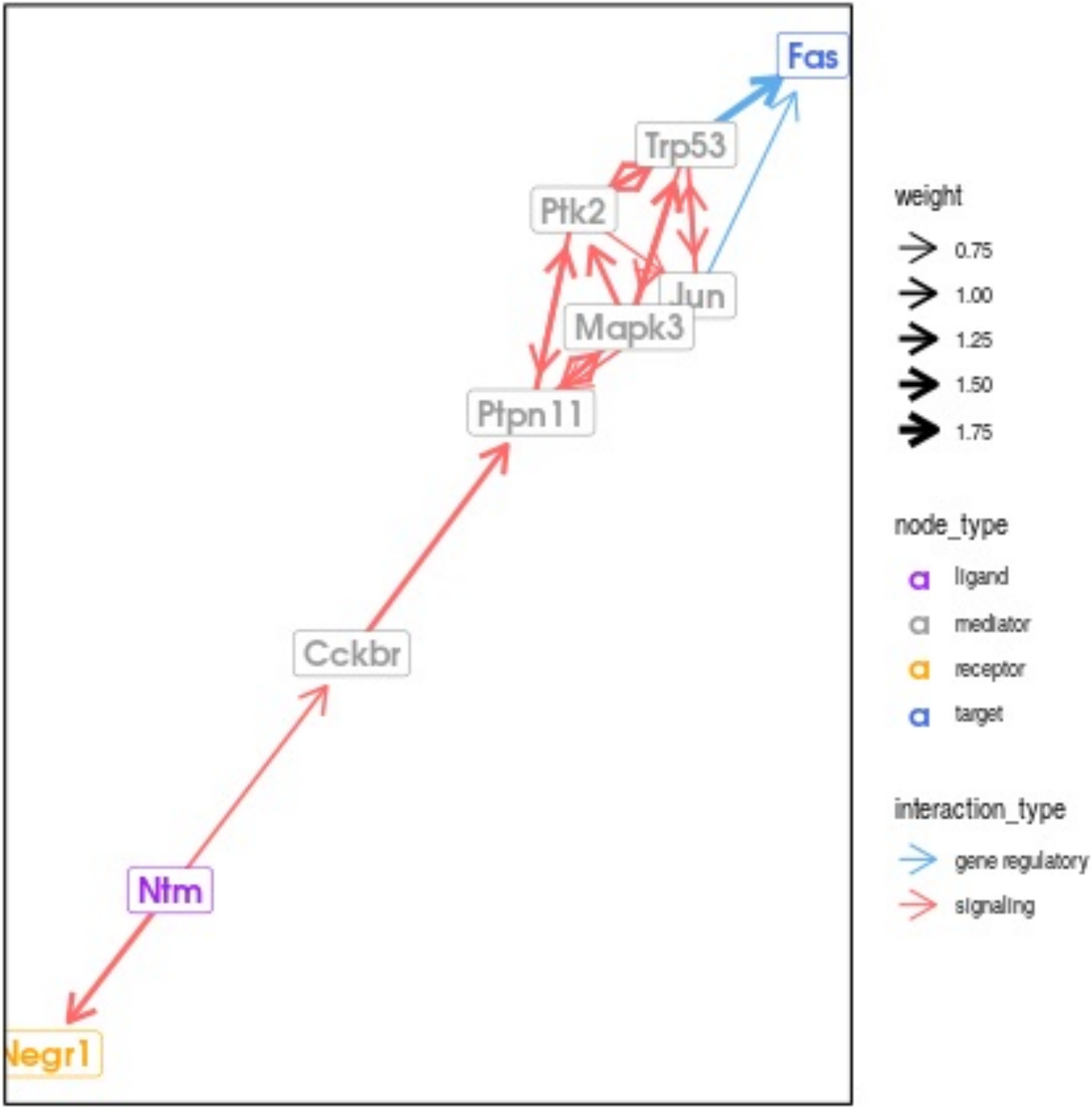
Fas
Trp53
Ptk2
Jun
Mapk3
Ptpn11
Cckbr
Ntm
Negr1
weight
0.75
1.00
1.25
1.50
1.75
node_type
ligand
mediator
receptor
target
interaction_type
gene regulatory
signaling

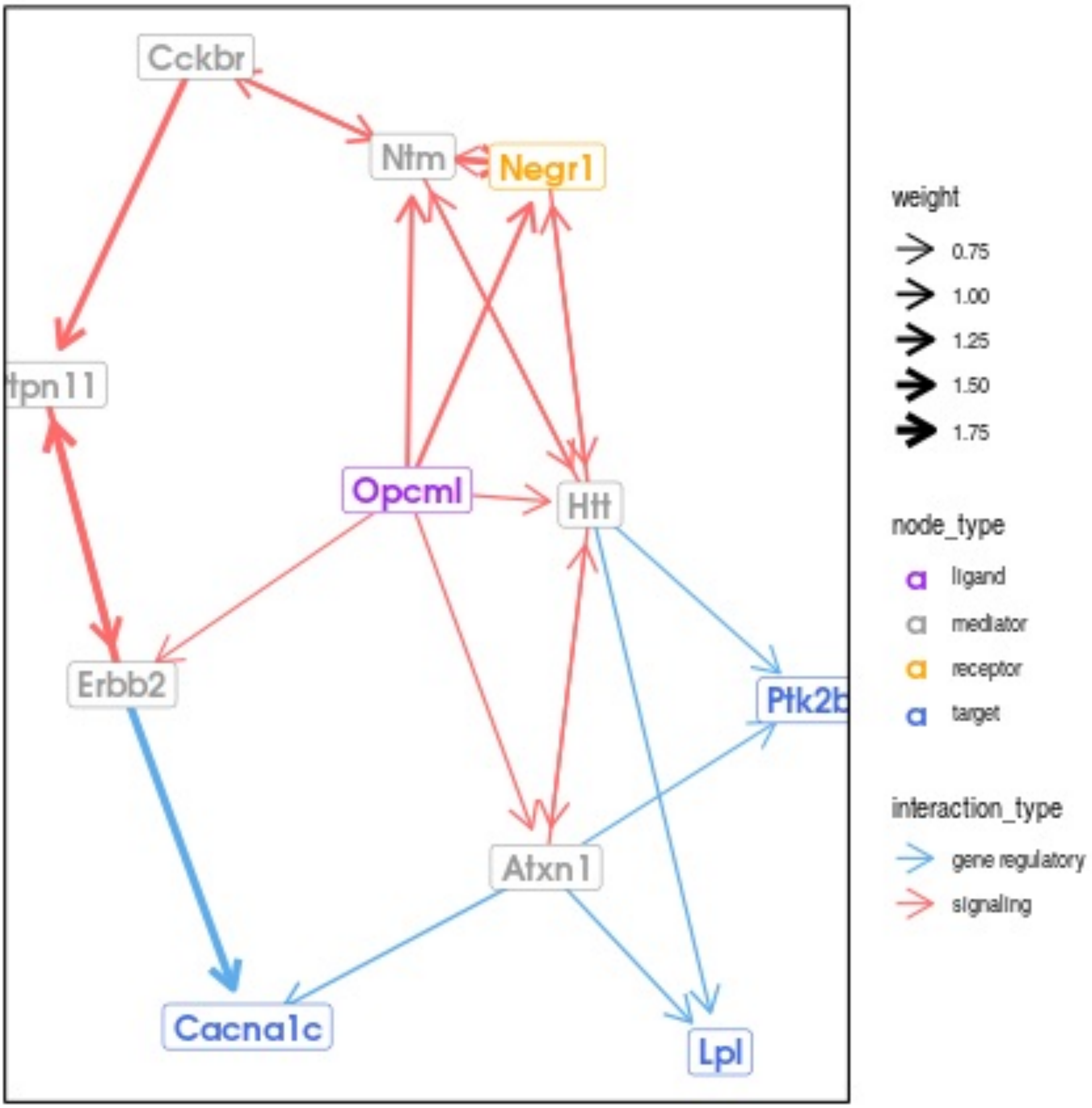

Cckbr
Ntm
Negr1
tpn11
Opcml
Htt
Erbb2
Ptk2b
Atxn1
Cacna1c
Lpl
weight
0.75
1.00
1.25
1.50
1.75
node_type
a ligand
a mediator
a receptor
a target
interaction_type
gene regulatory
signaling

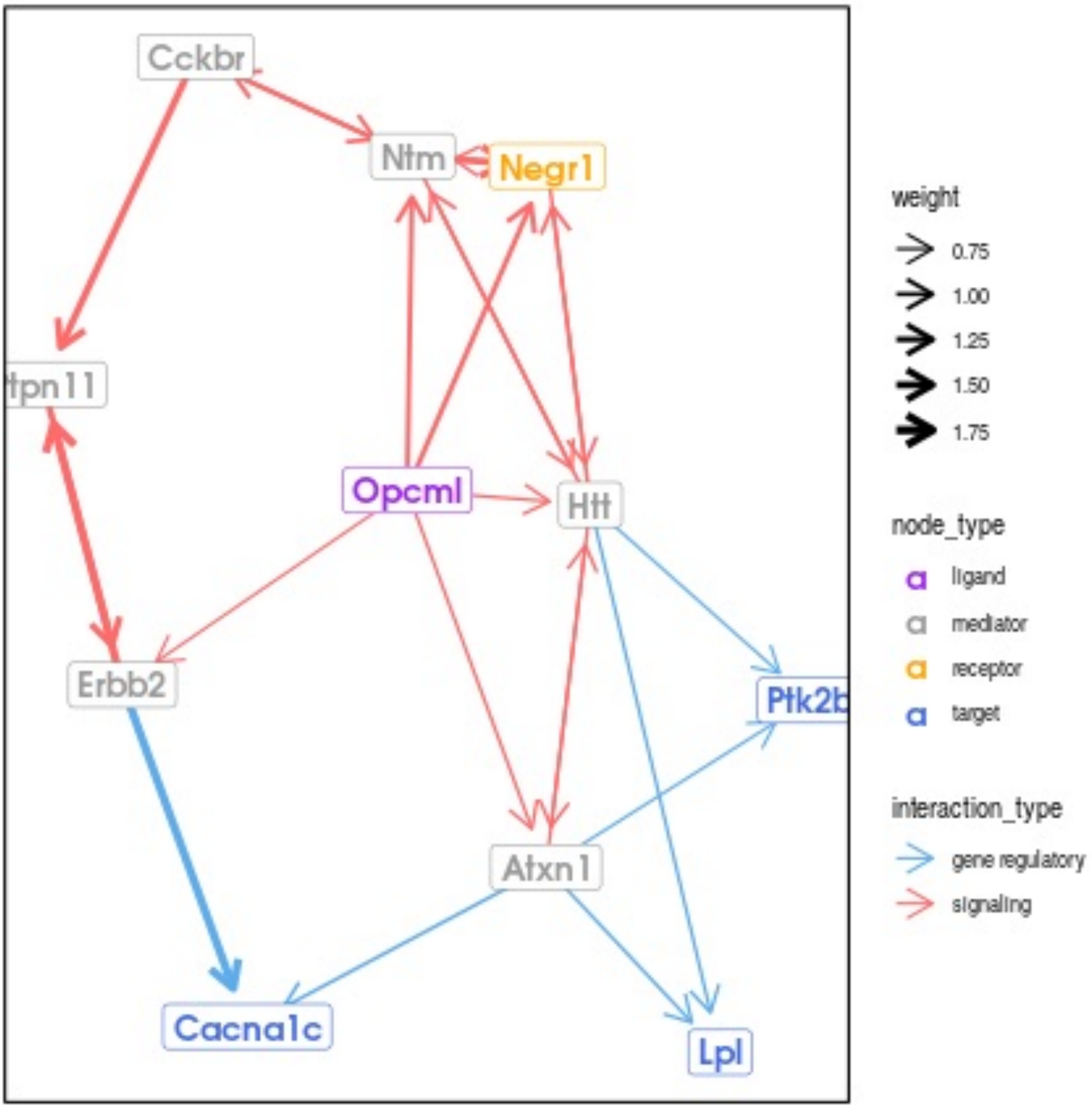
Cckbr
Ntm
Negr1
tpn11
Opcml
Htt
Erbb2
Ptk2b
Atxn1
Cacna1c
Lpl
weight
0.75
1.00
1.25
1.50
1.75
node_type
a ligand
a mediator
a receptor
a target
interaction_type
gene regulatory
signaling

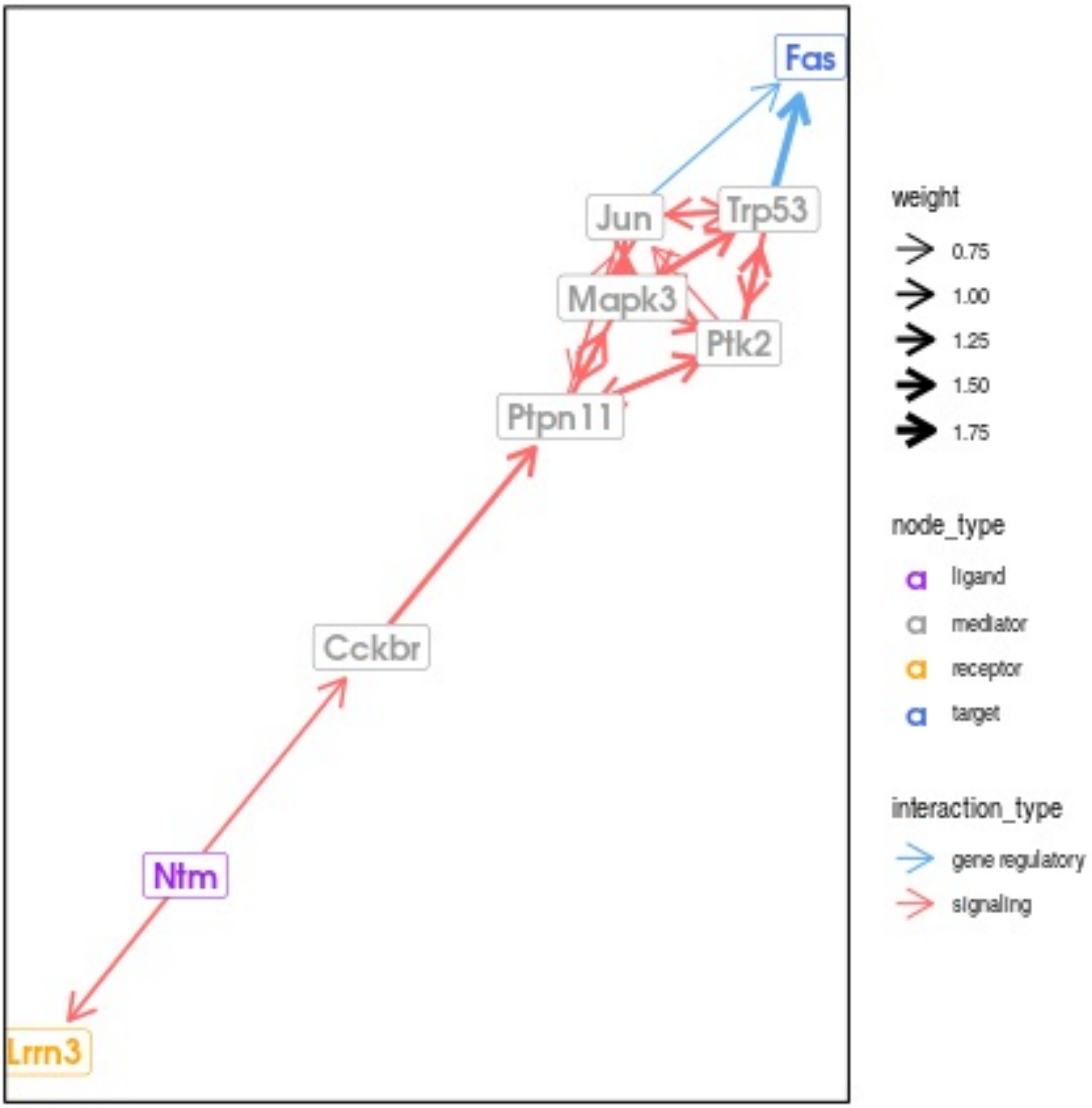
Fas
Jun
Trp53
Mapk3
Ptk2
Ptpn11
Cckbr
Ntm
Lrrn3
weight
0.75
1.00
1.25
1.50
1.75
node_type
ligand
mediator
receptor
target
interaction_type
gene regulatory
signaling

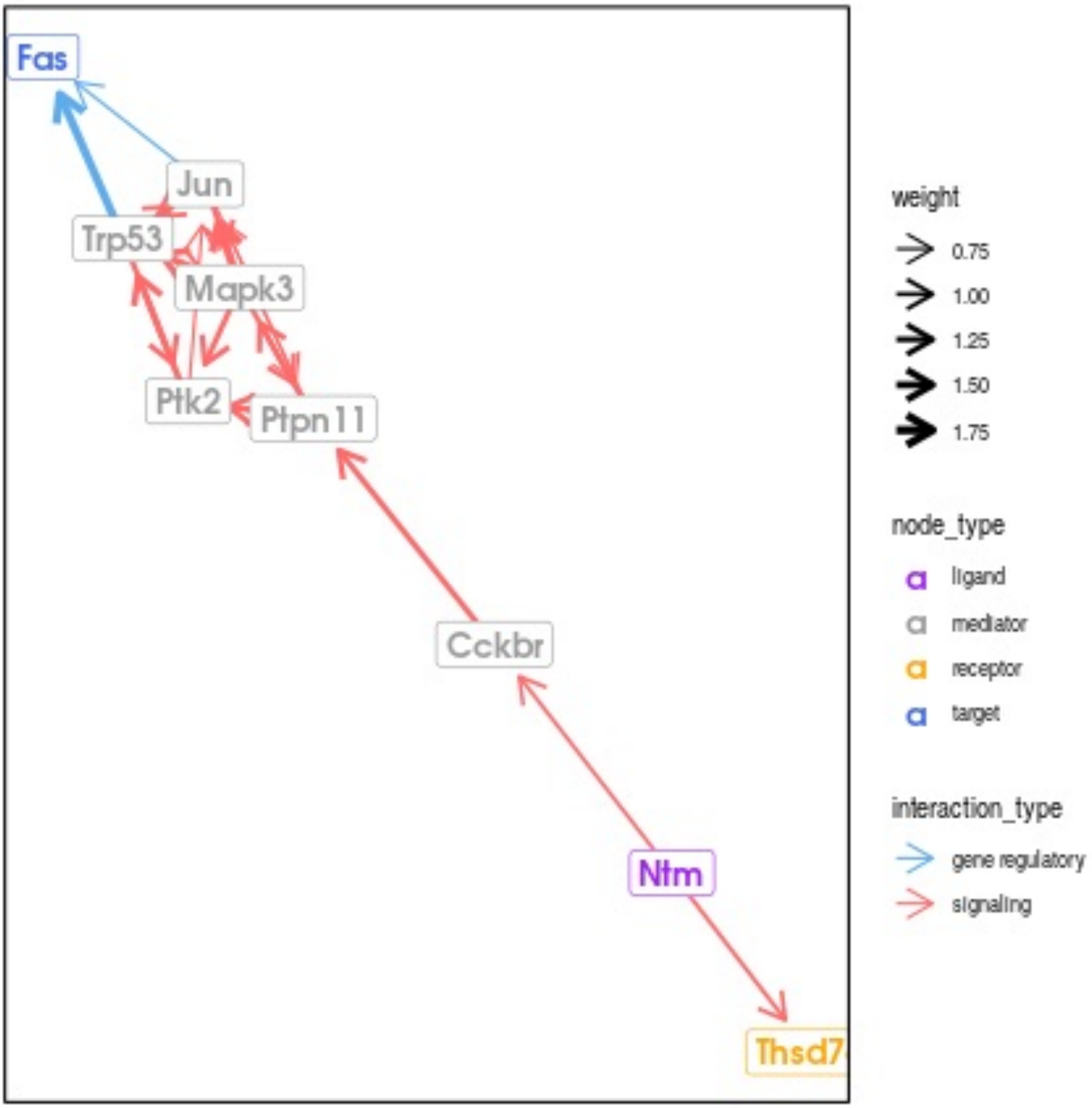

Fas
Jun
Trp53
Mapk3
Ptk2
Ptpn11
Cckbr
Ntm
Thsd7
weight
0.75
1.00
1.25
1.50
1.75
node_type
ligand
mediator
receptor
target
interaction_type
gene regulatory
signaling

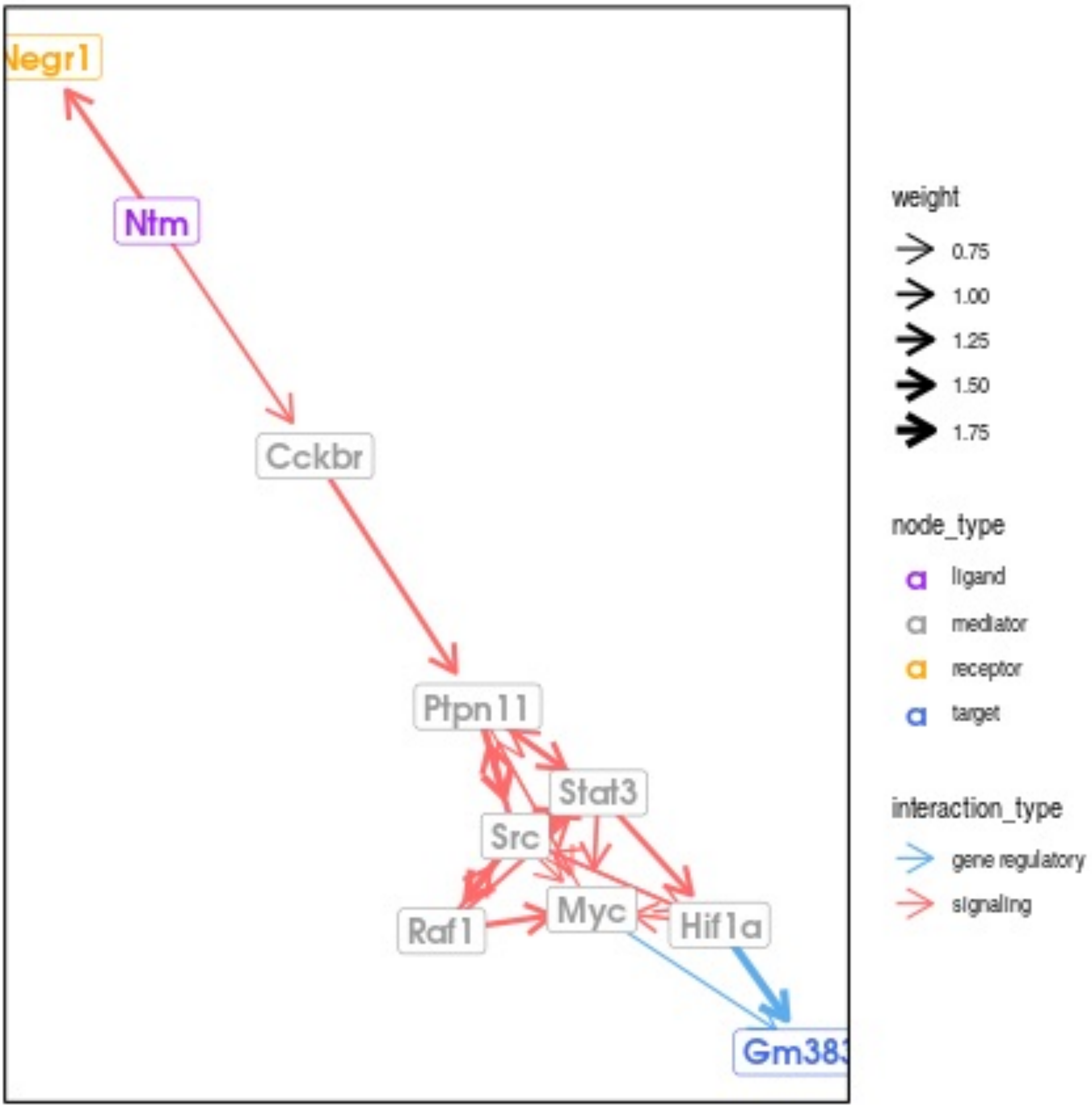

Negr1
Ntm
Cckbr
Ptpn11
Stat3
Src
Myc
Raf1
Hif1a
Gm383
weight
0.75
1.00
1.25
1.50
1.75
node_type
ligand
mediator
receptor
target
interaction_type
gene regulatory
signaling

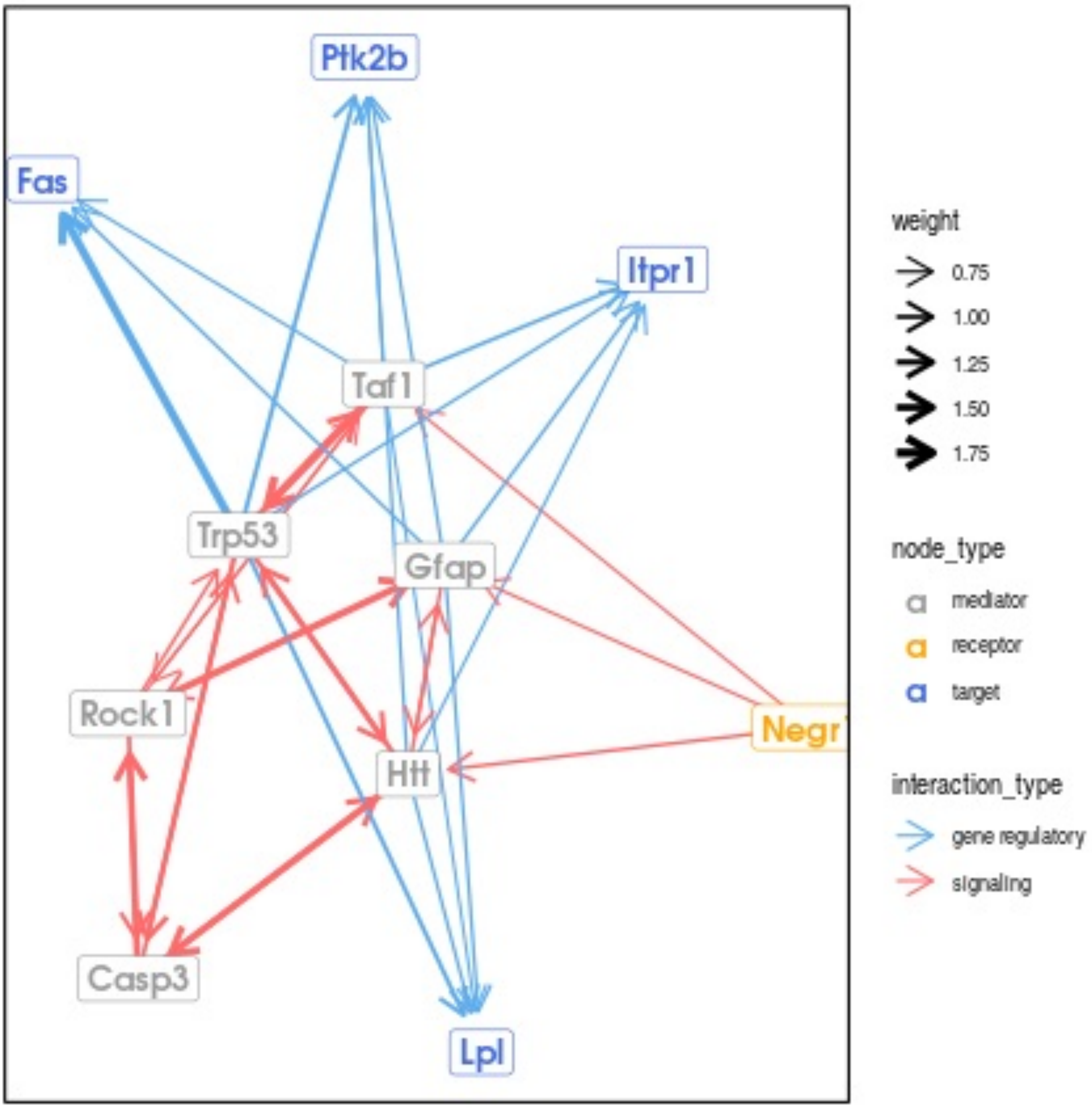

Ptk2b
Fas
Itpr1
Taf1
Trp53
Gfap
Rock1
Negr1
Htt
Casp3
Lpl
weight
0.75
1.00
1.25
1.50
1.75
node_type
mediator
receptor
target
interaction_type
gene regulatory
signaling

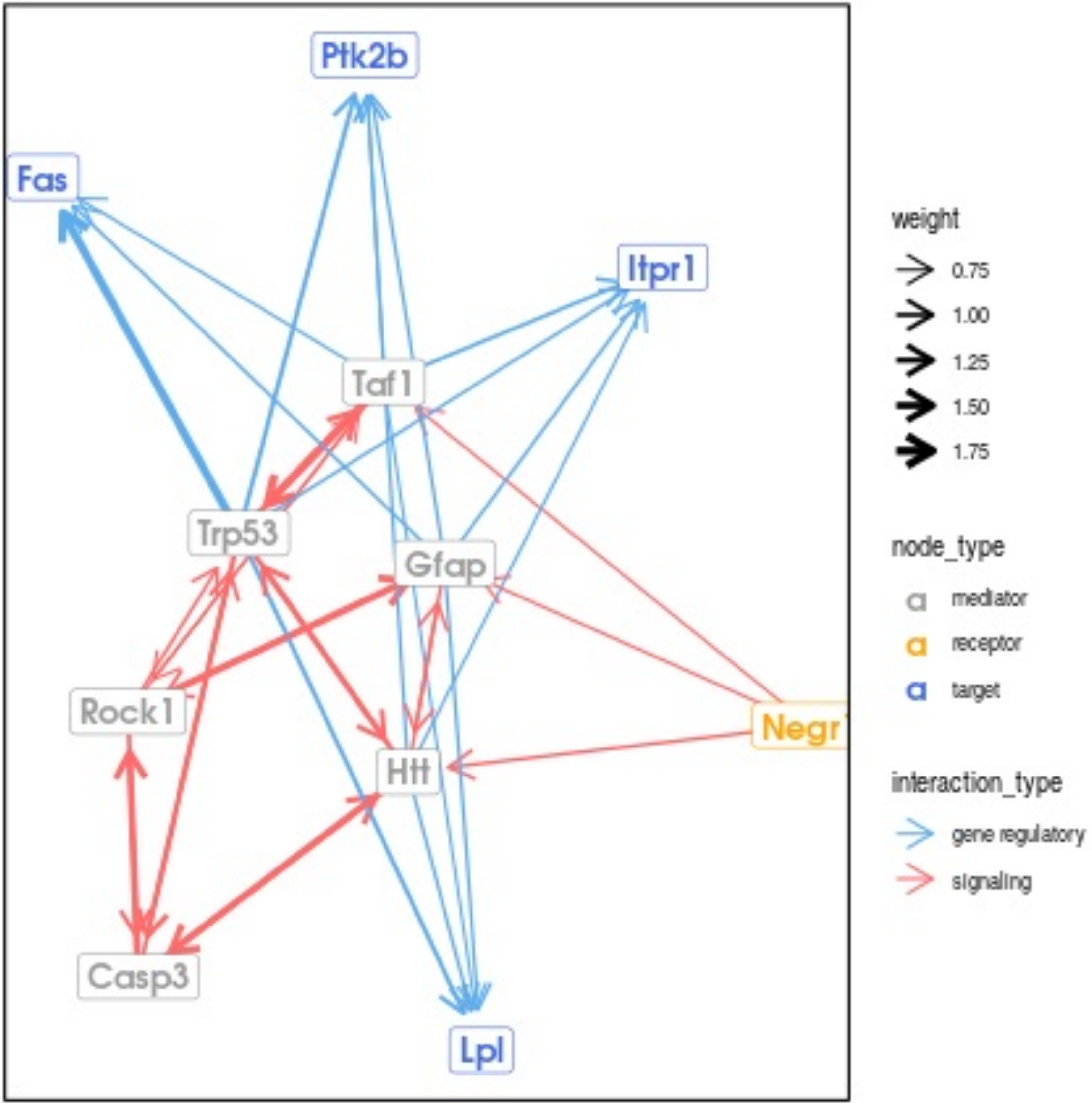
Ptk2b
Fas
Itpr1
Taf1
Trp53
Gfap
Rock1
Negr1
Htt
Casp3
Lpl
weight
0.75
1.00
1.25
1.50
1.75
node_type
mediator
receptor
target
interaction_type
gene regulatory
signaling

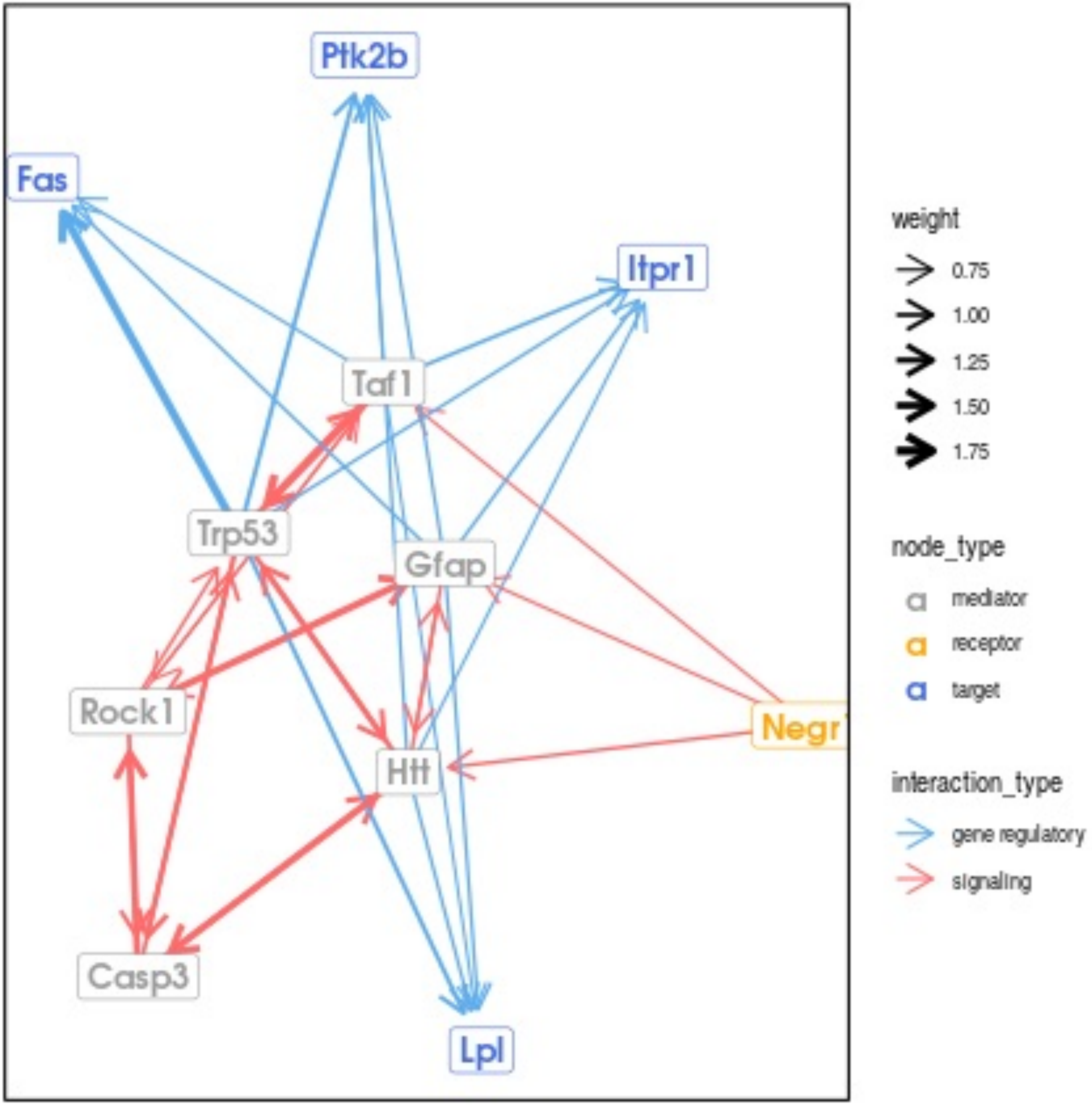

Ptk2b
Fas
Itpr1
Taf1
Trp53
Gfap
Rock1
Negr1
Htt
Casp3
Lpl
weight
0.75
1.00
1.25
1.50
1.75
node_type
mediator
receptor
target
interaction_type
gene regulatory
signaling

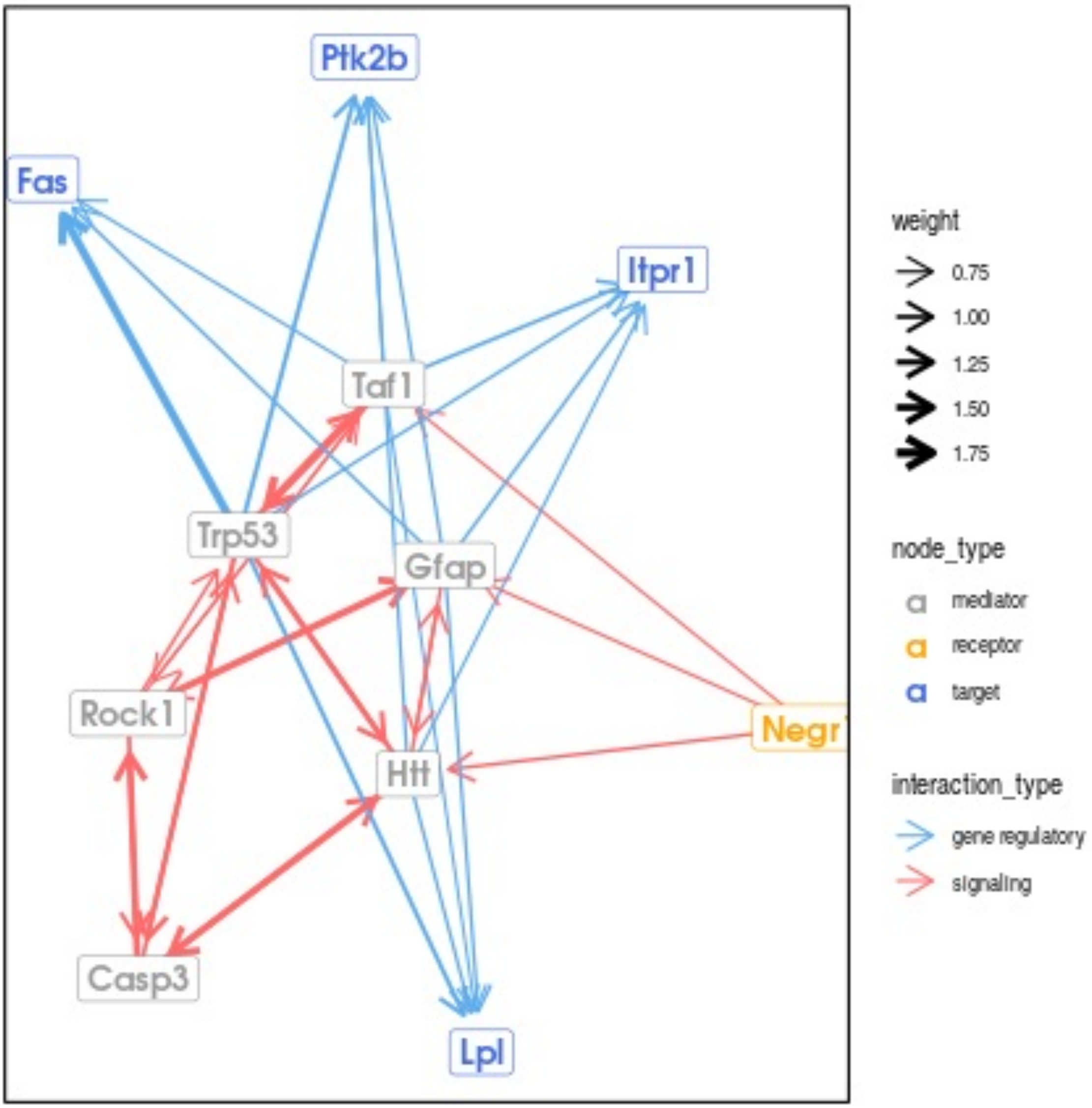

Ptk2b
Fas
Itpr1
Taf1
Trp53
Gfap
Rock1
Negr1
Htt
Casp3
Lpl
weight
0.75
1.00
1.25
1.50
1.75
node_type
mediator
receptor
target
interaction_type
gene regulatory
signaling

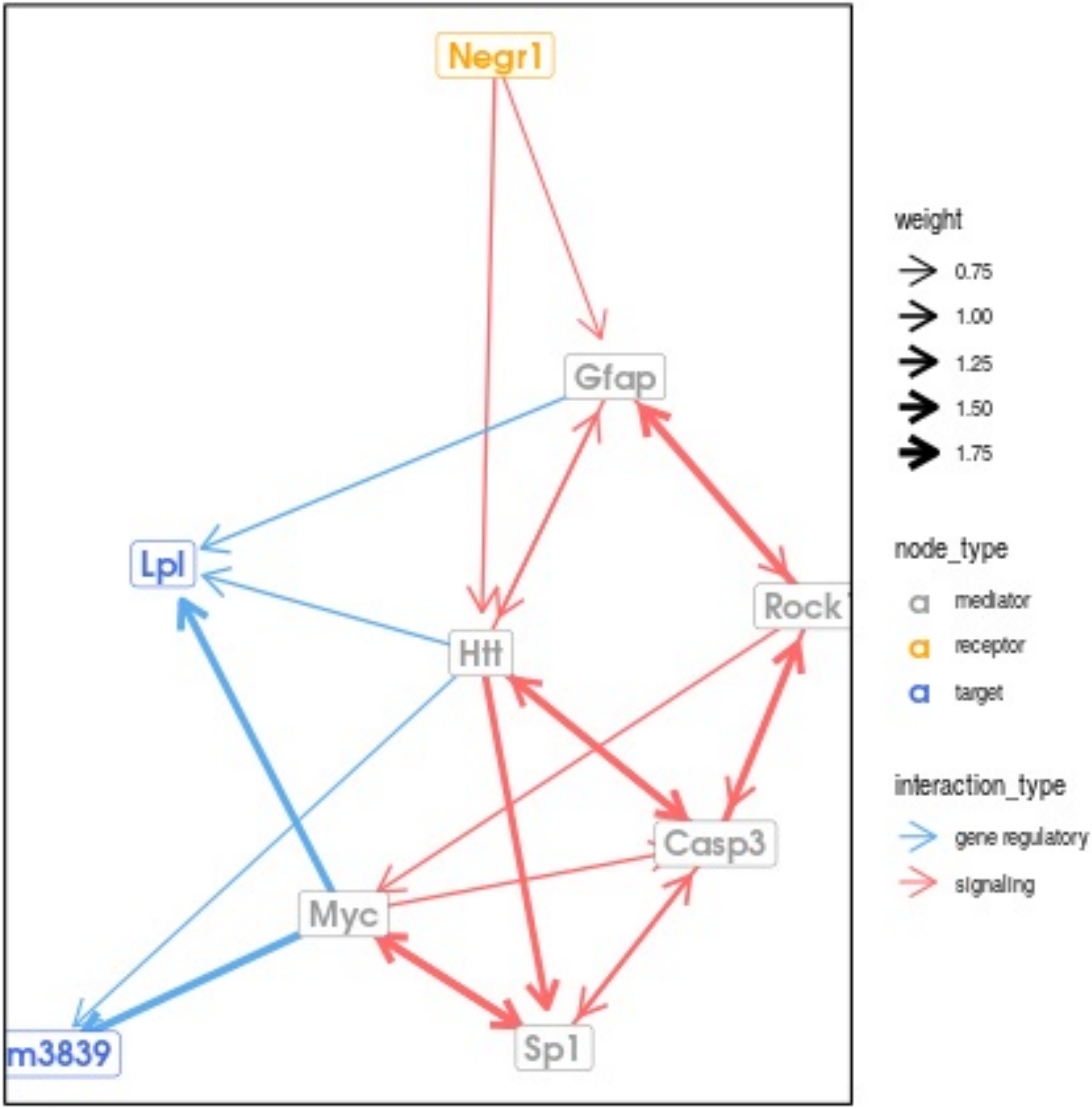

Negr1
Gfap
Lpl
Rock
Htt
Casp3
Myc
m3839
Sp1
weight
0.75
1.00
1.25
1.50
1.75
node_type
mediator
receptor
target
interaction_type
gene regulatory
signaling

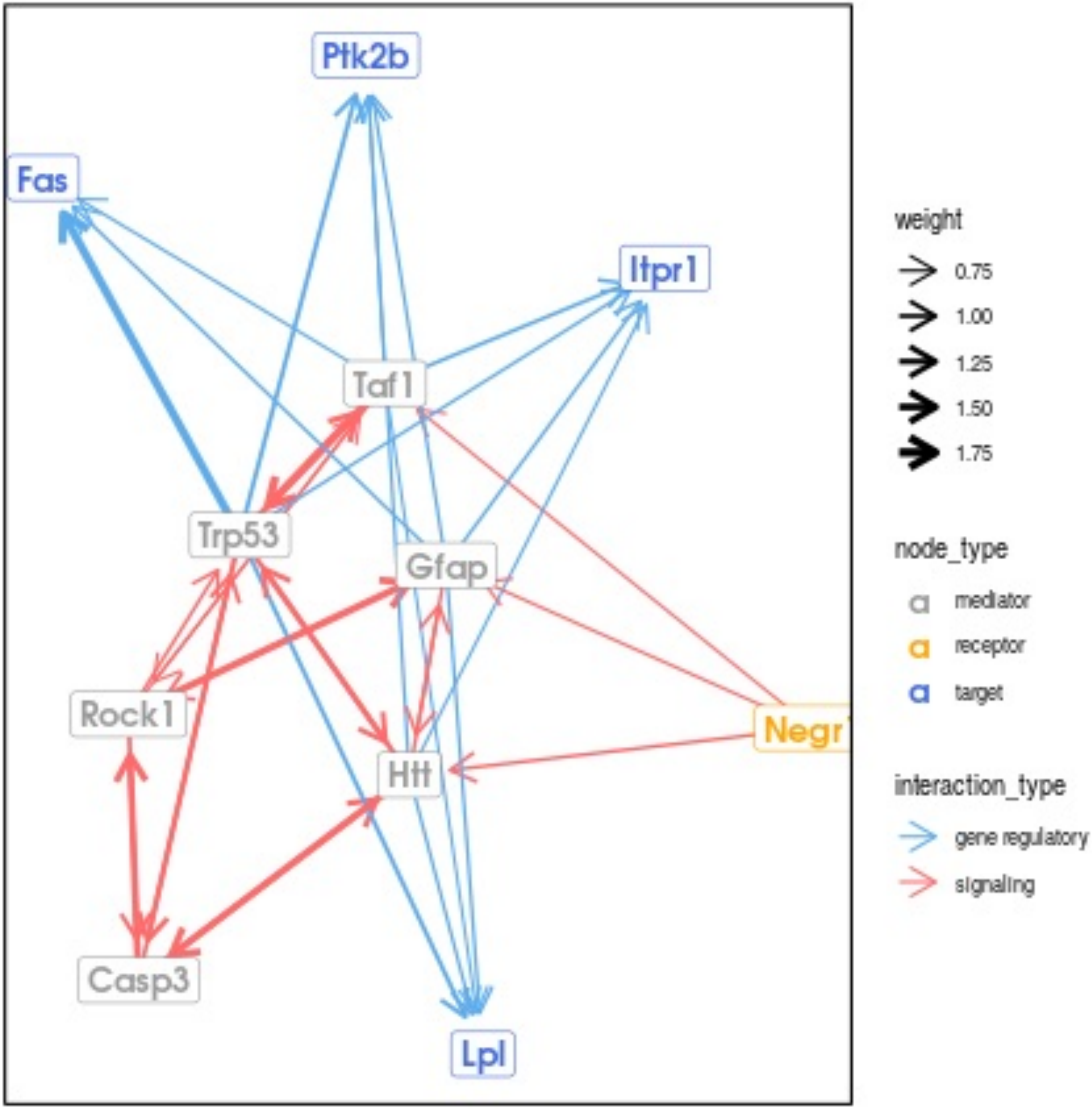

Ptk2b
Fas
Itpr1
Taf1
Trp53
Gfap
Rock1
Negr1
Htt
Casp3
Lpl
weight
0.75
1.00
1.25
1.50
1.75
node_type
mediator
receptor
target
interaction_type
gene regulatory
signaling

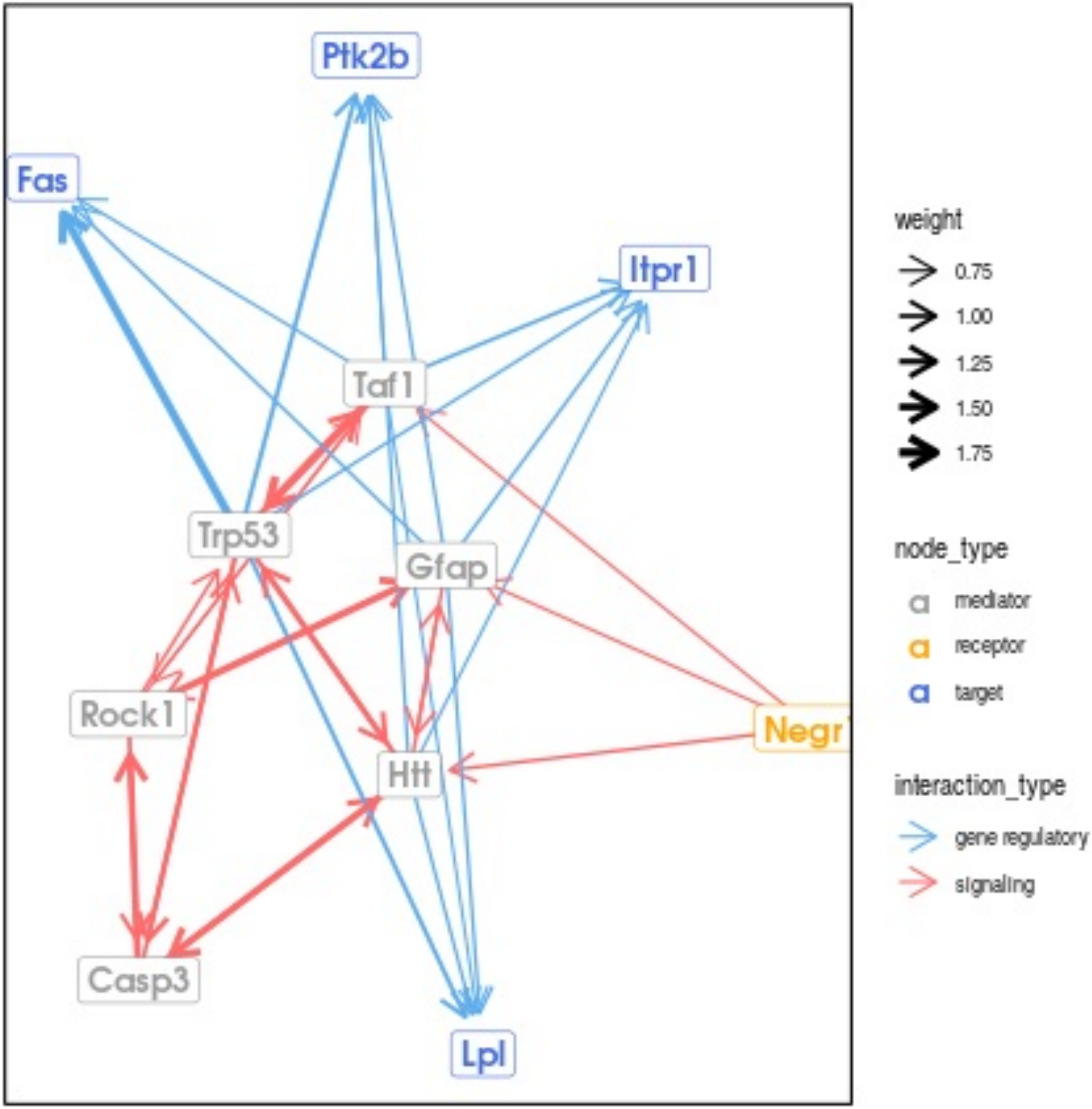

Ptk2b
Fas
Itpr1
Taf1
Trp53
Gfap
Rock1
Htt
Negr1
Casp3
Lpl
weight
0.75
1.00
1.25
1.50
1.75
node_type
mediator
receptor
target
interaction_type
gene regulatory
signaling

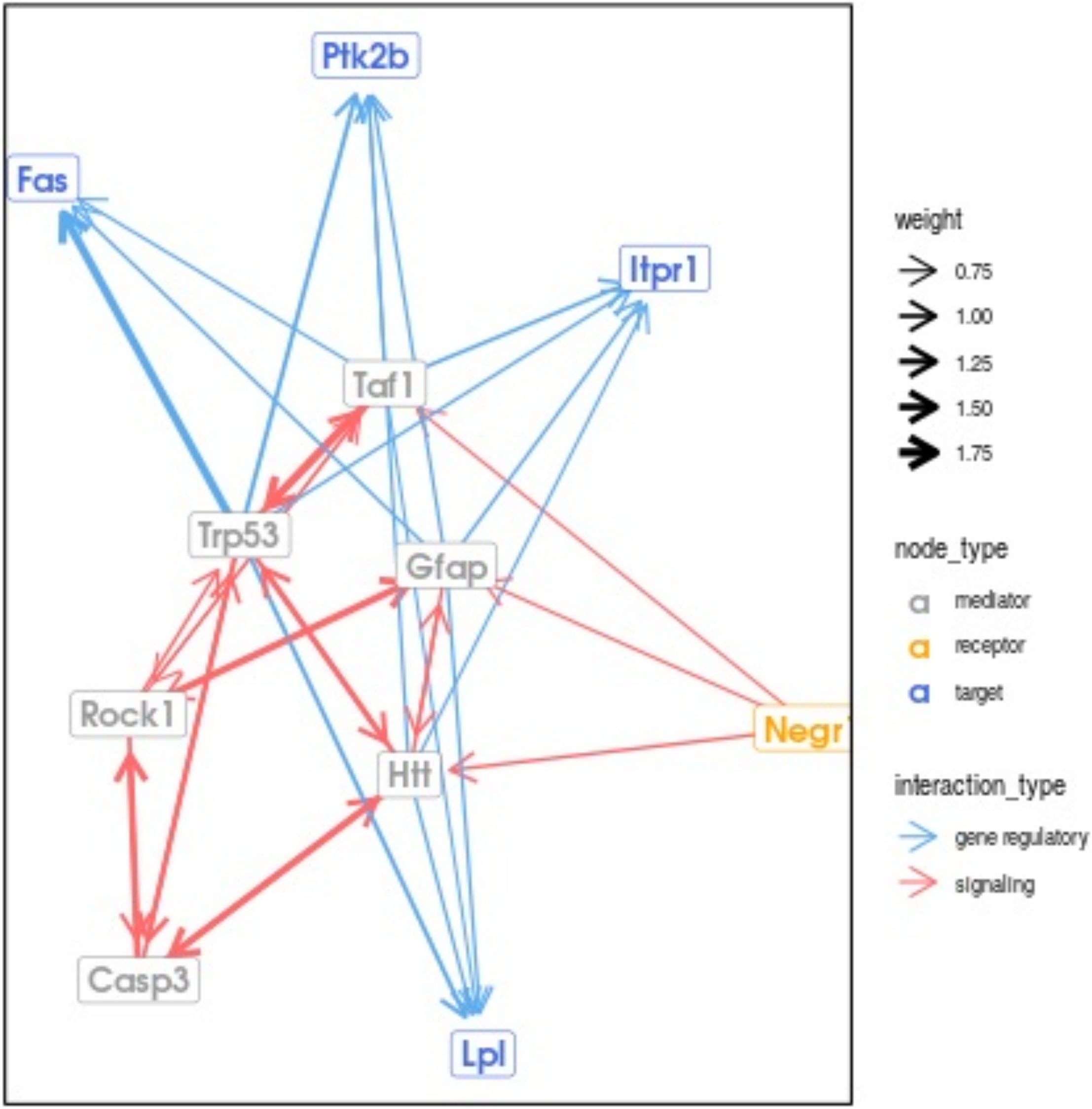
Ptk2b
Fas
Itpr1
Taf1
Trp53
Gfap
Rock1
Negr1
Htt
Casp3
Lpl
weight
0.75
1.00
1.25
1.50
1.75
node_type
mediator
receptor
target
interaction_type
gene regulatory
signaling

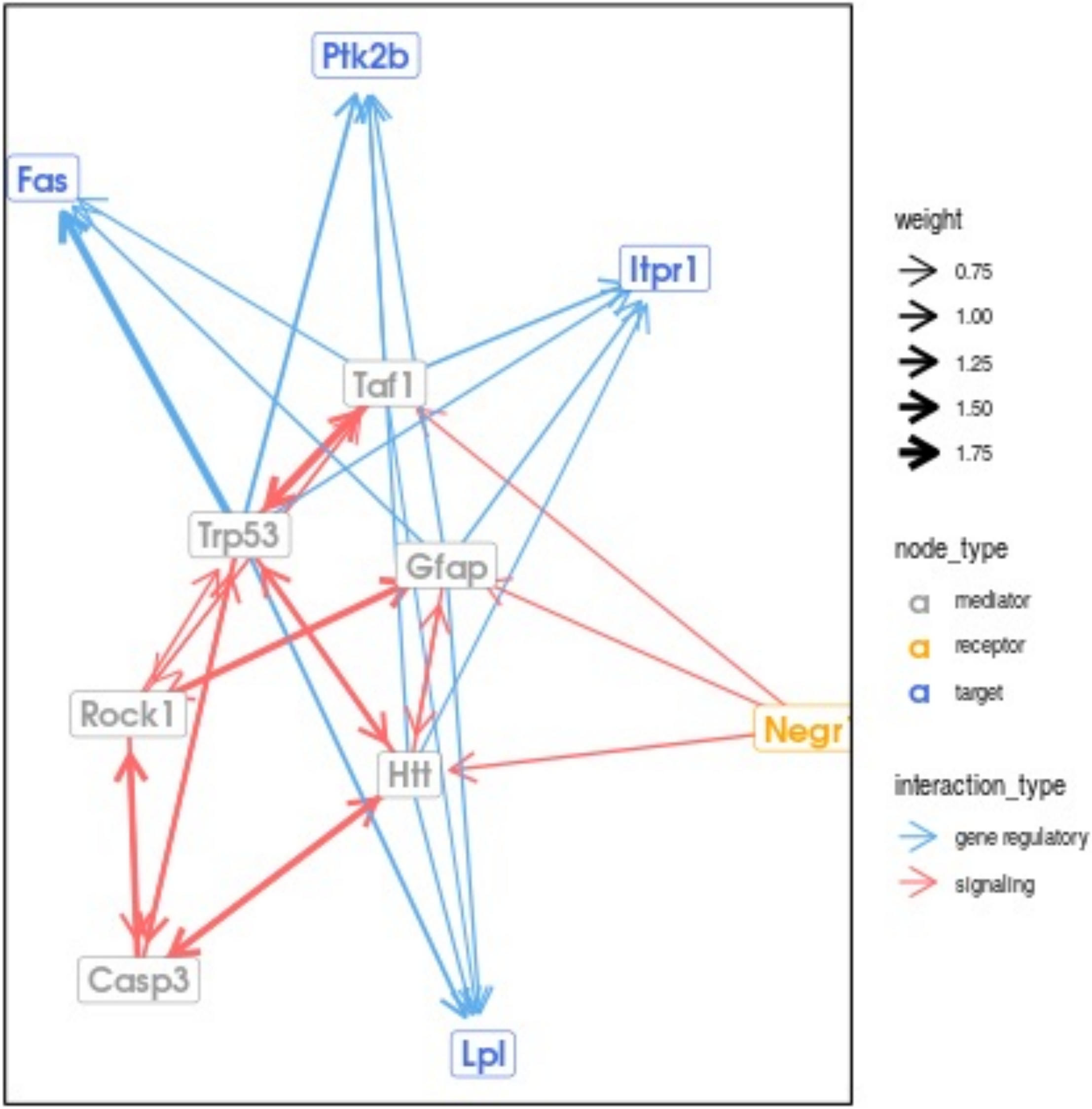
Ptk2b
Fas
Itpr1
Taf1
Trp53
Gfap
Rock1
Negr
Htt
Casp3
Lpl
weight
0.75
1.00
1.25
1.50
1.75
node_type
mediator
receptor
target
interaction_type
gene regulatory
signaling

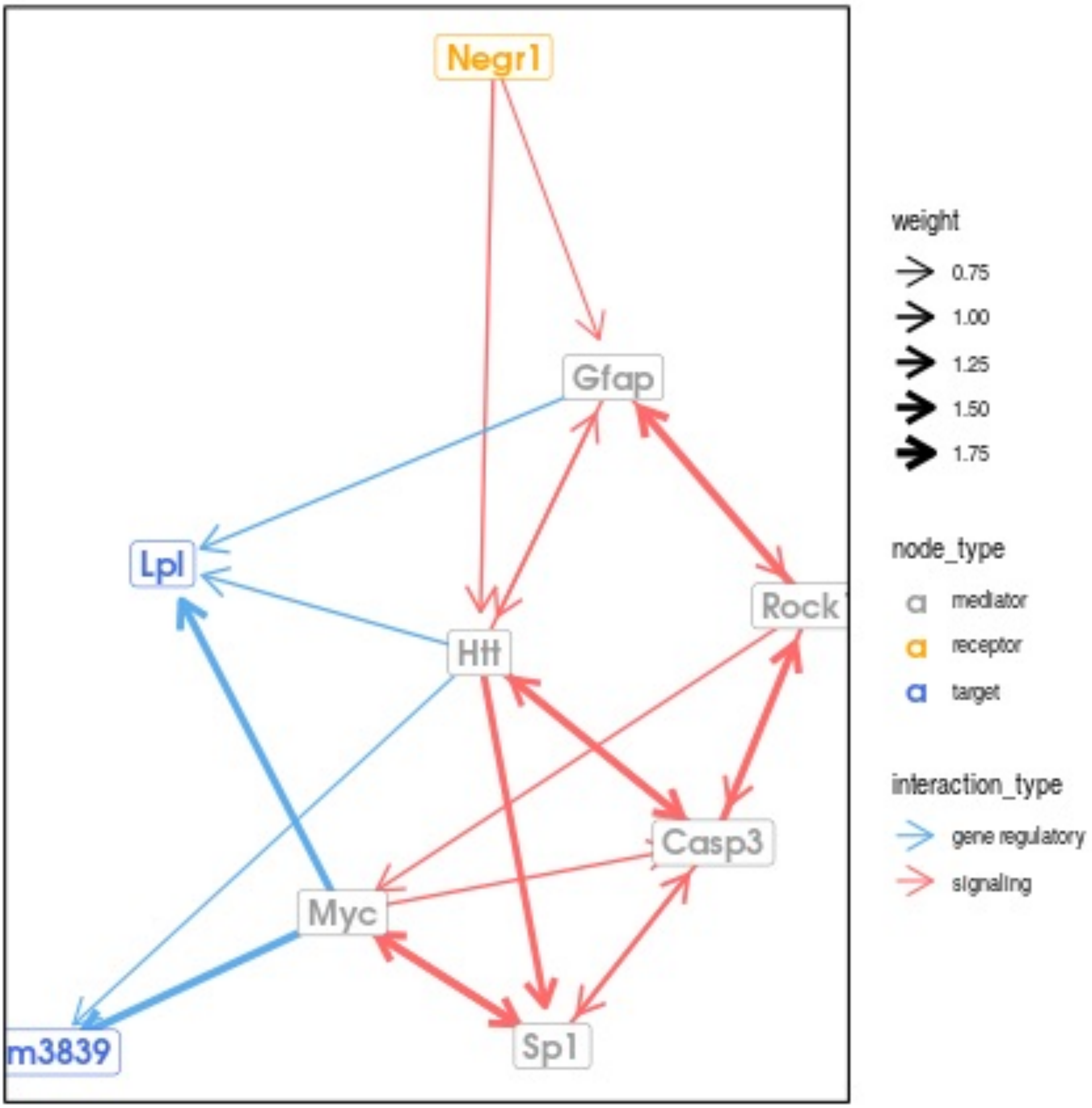
Negr1
Gfap
Lpl
Rock
Htt
Casp3
Myc
m3839
Sp1
weight
0.75
1.00
1.25
1.50
1.75
node_type
mediator
receptor
target
interaction_type
gene regulatory
signaling

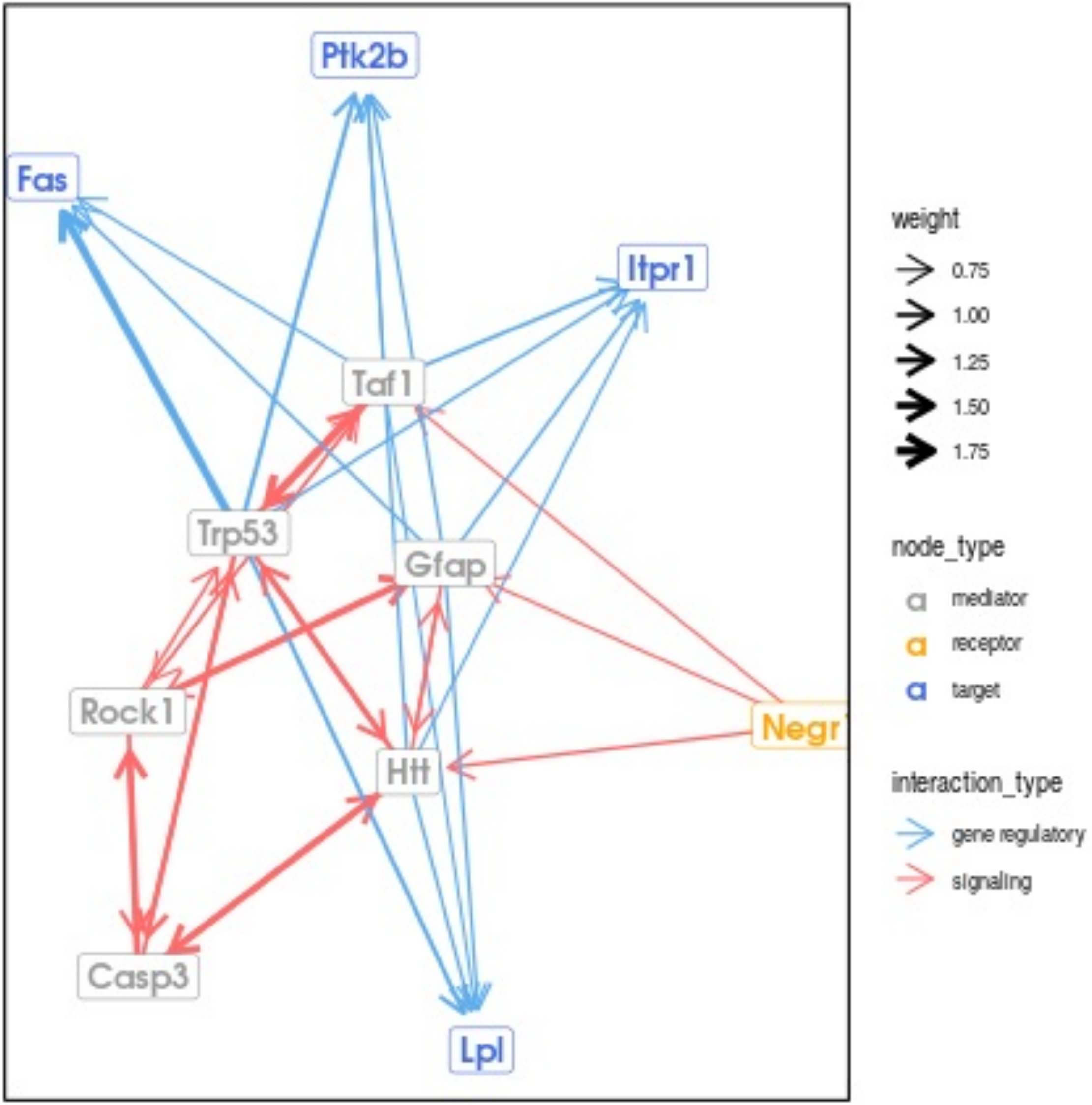

Ptk2b
Fas
Itpr1
Taf1
Trp53
Gfap
Rock1
Negr1
Htt
Casp3
Lpl
weight
0.75
1.00
1.25
1.50
1.75
node_type
mediator
receptor
target
interaction_type
gene regulatory
signaling

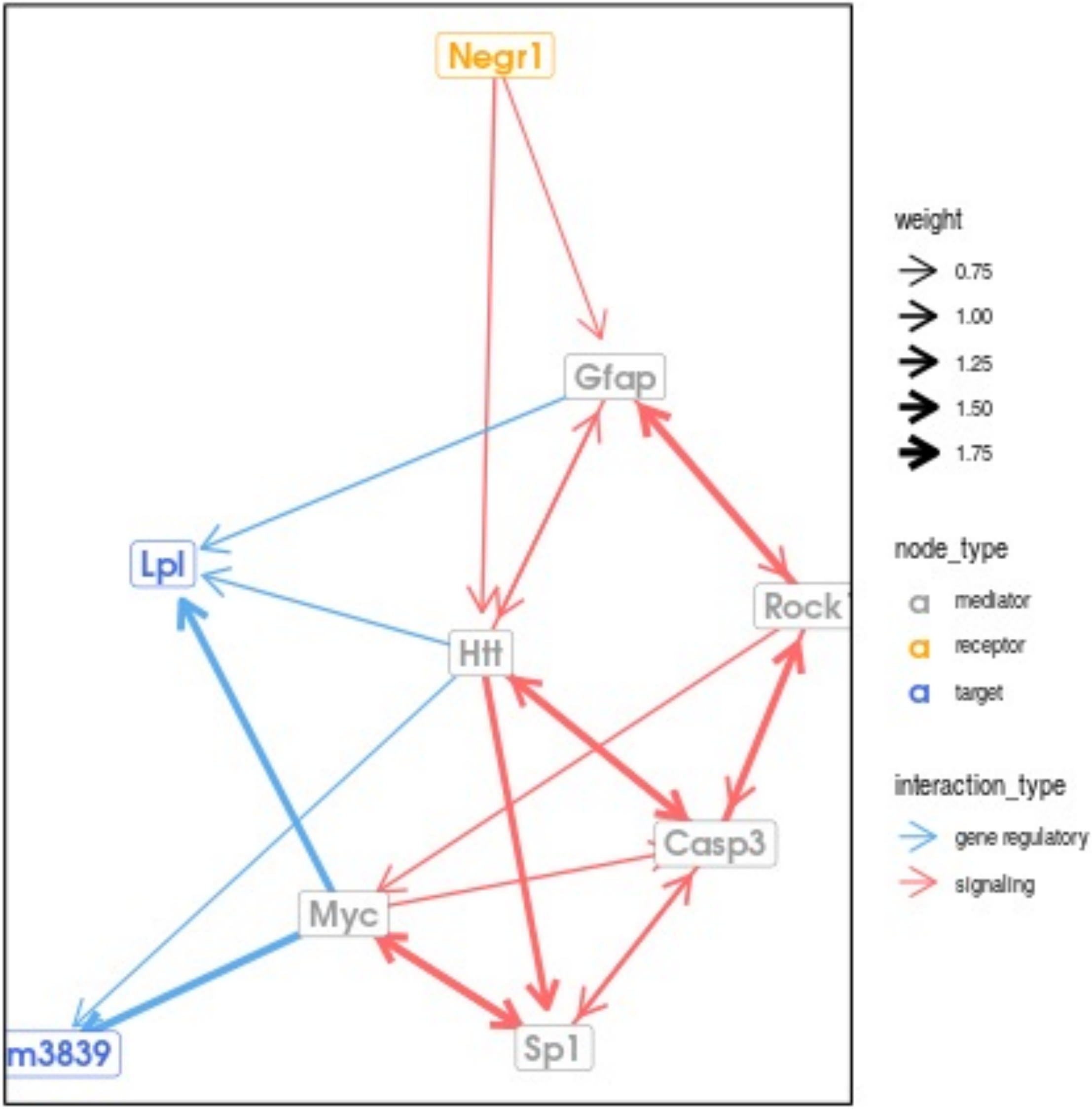

Negr1
Gfap
Lpl
Rock1
Htt
Casp3
Myc
m3839
Sp1
weight
0.75
1.00
1.25
1.50
1.75
node_type
mediator
receptor
target
interaction_type
gene regulatory
signaling

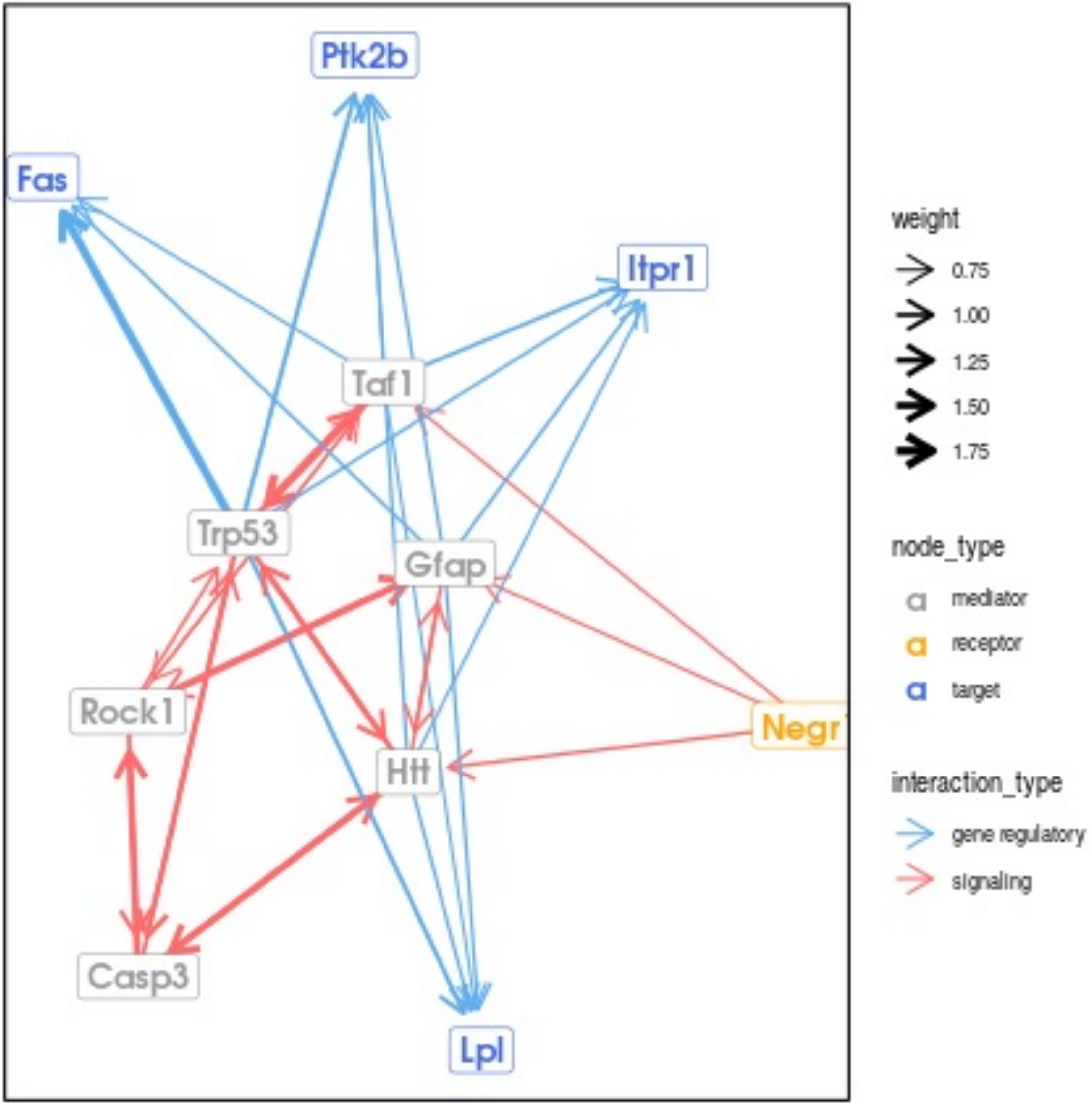

Ptk2b
Fas
Itpr1
Taf1
Trp53
Gfap
Rock1
Negr1
Htt
Casp3
Lpl
weight
0.75
1.00
1.25
1.50
1.75
node_type
mediator
receptor
target
interaction_type
gene regulatory
signaling

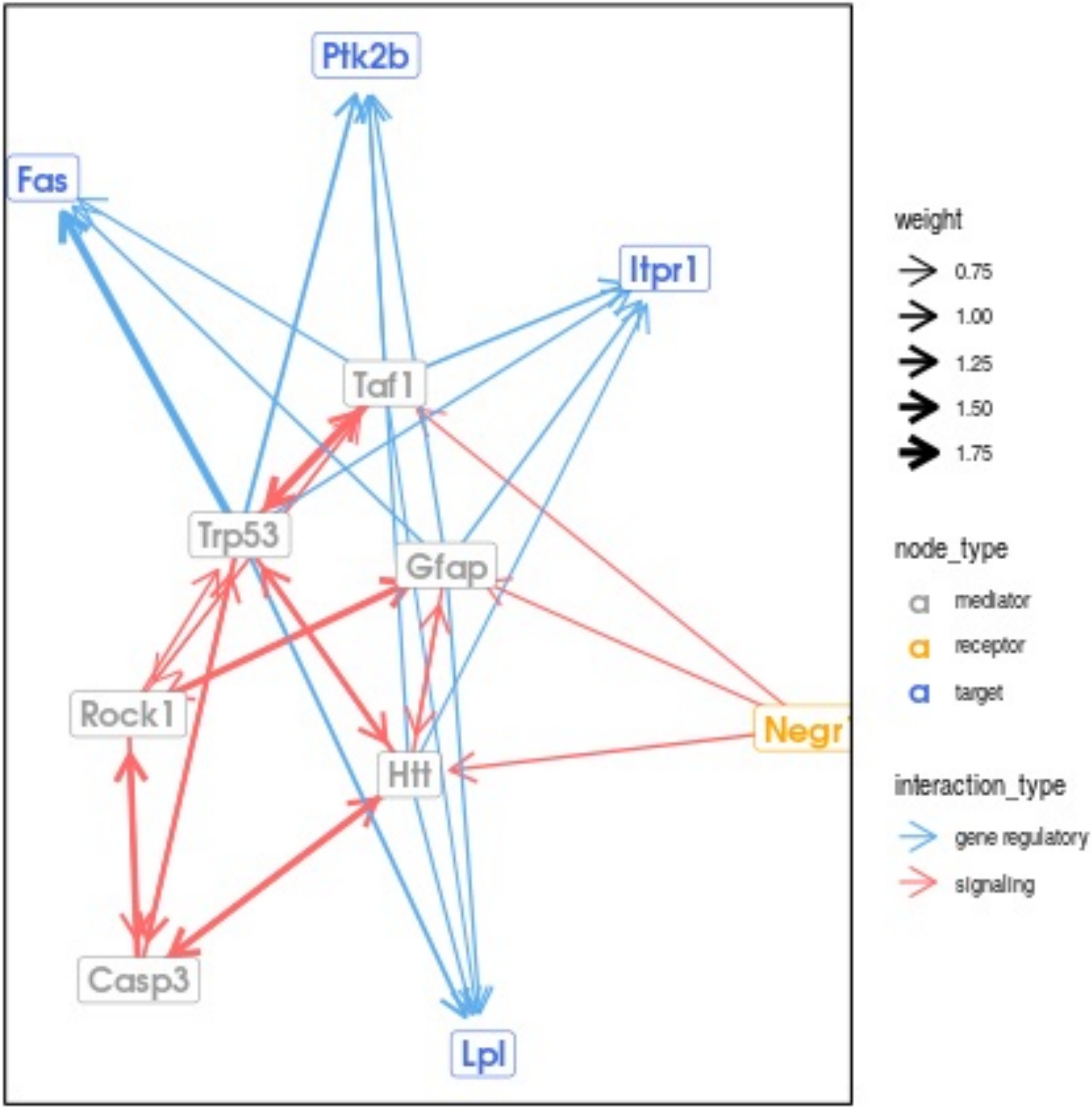
Ptk2b
Fas
Itpr1
Taf1
Trp53
Gfap
Rock1
Negr1
Htt
Casp3
Lpl
weight
0.75
1.00
1.25
1.50
1.75
node_type
mediator
receptor
target
interaction_type
gene regulatory
signaling

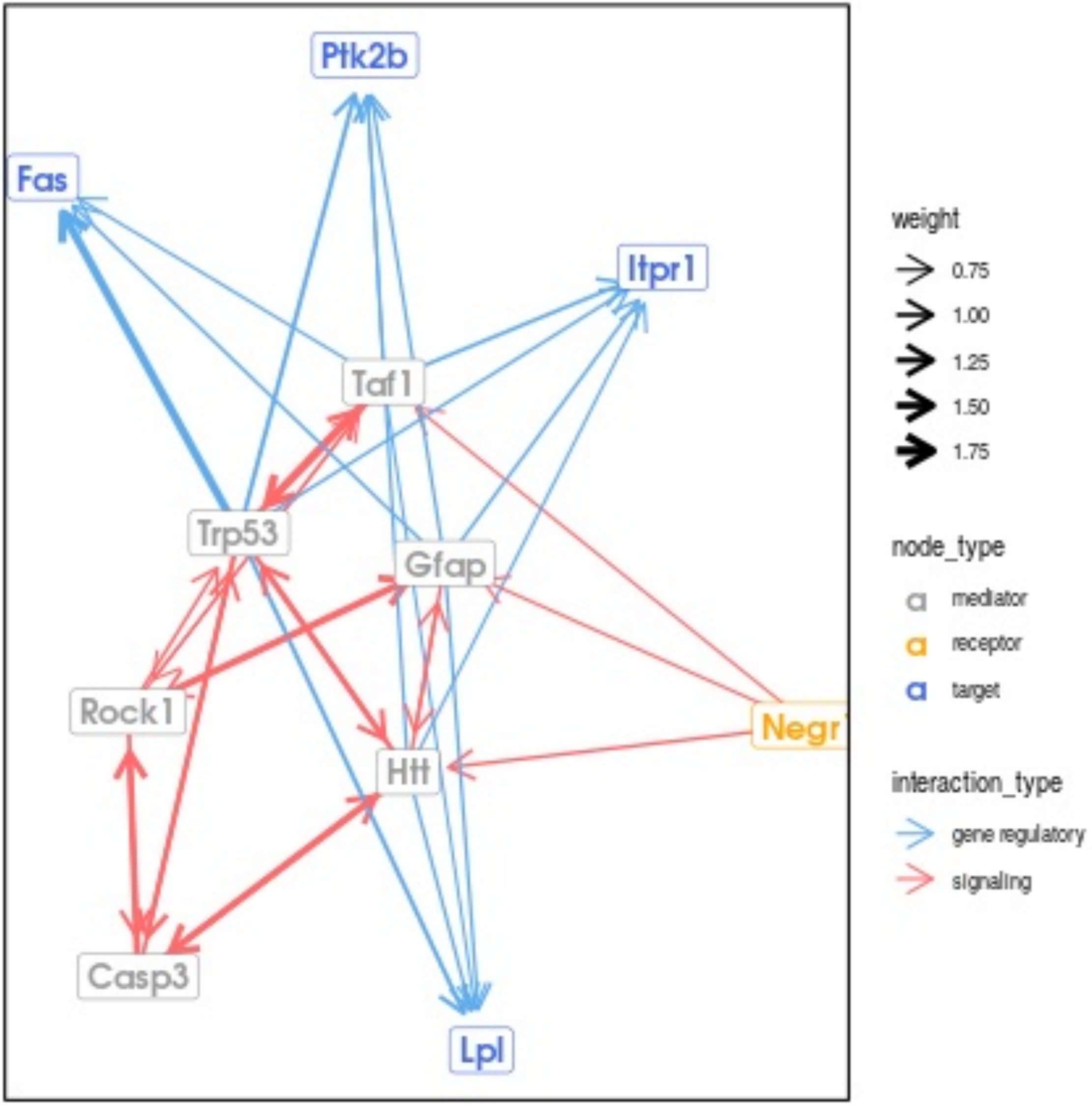

Ptk2b
Fas
Itpr1
Taf1
Trp53
Gfap
Rock1
Negr1
Htt
Casp3
Lpl
weight
0.75
1.00
1.25
1.50
1.75
node_type
mediator
receptor
target
interaction_type
gene regulatory
signaling

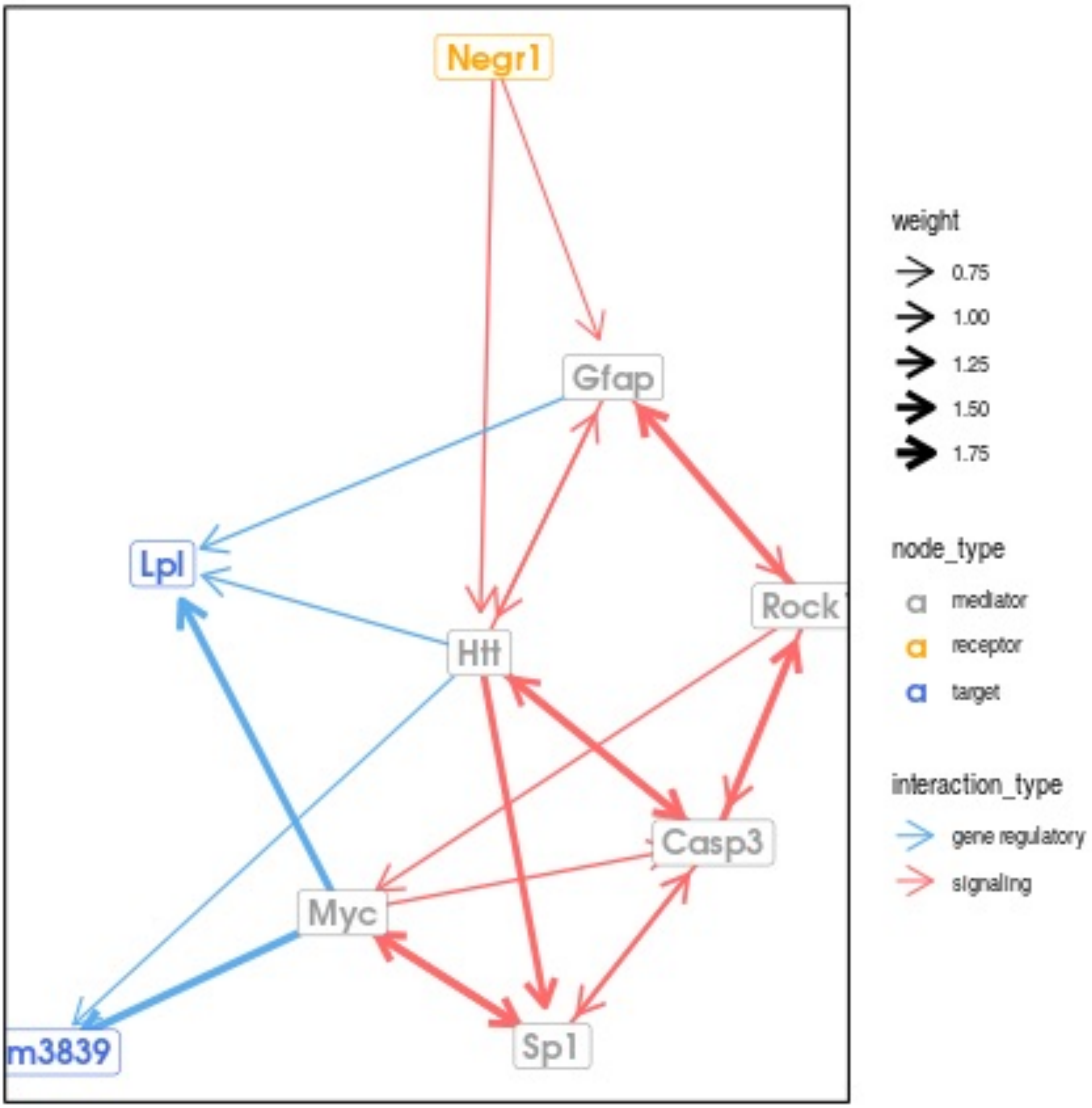

Negr1
Gfap
Lpl
Rock
Htt
Casp3
Myc
m3839
Sp1
weight
0.75
1.00
1.25
1.50
1.75
node_type
mediator
receptor
target
interaction_type
gene regulatory
signaling

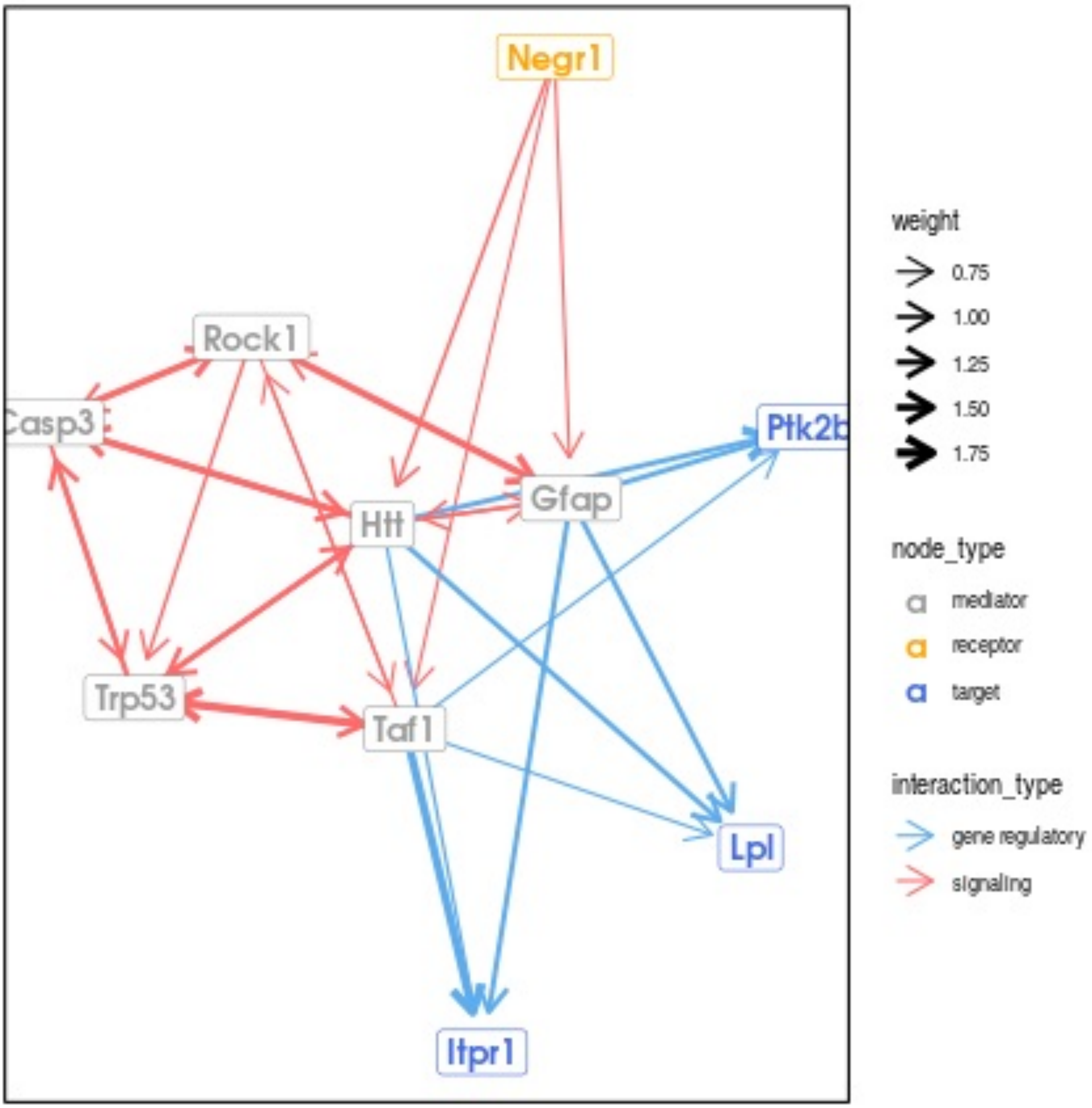
Negr1
Rock1
Casp3
Ptk2b
Gfap
Htt
Trp53
Taf1
Lpl
Itpr1
weight
0.75
1.00
1.25
1.50
1.75
node_type
mediator
receptor
target
interaction_type
gene regulatory
signaling

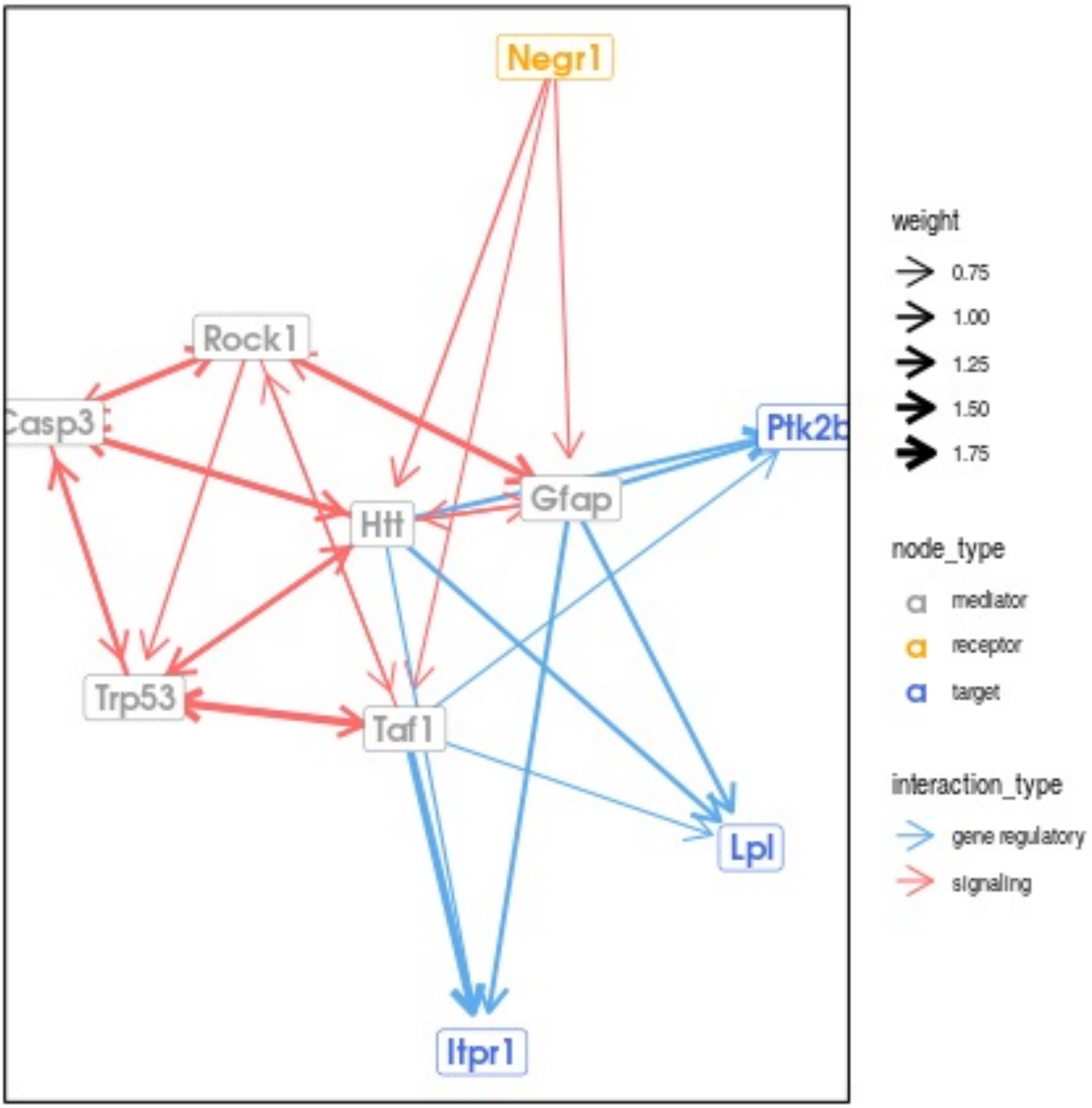
Negr1
Rock1
Casp3
Ptk2b
Gfap
Htt
Trp53
Taf1
Lpl
Itpr1
weight
0.75
1.00
1.25
1.50
1.75
node_type
mediator
receptor
target
interaction_type
gene regulatory
signaling

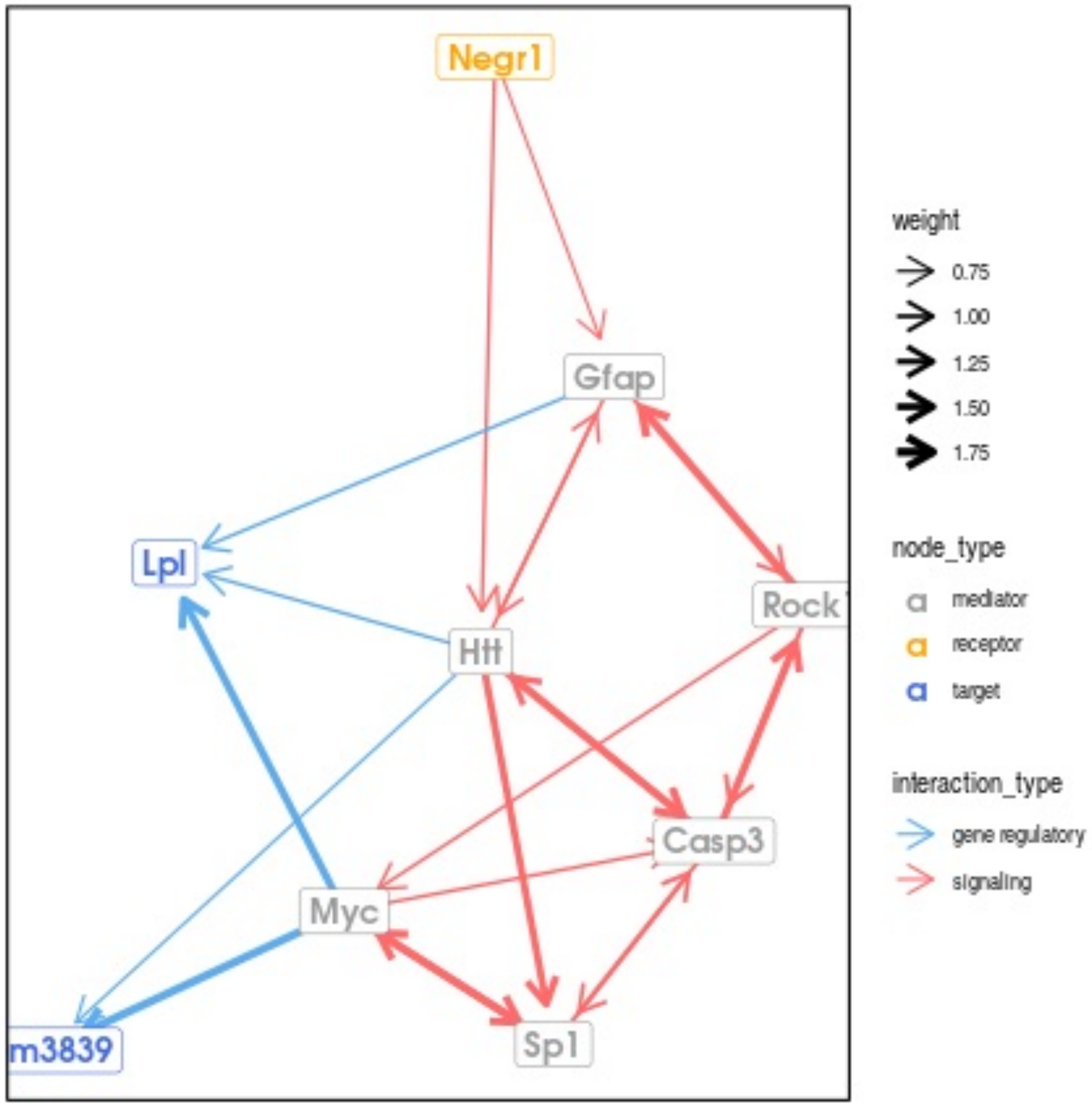

Negr1
Gfap
Lpl
Htt
Rock1
Casp3
Myc
Sp1
m3839
weight
0.75
1.00
1.25
1.50
1.75
node_type
mediator
receptor
target
interaction_type
gene regulatory
signaling

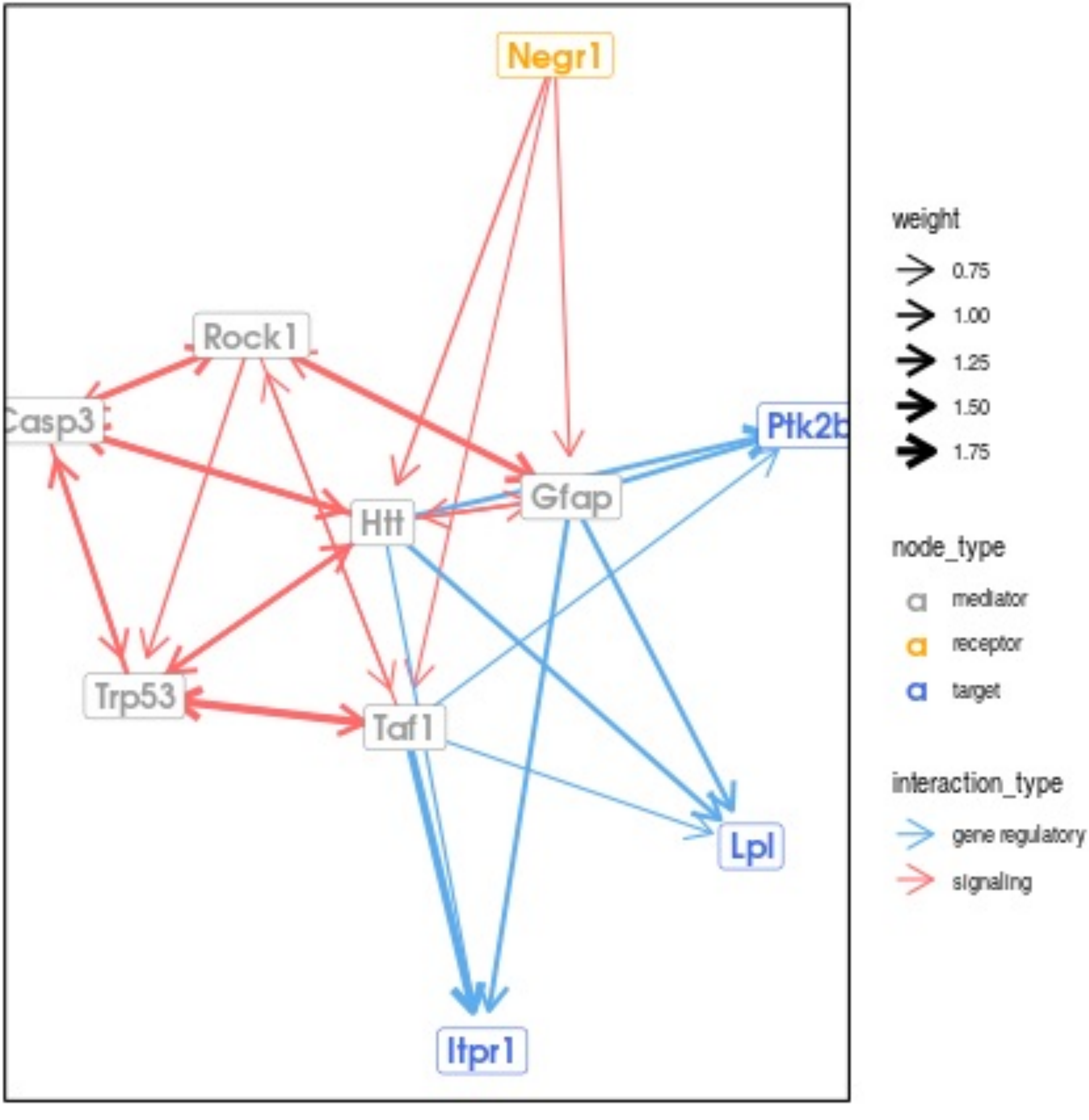
Negr1
Rock1
Casp3
Ptk2b
Gfap
Htt
Trp53
Taf1
Lpl
Itpr1
weight
0.75
1.00
1.25
1.50
1.75
node_type
mediator
receptor
target
interaction_type
gene regulatory
signaling

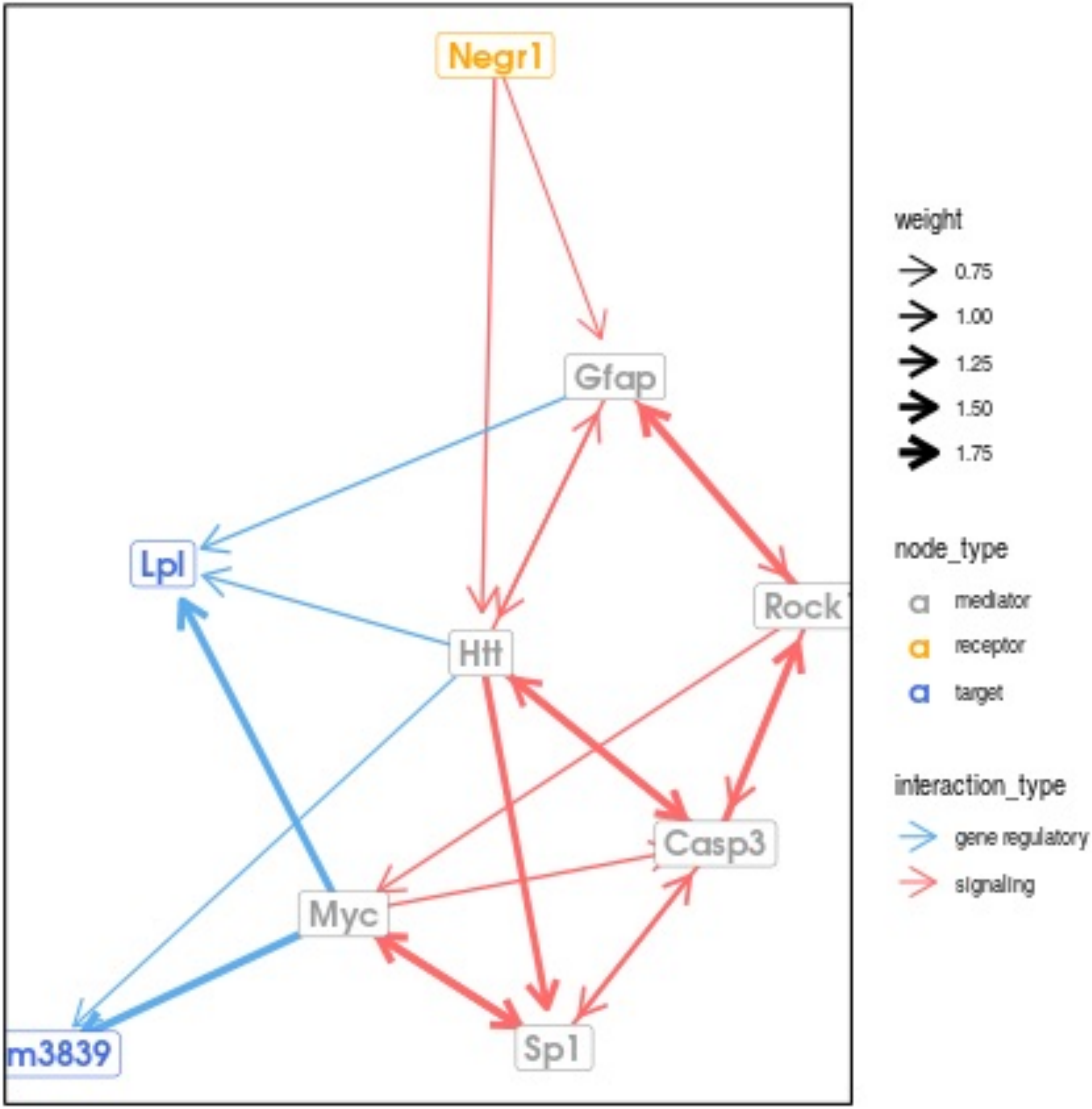
Negr1
Gfap
Lpl
Rock1
Htt
Casp3
Myc
m3839
Sp1
weight
0.75
1.00
1.25
1.50
1.75
node_type
mediator
receptor
target
interaction_type
gene regulatory
signaling

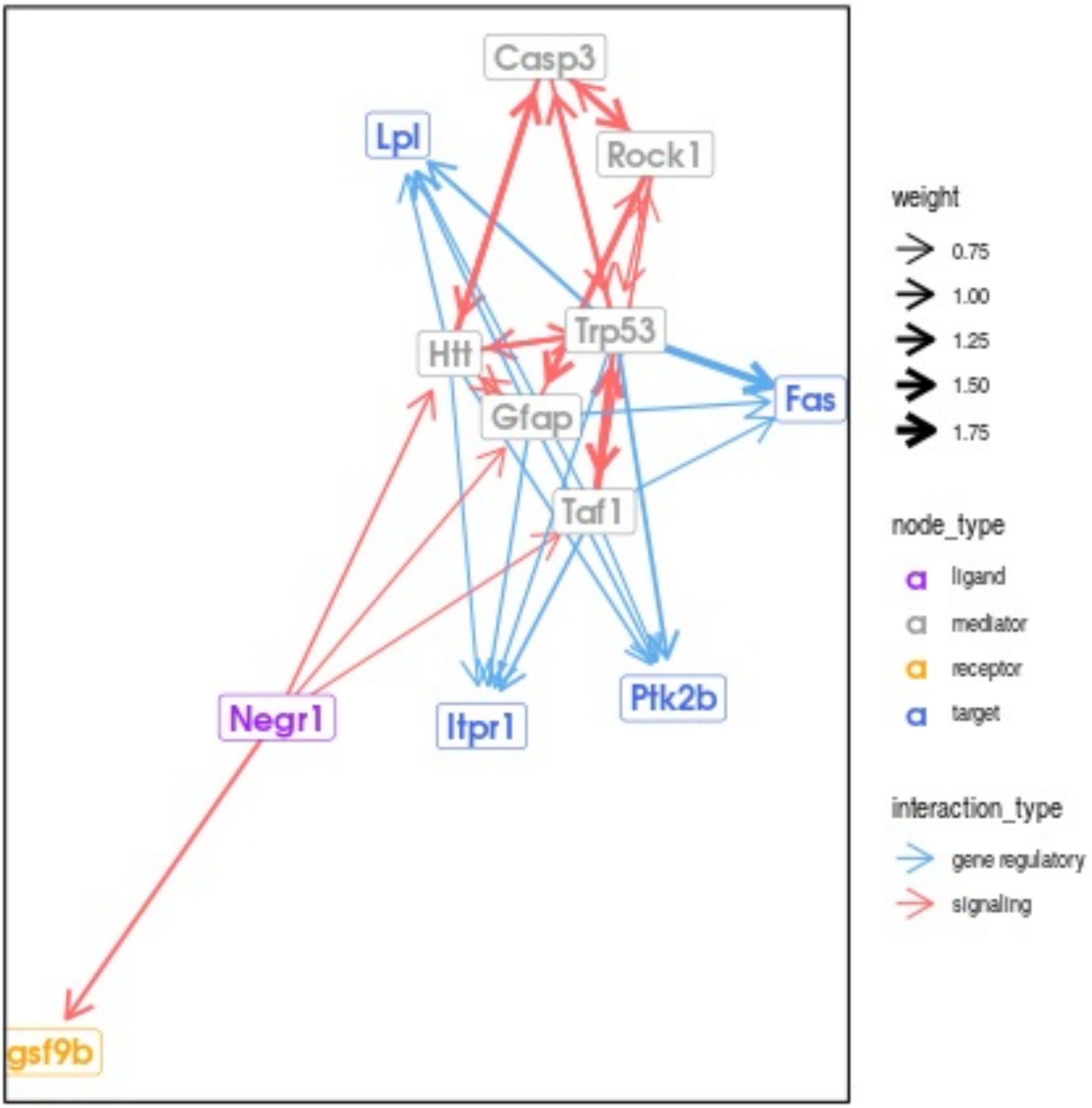
Casp3
Lpl
Rock1
Trp53
Htt
Gfap
Fas
Taf1
Negr1
Itpr1
Ptk2b
gsf9b
weight
0.75
1.00
1.25
1.50
1.75
node_type
ligand
mediator
receptor
target
interaction_type
gene regulatory
signaling

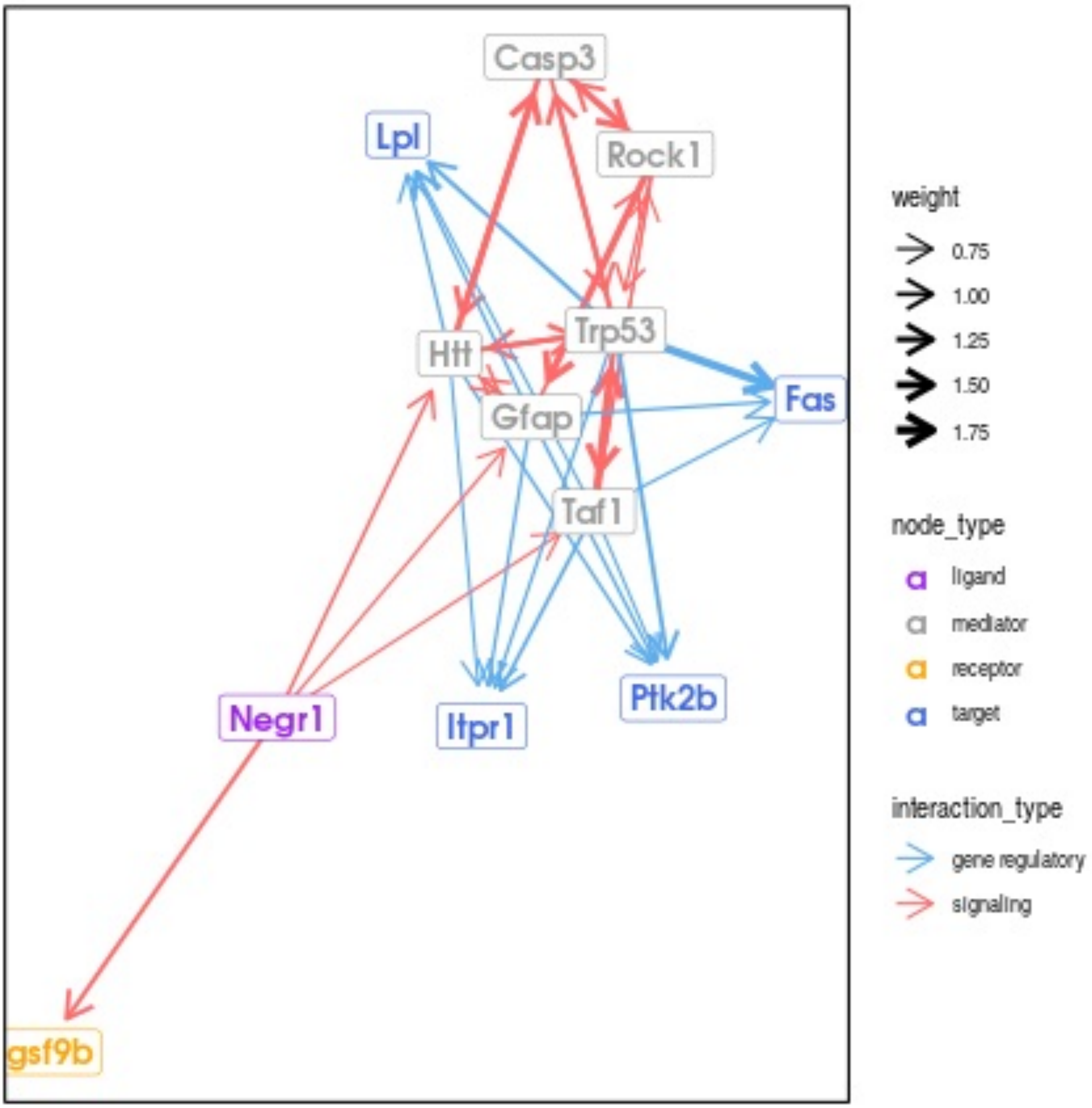
Casp3
Lpl
Rock1
Trp53
Htt
Gfap
Fas
Taf1
Negr1
Itpr1
Ptk2b
gsf9b
weight
0.75
1.00
1.25
1.50
1.75
node_type
ligand
mediator
receptor
target
interaction_type
gene regulatory
signaling

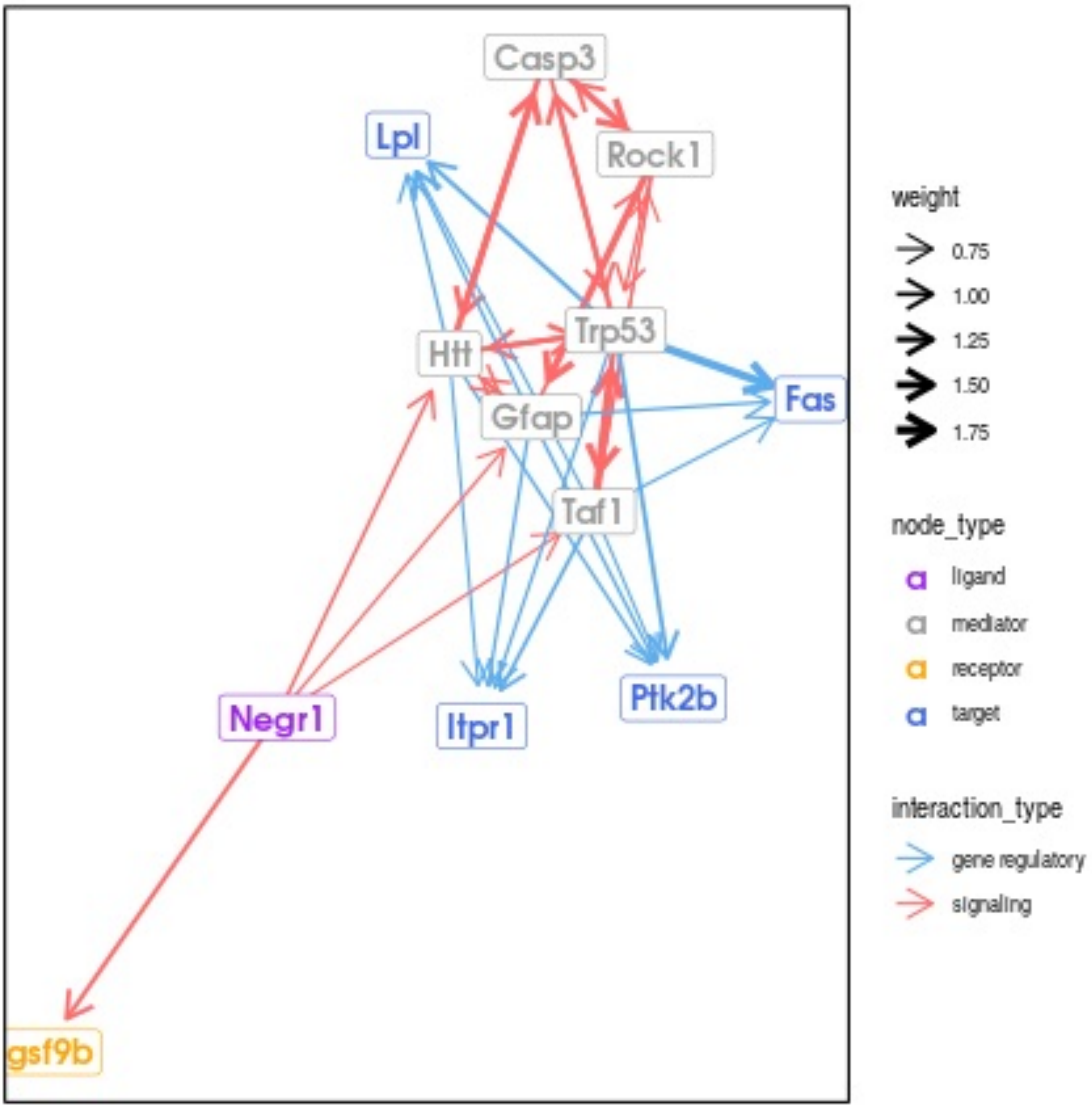

Casp3
Lpl
Rock1
Trp53
Htt
Gfap
Fas
Taf1
Negr1
Itpr1
Ptk2b
gsf9b
weight
0.75
1.00
1.25
1.50
1.75
node_type
ligand
mediator
receptor
target
interaction_type
gene regulatory
signaling

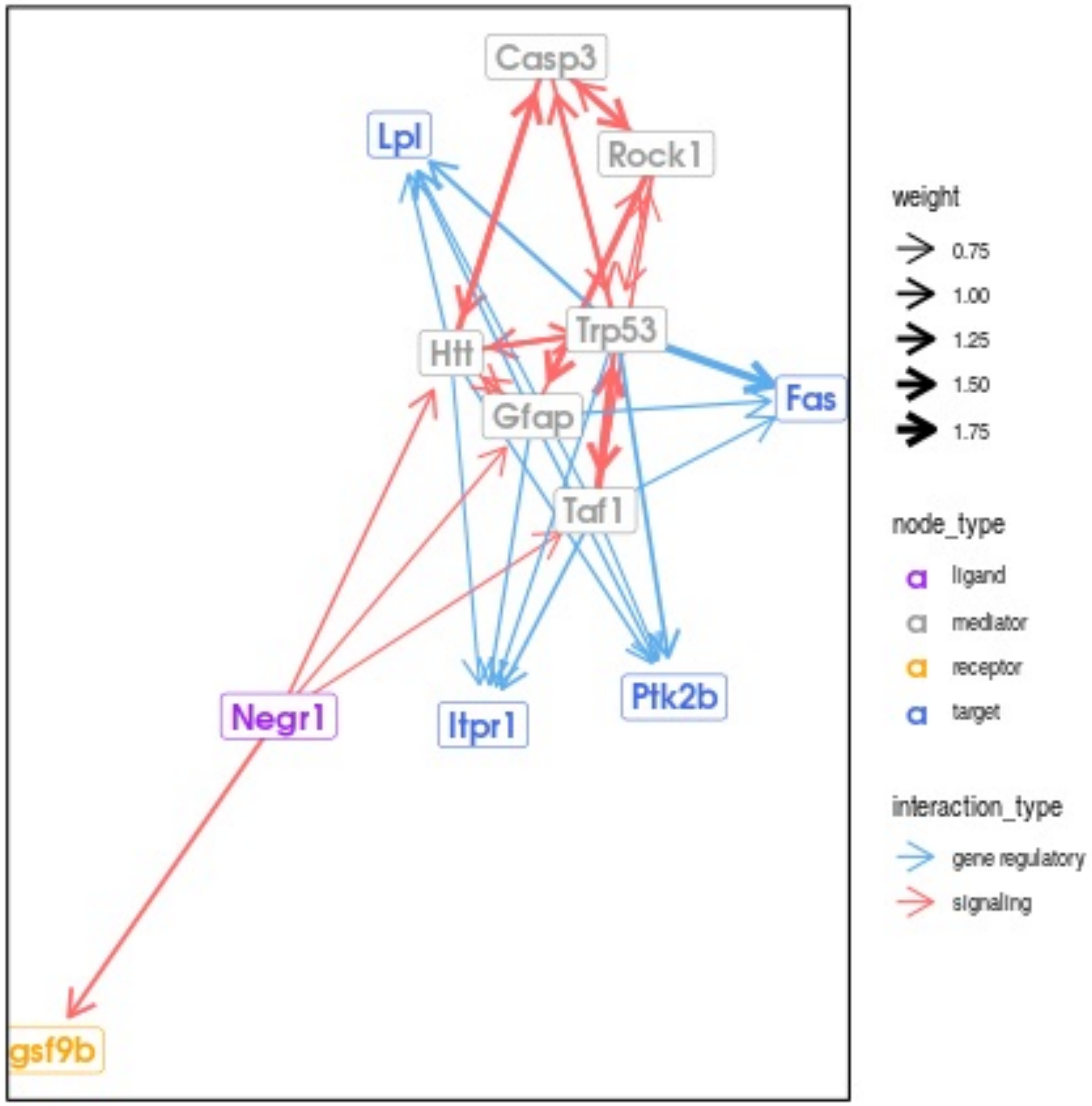

Casp3
Lpl
Rock1
Trp53
Htt
Gfap
Fas
Taf1
Negr1
Itpr1
Ptk2b
gsf9b
weight
0.75
1.00
1.25
1.50
1.75
node_type
ligand
mediator
receptor
target
interaction_type
gene regulatory
signaling

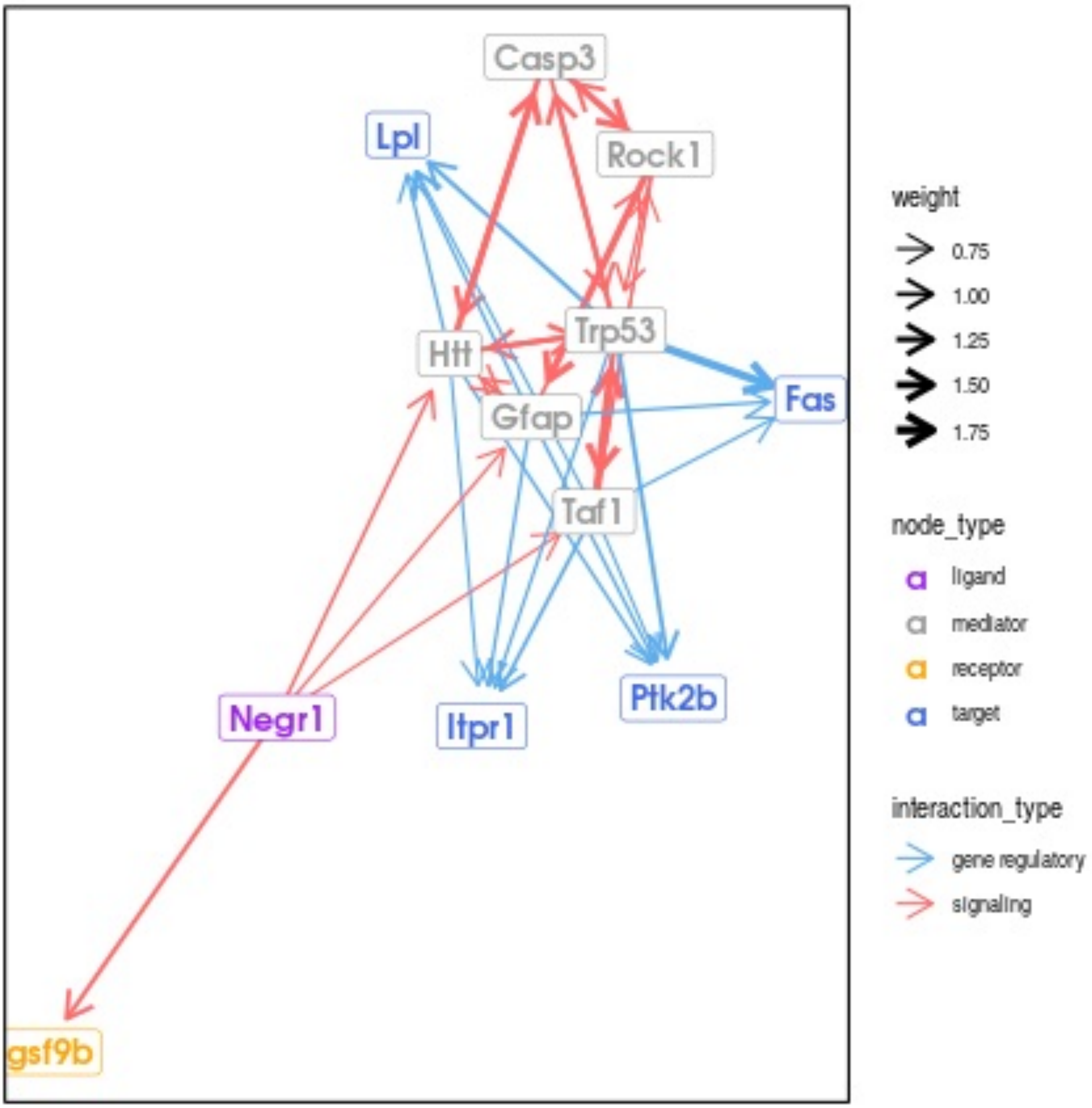

Casp3
Lpl
Rock1
Trp53
Htt
Gfap
Fas
Taf1
Negr1
Itpr1
Ptk2b
gsf9b
weight
0.75
1.00
1.25
1.50
1.75
node_type
ligand
mediator
receptor
target
interaction_type
gene regulatory
signaling

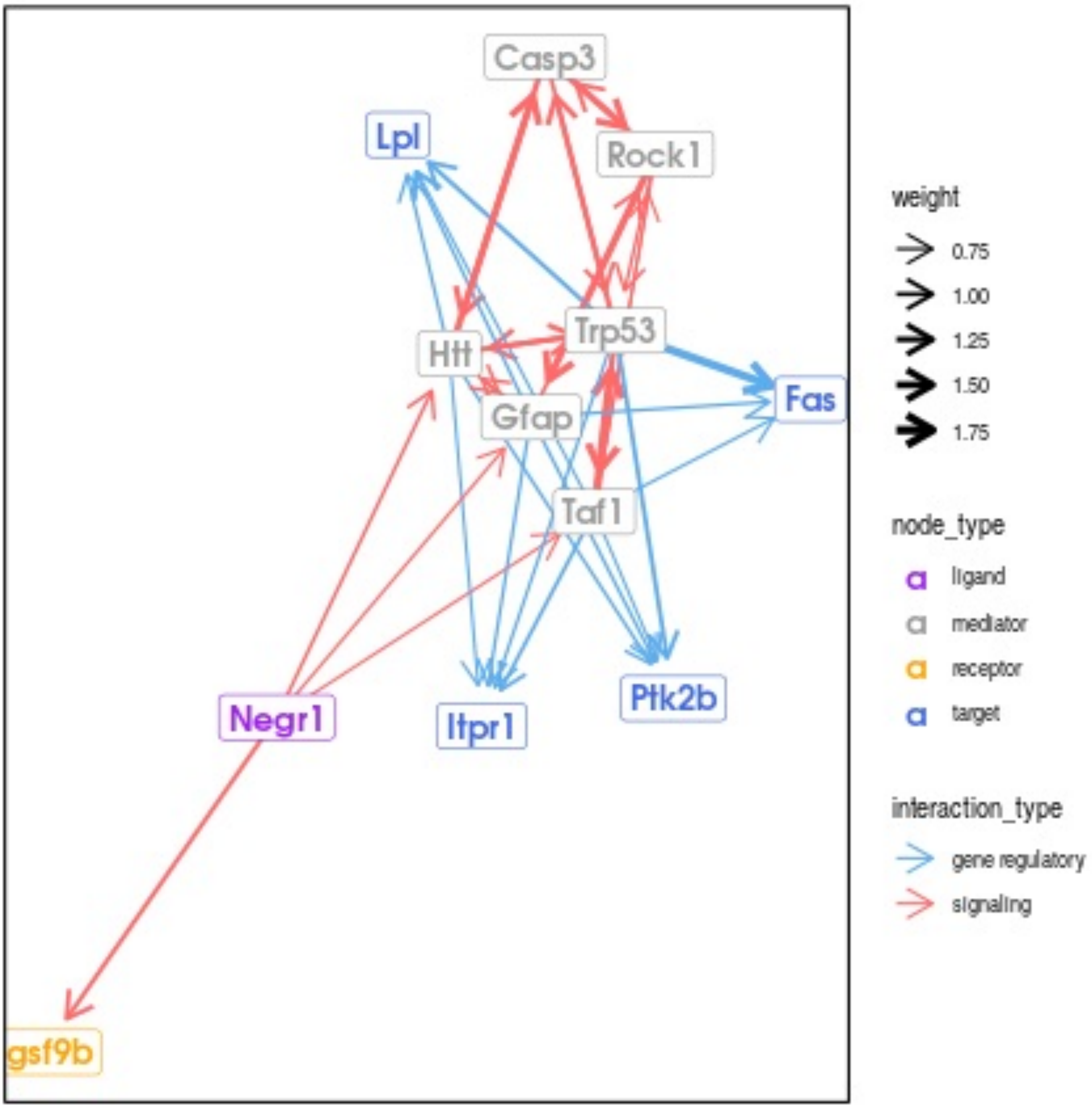
Casp3
Lpl
Rock1
Trp53
Htt
Gfap
Fas
Taf1
Negr1
Itpr1
Ptk2b
gsf9b
weight
0.75
1.00
1.25
1.50
1.75
node_type
ligand
mediator
receptor
target
interaction_type
gene regulatory
signaling

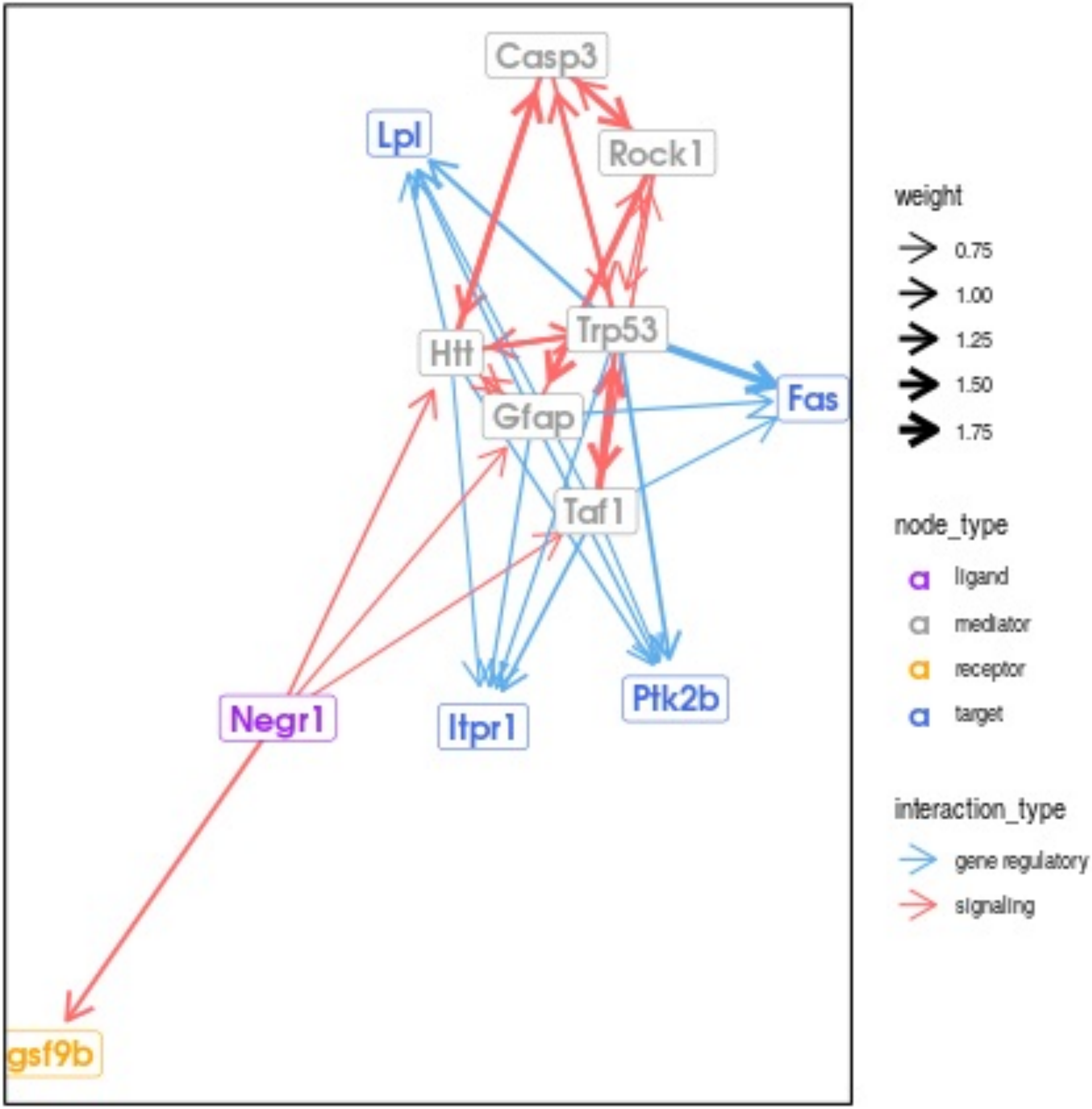

Casp3
Lpl
Rock1
Trp53
Htt
Gfap
Fas
Taf1
Negr1
Itpr1
Ptk2b
gsf9b
weight
0.75
1.00
1.25
1.50
1.75
node_type
ligand
mediator
receptor
target
interaction_type
gene regulatory
signaling

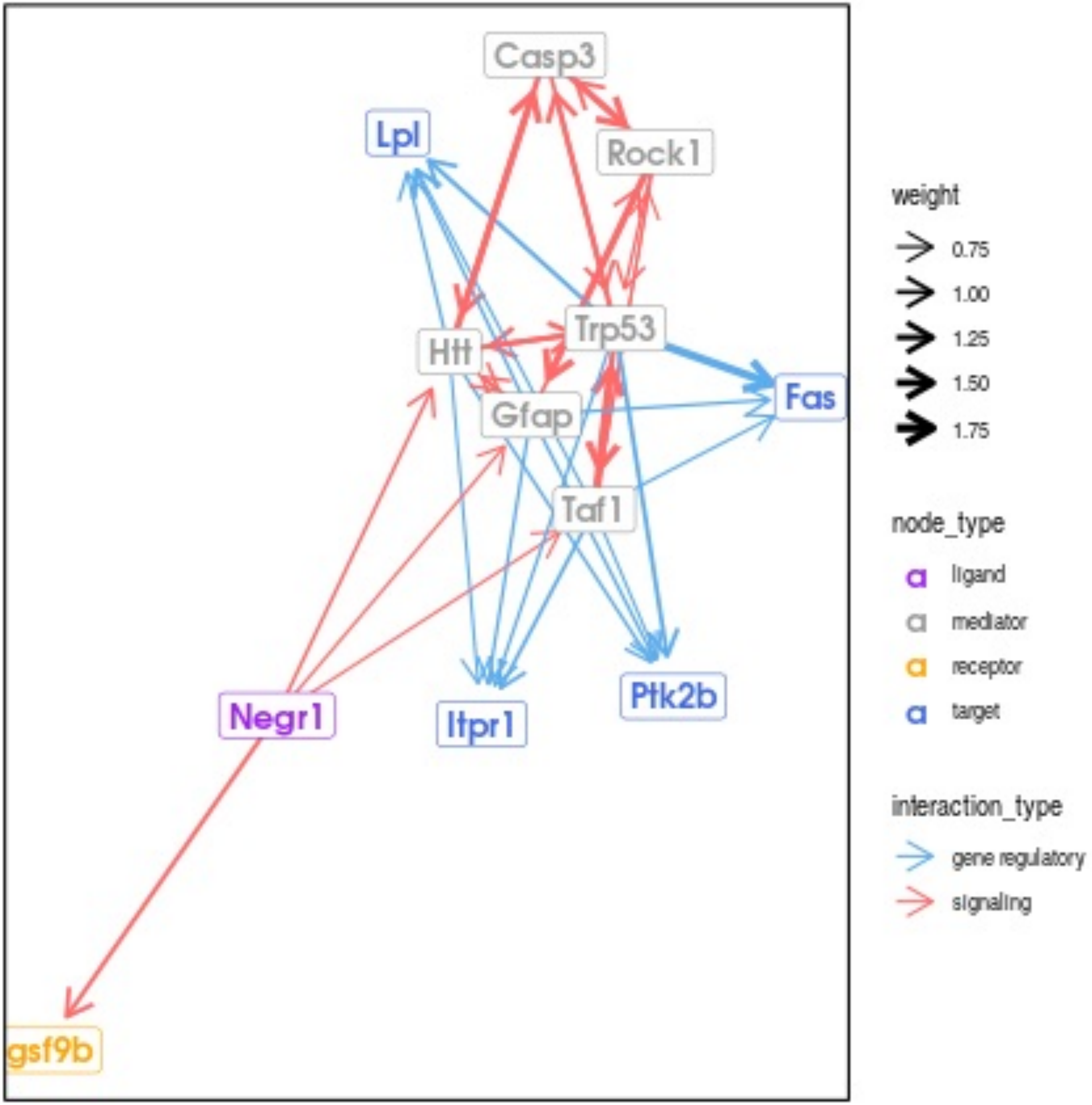

Casp3
Lpl
Rock1
Trp53
Htt
Gfap
Fas
Taf1
Negr1
Itpr1
Ptk2b
gsf9b
weight
0.75
1.00
1.25
1.50
1.75
node_type
ligand
mediator
receptor
target
interaction_type
gene regulatory
signaling

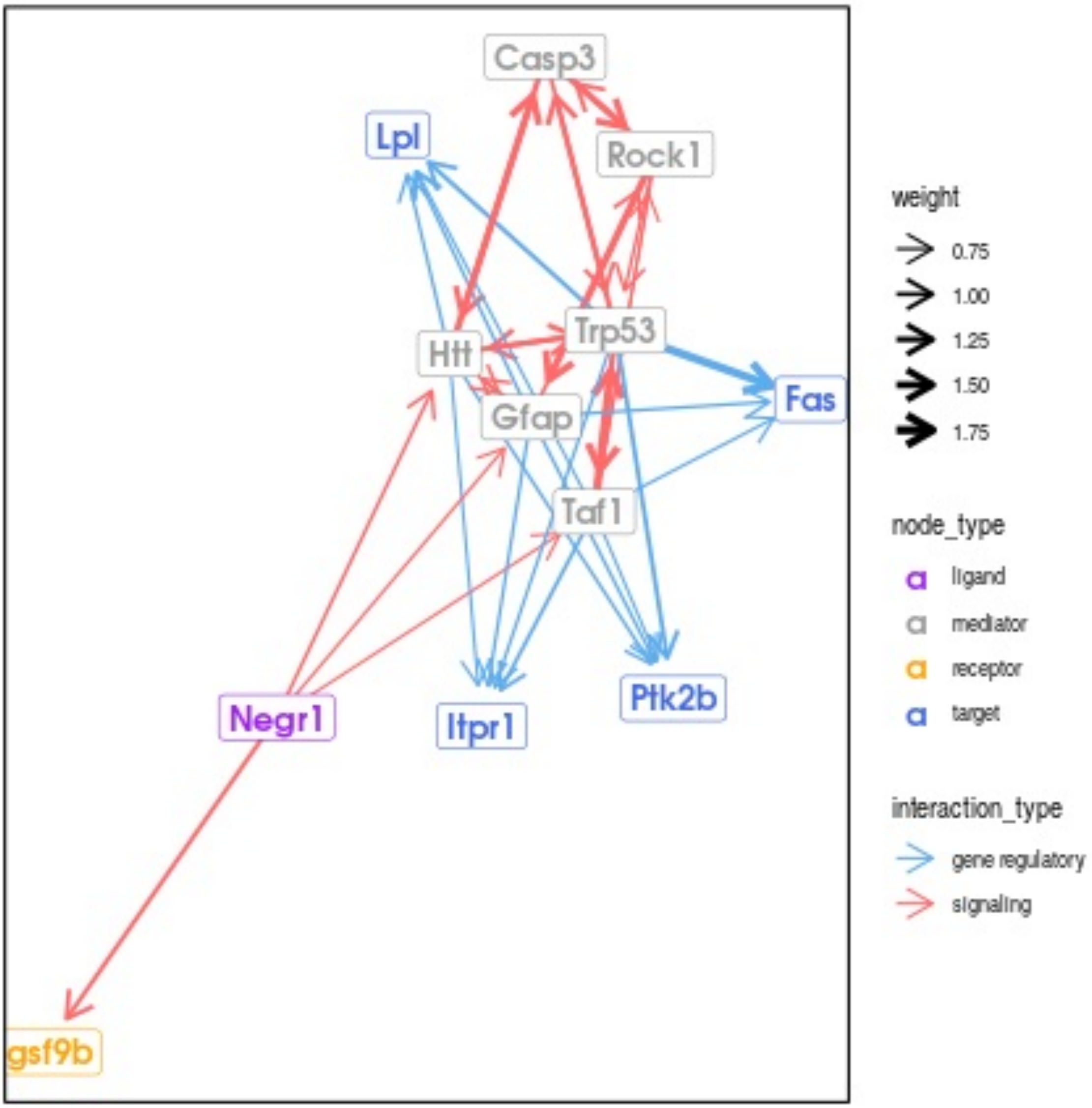

Casp3
Lpl
Rock1
Trp53
Htt
Fas
Gfap
Taf1
Negr1
Itpr1
Ptk2b
gsf9b
weight
0.75
1.00
1.25
1.50
1.75
node_type
ligand
mediator
receptor
target
interaction_type
gene regulatory
signaling

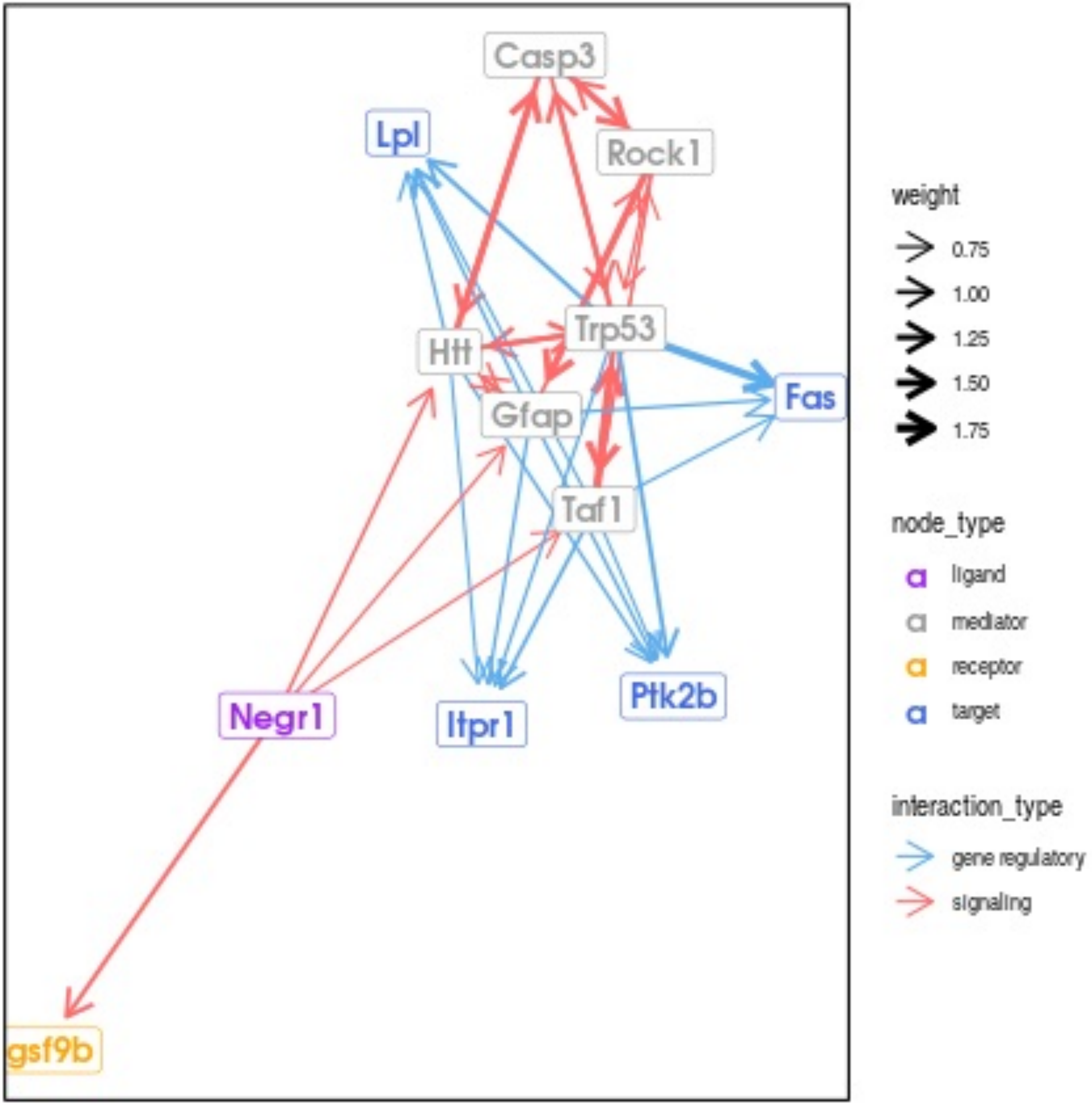

Casp3
Lpl
Rock1
Trp53
Htt
Gfap
Fas
Taf1
Negr1
Itpr1
Ptk2b
gsf9b
weight
0.75
1.00
1.25
1.50
1.75
node_type
ligand
mediator
receptor
target
interaction_type
gene regulatory
signaling

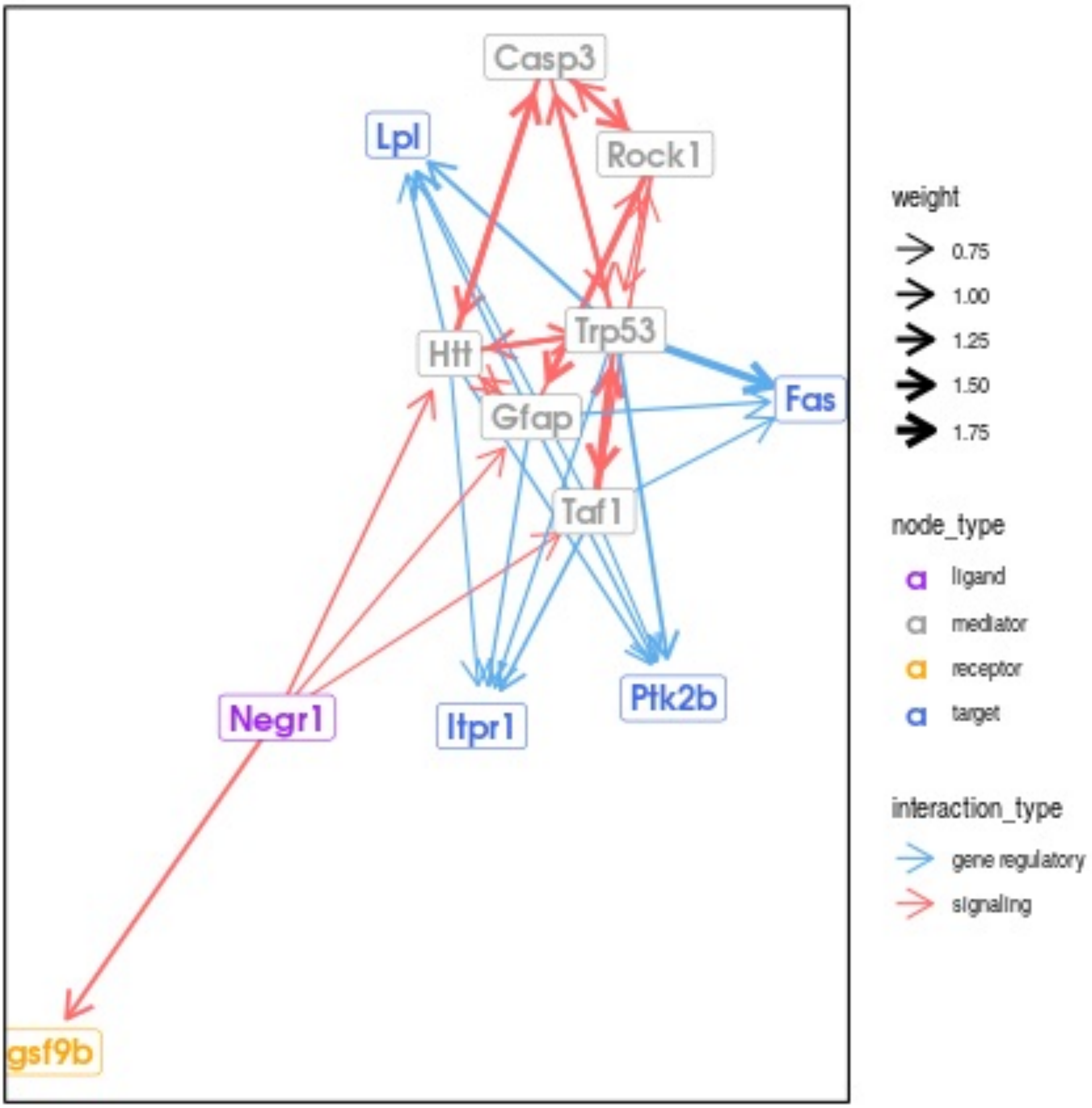
Casp3
Lpl
Rock1
Trp53
Htt
Gfap
Fas
Taf1
Negr1
Itpr1
Ptk2b
gsf9b
weight
0.75
1.00
1.25
1.50
1.75
node_type
ligand
mediator
receptor
target
interaction_type
gene regulatory
signaling

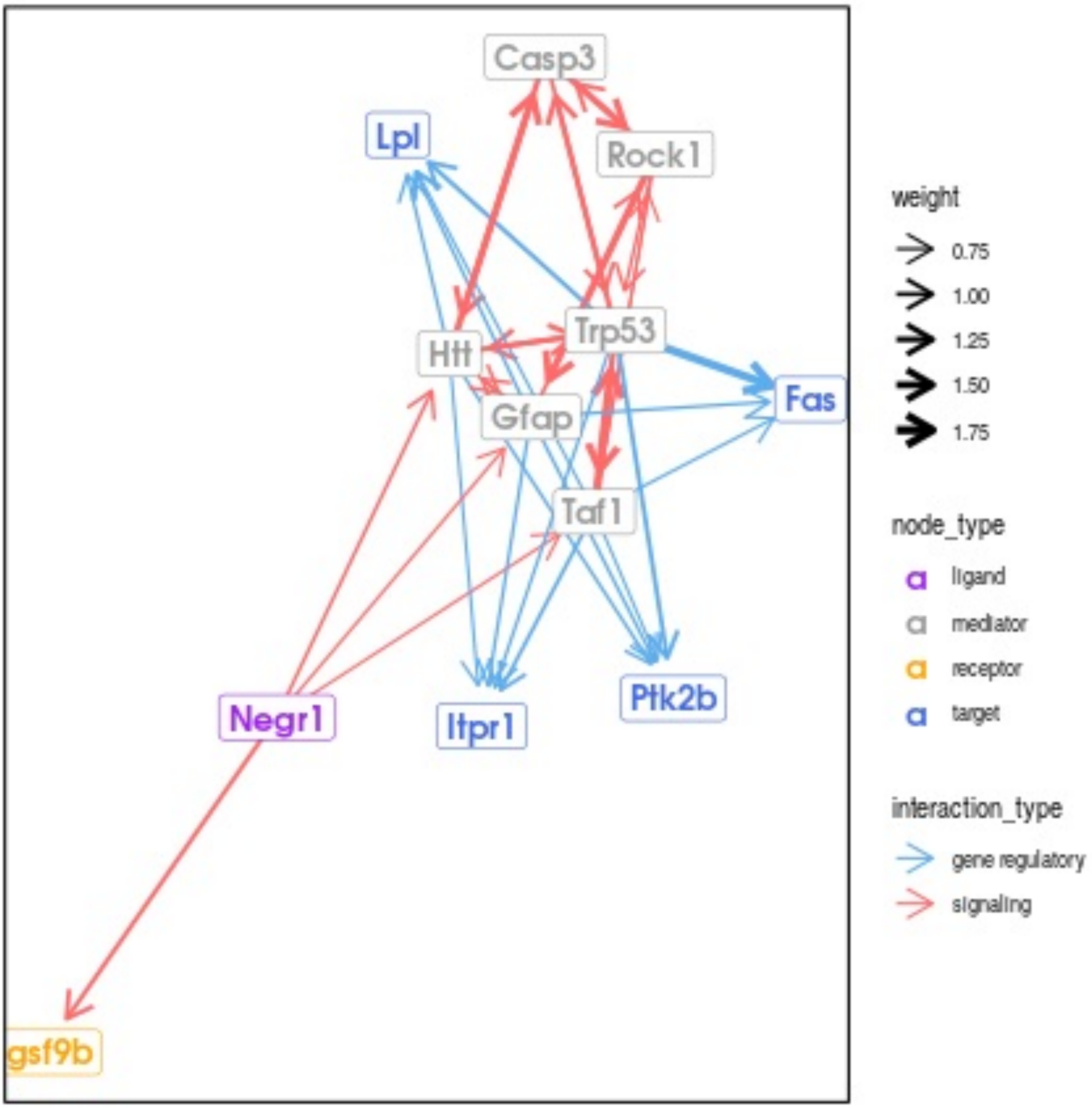

Casp3
Lpl
Rock1
Trp53
Htt
Gfap
Fas
Taf1
Negr1
Itpr1
Ptk2b
gsf9b
weight
0.75
1.00
1.25
1.50
1.75
node_type
ligand
mediator
receptor
target
interaction_type
gene regulatory
signaling

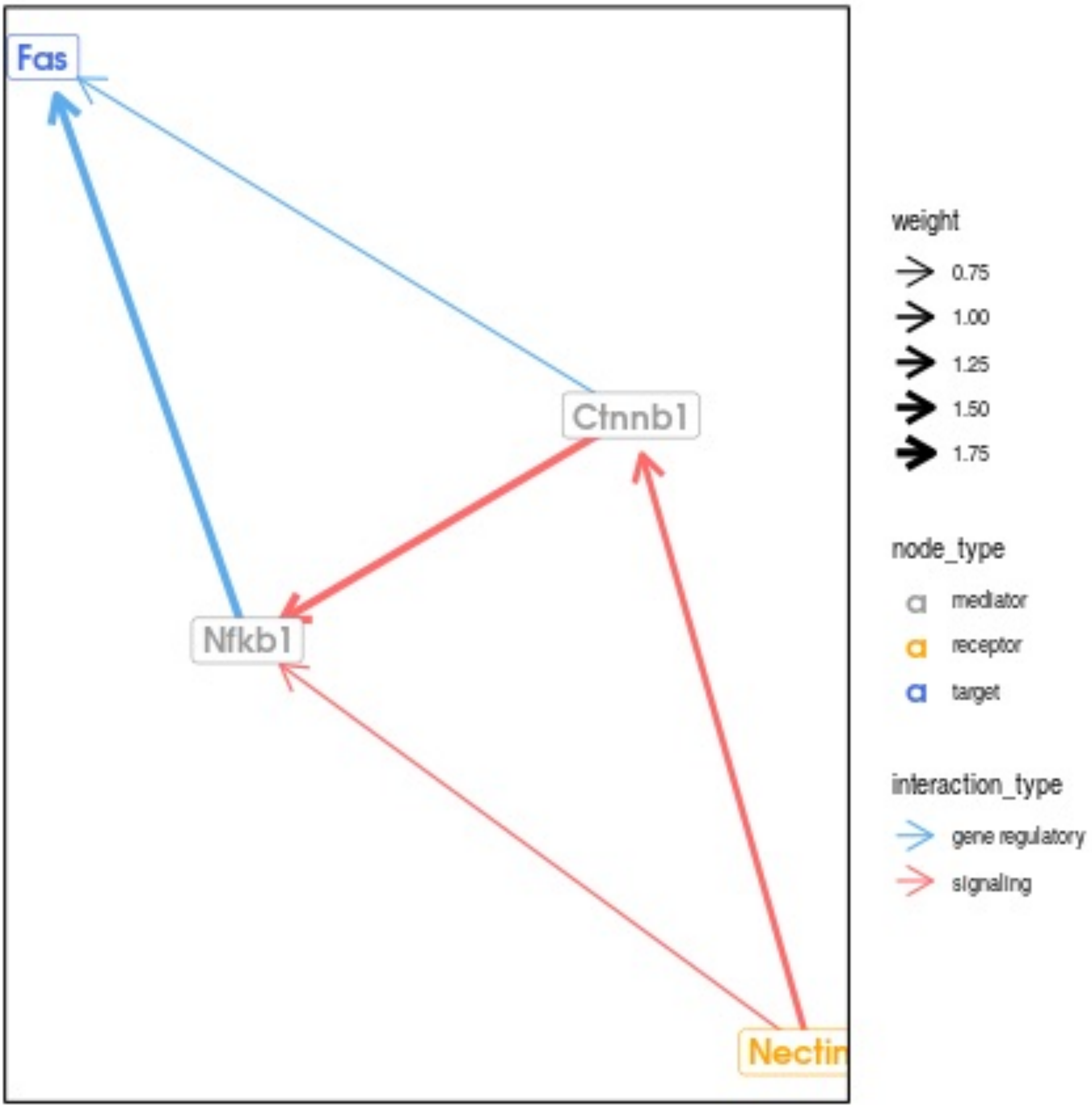
Fas
Ctnnb1
Nfkb1
Nectin
weight
0.75
1.00
1.25
1.50
1.75
node_type
mediator
receptor
target
interaction_type
gene regulatory
signaling

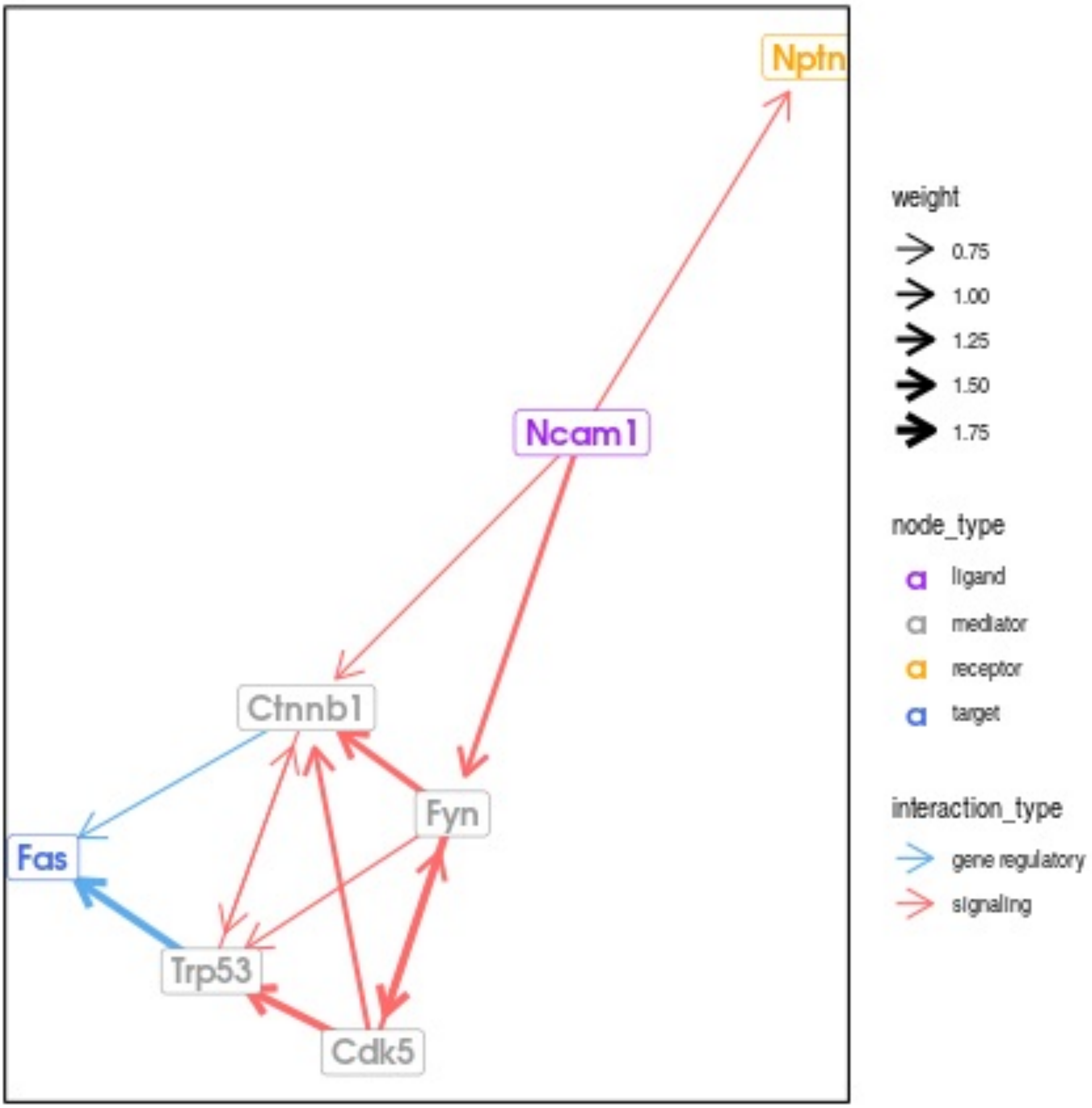

Nptn
Ncam1
Ctnnb1
Fyn
Fas
Trp53
Cdk5
weight
0.75
1.00
1.25
1.50
1.75
node_type
ligand
mediator
receptor
target
interaction_type
gene regulatory
signaling

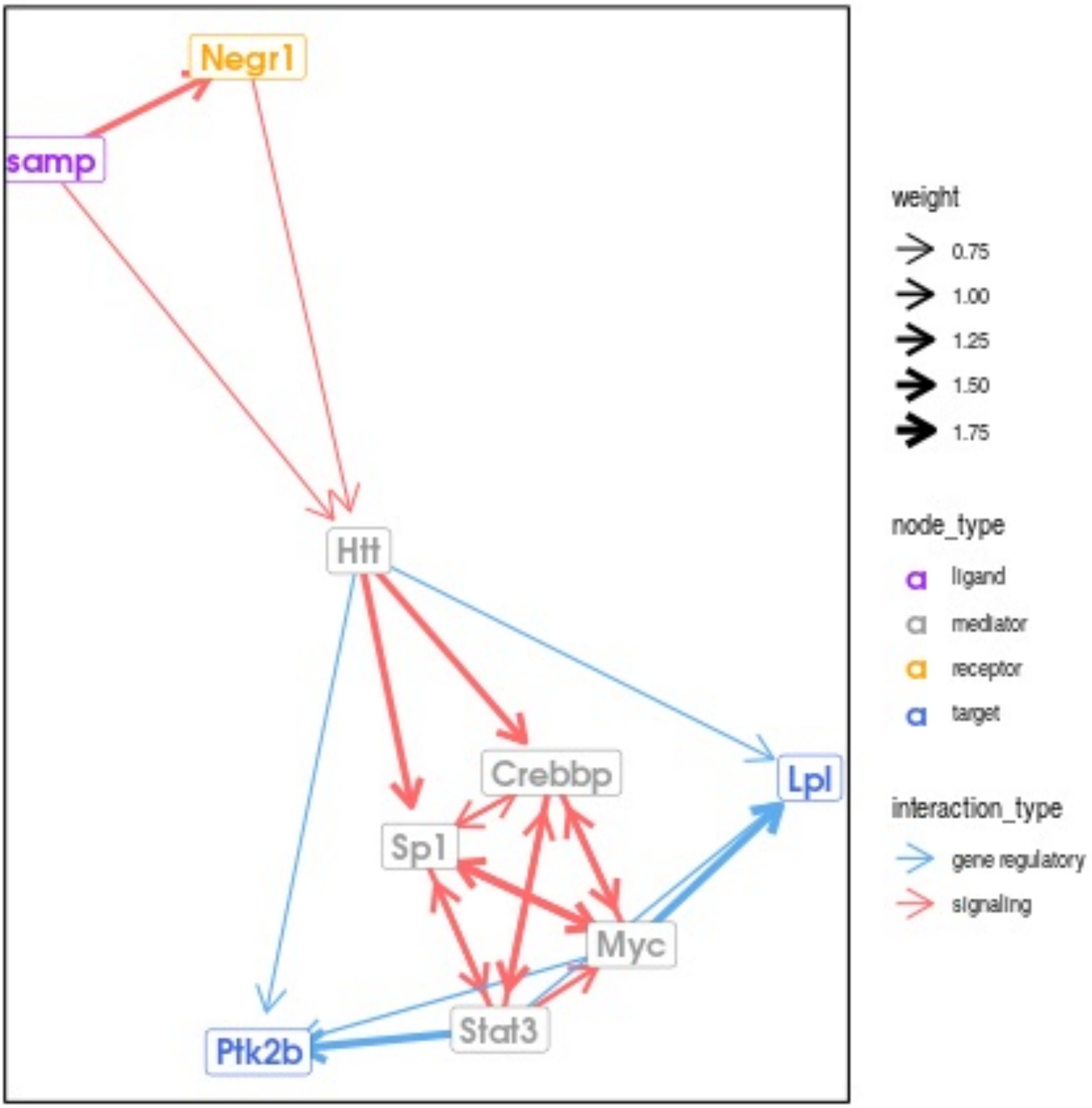

Negr1
samp
Htt
Crebbp
Lpl
Sp1
Myc
Stat3
Ptk2b
weight
0.75
1.00
1.25
1.50
1.75
node_type
ligand
mediator
receptor
target
interaction_type
gene regulatory
signaling

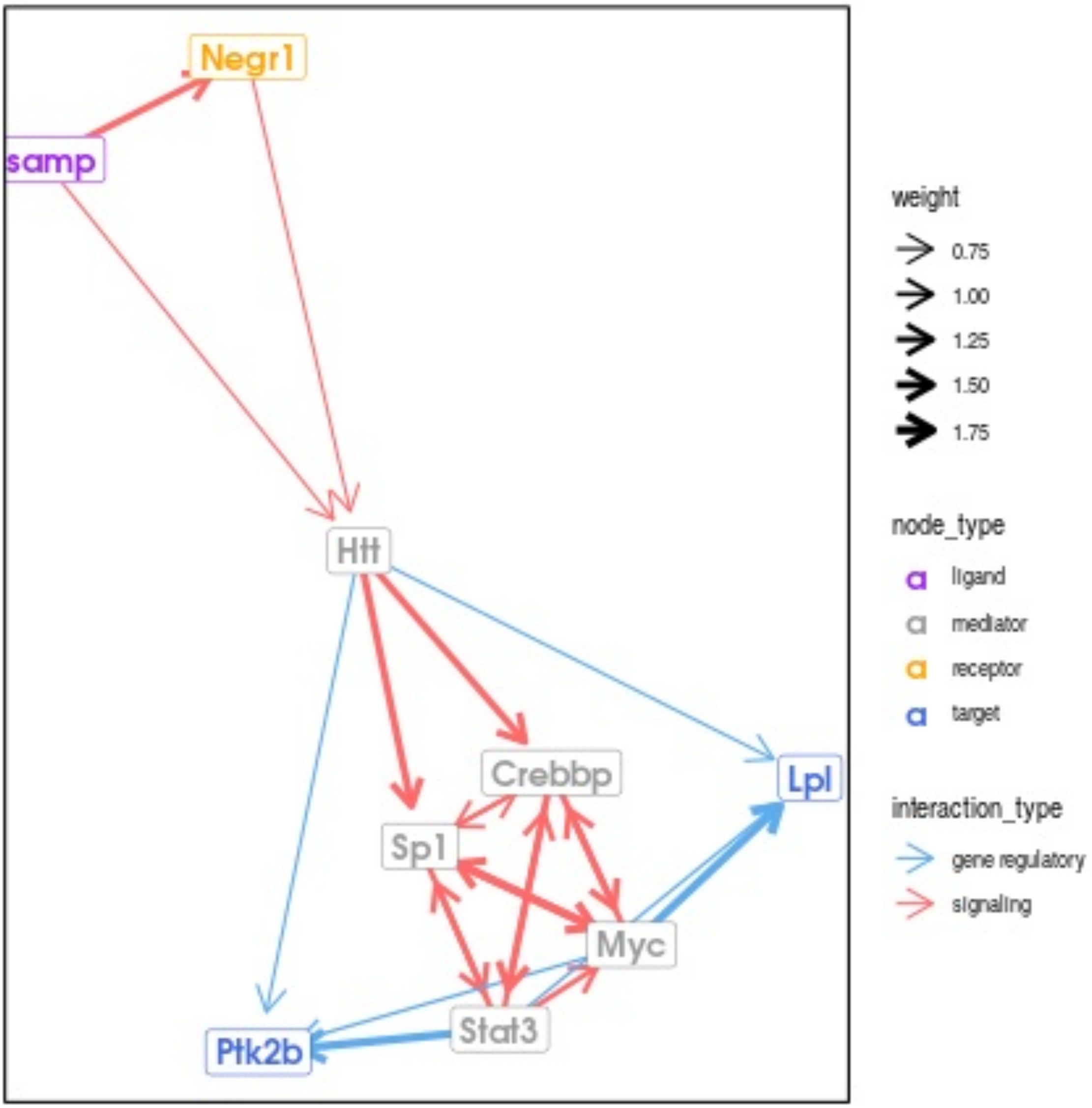

Negr1
samp
Htt
Crebbp
Sp1
Myc
Lpl
Stat3
Ptk2b
weight
0.75
1.00
1.25
1.50
1.75
node_type
ligand
mediator
receptor
target
interaction_type
gene regulatory
signaling

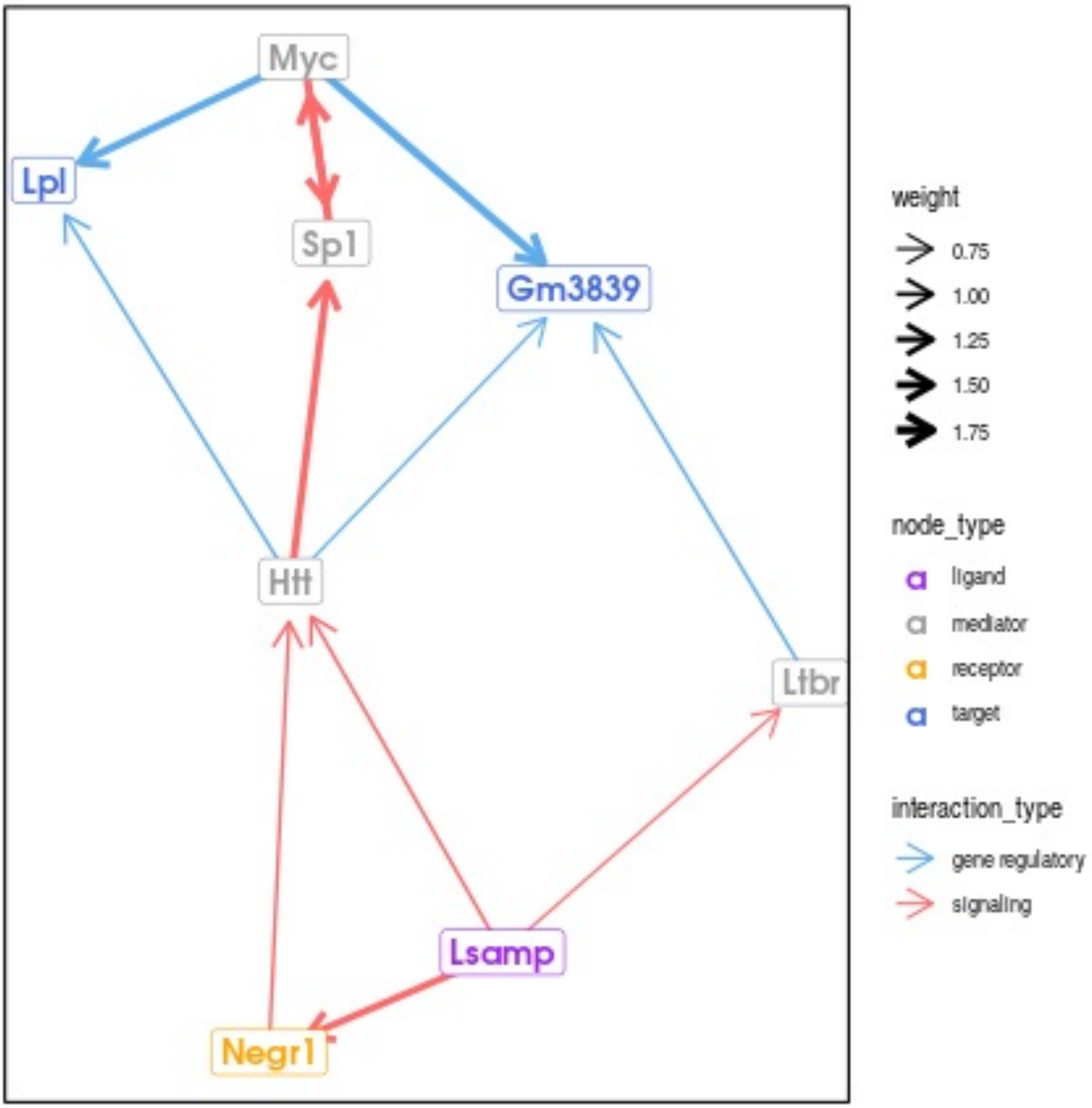
Myc
Lpl
Sp1
Gm3839
Htt
Ltbr
Lsamp
Negr1
weight
0.75
1.00
1.25
1.50
1.75
node_type
ligand
mediator
receptor
target
interaction_type
gene regulatory
signaling

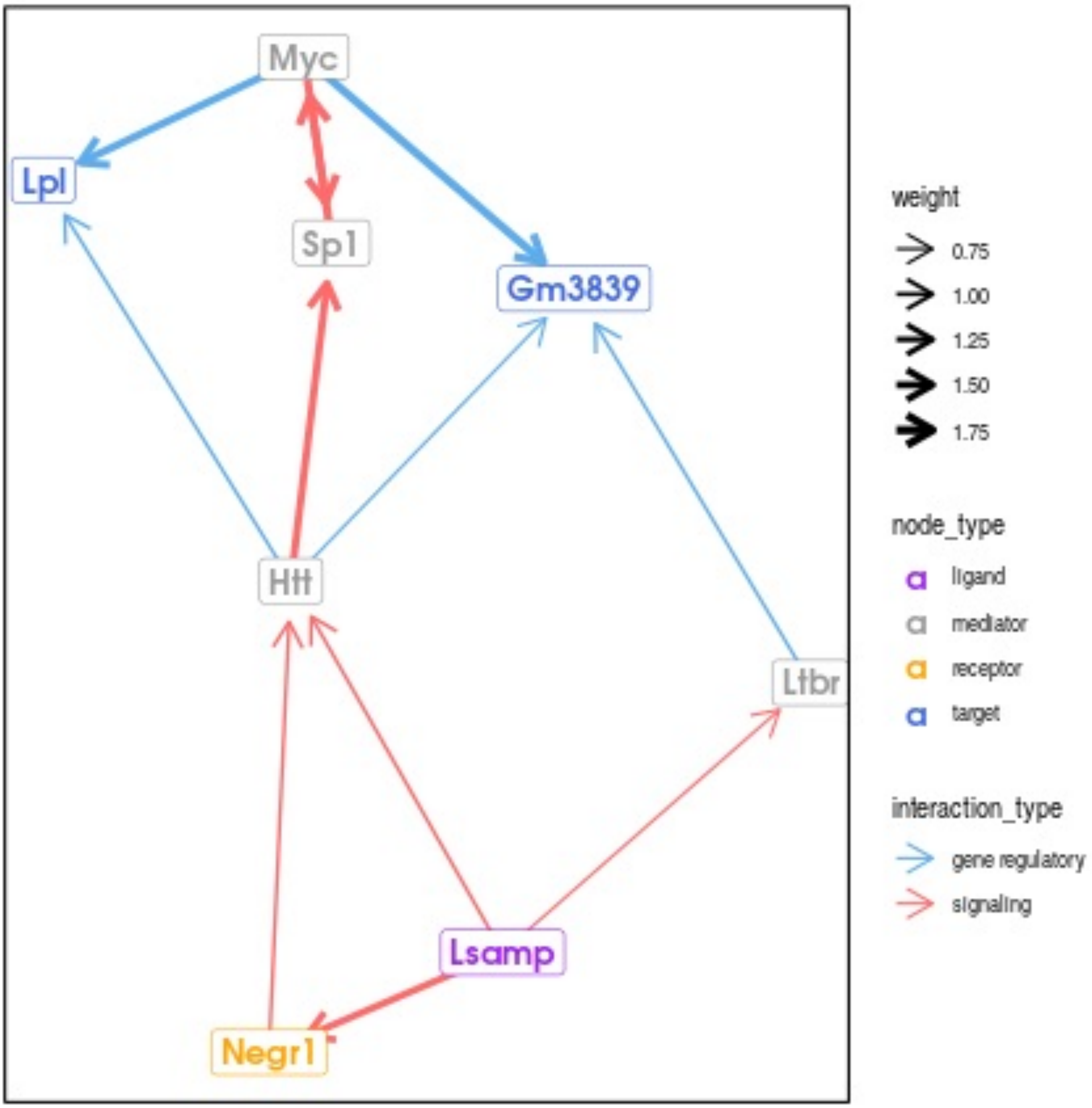
Myc
Lpl
Sp1
Gm3839
Htt
Ltbr
Lsamp
Negr1
weight
0.75
1.00
1.25
1.50
1.75
node_type
ligand
mediator
receptor
target
interaction_type
gene regulatory
signaling

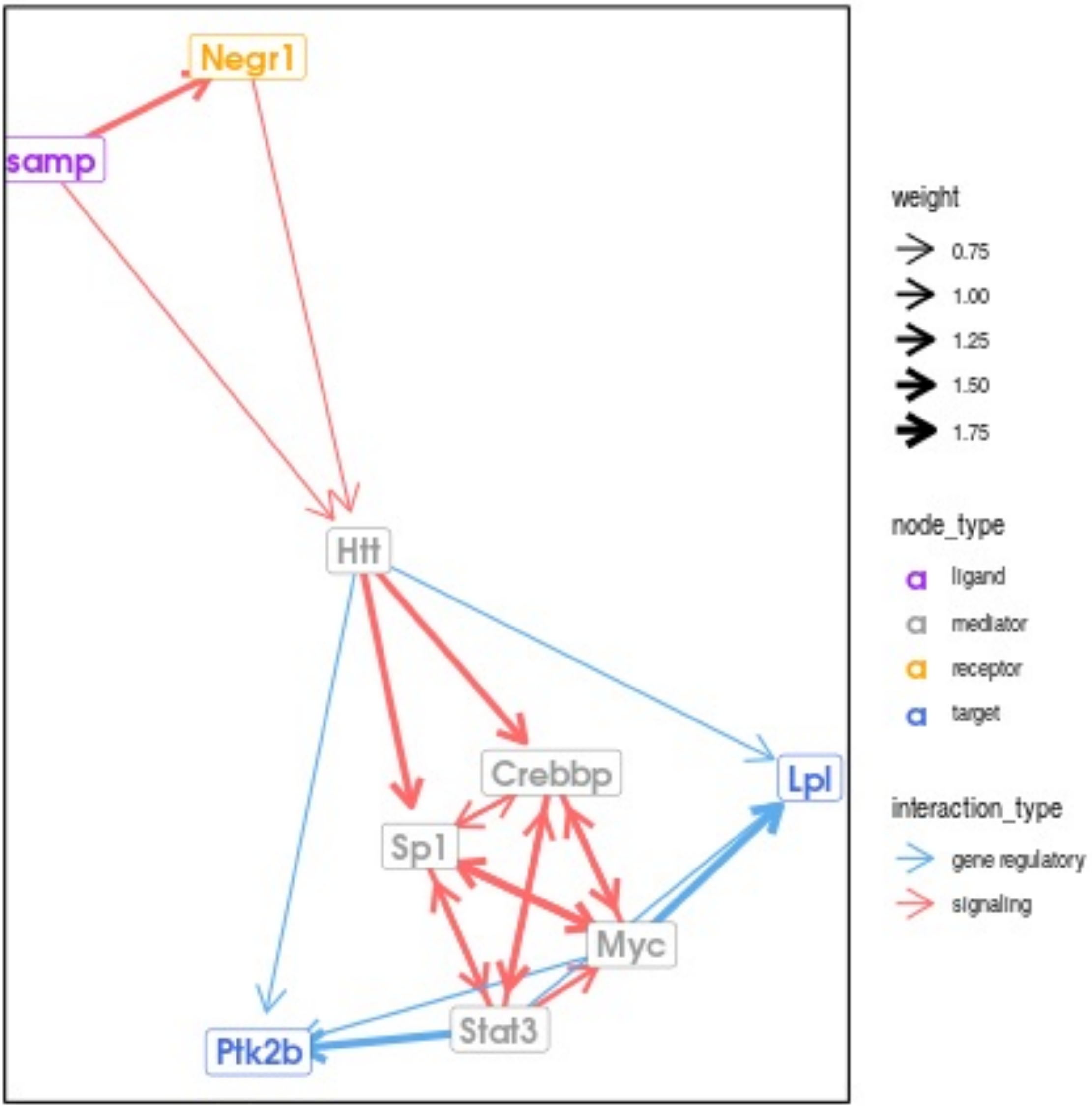

Negr1
samp
Htt
Crebbp
Sp1
Lpl
Myc
Stat3
Ptk2b
weight
0.75
1.00
1.25
1.50
1.75
node_type
ligand
mediator
receptor
target
interaction_type
gene regulatory
signaling

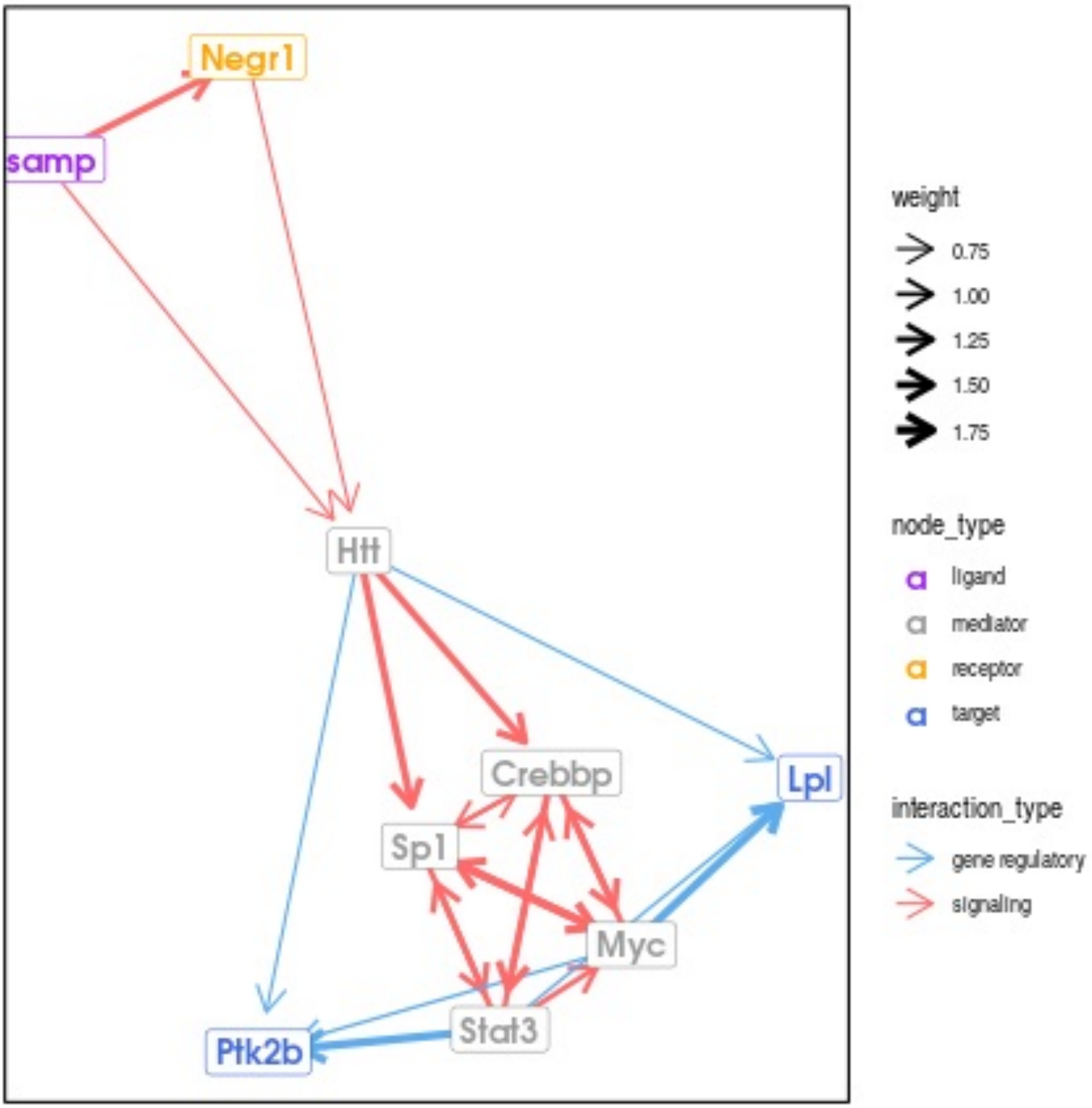

Negr1
samp
Htt
Crebbp
Lpl
Sp1
Myc
Stat3
Ptk2b
weight
0.75
1.00
1.25
1.50
1.75
node_type
ligand
mediator
receptor
target
interaction_type
gene regulatory
signaling

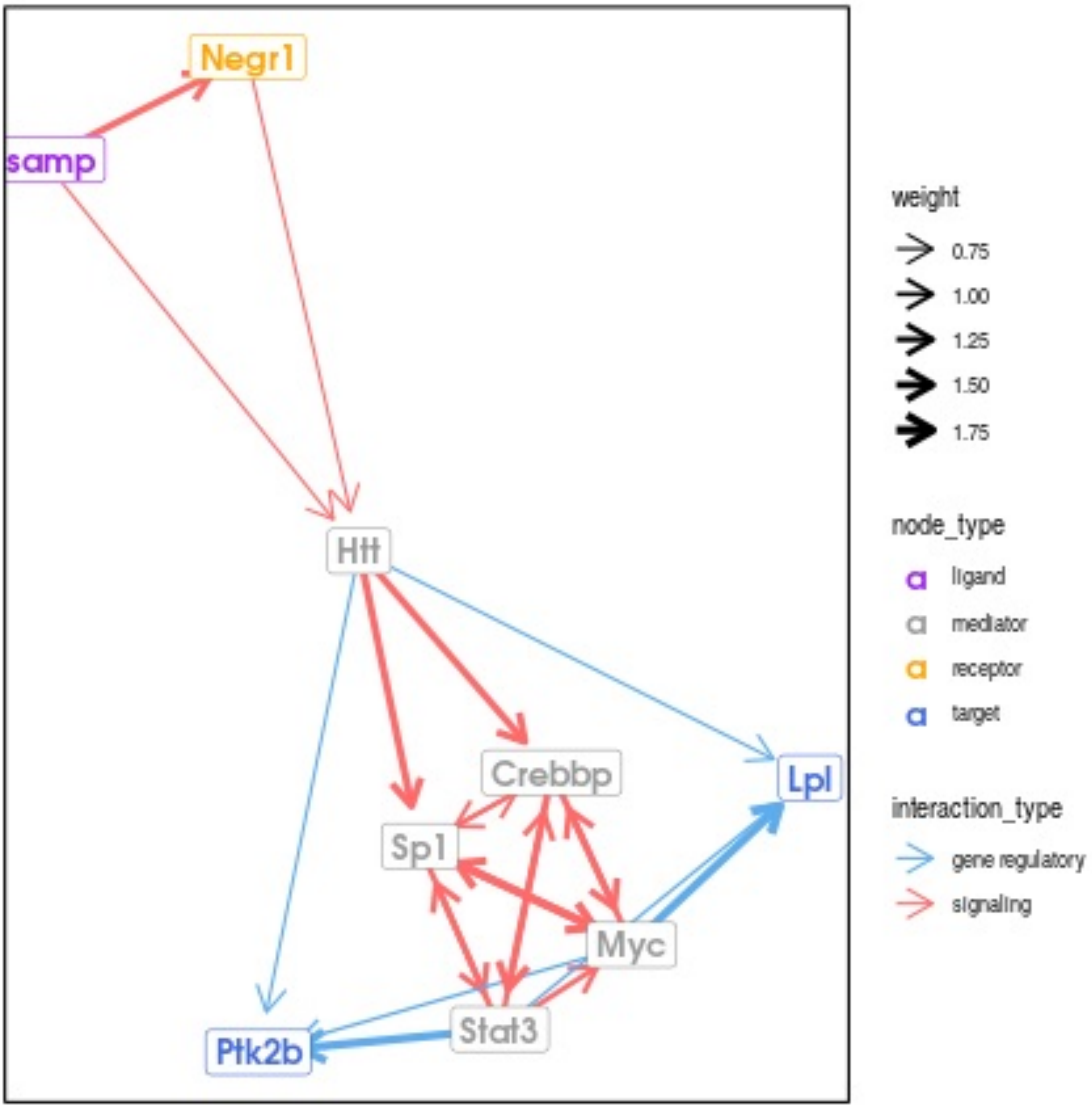

Negr1
samp
Htt
Crebbp
Lpl
Sp1
Myc
Stat3
Ptk2b
weight
0.75
1.00
1.25
1.50
1.75
node_type
ligand
mediator
receptor
target
interaction_type
gene regulatory
signaling

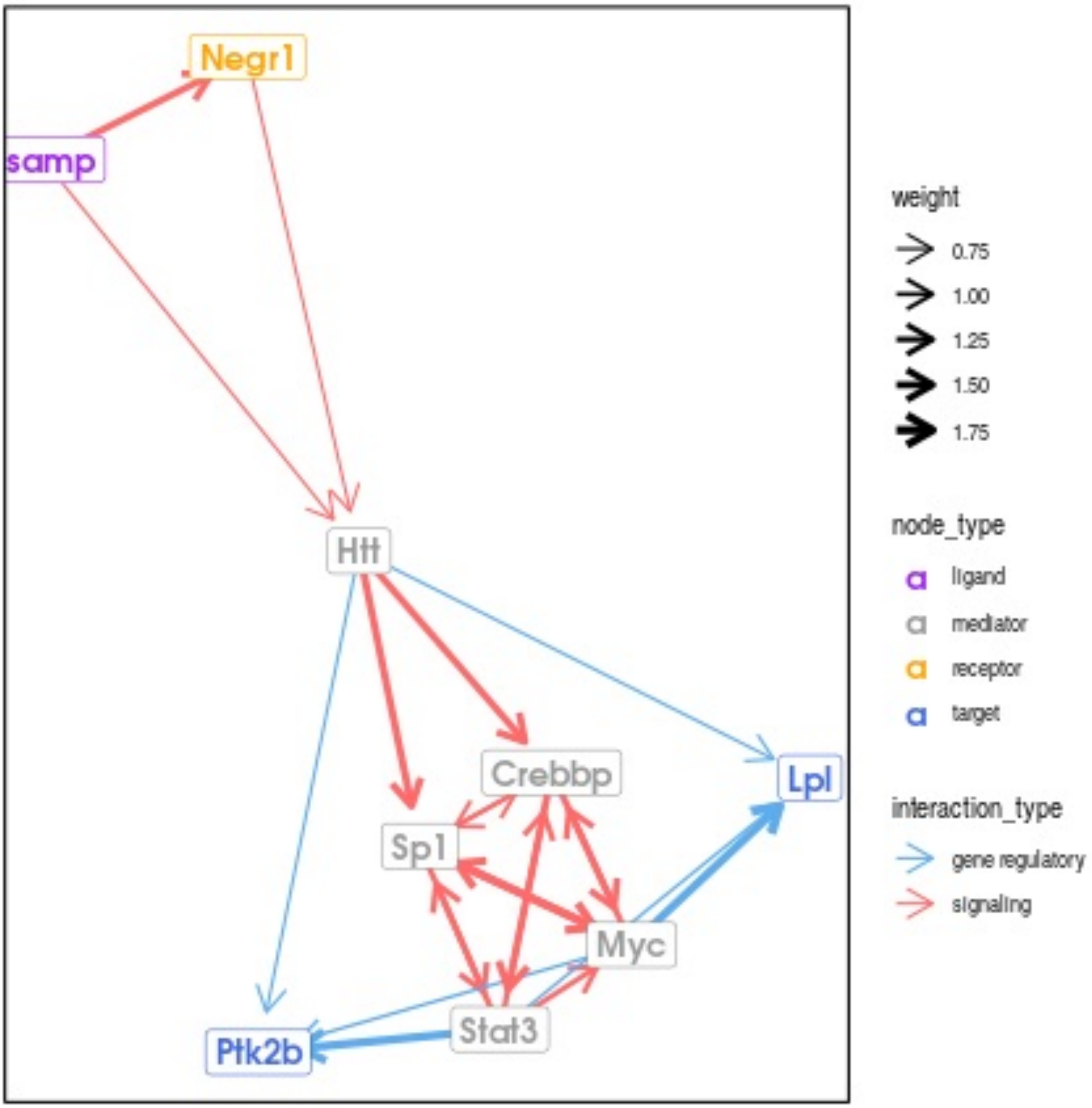

Negr1
samp
Htt
Crebbp
Lpl
Sp1
Myc
Stat3
Ptk2b
weight
0.75
1.00
1.25
1.50
1.75
node_type
ligand
mediator
receptor
target
interaction_type
gene regulatory
signaling

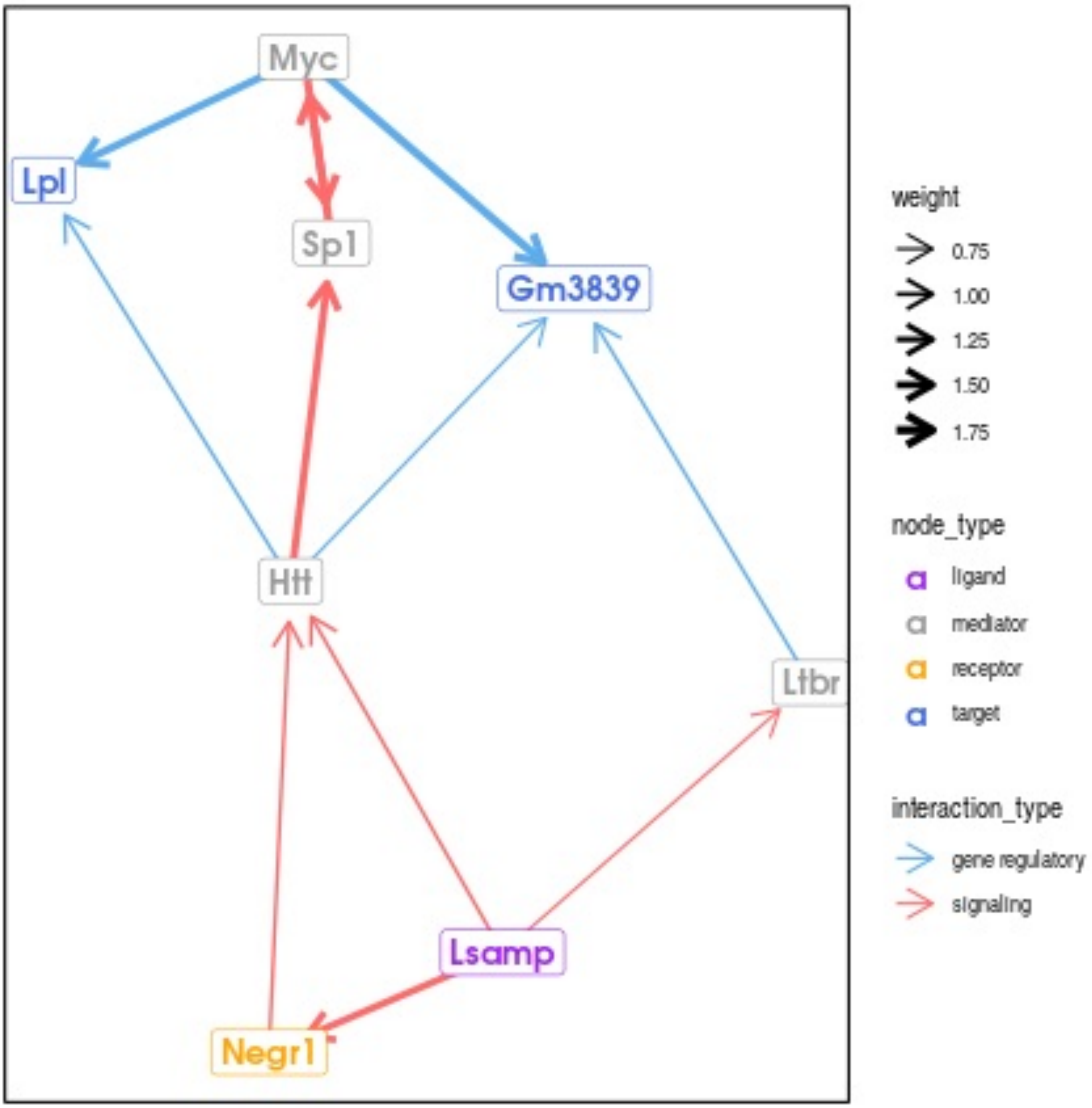

Myc
Lpl
Sp1
Gm3839
Htt
Ltbr
Lsamp
Negr1
weight
0.75
1.00
1.25
1.50
1.75
node_type
ligand
mediator
receptor
target
interaction_type
gene regulatory
signaling

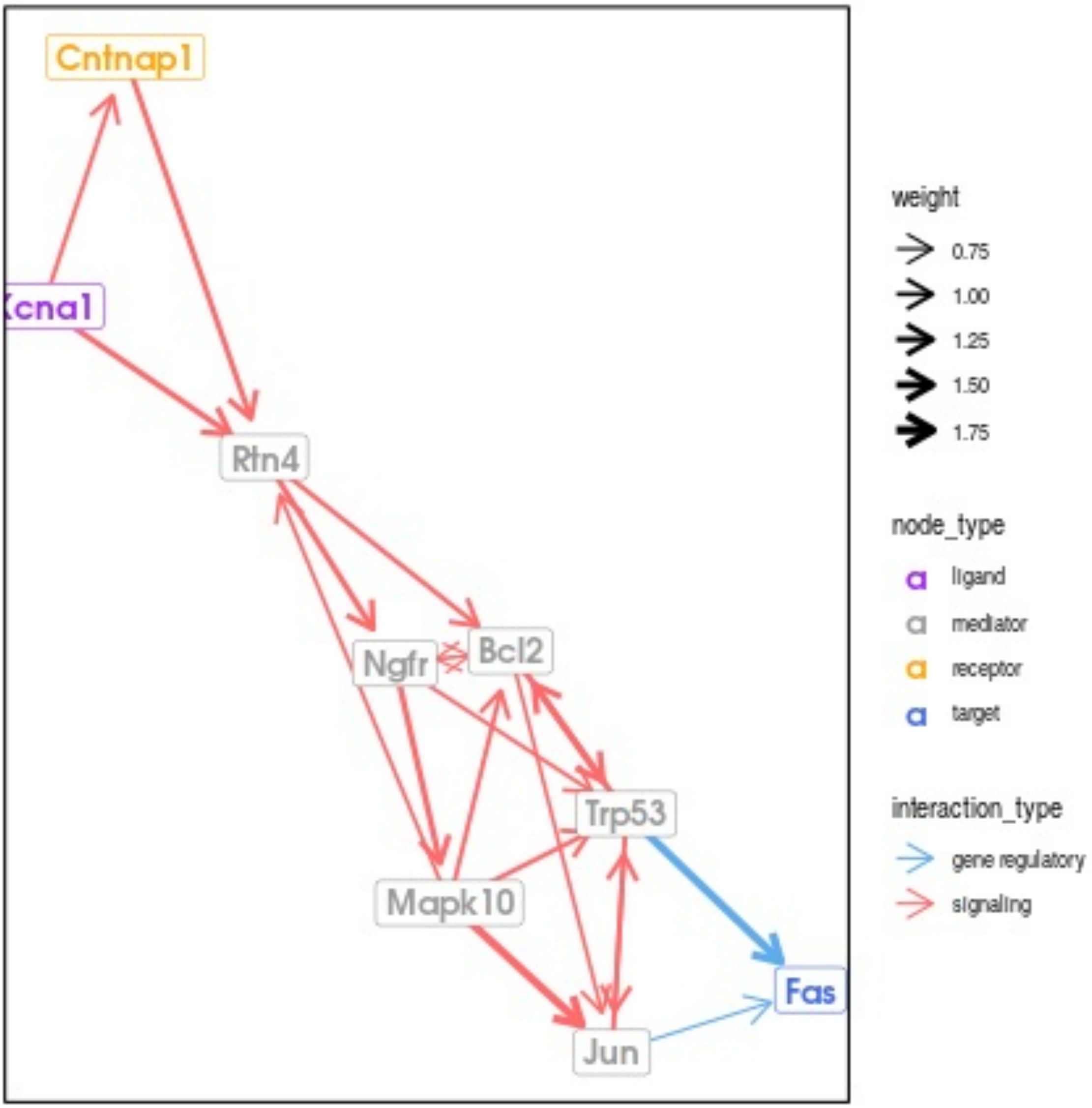
Cntnap1
Xcna1
Rtn4
Ngfr
Bcl2
Trp53
Mapk10
Jun
Fas
weight
0.75
1.00
1.25
1.50
1.75
node_type
ligand
mediator
receptor
target
interaction_type
gene regulatory
signaling

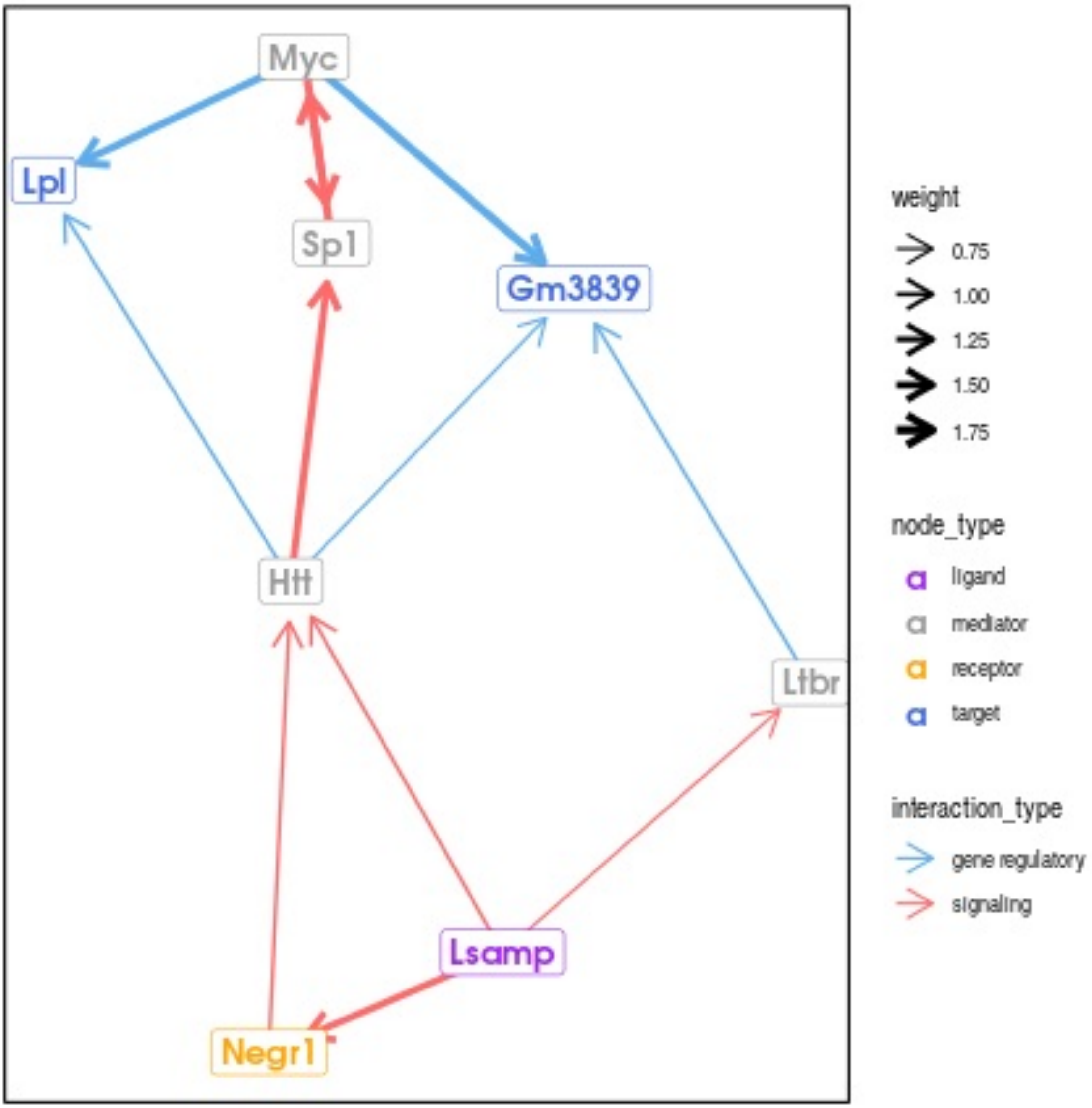
Myc
Lpl
Sp1
Gm3839
Htt
Ltbr
Lsamp
Negr1
weight
0.75
1.00
1.25
1.50
1.75
node_type
ligand
mediator
receptor
target
interaction_type
gene regulatory
signaling

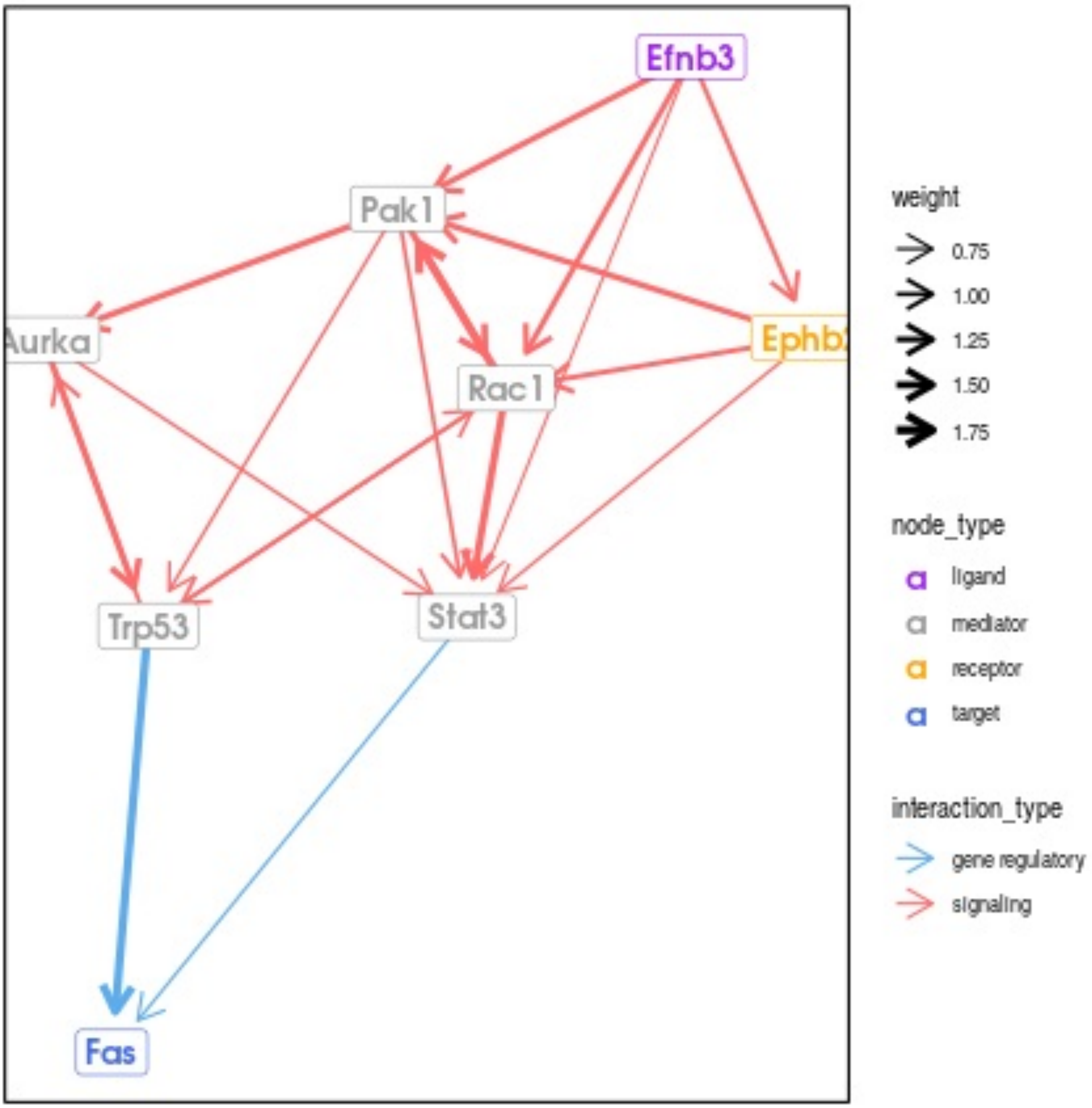
Efnb3
Pak1
Aurka
Ephb
Rac1
Trp53
Stat3
Fas
weight
0.75
1.00
1.25
1.50
1.75
node_type
ligand
mediator
receptor
target
interaction_type
gene regulatory
signaling

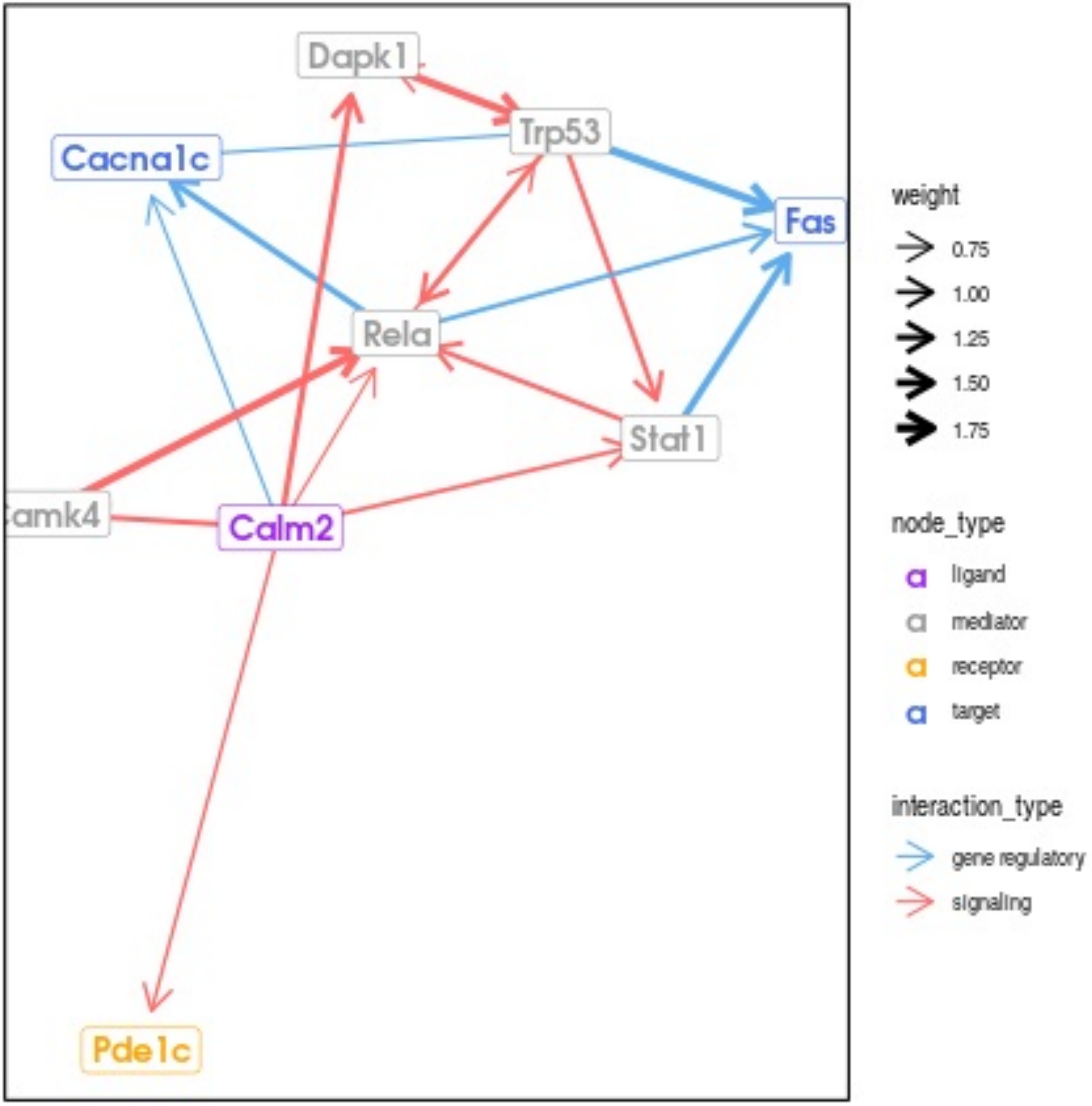
Dapk1
Trp53
Cacna1c
Fas
Rela
Stat1
Camk4
Calm2
Pde1c
weight
0.75
1.00
1.25
1.50
1.75
node_type
ligand
mediator
receptor
target
interaction_type
gene regulatory
signaling

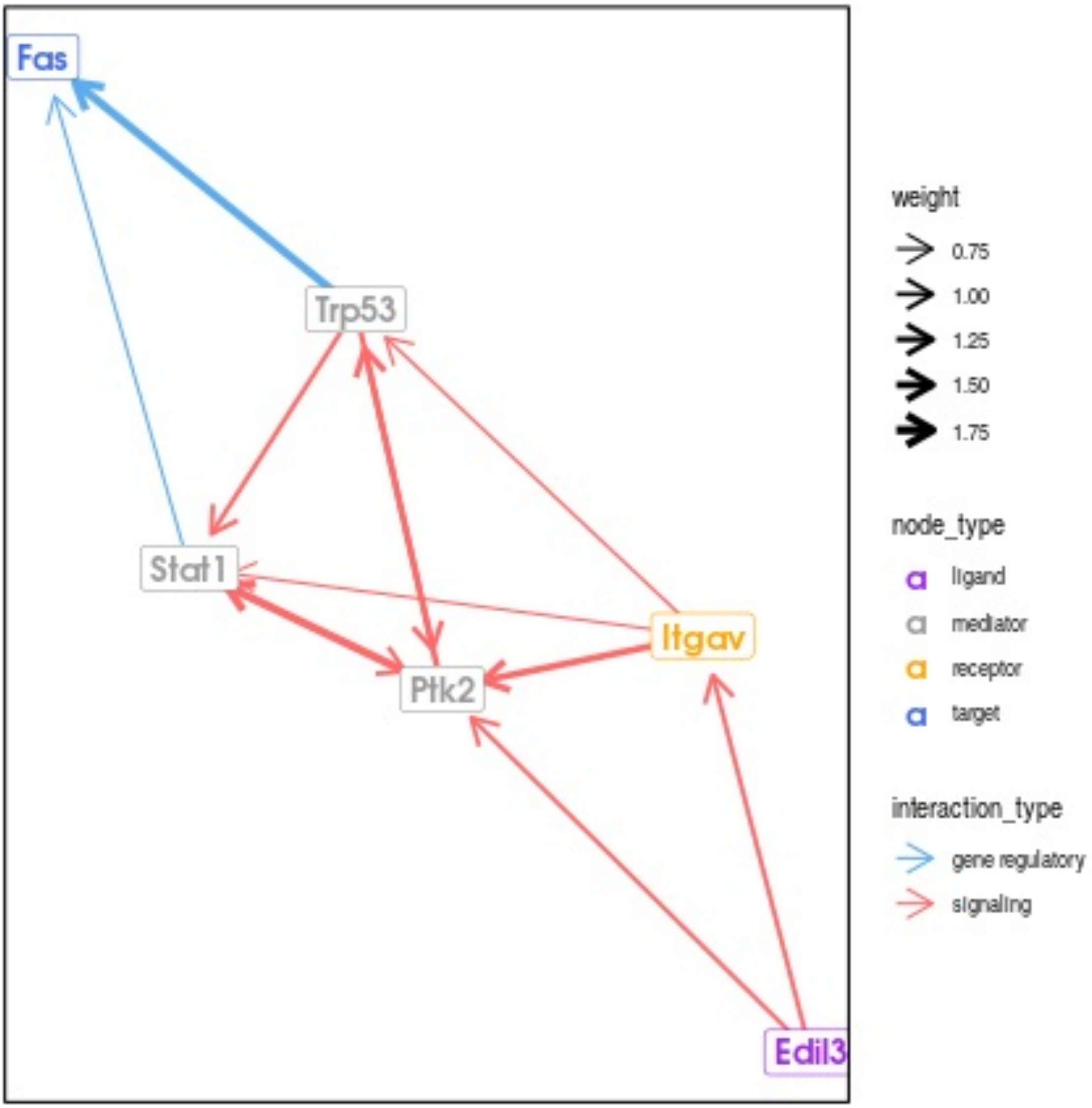
Fas
Trp53
Stat1
Itgav
Ptk2
Edil3
weight
0.75
1.00
1.25
1.50
1.75
node_type
ligand
mediator
receptor
target
interaction_type
gene regulatory
signaling

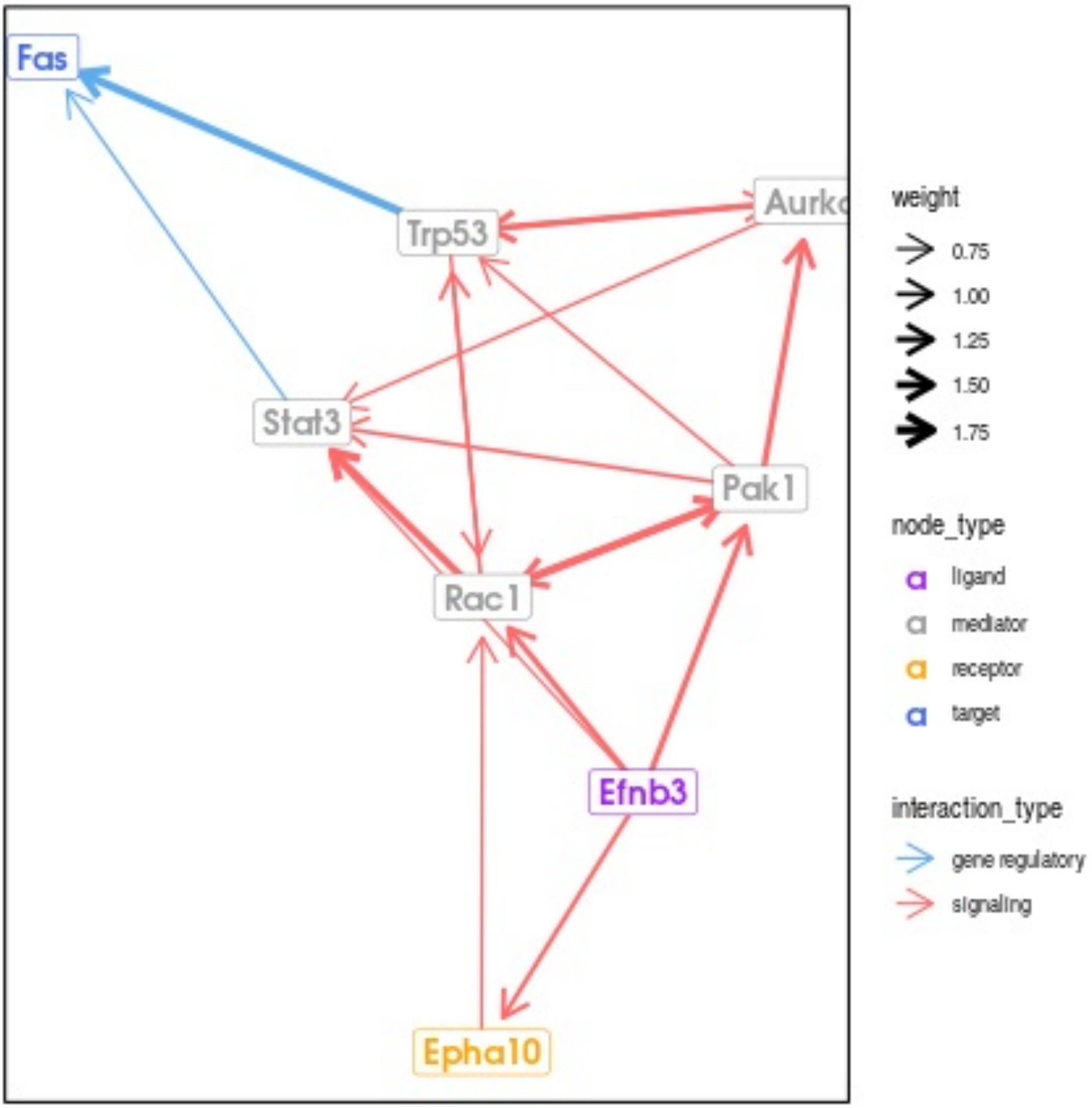

Fas
Trp53
Aurka
Stat3
Pak1
Rac1
Efnb3
Epha10
weight
0.75
1.00
1.25
1.50
1.75
node_type
ligand
mediator
receptor
target
interaction_type
gene regulatory
signaling

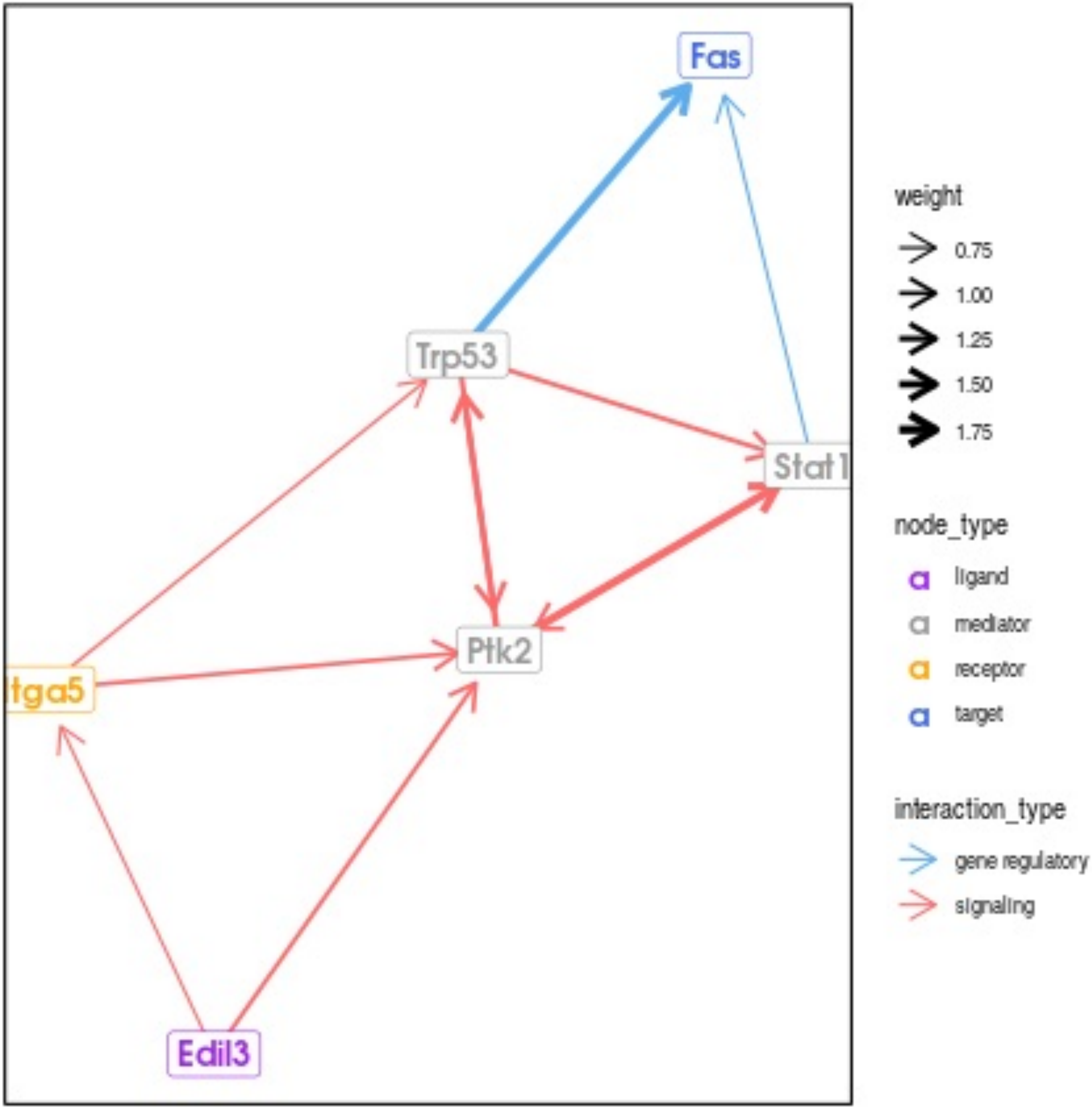

Fas
Trp53
Stat1
Ptk2
Itga5
Edil3
weight
0.75
1.00
1.25
1.50
1.75
node_type
ligand
mediator
receptor
target
interaction_type
gene regulatory
signaling

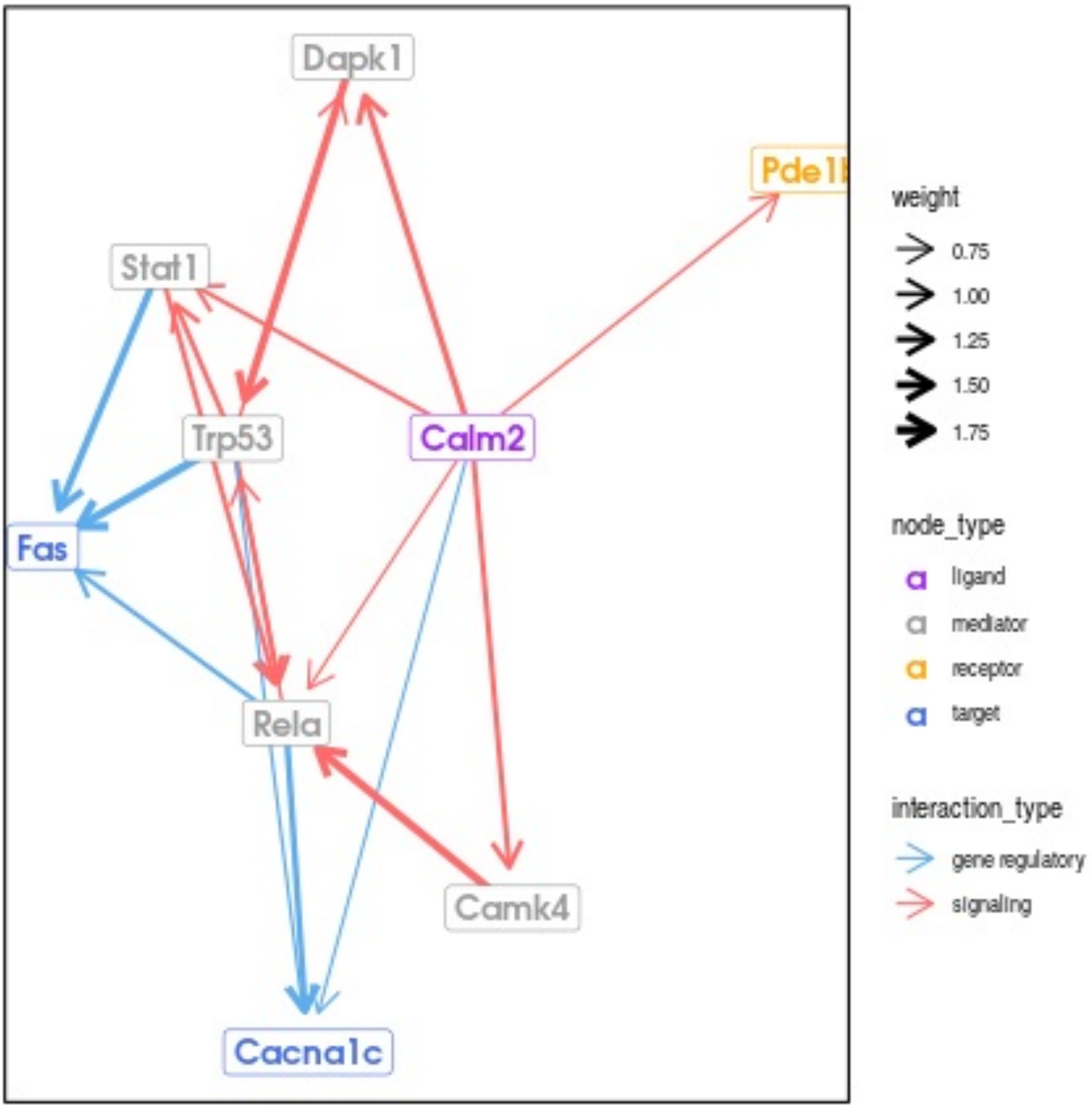

Dapk1
Pde1b
Stat1
Trp53
Calm2
Fas
Rela
Camk4
Cacna1c
weight
0.75
1.00
1.25
1.50
1.75
node_type
ligand
mediator
receptor
target
interaction_type
gene regulatory
signaling

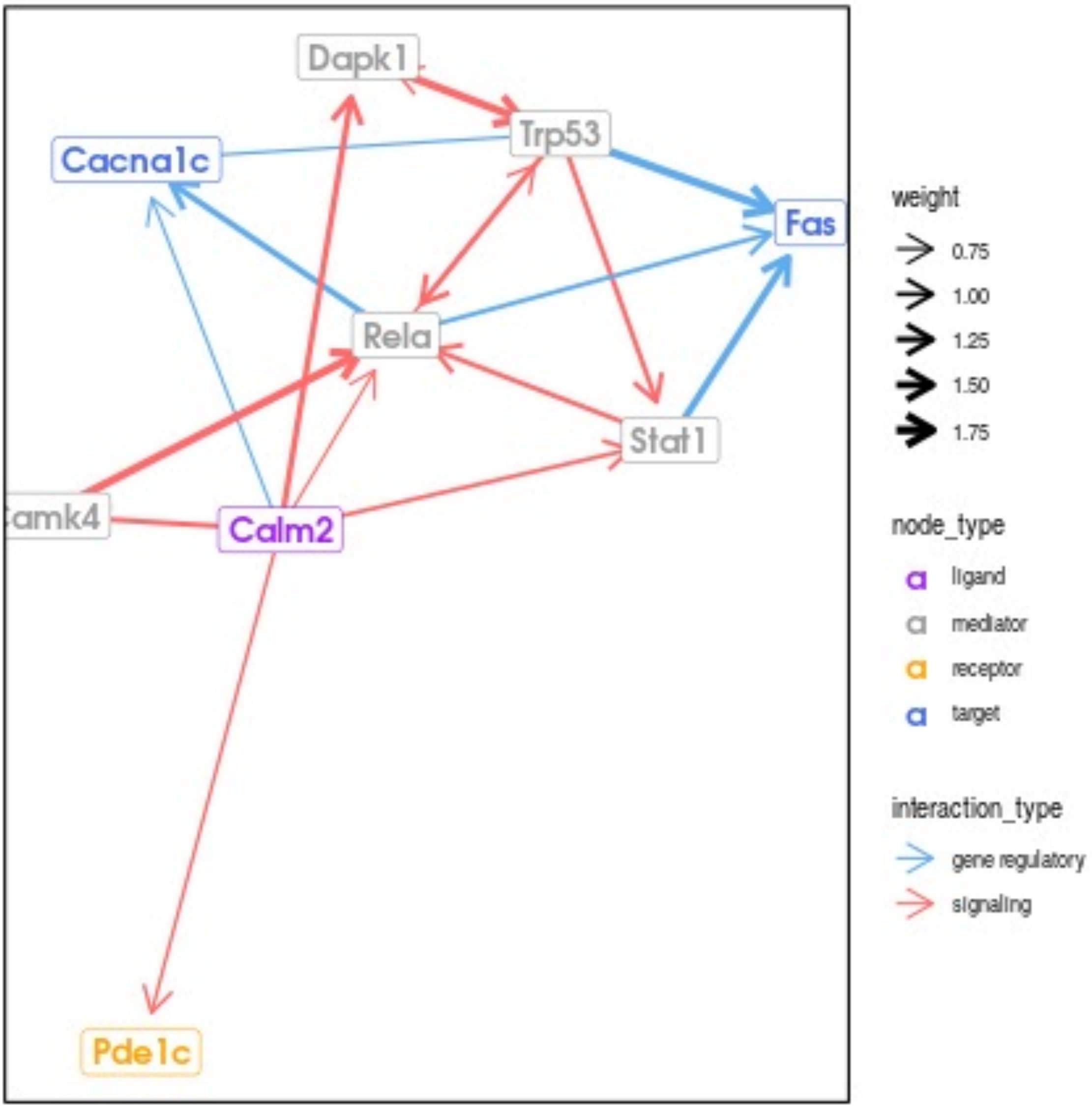

Dapk1
Trp53
Cacna1c
Fas
Rela
Stat1
amk4
Calm2
Pde1c
weight
0.75
1.00
1.25
1.50
1.75
node_type
ligand
mediator
receptor
target
interaction_type
gene regulatory
signaling

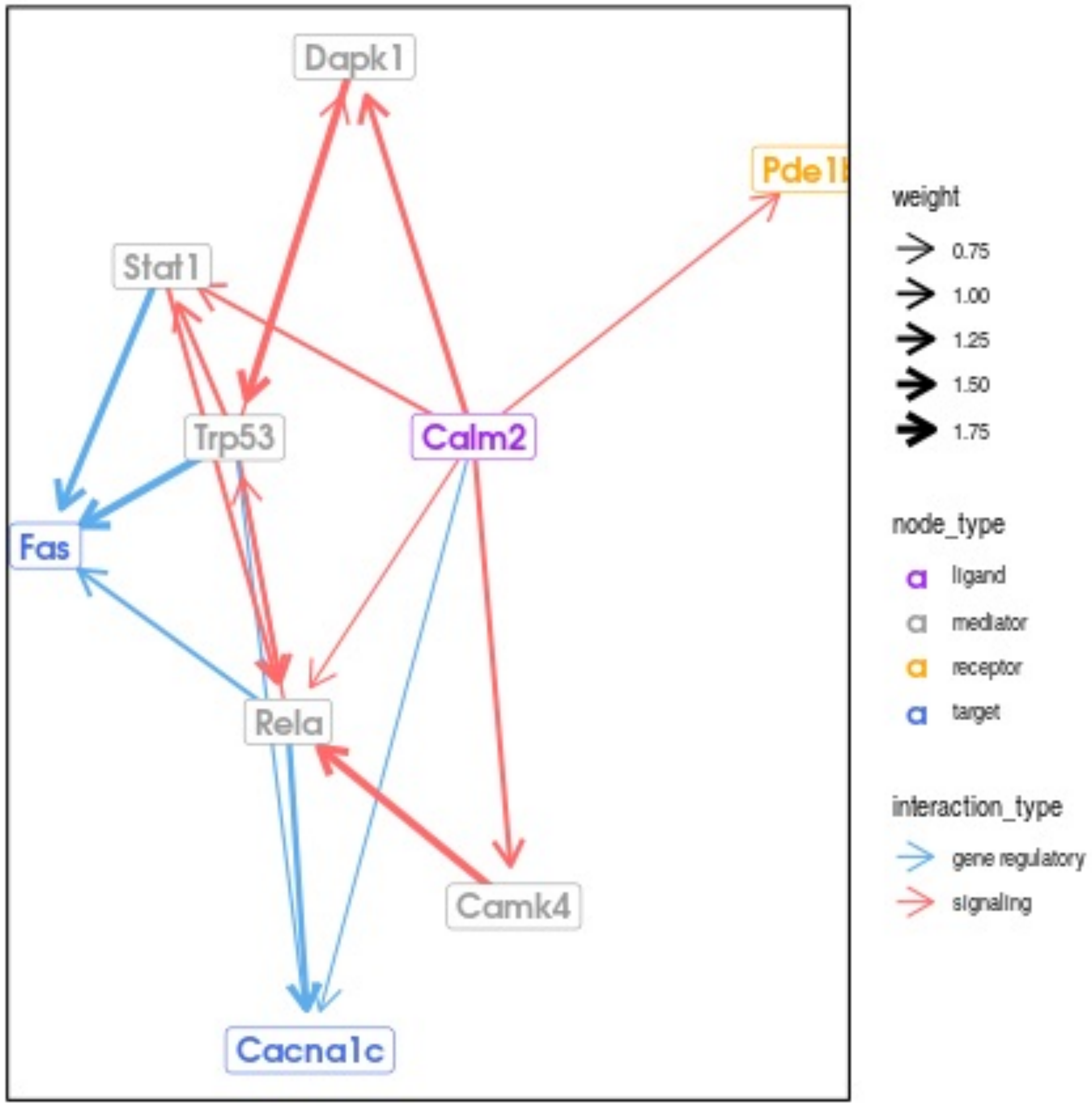
Dapk1
Pde1b
Stat1
Trp53
Calm2
Fas
Rela
Camk4
Cacna1c
weight
0.75
1.00
1.25
1.50
1.75
node_type
ligand
mediator
receptor
target
interaction_type
gene regulatory
signaling

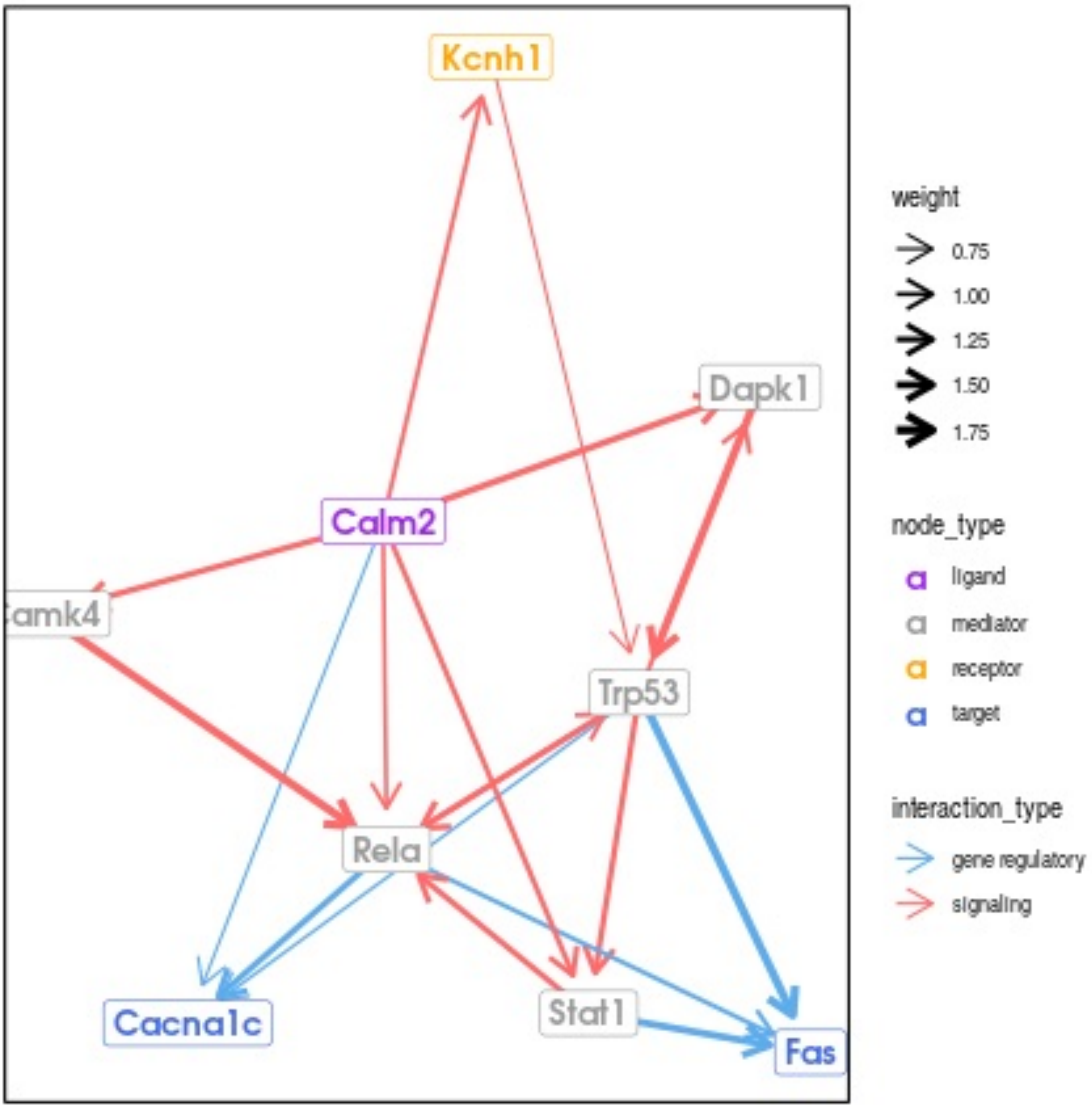
Kcnh1
Dapk1
Calm2
amk4
Trp53
Rela
Cacna1c
Stat1
Fas
weight
0.75
1.00
1.25
1.50
1.75
node_type
ligand
mediator
receptor
target
interaction_type
gene regulatory
signaling

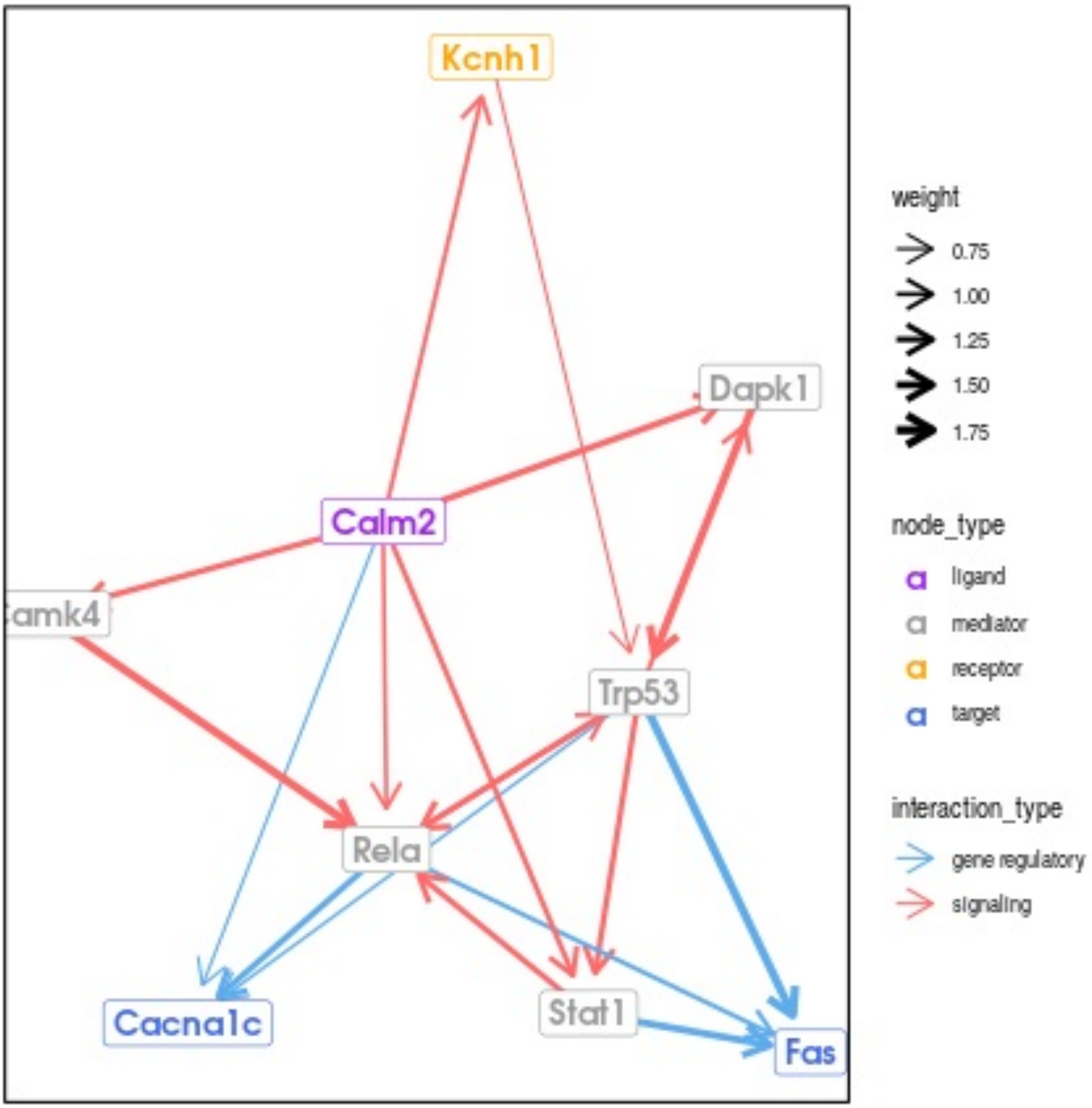
Kcnh1
Dapk1
Calm2
amk4
Trp53
Rela
Cacna1c
Stat1
Fas
weight
0.75
1.00
1.25
1.50
1.75
node_type
ligand
mediator
receptor
target
interaction_type
gene regulatory
signaling

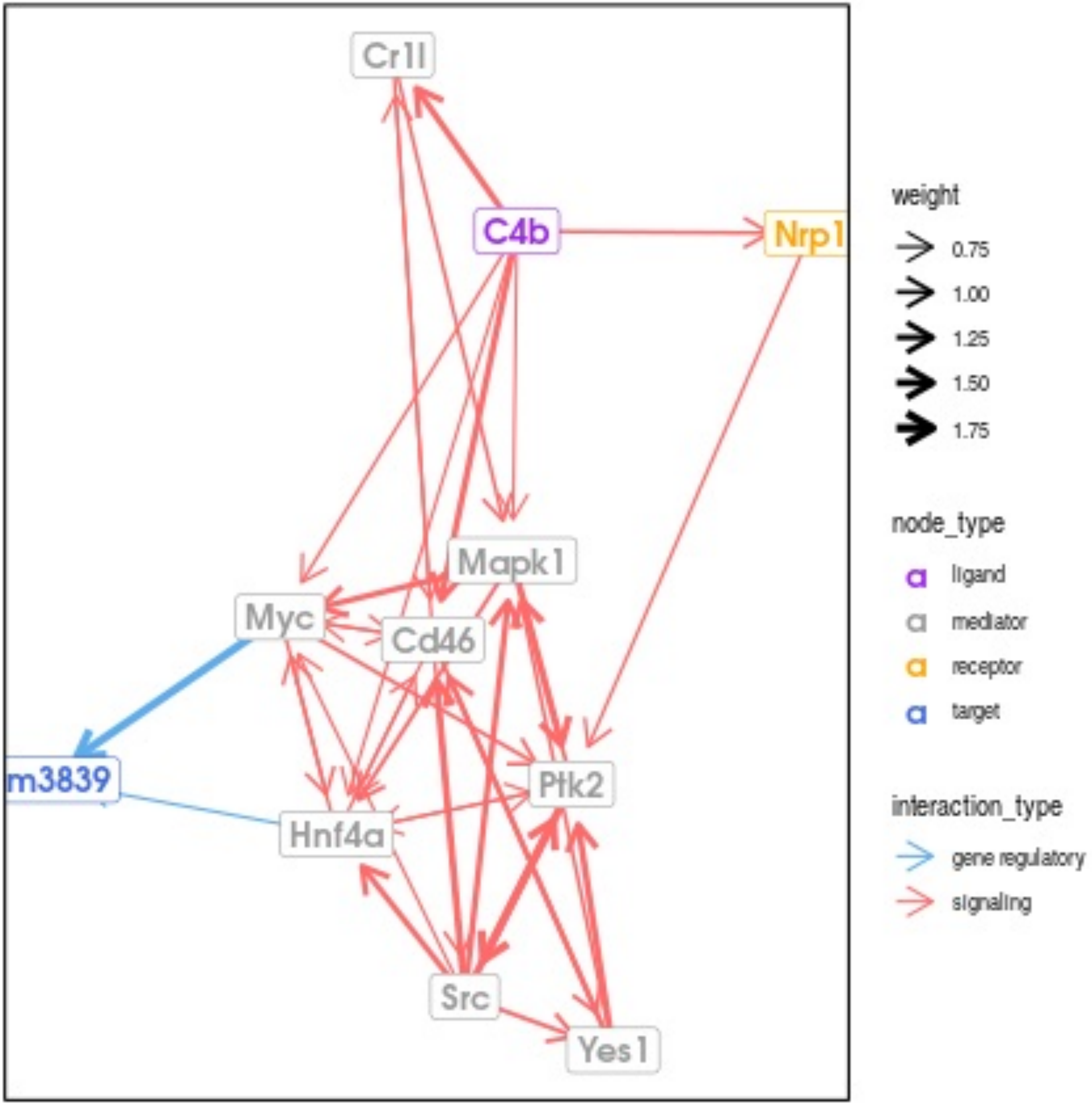
Cr1l
C4b
Nrp1
Mapk1
Myc
Cd46
m3839
Ptk2
Hnf4a
Src
Yes1
weight
0.75
1.00
1.25
1.50
1.75
node_type
ligand
mediator
receptor
target
interaction_type
gene regulatory
signaling
